# High-Throughput Screening and Initial SAR Studies Identify a Novel Sub-micromolar Potent Human cGAS Inhibitor

**DOI:** 10.1101/2025.08.18.670979

**Authors:** Jazmin Alarcón-Espósito, Ravi Kumar Nagiri, Siyu Wang, Chloe Larson, Loreto Carvallo-Torres, Vipin Kumar Singh, Fraser Glickman, Li Gan, Subhash C. Sinha

**Affiliations:** Helen and Robert Appel Alzheimer’s Disease Research Institute, Feil Family Brain and Mind Research Institute, Weill Cornell Medicine, New York, NY, USA; Fisher Drug Discovery Resource Center, Rockefeller University, New York, NY

**Keywords:** Cyclic GMP-AMP synthase (cGAS), cGAS-STING pathway, High throughput screening (HTS), cGAS inhibitors, Alzheimer’s disease (AD)

## Abstract

Cyclic GMP-AMP synthase (cGAS) has emerged as a promising therapeutic target of several human diseases, including Alzheimer’s disease (AD) and other neurodegenerative disorders. As a cytosolic DNA sensor, cGAS generates an innate immune response to promote neuroinflammation by producing an endogenous agonist of the stimulator of interferon genes (STING), 2’3’-cyclic GMP-AMP (cGAMP), which activates the cGAS–STING pathway. We have performed a high-throughput screening of a chemical library containing over 300K small molecules at the Fisher Drug Discovery Resource Center (DDRC), Rockefeller University (RU), to identify multiple hit inhibitors of human (h)-cGAS. We used a modified Kinase Glo® Luminescent Kinase assay, which was earlier developed at RU and later used by multiple groups, including ours, to perform primary screening of the library using h-cGAS. The hit candidates bearing novel scaffolds are structurally diverse and exhibited in vitro activity in the low micromolar range. **RU-0610270** or compound (cpd) **1**, a sulfonamide derivative, is one of the most potent hits (IC_50_=1.88 µM), selected for hit expansion and structure-activity relationship (SAR) analysis. We synthesized new analogs of **cpd 1** and evaluated them in vitro against h-cGAS to identify **cpd 6** (IC_50_=0.66 µM) as the most potent hit analog. We further profiled **cpd 6** and found that it modestly inhibited cGAMP levels by 29% at 30 µM in THP1 cells without detectable toxicity, and by 76% at 100 µM, albeit with a moderate decrease (∼20%) in cell viability. These results highlight a novel chemical series with promising in vitro activity, providing a starting point for the development of selective and potent human cGAS inhibitors for clinical use.

## Introduction

Cyclic GMP-AMP synthase (cGAS), a cytosolic double stranded DNA (ds-DNA) sensor enzyme, has emerged as a promising therapeutic target for numerous human diseases, including autoimmune diseases, Alzheimer’s disease (AD), and other neurodegenerative diseases (NDs).^1-7^ Upon its stabilization by ds-DNA, the 2:2 cGAS:ds-DNA dimer catalyzes the condensation of ATP with GTP to produce 2’3’-cyclic GMP-AMP, cGAMP, which is a low nanomolar potent agonist of stimulator of interferon genes (STING) (Figure 1).^8^ This activates STING and the STING-dependent neuroinflammatory responses characterized by elevated TBK1 phosphorylation, IRF3 activation, and type I interferons (IFN-I).^2, 5^ While activation of cGAS-STING pathway through STING agonists can have positive antiviral effects and may sensitize tumors to immunotherapy, chronic activation of the pathway leads to sustained expression of interferon-inducible genes, which perpetuates inflammation and tissue damage.^1^ In AD pathology, amyloid-β (Aβ) plaques and hyperphosphorylated Tau (pTau) induce oxidative mitochondrial stress and damage, causing the release of ds-mitochondrial DNA (mtDNA) into the cytoplasm and activating STING similarly, as described above.^5, 6^ Chronic activation of this cGAS-STING-IFN pathway drives microglial dysfunction, promotes neurotoxic astrocyte activation, and accelerates synaptic loss and cognitive decline.^1, 4, 7^

**Figure 1.**
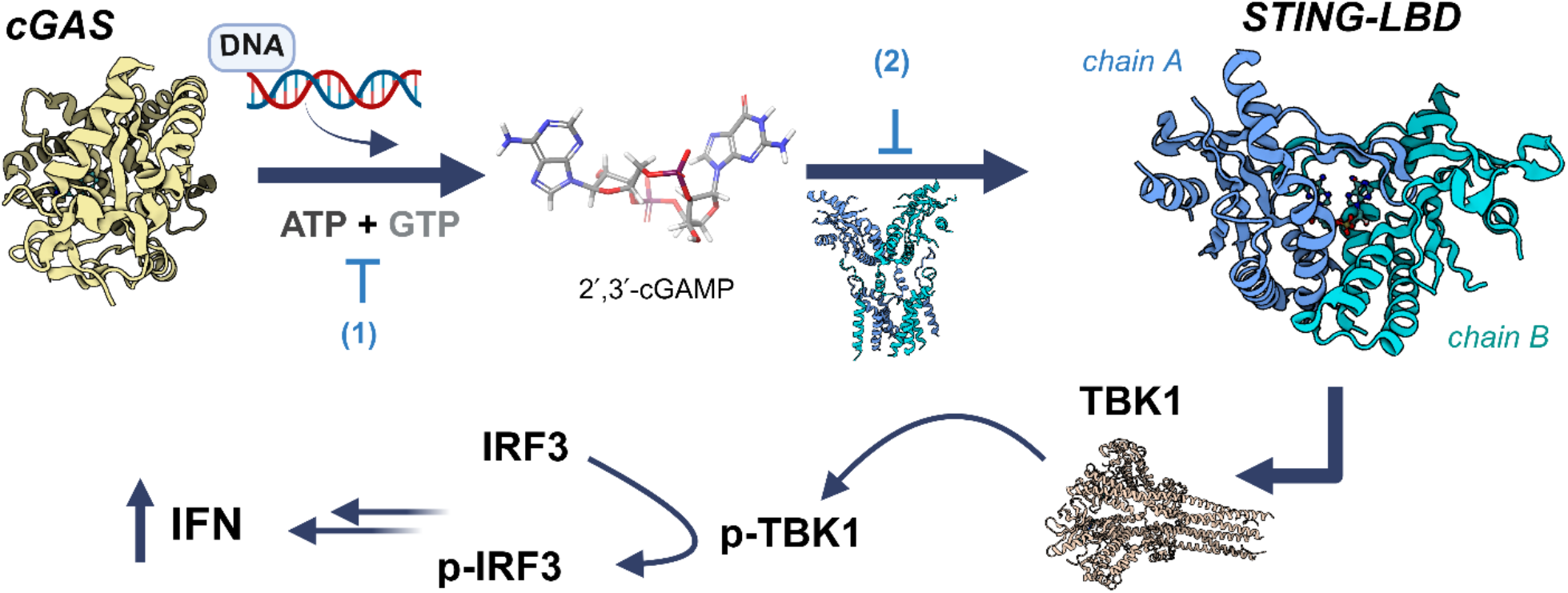
cGAS activation regulates innate immunity and the cGAS-STING-TBK1 pathway and the downstream signaling, which could be blocked using cGAS (1), or STING (2) inhibitors (LBD: ligand binding domain).

Recent studies by Udeochu *et al*.^*4*^, working with both postmortem human tissues from AD patients and transgenic (Tg) mouse models of tauopathy, including those partially or fully devoid of the cGAS gene or those carrying additional risk or protective genes, revealed numerous facts. First, it was demonstrated that cGAS is activated in both human AD brains and Tau P301S Tg (PS19) mouse brains. Second, both partial and full genetic ablation of cGAS and pharmacological inhibition using small-molecule inhibitors of cGAS, e.g., **RU.521** or **TDI-6570** for mouse (m)-cGAS) reduces Aβ pathology, suppresses neuroinflammation, and rescues memory deficits in PS19 mice.^4-6^ Similarly, STING inhibition using **H-151** reduces Aβ pathology, besides suppressing neuroinflammation and rescuing memory deficits in 5xFAD mice.^2, 3^ Third, Gan lab has shown that the risk factors–such as the APOE ε4 allele,^9^ the TREM2 R47H variant,^10-12^ and female sex promote microglial cGAS activation, leading to cellular senescence and neurodegeneration in tauopathy models, while the resilience allele, such as Christchurch (CC) APOE3/ R136S^13^ variant provides resilience against tau pathology by suppressing cGAS–STING–IFN signaling.

The above-described information and the promise that the pharmacological inhibition of cGAS could treat AD and other NDs prompted us to discover a novel h-cGAS inhibitor (cGASI). Toward this goal, we optimized the previously described cell-free cGAS activity assay and performed high-throughput screening (HTS) of a chemical library containing 300K small molecules at the Rockefeller University (RU), followed by hit validation and optimization of one of the identified potent h-cGASI hits. Here, we report that the optimized screening method conveniently screened the 300K small molecule library, identifying 38 hit inhibitors of h-cGAS in a relatively short period vis-a-vis the previous HTS campaign.^14^ Three distinct structural families (F), e.g., *F1* (bicyclic cpds), *F2* (sulfonamides), and *F3* (thiadiazole cpds), showed low micromolar IC_50_ (0.52 to 3.35 μM) in a cell-free assay. One of the sulfonamide derivatives, **cpd 6**, obtained by medicinal chemistry, was the most potent hit, **RU-0610270** (IC_50_ = 1.88 μM), which showed a slight improvement in its activity (IC_50_ = 1.38 μM) against h-cGASI, and inhibited cGAMP production modestly in THP1 monocytes.

## Results and discussion

### Assay development for cGAS high-throughput screening

The cGAS enzyme catalyzes the reaction of GTP and ATP to produce cGAMP and two equivalents of pyrophosphate (Scheme 1), providing at least three different methods to assess the activity of a cGAS inhibitor under cell-free conditions.^6^ These can be characterized as: *i)* ATP-depletion assay, *ii)* cGAMP production assay, and *iii)* pyrophosphate (PPi) production assay. In the ATP depletion assay, cGAS-catalysis is determined by measuring the amount of unreacted ATP in the reaction using a luminescence-based Kinase-Glo® kit.^15^ Measuring the ATP levels after approximately 50% conversion provides consistent results. On the other hand, in both the cGAMP and pyrophosphate production assays, cGAS catalysis is determined by measuring the levels of cGAMP and pyrophosphate, respectively. The cGAMP level can be determined using the Bell Brook cGAS/cGAMP Transcreener® kits, cGAMP ELISA, or the RapidFire-MS,^16^ and the pyrophosphate levels measured using inorganic pyrophosphatase coupled methods. All these methods have been previously used in HTS campaigns;^6^ we found that the ATP-depletion assay is simple and well-suited for large screens. Additionally, we used the cGAS/cGAMP Transcreener® for performing a confirmatory assay in cell-free conditions, and the cGAMP ELISA to determine cGAS activity in cell-based assays.^17, 18^

**Scheme 1.**
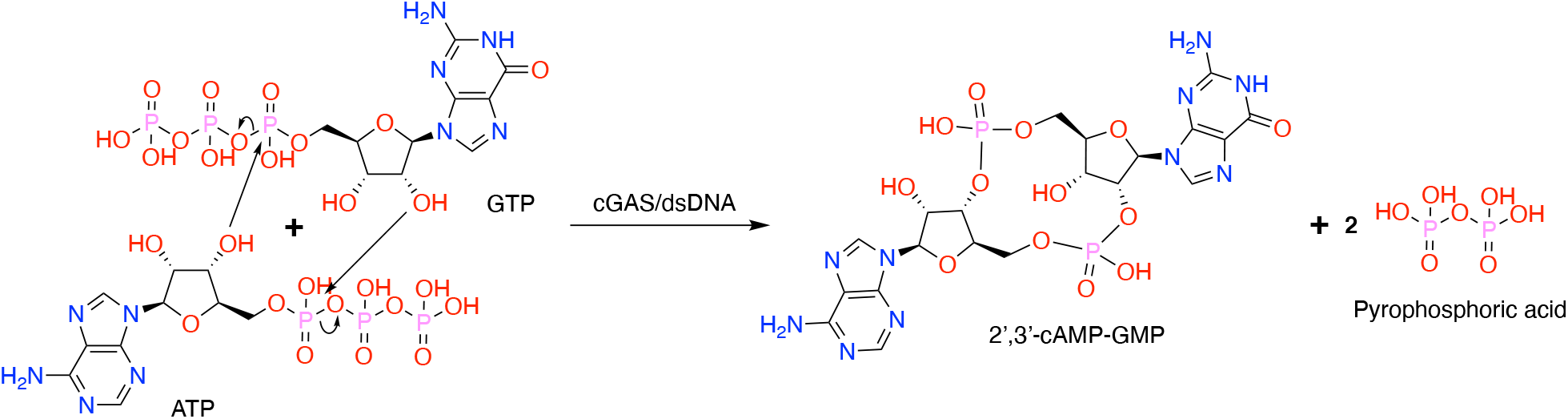
cGAS catalyzes the reaction of GTP and ATP to produce cGAMP. Key: dsDNA, double stranded DNA.

### Optimization of the ATP depletion assay for HTS

We used the previously described HTS method, developed by Lama *et al*.^*14*^, and optimized it further to increase the throughput and reduce the time required for screening. The key difference between the two methods, as described in Table 1, includes the concentration of NaCl that greatly affected cGAS activity, measured by determining unconsumed ATP concentration in the cGAS-catalyzed conversion of ATP+GTP→cGAMP using ‘Kinase-Glo® Max Luminescent kinase assay kit’ in 384-well plates. We evaluated three buffers: *B1* contained 5 mM MgCl_2_ and 1 µM ZnCl_2_ in Tris-HCl buffer with 150 mM NaCl, *B2* contained 5 mM MgCl_2_ in Tris-HCl buffer with 50 mM NaCl, and *B3* contained 2.5 mM MgCl_2_ and 0.1 mM MnCl_2_ in Tris-HCl buffer with 50 mM NaCl. *B2* has recently been used by Skeldon *et al*.^19^ in the cGAS assay. We examined *B3* containing MnCl_2_ because the latter has been shown to activate cGAS. It was also described that MnCl_2_ activated cGAS independently of ds-DNA.^20^ In addition to the modified buffer, we used HIS-tagged human cGAS (h-cGAS) instead of untagged cGAS and herring testes DNA (ht-DNA) to substitute for ds-DNA. These changes facilitate cGAS generation and purification, as ht-DNA is commercially available at a low cost; otherwise, these were not expected to make much difference.

**Table 1.** Comparison of buffer compositions for optimizing cGAS enzymatic assays.

| Reagents | Buffer 1<sup>14</sup> | Buffer 2 | Buffer 3 |
| --- | --- | --- | --- |
| Tris-HCl pH 7.4 | 20 mM | 20 mM | 20 mM |
| NaCl | 150 mM | 50 mM | 50 mM |
| MgCl<sub>2</sub> | 5 mM | 5 mM | 2.5 mM |
| ZnCl<sub>2</sub> | 1 μM | - | - |
| MnCl<sub>2</sub> | - | - | 0.1 mM |
| DTT | 1mM | 1mM | 1 mM |
| Tween20 | 0.01% | 0.01% | 0.01% |

We evaluated the activity of HIS-h-cGAS by incubating ATP and GTP (100 µM each) with the enzyme (100 nM) and ds-DNA (25 nM) or ht-DNA (5 µg/ml) in buffers *B1*-*B3* at room temperature for 3, 5 or 7 hours, keeping the ds-DNA (25 nM) and ht-DNA (5 µg/ml) concentration similar to that described in the original condition^14^ or from our study performed in 96-well plates. Identical reactions without cGAS or DNA were used as negative controls, and the progress of the reactions was measured using Kinase-Glo®, which determined the unconsumed ATP. The results, shown in Figure 2A (in the table) and 2B, confirmed that no ATP conversion was observed in the absence of cGAMP or DNA, as expected. Meanwhile, ATP levels decreased in all conditions over time. However, cGAS activity was significantly higher in *B2* compared to *B1*, and a 3-hour incubation was sufficient to achieve greater than 50% ATP conversion, making it already suitable for the HTS campaign. The cGAS activity increased further in buffer *B3* containing MnCl_2_ at low concentration (0.1 mM) likely because of a known synergistic effect between dsDNA and MnCl2 in cGAS activation and likely not independent of DNA as the latter require high MnCl_2_ concentration (2 mM).^20, 21^ Subsequently, evaluation of t h-cGASI, **G150**^14^ and **TDI-008246**^22^ showed that both possess low nanomolar IC_50_ values as expected. However, cGAS inhibition was found to be higher in *B1*, likely due to the low conversion. Based on these results, both *B2* and *B3* buffers together with a less costly and readily available ht-DNA instead of ds-DNA could be used for the HTS campaign. However, we used *B2* buffer to minimize changes in the already established HTS method that had already enabled the identification of numerous low nanomolar potent cGASI hits.^14^

**Figure 2.**
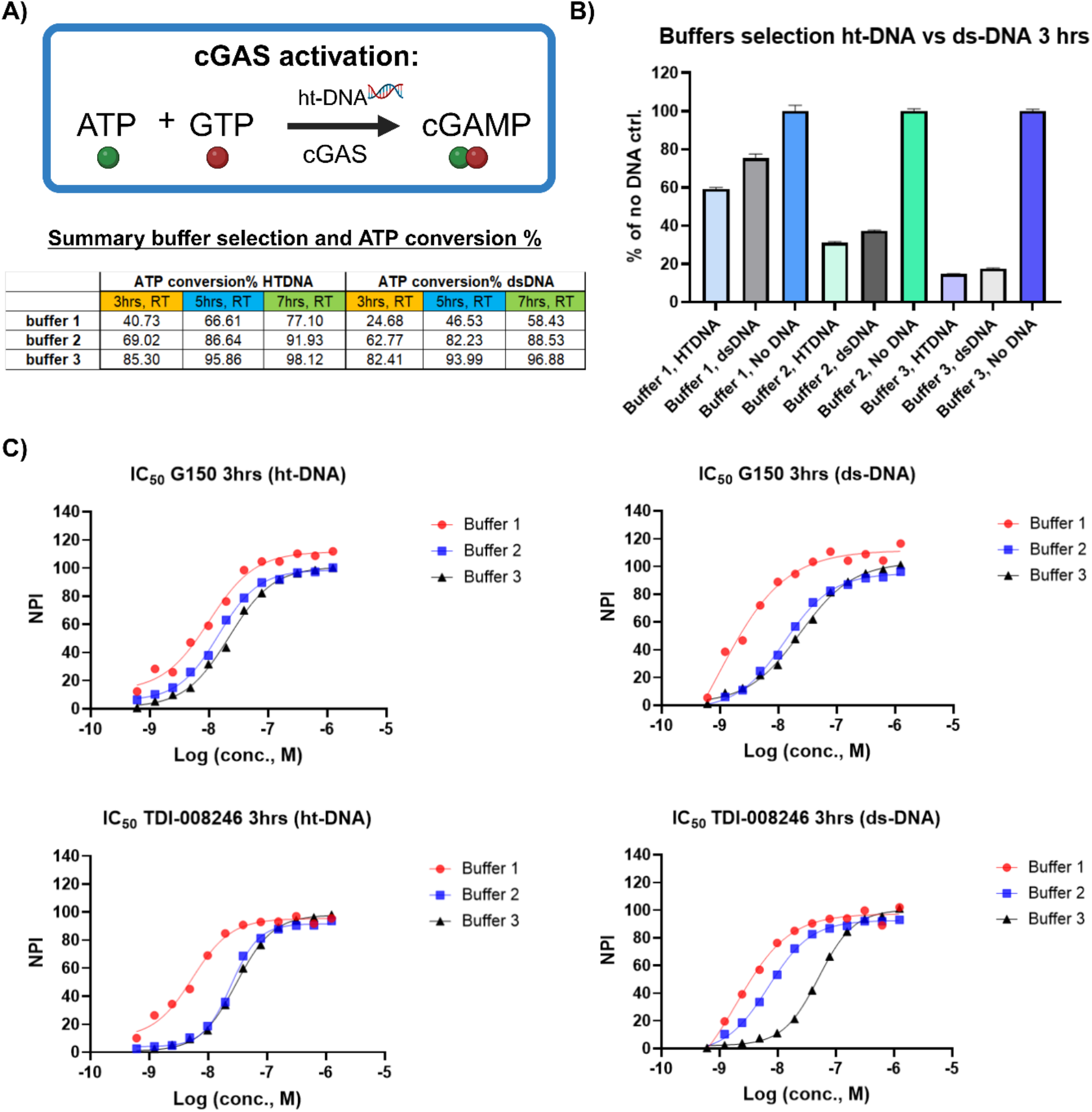
cGAS-catalysis is greatly enhanced at lower NaCl concentration and in the presence of MnCl_2_. A summary of buffers and ht-DNA vs. dsDNA showing effects on cGAS activity: **A)** over 3-, 5-, and 7-hours period depicted as a Table, and **B)** at 3-hour period depicted in a graph format. **C)** IC_50_ curves for known h-cGASI **G150** and **TDI008246** in three buffers using ht-DNA (5 µg/ml conc.) vs dsDNA (25 nM conc.) (n = 3; mean ± S.D.). ***Buffer composition***: ***B1***: 20 mM Tris-HCl pH 7.4, 150 mM NaCl, 5 mM MgCl_2_, 1 µM ZnCl_2_, 1 mM DTT and 0.01% Tween-20; ***B2***: 20 mM Tris-HCl pH 7.4, 50 mM NaCl, 5 mM MgCl_2_, 1 mM DTT and 0.01% Tween-20; ***B3***: 20 mM Tris-HCl pH 7.4, 50 mM NaCl, 2.5 mM MgCl_2_, 0.1 mM MnCl_2_, 1 mM DTT and 0.01% Tween-20.

After establishing buffer *B2* and selecting ht-DNA as the stimulatory ligand to consistently and effectively activate HIS-h-cGAS in the assay, we determined the Michaelis Mentes parameters establishing the optimal assay concentration of all the reagents, including cGAS and ht-DNA, required for cGAS catalysis of ATP+GTP→cGAMP reaction (see: Supporting Information (SI) and Figure 3 for details). Briefly, to determine optimal cGAS and ht-DNA concentration, we incubated ATP and GTP (100 µM each) in buffer *B2* at room temperature using ht-DNA (0-20 µg/ml) and h-cGAS (0-400 nM), respectively, in two separate experiments and measured unconsumed ATP using Kinase-Glo® at multiple time points (Figure 3). In these experiments, reactions without ht-DNA or without cGAS were used as the negative control. We found that ht-DNA efficiently activated HIS-h-cGAS at a concentration of 0.625 µg/ml (Figure 3A), while 50 nM HIS-h-cGAS catalyzed a reaction greater than 50% as measured by ATP conversion (Figure 3C). Michaelis Menten kinetics showed that ht-DNA activated cGAS catalysis with high affinity, K_m_, a parameter indicative of the substrate concentration at which the reaction velocity is half of the maximum velocity (V_max_/2).^23^ We selected a time point from our experiment between 2-4 hours for the calculation and obtained a K_m_ = 0.25 µg/ml of ht-DNA titration. For the HTS, we used approximately 10 times the K_m_ value (2 µg/ml) to ensure the enzyme was nearly saturated with DNA, resulting in a robust assay and reduced sensitivity to substrate fluctuations. Additionally, we performed a DMSO tolerance experiment, which revealed that the luminescence signal was only slightly affected by the concentration of DMSO (Figure 3D). In all these experiments, the negative control conditions (‘NO DNA’ or ‘NO cGAS’) showed no activity, regardless of incubation time, providing us with the opportunity to use any of them as a negative control. We proceed with No DNA as a control, besides using a lower DMSO concentration (0.2%) in our future experiments.

**Figure 3.**
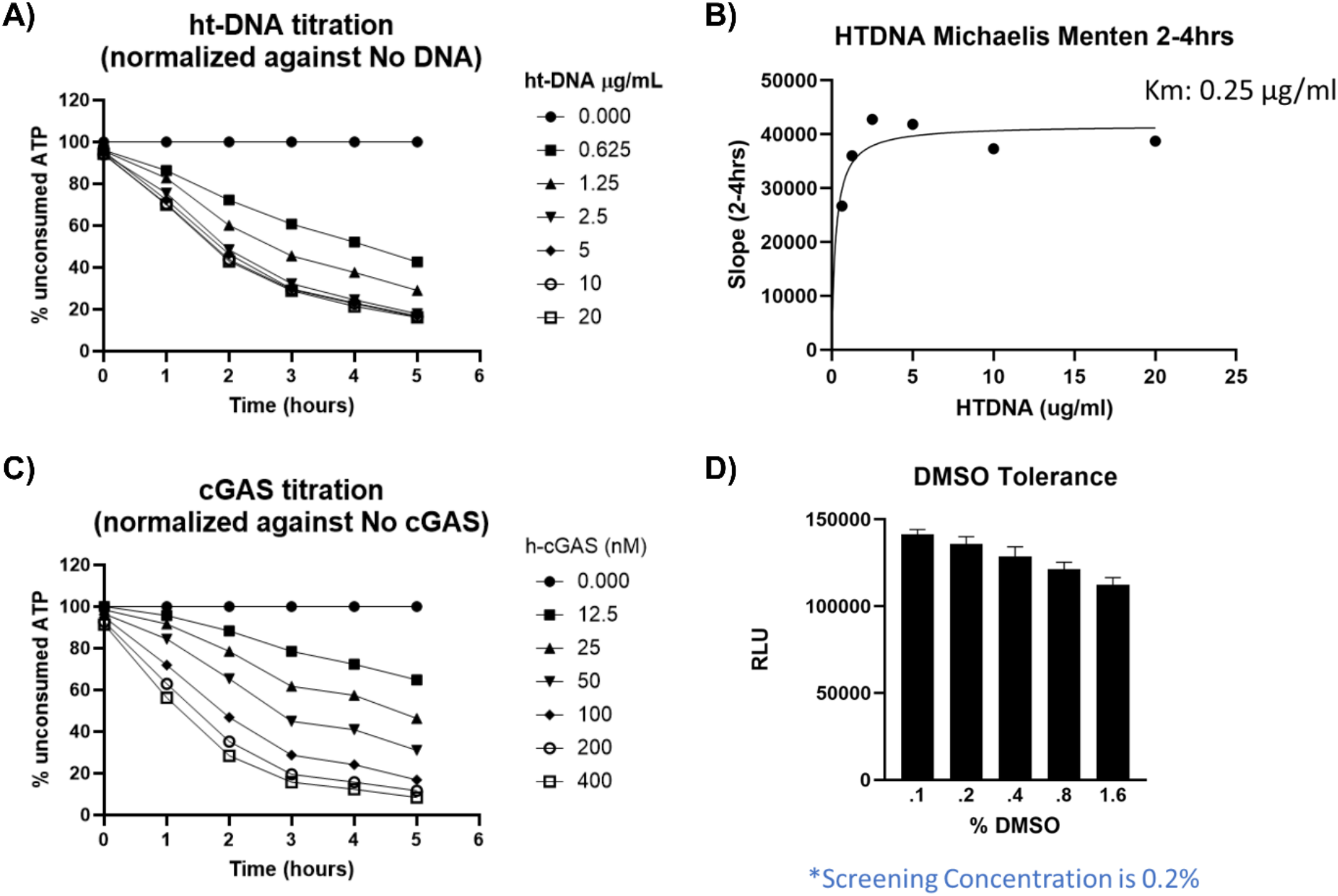
**A)** Titration of ht-DNA, normalized against ‘NO DNA condition’. **B)** Michaelis-Menten kinetics of ht-DNA. **C)** Titration of cGAS, normalized to the ‘NO cGAS condition’. The experiments were performed using ht-DNA (5 µg/ml conc.) and *B2*. **D)** DMSO tolerance assessment. ***Buffer composition***: ***B2***: 20 mM Tris-HCl pH 7.4, 50 mM NaCl, 5 mM MgCl_2_, 1 mM DTT and 0.01% Tween-20 (n = 4; mean ± S.D.).

### Pilot screening using the LOPAC library

To determine and confirm that the screening method developed above is appropriate for a large HTS campaign, we performed pilot experiments using the LOPAC library, comprising of 1,266 diverse, active pharmaceutical compounds on two different days and determined the Z’ value of the results. The Z’ evaluates the statistical reliability and robustness of an experimental design by determining the standard deviations (σ) and means (µ) of both positive and negative control sets (p and n, respectively) and feeding into *Eq. 1*.^24^ Ideally, Z’ values between 1 and 0.5 are preferred, with lower standard deviations indicating uniform controls and a Z’ value closer to one denoting optimal performance. We found that the hit rates from the pilot experiment were low, the hits identified each day were reproducible, and our screening yielded a Z’-prime value of > 0.8, providing a high degree of confidence in our screening method (see Figure 4).

**Figure 4.**
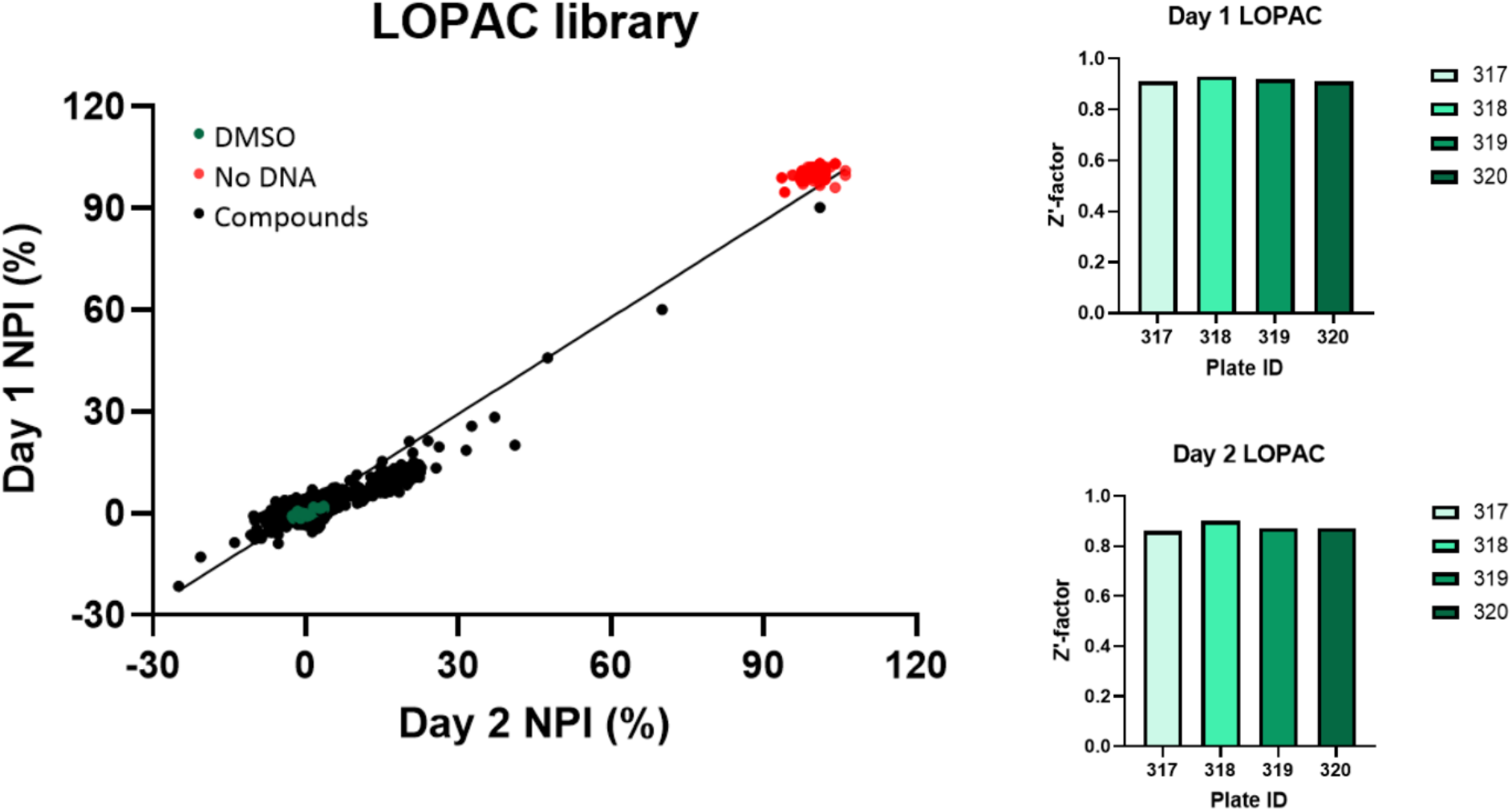
Reproducibility of the LOPAC library analyzed over two different days (R^2^ = 0.97). ***Assay condition***: 100 nM h-cGAS, 2 µg/ml ht-DNA, 100 µM ATP/GTP in buffer *B2*, RT, 3 hours.

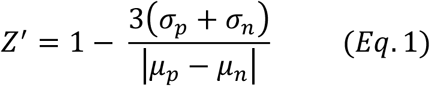

### Primary Screening and Secondary confirmatory assays

Subsequently, we performed HTS of a new collection of chemical library acquired by DDRC at the Rockefeller University. The library contains a collection of 302,000 compounds, obtained from various commercially available libraries (for more details, see the SI), and that had not been previously screened against human and mouse cGAS.^14, 25^ We initiated the primary screening using our modified HTS protocol, described above, and setting the concentration of the screening compounds at 10 µM and setting the minimum threshold to 50% inhibition of cGAS, otherwise following the general methods and the Kinase-Glo® assay for detection as described previously.^14^ We anticipated that a lower concentration (10 µM vs. 12.5 µM) and higher cGAS inhibition threshold would eliminate weak hit candidates in the primary screening.

From the initial screening campaign, all compounds exhibiting greater than 40% inhibition of cGAS activity were considered for further evaluation. Several less active compounds bearing similar pharmacophores were also included to facilitate clustering into structural families and gather data for preliminary structure–activity relationship (SAR) analysis. Following visual inspection, scaffold diversity assessment, and prioritization of chemical novelty, a total of 51 hit compounds were selected for cherry-picking and follow-up studies. These hits were divided into 6 families, 5 of which present unique pharmacophores in full or part with 4-9 members, while family 6 contains 10 hits each with a unique pharmacophore (*F1-F6*, Figure S2 and Table S1 in SI). We re-evaluated all 51 hits using both Kinase Glo® assay as well as BellBrooks’s Tanscreener® cGAMP cGAS TR-FRET assay kits.^26^ The latter determines GAMP levels in the reaction mixture, unlike the Kinase Glo® assay, which measures only the consumption of ATP, thereby providing a secondary proof and confirmation for the identified hits. From this selection, 12 top hits were chosen as confirmed primary hits for further evaluation and analysis.

### Representative Compounds from HTS

Next, all 12 hits were assessed for their identities and purities by subjecting them to LC-MS analysis and determining the enzyme inhibitory constant (IC_50_) using both the kinase Glo® and the Transcreener® assays, as described above. We found that all 12 hits have a purity of greater than 90% and possess the desired mass, and but only 6 hits showed a measurable IC_50_ below 16.67 µM, the maximum concentration used in this experiment. The results of LC-MS analysis and IC_50_ determination are provided in SI Table S2. Finally, we performed a DNA intercalation assay to identify and exclude hits that exhibited DNA intercalation (Table S3 in SI). Such hits may or may not target the enzyme’s catalytic site, but the DNA intercalation can cause toxicity. We found all selected hits did not or minimally intercalate to DNA. We prioritized top two hits **RU-0350627** (JA-1, IC_50_ = 0.52 µM) and **RU-0610270** (JA-2, IC_50_ = 1.88 µM) belonging to families *F1* and *F2* (Figure 5). Both hits possess drug-like properties and are readily available synthetically. Interestingly, we have also identified JA-1 as a potent h-cGAS inhibitor hit independently using the virtual HTS screening strategy and screening of the virtual primary hits, including numerous analogs of JA-1. A comprehensive study of JA-1 is currently underway and will be reported in a separate manuscript from our group. In this study, we focused on JA-2 as an h-cGAS inhibitor hit bearing a novel sulfonamide scaffold.

**Figure 5.**
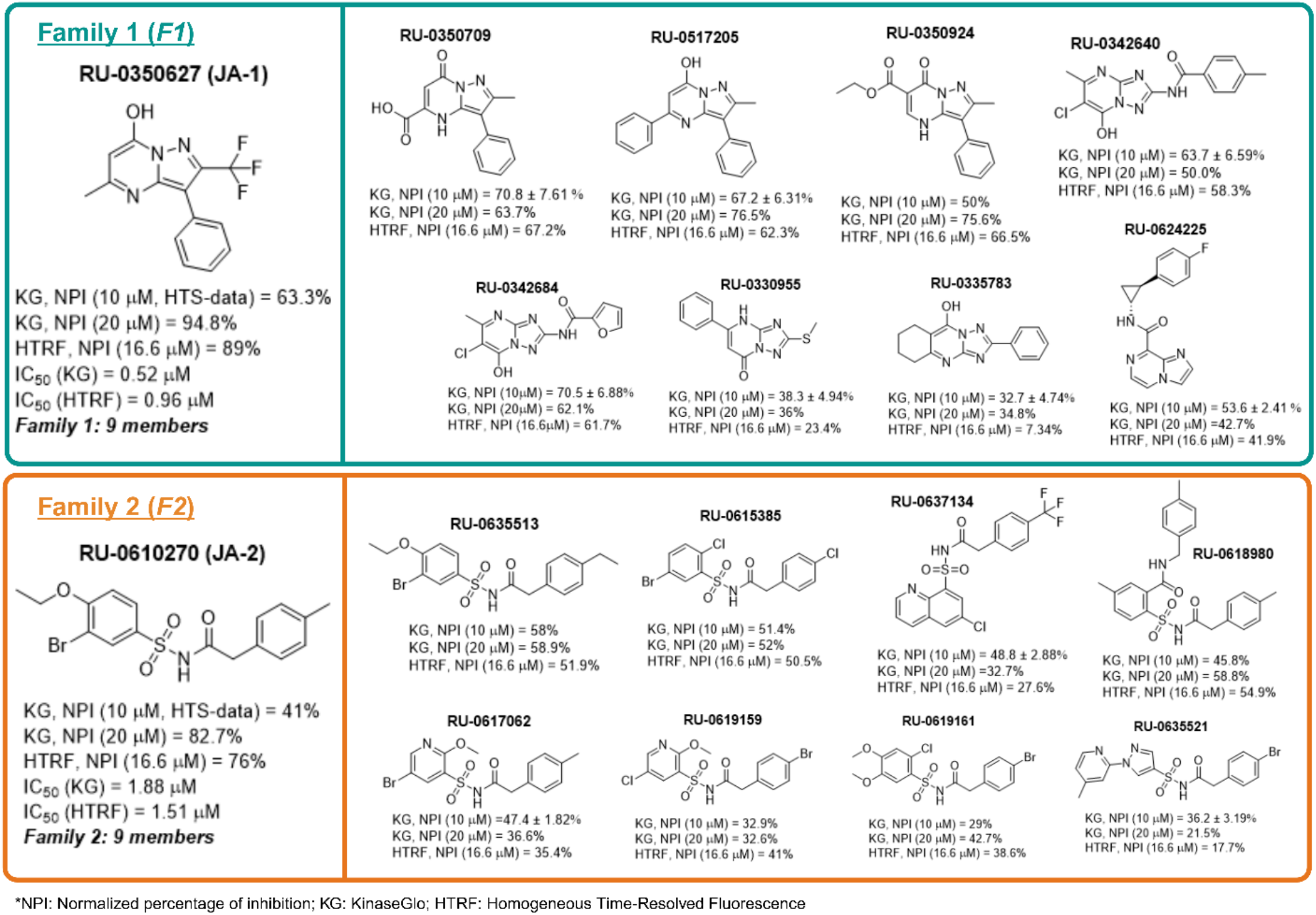
Top h-cGAS inhibitor hits identified from HTS of 300K compounds. Shown are structure of the most potent hits from two families (*F1* and *F2*) and analogs present in the HTS compound collection.

With the hit compound JA-2 selected for further evaluation, we first synthesized it using the commercially available starting materials and confirmed its activity in a cell-free assay. As the sulfonamide scaffold in JA-2 presents a new opportunity to develop cGAS inhibitors and explore their fit within the catalytic binding site of the enzyme, we decided to gather primary SAR information by evaluating the commercially available analogs. For this, we obtained 149 analogs from the ENAMINE collection and screened in vitro using the Kinase Glo® assay. Surprisingly, none of the commercially available analogs, which differ considerably from JA-2, demonstrated better activity than the original hit (data not shown). Therefore, we designed and prepared 49 new JA-2 analogs that retain the sulfonamide linker and possess new substituents in one or both phenyl rings, 1 and 2 (Scheme 2). Some compounds also lacked the -CH_2_-spacer between ring 2 and the -CO-function. All 49 analogs of JA-1 were synthesized using the commercially available sulfonamide (**I**) and acid (**II**) starting materials as outlined in Scheme 2.

**Scheme 2.**
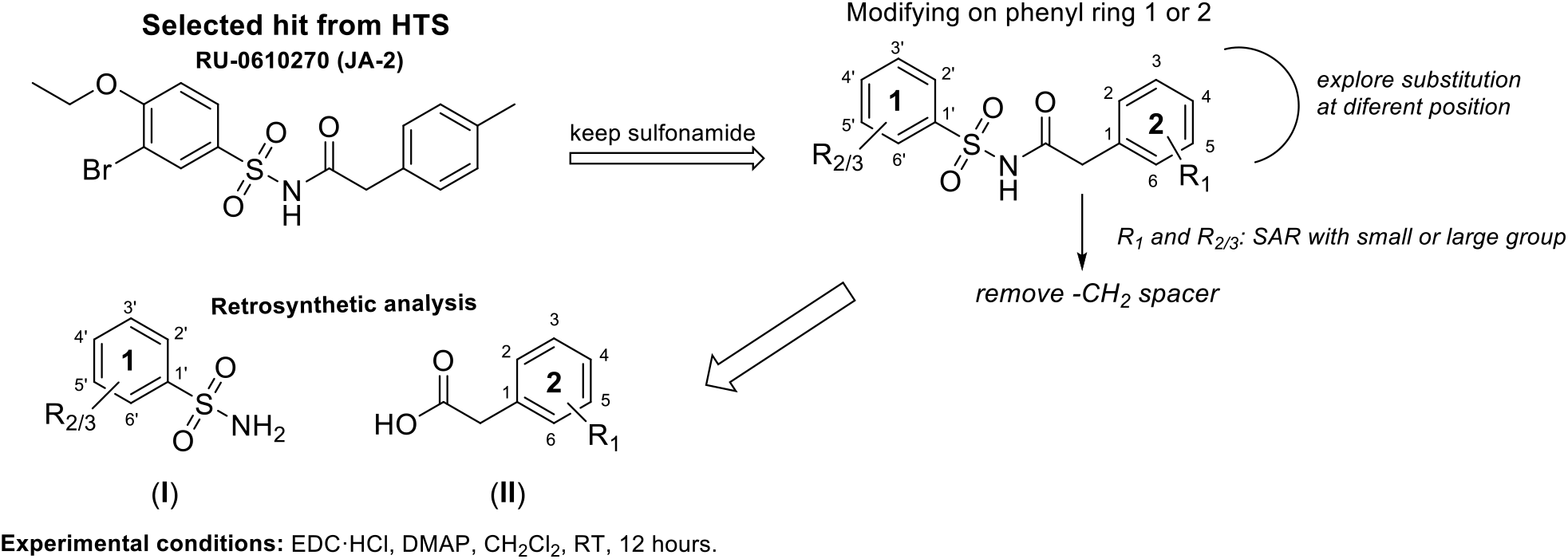
Sulfonamide derivatives design and retrosynthesis.

We assessed the in vitro activity of the resulting JA-2 analogs using Kinase glo® assay, as described previously. Initially, we evaluated all 49 compounds by incubating with 100 µM ATP and GTP, 100 nM h-cGAS and 2 µg/ml ht-DNA at room temperature, as used in the HTS campaign. The results of these experiments are summarized in Figure 6 and Table 2. Thus, we identified five active compounds with IC_50_ values between 1.1 and 5.8 µM, and two with IC_50_ values of 24.5 and 48.6 µM. We also tested the activity of five active compounds by performing experiments using a lower cGAS concentration (25 nM) at physiological temperature, 37°C. We found no significant difference in the IC_50_ of these compounds.

**Table 2.**
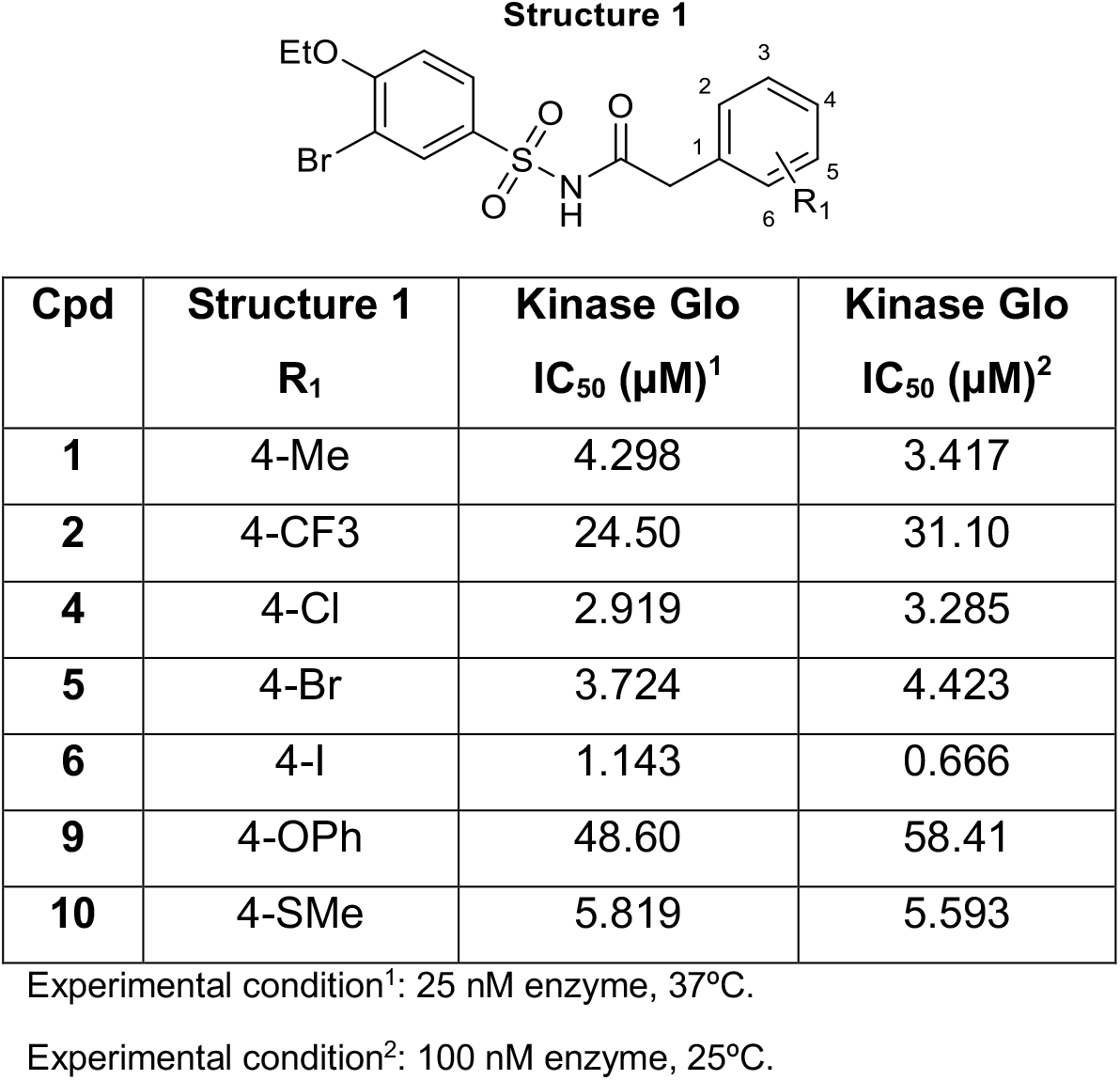
IC_50_ data for selected sulfonamide derivatives in Kinase Glo under different experimental conditions.

| Cpd | Structure 1<br>R<sub>1</sub> | Kinase Glo<br>IC<sub>50</sub> (μM)<sup>1</sup> | Kinase Glo<br>IC<sub>50</sub> (μM)<sup>2</sup> |
| --- | --- | --- | --- |
| **1** | 4-Me | 4.298 | 3.417 |
| **2** | 4-CF<sub>3</sub> | 24.50 | 31.10 |
| **4** | 4-Cl | 2.919 | 3.285 |
| **5** | 4-Br | 3.724 | 4.423 |
| **6** | 4-I | 1.143 | 0.666 |
| **9** | 4-OPh | 48.60 | 58.41 |
| **10** | 4-SMe | 5.819 | 5.593 |

Experimental condition<sup>1</sup>: 25 nM enzyme, 37°C.
Experimental condition<sup>2</sup>: 100 nM enzyme, 25°C.

**Figure 6.**
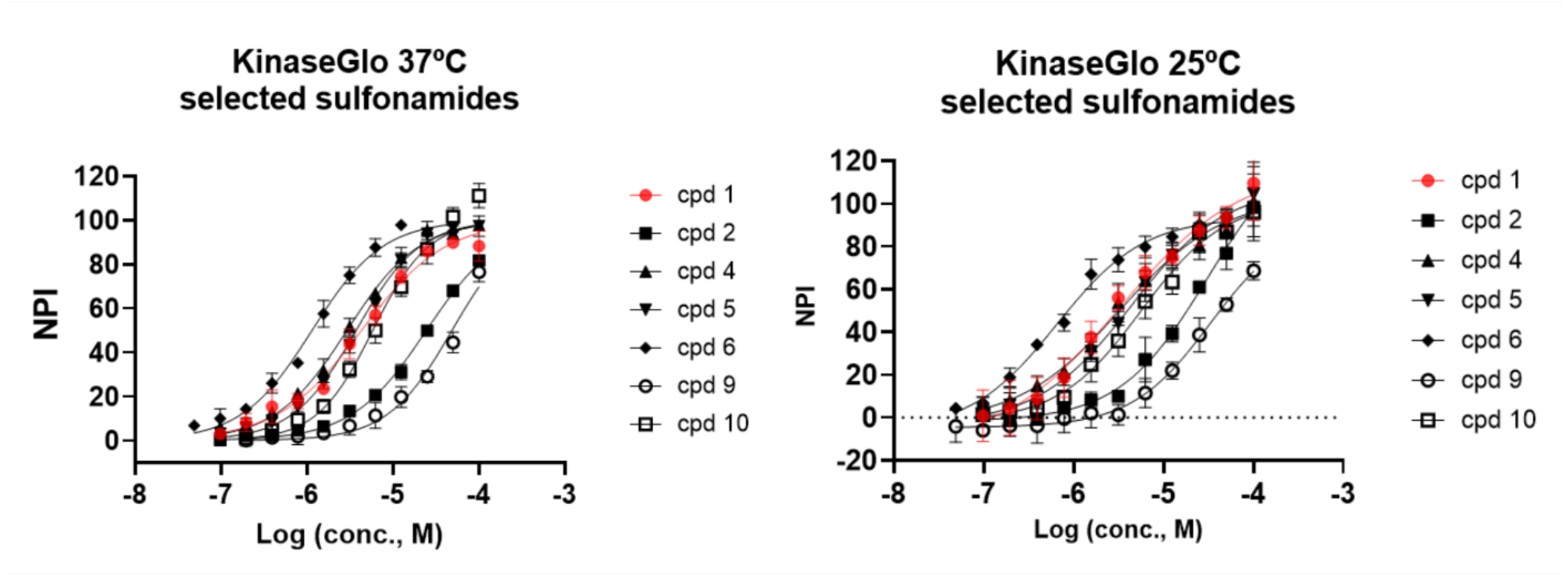
IC_50_ plots selected sulfonamide analogs. Kinase Glo (left, 37ºC and 25 nM enzyme) and (right, 25ºC and 100 nM enzyme). Corresponding IC_50_ values are listed in Table 1.

### SAR analysis of the compounds

Initially, we modified the substituents on ring 2, keeping ring 1 unchanged in **compounds (cpds) 1-24** and assessed their effects on activity. We found that an ortho -CH_3_ group, as present in **cpd 13**, exhibited reduced activity, while substituting halogens in the para position revealed the following activity trend: I (**cpd 6**) > Br (**cpd 5**) > Cl (**cpd 4**) > F (**cpd 3**). Several factors can explain this trend: *(i)* Iodine’s large size and high polarizability enhance Van der Waals interactions, increasing activity, whereas fluorine, with low polarizability, contributes weakly. *(ii)* Although fluorine’s high electronegativity reduces phenyl ring electron density, polarizability and interaction potential often outweigh this effect. *(iii)* Iodine’s higher hydrophobicity improves binding within hydrophobic pockets. *(iv)* A larger size of iodine is likely needed to create a stronger Van der Waals interaction of the molecule in the protein active site. *(v)* Substituent position is critical, as evidenced by reduced activity for ortho-iodine (**cpd 16**) compared to para-iodine (**cpd 6**), highlighting the importance of steric compatibility with the binding pocket.^27-31^

We further investigated modifications in ring 2 by focusing on para substituents -CF_3_ (**cpd 2**), -CH_3_ (**cpd 1**), and -SCH_3_ (**cpd 10**). Notably, the activity followed the trend -CH_3_ > -SCH_3_ > - CF_3_ (see Table S2 in SI), which could be influenced by steric and electronic effects as well as their lipophilicity. Presumably, the small -CH_3_ substituent in **cpd 1** minimizes its steric hindrance and fits well in the binding pocket. In contrast, the bulkier -CF_3_ and -SCH_3_ substituents in **cpds 2** and **10** could cause potential steric clashes and the substituent-induced conformational changes, thereby disrupting ligand-target interactions and reducing their activities.^32, 33^ Alternatively, a moderate lipophilicity and electron-donating property of the -CH_3_ group could be optimal for supporting both hydrophobic and electronic interactions, unlike the most lipophilic and electronegative substituent -CF_3_ among the three substituents present in **cpds 1, 2**, and **10**.^32, 33^

Subsequently, we modified ring 1 by altering its substituents, positions, or aromatic ring type, while keeping the rest of the molecule constant. Surprisingly, none of the compounds exhibited higher activity than **cpd 1** (see Table S2 in the SI for more details). For instance, replacing -Br with hydrogen (**cpd 26**) led to a loss of activity, as did changing -OCH_2_CH_3_ to -OCH_3_ (**cpd 27**) or removing all substituents on ring 1 (**cpd 25**). Similarly, para substituents: -OCF_3_ in **cpd 28**, -CCH_3_ in **cpd 29**, and -CF_3_ in **cpd 30** showed negligible activity compared to **cpd 1**, which features -Br and -OCH_2_CH_3_. Substituting phenyl ring 1 with benzo[d]thiazole (**cpd 47**) or quinoline (**cpd 48**) or phenyl ring 2 with furan (**cpd 41**) also resulted in inactive analogs. These findings highlight the importance of the -Br and -OCH_2_CH_3_ substituents in ring 1 as well as of the phenyl scaffold for maintaining activity. Attempts to modify the substituents, alter their positions, or replace the aromatic ring consistently resulted in reduced or abolished activity compared to **cpd 1**.

### Cell based assay

Finally, we assessed the inhibitory effects of **cpds 1, 4, 5, 6** and **10** on h-cGAS activity at 10 or 30 µM compound concentration in a cell-based assay using THP1 Dual cells, a human monocytic reporter cell line engineered to simultaneously monitor NF-κB and IFN pathway activation via secreted luciferase and alkaline phosphatase reporters, respectively, as described previously.^14^ Briefly, cells plated in 96-well plates were incubated with the test compounds at 10 or 30 µM concentration and ht-DNA/lipofectamine for 16 hours and their effects on IFN and NF-kB expression were determined by measuring the luciferase and phosphatase activity in supernatants and on cell viability by measuring the ATP production from cell pellets. While all the above-tested h-cGAS inhibitors proved nontoxic at both 10 and 30 µM concentrations (Figure 7A), they also showed no inhibitory effects on IFN and NF-kB expression, as determined using the luciferase and phosphatase activity data (Figures 7C and 7D). This is not surprising because most cGAS inhibitors have shown >50-100 difference between cell-free cGAS inhibition vs. THP1 cell-based IFN expression. Indeed, we found that **cpd 6**, the most potent inhibitor (based on the cell-free assay) of all sulfonamide derivatives described above, reduced IFN levels by 75.52%. However, it did not affect NF-kB levels at 100 µM, but it also showed some toxicity (∼20%) at this concentration. The compound also reduced cGAMP levels by 71.49% at a 100 µM concentration and 26.68% at a 30 µM concentration in THP1 cell lysates (Figure 7E).

**Figure 7.**
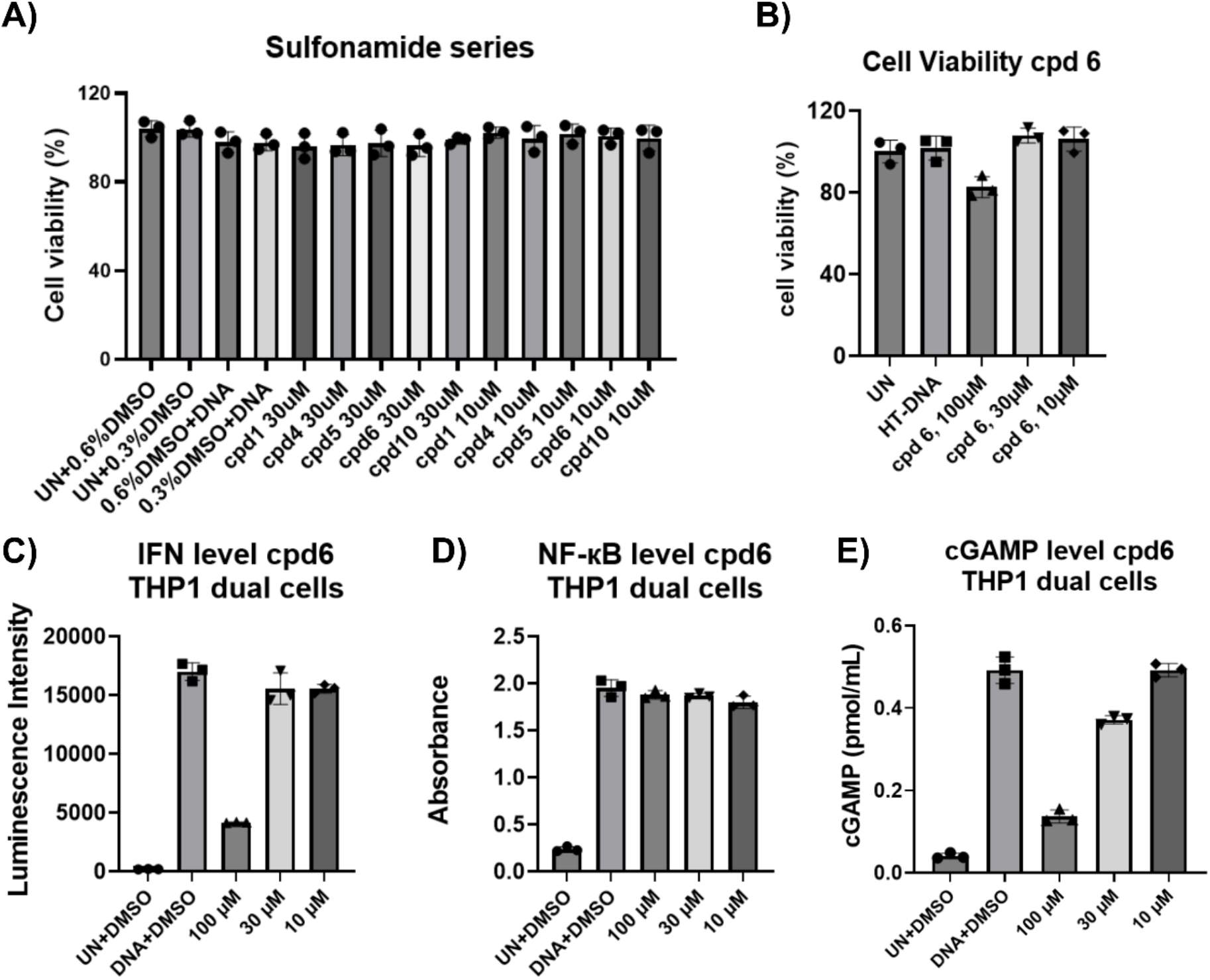
**A)** Cell viability (%) of selected compounds in THP1 dual cells. Effect of **cpd 6** (10– 100 µM) on **B)** Cell viability, **C)** IFN, and **D)** NF-κB activity in THP1-Dual cells. Untreated and DNA cells were included as controls. **E)** cGAMP ELISA of **cpd 6**. (n = 3; mean ± S.D.).

Finally, we evaluated the permeability of **cpd 6** using the MCDKII-MDR1 cells, which not only estimate cell and brain permeability of a compound but also measure efflux due to P-glycoprotein (Pgp). The results, summarized in Table 3, indicate that **cpd 6** exhibits high cell permeability and low efflux, supporting its suitability for further optimization. This data also confirms that the low potency in cellular assays is not due to issues with cellular permeability.

**Table 3.** Permeability results of cpd 6 and control compounds in MDCKII-MDR1 cell line.

| Compound ID | $P_{app}$ (A-B)<br>($10^{-6}$, cm/s) | $P_{app}$ (B-A)<br>($10^{-6}$, cm/s) | Efflux<br>Ratio | Recovery<br>(%) A-B | Recovery<br>(%) B-A |
| --- | --- | --- | --- | --- | --- |
| Metoprolol | 28.56 | 27.96 | 0.98 | 109.14 | 97.41 |
| Digoxin | 0.46 | 11.12 | 24.89 | 95.47 | 101.32 |
| **Cpd 6** | 17.30 | 23.38 | 1.35 | 95.95 | 97.54 |

## Conclusion

In summary, we found sulfonamide as a new scaffold in our HTS targeting human cGAS. The selected family allowed us to explore new modifications and their effects on the observed activity. We successfully obtained compounds with sub-micromolar to low micromolar activity. Evaluation using THP1 dual cells showed that the top analog, **cpd 6**, inhibited both cGAMP production and IFN levels, albeit at a high concentration. Further analysis of the SAR generated in this study is under progress and will be used with other ongoing studies to develop more potent cGAS inhibitors.

### Experimental section

#### Reagents and materials

All reagents are of the highest purity commercially available. ATP, GTP, and HTDNA were purchase from sigma Aldrich. The cGAS enzyme was costume made by Genscript, human cGAS (**type:** Recombinant enzyme; **lot number:** U6746IA020-3/P3IB001; **expression host:** E.coli; **purification:** Protein was obtained from supernatant of cell lysate, Ni column + superdex 200; **concentration:** 1.11 mg/ml (by Bradford); **purity:** ≥ 90% (SDS-PAGE under reducing condition); **mass:** Confirmed by LC-MS; s**torage Buffer:** 50 mM Tris-HCl, 500 mM NaCl, 10% Glycerol, pH 8.0). **cGAS Protein Length** = 528 amino acids. **Sequency**: MHHHHHHQPWHGKAMQRASEAGA TAPKASARNARGAPMDPTESPAAPEAALPKAGKFGPARKSGSRQKKSAPDTQERPPVRATGARAKKAPQRAQDTQPSDATSAPGAEGLEPPAAREPALSRAGSCRQRGARCSTKPRPPPGPWDVPSPGLPVSAPILVRRDAAPGASKLRAVLEKLKLSRDDISTAAGMVKGVVDHLLLRLKCDSAFRGVGLLNTGSYYEHVKISAPNEFDVMFKLEVPRIQLEEYSNTRAYYFVKFKRNPKENPLSQFLEGEILSASKMLSKFRKIIKEEINDIKDTDVIMKRKRGGSPAVTLLISEKISVDITLALESKSSWPASTQEGLRIQNWLSAKVRKQLRLKPFYLVPKHAKEGNGFQEETWRLSFSHIEKEILNNHGKSKTCCENKEEKCCRKDCLKLMKYLLEQLKERFKDKKHLDKFSSYHVKTAFFHVCTQNPQDSQWDRKDLGLCFDNCVTYFLQCLRTEKLENYFIPEFNLFSSNLIDKRSKEFLTKQIEYERNNEFPVFDEF.

#### Buffer compositions

***B1***: 20 mM Tris-HCl pH 7.4, 150 mM NaCl, 5 mM MgCl_2_, 1 µM ZnCl_2_, 1 mM DTT and 0.01% Tween-20.

***B2***: 20 mM Tris-HCl pH 7.4, 50 mM NaCl, 5 mM MgCl_2_, 1 mM DTT and 0.01% Tween-20.

***B3***: 20 mM Tris-HCl pH 7.4, 50 mM NaCl, 2.5 mM MgCl_2_, 0.1 mM MnCl_2_, 1 mM DTT and 0.01% Tween-20.

#### Kinase-Glo assay

The experimental conditions followed Lama *et al*.^*14*^ with modifications. Briefly, reactions were performed in 384-well plates using the cGAS enzyme, ht-DNA, ATP, and GTP at the indicated concentrations (100 nM, 2 µg/mL, 100 µM, respectively), with compounds tested at 10 µM in *B2*. After a 3-hour incubation, the reactions were stopped with Kinase-Glo Max, and luminescence was measured. Compounds showing greater than 50% inhibition were further tested in dose-response studies. Full details are provided in the SI.

#### TR-FRET assay

A brief description of the TR-FRET assay is provided here. In summary, the assay detects cGAMP production by displacing a fluorescent tracer, resulting in a decrease in the TR-FRET signal. Reactions were set up in 384-well plates with cGAS enzyme (25 nM), ht-DNA (2 μg/ml) or dsDNA (60 nM), ATP (100 μM), and GTP (100 μM) under specified conditions. Compounds were added at the desired concentrations and incubated; the reactions were then stopped before the TR-FRET signal was read. Calibration curves were performed with cGAMP standards. Complete details, including buffer composition, controls, and modifications, are provided in the SI.

#### DNA intercalator assay

We performed a DNA intercalator assay using fluorescence polarization (FP) to evaluate HTS hits and exclude those with intercalation effects following the method described by Lama *et al*.^*14*^ Briefly, reactions were set up in 384-well plates using dsDNA (200 nM) or ht-DNA (40 μg/ml), Acridine Orange (50 nM), and Mitoxantrone (20 μM, positive control). Compounds were tested at 33.3 µM. After 30 min incubation at room temperature, FP was measured with excitation at 485 nm and emission at 530 nm. Full experimental details are provided in the SI.

#### Cell culture

THP1-Dual™ cells (InvivoGen) were cultured in RPMI 1640 (Gibco™) supplemented with 10% FBS and 1% penicillin/streptomycin. To keep luciferase expression, 100 µg/ml of zeocin and 10 µg/ml of blasticidin were added to the growth medium every other passage.

#### Cell-based IRF-Lucia luciferase and cell viability

THP1-Dual cells were seeded in 96-well plates and treated with compounds or DMSO control for 4 hr. Cells were transfected with ht-DNA/Lipofectamine 2000 overnight. Luciferase activity in supernatants was measured using QUANTI-Luc reagent. % Inhibition was calculated relative to controls. Cell viability was assessed with CellTiter-Glo® 2.0. Full details are provided in the SI.

#### cGAMP ELISA assay

0.25 million THP1-dual cells were treated with serial dilutions of compounds (100–1 µM) for 4 hrs, then transfected with ht-DNA/Lipofectamine 2000 and incubated overnight. Cells were lysed in RIPA buffer containing protease and phosphatase inhibitors. cGAMP levels in cell lysates were measured using a 2,3-cGAMP ELISA kit (Arbor) following the manufacturer’s protocol. Full details are provided in the SI.

#### Preparation of final compounds

All the NMR and LC-MS data can be found in supporting information.

**Scheme 3.**
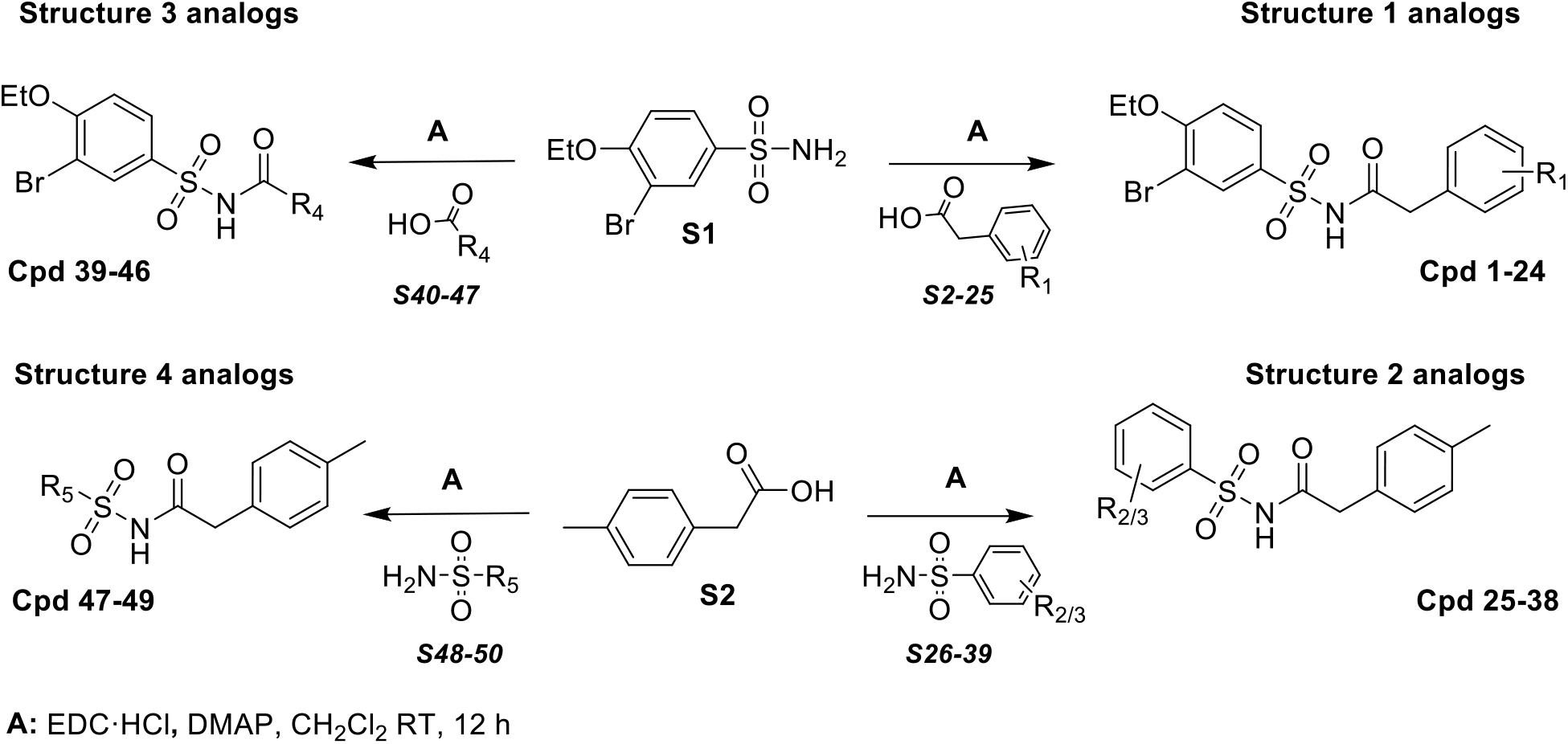
Synthesis of **structures 1-4** sulfonamide analogs.

##### Cpds 1-24 and 39-46

A mixture of 3-bromo-4-ethoxybenzenesulfonamide **S1** (50 mg, 0.178 mmol), substituted carboxylic acids **S2-25** or **S40-47** (0.196 mmol), EDC·HCl (0.357 mmol) and DMAP (0.375 mmol) in dichloromethane (2 mL) was stirred at room temperature for 12 h. The reaction was monitored by TLC and LC-MS. After completion of the reaction, the reaction mixture was diluted with water and acidified to a pH of 1 with the addition of 10% aq. HCl and followed by extracted with dichloromethane. The organic layer was dried over Na_2_SO_4,_ and the organic solvent was evaporated on a rotary evaporator. The crude residue was purified by column chromatography on silica gel to afford the desired products, **cpds 1-24** and **39-46**.

##### Cpds 25-38

A mixture of 2-(*p*-tolyl)acetic acid **S2** (30 mg, 0.196 mmol), substituted sulfonamides **S26-39** or **S48-50** (0.178 mmol), EDC·HCl (0.357 mmol) and DMAP (0.375 mmol) was similarly treated in dichloromethane (2 mL), as above, and worked-up and purified by column chromatography to afford the desired products, **cpds 25-38** and **47-49**.

## Acknowledgments

We are thankful to NIH/NIA (R01AG074541 to SCS, LG; R01AG072758 to LG; R43AG080954 to LG and AS of Aeton Therapeutics), Rainwater and JPB Foundations, and Cure Alzheimer’s fund and Daedalus fund for Innovation (to L.G., S.C.S.) for financial support. We are also thankful to IRBM teams for performing some of the screening of compounds and helpful discussions, Pharmaron for performing ADME experiments, and Dr. Anjana Sinha of Aeton Therapeutics and especially to the scientific advisors from Rainwater foundation for the discussions and guidance.

## Author contributions

S.C.S. and L.G. conceived the project and planned experiments. S.C.S and J.A.E designed experiments. J.A.E. performed experiments, developed experimental protocols, and analyses data. R.K.N., S.W. and V.K.S. performed the synthesis of the compounds, biochemical experiments and analysis of the results. C.L., L.C.T and F.G. provided technical support and troubleshooting. J.A.E. and S.C.S. wrote the manuscript. All authors read, reviewed and approved the paper.

## REFERENCES

(1) Chung, S.; Jeong, J.-H.; Park, J.-C.; Han, J. W.; Lee, Y.; Kim, J.-I.; Mook-Jung, I. Blockade of STING activation alleviates microglial dysfunction and a broad spectrum of Alzheimer’s disease pathologies. Experimental & Molecular Medicine 2024, 56 (9), 1936–1951. DOI: 10.1038/s12276-024-01295-y.

(2) Govindarajulu, M.; Ramesh, S.; Beasley, M.; Lynn, G.; Wallace, C.; Labeau, S.; Pathak, S.; Nadar, R.; Moore, T.; Dhanasekaran, M. Role of cGAS-Sting Signaling in Alzheimer’s Disease. Int J Mol Sci 2023, 24 (9). DOI: 10.3390/ijms24098151 From NLM.

(3) Xie, X.; Ma, G.; Li, X.; Zhao, J.; Zhao, Z.; Zeng, J. Activation of innate immune cGAS-STING pathway contributes to Alzheimer’s pathogenesis in 5×FAD mice. Nature Aging 2023, 3 (2), 202–212. DOI: 10.1038/s43587-022-00337-2.

(4) Udeochu, J. C.; Amin, S.; Huang, Y.; Fan, L.; Torres, E. R. S.; Carling, G. K.; Liu, B.; McGurran, H.; Coronas-Samano, G.; Kauwe, G.; et al. Tau activation of microglial cGAS–IFN reduces MEF2C-mediated cognitive resilience. Nature Neuroscience 2023, 26 (5), 737–750. DOI: 10.1038/s41593-023-01315-6.

(5) Huang, Y.; Liu, B.; Sinha, S. C.; Amin, S.; Gan, L. Mechanism and therapeutic potential of targeting cGAS-STING signaling in neurological disorders. Mol Neurodegener 2023, 18 (1), 79. DOI: 10.1186/s13024-023-00672-x From NLM.

(6) Alarcón-Espósito, J.; Nagiri, R. K.; Gan, L.; Sinha, S. C. Identification and development of cGAS inhibitors and their uses to treat Alzheimer’s disease. NeurotherapeuBcs 2025, e00536. DOI: 10.1016/j.neurot.2025.e00536.

(7) Gulen, M. F.; Samson, N.; Keller, A.; Schwabenland, M.; Liu, C.; Glück, S.; Thacker, V. V.; Favre, L.; Mangeat, B.; Kroese, L. J.; et al. cGAS–STING drives ageing-related inflammation and neurodegeneration. Nature 2023, 620 (7973), 374–380. DOI: 10.1038/s41586-023-06373-1.

(8) Sun, Z.; Hornung, V. cGAS–STING signaling. Current Biology 2022, 32 (13), R730–R734. DOI: 10.1016/j.cub.2022.05.027 (acccessed 2025/08/06).

(9) Raulin, A.-C.; Doss, S. V.; Trorer, Z. A.; Ikezu, T. C.; Bu, G.; Liu, C.-C. ApoE in Alzheimer’s disease: pathophysiology and therapeutic strategies. Molecular NeurodegeneraBon 2022, 17 (1), 72. DOI: 10.1186/s13024-022-00574-4.

(10) Guerreiro, R.; Wojtas, A.; Bras, J.; Carrasquillo, M.; Rogaeva, E.; Majounie, E.; Cruchaga, C.; Sassi, C.; Kauwe, J. S.; Younkin, S.; et al. TREM2 variants in Alzheimer’s disease. N Engl J Med 2013, 368 (2), 117–127. DOI: 10.1056/NEJMoa1211851 From NLM.

(11) Jonsson, T.; Stefansson, K. TREM2 and neurodegenerative disease. N Engl J Med 2013, 369 (16), 1568–1569. DOI: 10.1056/NEJMc1306509 From NLM.

(12) Sayed, F. A.; Kodama, L.; Fan, L.; Carling, G. K.; Udeochu, J. C.; Le, D.; Li, Q.; Zhou, L.; Wong, M. Y.; Horowitz, R.; et al. AD-linked R47H-TREM2 mutation induces disease-enhancing microglial states via AKT hyperactivation. Sci Transl Med 2021, 13 (622), eabe3947. DOI: 10.1126/scitranslmed.abe3947 From NLM.

(13) Arboleda-Velasquez, J. F.; Lopera, F.; O’Hare, M.; Delgado-Tirado, S.; Marino, C.; Chmielewska, N.; Saez-Torres, K. L.; Amarnani, D.; Schultz, A. P.; Sperling, R. A.; et al. Resistance to autosomal dominant Alzheimer’s disease in an APOE3 Christchurch homozygote: a case report. Nat Med 2019, 25 (11), 1680–1683. DOI: 10.1038/s41591-019-0611-3 From NLM.

(14) Lama, L.; Adura, C.; Xie, W.; Tomita, D.; Kamei, T.; Kuryavyi, V.; Gogakos, T.; Steinberg, J. I.; Miller, M.; Ramos-Espiritu, L.; et al. Development of human cGAS-specific small-molecule inhibitors for repression of dsDNA-triggered interferon expression. Nature CommunicaBons 2019, 10 (1), 2261. DOI: 10.1038/s41467-019-08620-4.

(15) Koresawa, M.; Okabe, T. High-Throughput Screening with Quantitation of ATP Consumption: A Universal Non-Radioisotope, Homogeneous Assay for Protein Kinase. ASSAY and Drug Development Technologies 2004, 2 (2), 153–160. DOI: 10.1089/154065804323056495 (acccessed 2024/07/10).

(16) Dueñas, M. E.; Peltier-Heap, R. E.; Leveridge, M.; Annan, R. S.; Bülner, F. H.; Trost, M. Advances in high-throughput mass spectrometry in drug discovery. EMBO Molecular Medicine 2023, 15 (1), e14850. DOI: 10.15252/emmm.202114850 (acccessed 2024/08/31).

(17) Li, Q.; Tian, S.; Liang, J.; Fan, J.; Lai, J.; Chen, Q. Therapeuytic Development by Targeting the cGAS-STING Pathway in Autoimmune Disease and Cancer. FronBers in Pharmacology 2021, Volume 12 - 2021, Review. DOI: 10.3389/fphar.2021.779425.

(18) van Campen, N.; Mekers, V. E.; Looman, M. W.; van den Bogaard, L.; Kers-Rebel, E. D.; Peeters, W. J. M.; Merino, E. F.; Schuurmans, F.; Smeenk, R. J.; Verheij, M.; et al. ATM and ATR inhibition increases radiosensitivity and cGAS-STING activation in prostate cancer. Cytokine 2025, 193, 156980. DOI: 10.1016/j.cyto.2025.156980.

(19) Skeldon, A. M.; Wang, L.; Sgarioto, N.; Beveridge, R. E.; Chan, S.; Dorich, S.; Dumais, V.; Fradet, N.; Gaudreault, S.; LeGros, P.; et al. Structural insight into the cGAS active site explains di?erences between therapeutically relevant species. CommunicaBons Chemistry 2025, 8 (1), 88. DOI: 10.1038/s42004-025-01481-7.

(20) Zhao, Z.; Ma, Z.; Wang, B.; Guan, Y.; Su, X. D.; Jiang, Z. Mn(2+) Directly Activates cGAS and Structural Analysis Suggests Mn(2+) Induces a Noncanonical Catalytic Synthesis of 2’3’-cGAMP. Cell Rep 2020, 32 (7), 108053. DOI: 10.1016/j.celrep.2020.108053 From NLM.

(21) Zhang, T.; Hu, C.; Zhang, W.; Ruan, Y.; Ma, Y.; Chen, D.; Huang, Y.; Fan, S.; Lin, W.; Huang, Y.; et al. Advances of MnO2 nanomaterials as novel agonists for the development of cGAS-STING-mediated therapeutics. FronBers in Immunology 2023, Volume 14 - 2023, Review. DOI: 10.3389/fimmu.2023.1156239.

(22) Lama, L.; Tuschl, T.; Tomita, D.; Patel, D.; Glickman, J. F.; Kamei, T.; Miller, M.; Asano, Y.; Okamoto, R. E. I.; Hashizume, S.; et al. 2,3,4,5-TETRAHYDRO-1H-PYRIDO[4, 3-B]INDOLE INHIBITORS OF CGAS FOR TREATING AUTOINFLAMMATORY DISEASES. WO WO 2019/153002 A1, 2019.

(23) Brooks, H. B.; Geeganage, S.; Kahl, S. D.; Montrose, C.; Silampalam, S.; Smith, M. C.; Weidner, J.R. Basics of Enzymatic Assays for HTS. In Assay Guidance Manual, Markossian, S., Grossman, A., Baskir, H., Arkin, M., Auld, D., Austin, C., Baell, J., Brimacombe, K., Chung, T. D. Y., Coussens, N. P., et al. Eds.; Eli Lilly & Company and the National Center for Advancing Translational Sciences, 2004.

(24) Zhang, J. H.; Chung, T. D.; Oldenburg, K. R. A Simple Statistical Parameter for Use in Evaluation and Validation of High Throughput Screening Assays. J Biomol Screen 1999, 4 (2), 67–73. DOI: 10.1177/108705719900400206 From NLM.

(25) Vincent, J.; Adura, C.; Gao, P.; Luz, A.; Lama, L.; Asano, Y.; Okamoto, R.; Imaeda, T.; Aida, J.; Rothamel, K.; et al. Small molecule inhibition of cGAS reduces interferon expression in primary macrophages from autoimmune mice. Nature CommunicaBons 2017, 8 (1), 750. DOI: 10.1038/s41467-017-00833-9.

(26) Lea, W. A.; Simeonov, A. Fluorescence polarization assays in small molecule screening. Expert Opinion on Drug Discovery 2011, 6 (1), 17–32. DOI: 10.1517/17460441.2011.537322.

(27) Metrangolo, P.; Resnati, G. Halogen Bonding: A Paradigm in Supramolecular Chemistry. Chemistry – A European Journal 2001, 7 (12), 2511-2519. DOI: 10.1002/1521-3765(20010618)7:12<2511::AID-CHEM25110>3.0.CO;2-T.

(28) O’Hagan, D. Understanding organofluorine chemistry. An introduction to the C–F bond. Chemical Society Reviews 2008, 37 (2), 308-319, 10.1039/B711844A. DOI: 10.1039/B711844A.

(29) Cavallo, G.; Metrangolo, P.; Milani, R.; Pilati, T.; Priimagi, A.; Resnati, G.; Terraneo, G. The Halogen Bond. Chemical Reviews 2016, 116 (4), 2478–2601. DOI: 10.1021/acs.chemrev.5b00484.

(30) Bissantz, C.; Kuhn, B.; Stahl, M. A Medicinal Chemist’s Guide to Molecular Interactions. Journal of Medicinal Chemistry 2010, 53 (14), 5061–5084. DOI: 10.1021/jm100112j.

(31) Lu, Y.; Liu, Y.; Xu, Z.; Li, H.; Liu, H.; Zhu, W. Halogen bonding for rational drug design and new drug discovery. Expert Opinion on Drug Discovery 2012, 7 (5), 375–383. DOI: 10.1517/17460441.2012.678829.

(32) Purser, S.; Moore, P. R.; Swallow, S.; Gouverneur, V. Fluorine in medicinal chemistry. Chemical Society Reviews 2008, 37 (2), 320-330, 10.1039/B610213C. DOI: 10.1039/B610213C.

(33) Hagmann, W. K. The Many Roles for Fluorine in Medicinal Chemistry. Journal of Medicinal Chemistry 2008, 51 (15), 4359–4369. DOI: 10.1021/jm800219f.

