## Supporting Information for "High-Throughput Screening and Initial SAR Studies Identify a Novel Sub-micromolar Potent Human cGAS Inhibitor"

**Table of content**

| **Experimental methods** |
|---|
| **Plot S1.** Buffer optimization using HTDNA (5 µg/ml) vs. dsDNA (25 nM) after 3 h incubation: Summary of ATP conversion efficiency (raw data in RLU). |
| **Supplementary Figure S1A.** Primary screening workflow used in our HTS for the identification of human cGAS inhibitors. Numbers in parentheses indicate the column position in the 384-well plate and the volume dispensed in µL. **S1B.** Secondary screening. Direct method to measure cGAMP and enzyme activity by TR-FRET assay (adapted from Bellbrook protocol). **S1C**. Screening cascade cGAS inhibitors. |
| **Figure S2.** Representative and selected hits grouped by chemical families: molecular surface and in vitro properties. Molecular surfaces were generated for selected hits using Schrödinger software. Atoms are color-coded as follows: Hydrogen (H), white; Carbon (C), gray; Oxygen (O), network; Nitrogen (N), blue; Sulfur (S), yellow; Chlorine (Cl), dark green; Fluorine (F), light green. |
| **Supplementary Figure S3.** Sample preparation for cGAMP ELISA in THP1 dual cells. |
| **Supplementary Table S1.** Activity data of selected compounds from screening against Kinase Glo, TR-FRET assays and cytotoxicity. |
| **Supplementary Table S2.** Confirmation of selected hits through LC-MS. |
| **Supplementary Table S3.** DNA Intercalation Assay: Intercalator at 33.3 µM, Expressed as % Intercalation |

| |
|---|
| **Supplementary Table S4.** Library information screening. |
| **Supplementary Table S5.** Library composition screen in Fisher Drug Discovery Resource Center at the Rockefeller University. |
| **Supplementary Table S6.** Activity data in Kinase Glo and TR-FRET assay from analogs of sulfonamide selected hit. Percentage inhibition at 12.5 μM and 10.4 μM respectively. |
| **General synthesis methods**<br>**Supplementary Scheme S1**. Synthesis of Structure 1-4 sulfonamide analogs. |
| **$^1$H-NMR data and LC-MS data sulfonamide derivatives** |

### Experimental Methods

#### *Kinase-Glo assay*

The condition of the experiment was carried out based on Lama *et al.*[1] with some modifications. The final concentrations of cGAS enzyme, ht-DNA, ATP, and GTP for the screening were 100 nM, 2 μg/mL, 100 μM, and 100 μM, respectively. The final volume was 20 μl in 384 white solid bottom plates (Greiner 781075). The reaction started with 10 μl of reaction buffer that contained 20 mM Tris-HCl pH 7.4, 50 mM NaCl, 5 mM Mg*Cl*$_2$*,* and 0.01% Tween-20 were dispensed using a Liquid Dispenser MultiFlo with Stacker (BioTek, Agilent). After adding reagents, the plates were centrifuged at 180 x G for 60s. The stock solution of each compound was prepared in DMSO at 10 mM, then 40 nl of 10 mM stock compounds were dispensed for screening and 200 nl for $IC_{50}$ with a Janus 384 MDT Nano Head (PerkinElmer) into microplates. The working concentration for the primary screening was 10 μM and represented 0.2% DMSO in the final volume. The DMSO constitution did not interfere with cGAMP production from recombinant cGAS. 5 μl of master mix (4X) of 400 μM ATP, 400 μM GTP, 8 μg/ml ht-DNA prepared in 1X reaction buffer with 2 mM DTT was added using a Liquid Dispenser MultiFlo. Additionally, column 24 corresponds to the "No ht-DNA" condition (control for no enzymatic activity), 5 μl of master mix (4X) of 400 μM ATP, 400 μM GTP prepared in 1X reaction buffer with 2 mM DTT was added with a multichannel into this column. The reaction started by adding 5 μl (4X) cGAS solution prepared in 1X reaction buffer with 2 mM DTT in a concentration of 400 nM into all the columns and followed by an incubation of 3 hours at RT, with the plate sealed. After 3 hours, the reaction was stopped by adding 20 μl Kinase-Glo® Max (Promega, Madison, WI) reagent per well into all the columns, centrifuge plate at 180 x G for 60s, incubate for 10 minutes with shake, and reading the luminescence signal with a Biotek Synergy Neo plate reader (BioTek, Agilent). Compounds that exhibited more than 50% inhibition at 10 μM were re-checked in a dose-response manner to determine the half inhibitory concentration ($IC_{50}$). Compounds were serially diluted in half concentration in DMSO to complete 8-10 points and evaluated in triplicate. The high and low concentrations were 100 and 0.048 μM, respectively. Additionally, the reaction was incubated at 37°C, the final concentrations of cGAS enzyme, ht-DNA, ATP, and GTP for the screening were 25 nM, 2 μg/mL, 100 μM, and 100 μM, respectively. The data were analyzed using GraphPad Prism (10) software, and the standard deviation was included.

***TR-FRET assay***

The Transcreener® cGAMP detection mixture contains a cGAMP ATTO 647 tracer bound to a cGAMP antibody conjugated to terbium (Tb). Excitation of the Tb complex in the UV range (~330nm) results in energy transfer to the tracer and emission at a higher wavelength (665 nm) after a delay. cGAMP produced by the cGAS enzyme displaces the tracer, causing a decrease in TR-FRET. The experiment was done following the kit instructions, including some modifications related to the reaction buffer. Additionally, titration of the amount of enzyme used was performed. In addition, the original method used dsDNA, and for comparative purposes, we compared the controls' signals using dsDNA vs. ht-DNA. The final concentrations were 25 nM h-cGAS (GENSCRIPT), 60 nM dsDNA or 2 μg/ml ht-DNA, 100 μM ATP, and 100 μM GTP. The final volume was 12 μl in 384 small-volume white plates (Greiner 784075). The reaction started with 4 μl 1X reaction buffer that contained 20 mM Tris-HCl pH 7.4, 50 mM NaCl, 5 mM $MgCl_2$, 1 mM DTT, and 0.01% Tween-20 were dispensed using an automatic multichannel. Then 100nL of serial diluted stock compounds were dispensed with a Janus 384 MDT NanoHead (PerkinElmer) into microplates. 4 μl of master mix (3X) 300 μM ATP, 300 μM GTP, 6 μg/ml ht-DNA prepared in 1X reaction buffer with 1 mM DTT was added to all test wells using a multichannel. Column 24, which corresponds to the Kinase Glo assay "No ht-DNA" condition, control for no enzymatic activity in Kinase Glo assay, was changed for **G150** 5 μM, 4 μl of this master mix (3X) 300 μM ATP, 300 μM GTP, 6 μg/ml ht-DNA and 15 μM **G150** prepared in 1X reaction buffer with 1 mM DTT was added with multichannel into column 24. The reaction started by adding 4 μl (3X) cGAS solution in 1X reaction buffer with 1 mM DTT in a concentration of 300 nM into all the columns and followed by an incubation of 1 hour at RT. The plate was sealed and centrifuged. The reaction mixture was incubated at RT for 1 hour in the dark. After 1 hour, the reaction was stopped by adding 12 μl stop solution to all test wells using a multichannel and incubating for 1 hour more. The plate was sealed and centrifuged for 180 x G for 60 seconds. For the reading, we used a Biotek Synergy Neo plate reader (BioTek, Agilent). Further, a calibration curve is carried out with cGAMP to know the amount of cGAMP produced in the enzymatic activity.

***DNA intercalator assay***

We performed a DNA intercalator assay using FP to validate the hits we obtained from the HTS and discard those that showed some intercalation effect. The assay was performed in black 384-solid bottom opaque plates following the method previously described by Lama *et al*.[1] The final concentration of the reagents was 200 nM dsDNA or 40 ug/ml ht-DNA substrate, 50 nM Acridine Orange, 20 μM Mitoxantrone (known dsDNA intercalator). The reaction starts with 10 μl of 1X Reaction Buffer (20 mM

Tris-HCl pH 7.4, 50 mM NaCl, 5 mM MgCl2, 1 mM DTT, and 0.01% Tween-20) to all wells in Columns 1-23. Next, 0.2 µl of compound (screened at 33.3 µM) were dispensed per well with a Janus 384 MDT NanoHead (Perkin Elmer) to the corresponding wells in Columns 1-22. Preparation of the control wells in Column 24: for Blank (Wells A/B), add 30 µl of 1X Reaction Buffer; for the Acridine Orange control (Wells C/D), add 30 µl of 50 nM Acridine Orange in 1X Reaction Buffer; and for the Mitoxantrone control (Wells E-P), add 10 µl of 60 µM Mitoxantrone (3X) followed by 20 µl of 75 nM Acridine Orange and 300 nM dsDNA (1.5X). Finally, add 20 µl of 1.5X AO/dsDNA (75 nM Acridine Orange, 300 nM dsDNA) to all columns 1-23, centrifuge the plate for 30 seconds, and incubate at room temperature for 30 minutes. Read Fluorescence polarization using the BioTek Synergy Neo, Agilent. Samples were excited at 485 nm, and fluorescence emission was detected through a 530 nm filter.

***Cell culture***

THP1-Dual™ cells (InvivoGen) were cultured in RPMI 1640 (Gibco™) supplemented with 10% FBS and 1% penicillin/streptomycin. To keep luciferase expression, 100 µg/ml of zeocin and 10 µg/ml of blasticidin were added to the growth medium every other passage.

***Cell-based IRF-Lucia luciferase and cell viability***

THP1-Dual cells were seeded in 96-well plates ($5 \times 10^5$ cells/mL, 100 µl per well) and incubated overnight. Pretreated cells with indicated concentrations of compounds for 4 hrs. DMSO was added as negative control. Transfected cells with 2 µg/ml of herring testes DNA (ht-DNA) in complex with Lipofectamine 2000 (Invitrogen) overnight. Transfection mixture complex was prepared by combining 0.35 µg of ht-DNA in 25 µl OptiMEM (Gibco) with 0.5 µl of Lipofectamine 2000 in 25 µl Opti-MEM and adding the 50 µl combined mixture for each well containing cells. Luciferase luminescence was measured by QUANTI-Luc luciferase reagent (InvivoGen). 30 µl of cell culture supernatant per well was transferred into a 96-well plate and then add 70 µl QUANTI-Luc luciferase reagent. Luminescence was measured immediately using a Biotek Synergy Neo plate reader (BioTek, Agilent). Inhibition% of compounds against cGAS is calculated by relative luciferase activity or relative SEAP activity using DMSO-treated sample as negative control and Lipofectamine 2000: HTDNA complex-treated sample without compound as positive control. Inhibition% by luminescence = $100-(LU_{sample} - LU_{negative\ control})/(LU_{positive\ control} - LU_{negative\ control}) \times 100$, where LU indicates luminescence signal, or Inhibition% by relative SEAP activity = $100-(Abs_{sample} - Abs_{negative\ control})/(Abs_{positive\ control} - Abs_{negative\ control}) \times 100$. Cell viability was measured by CellTiter-Glo® 2.0 (Promega) following the manufacturer's protocol.

***cGAMP ELISA assay***

0.25 million THP1-dual cells in 500 µl media were dispensed into a 24-well plate, where the compounds in DMSO in serial dilution in a final concentration between 100 µM to 1 µM were duplicated. A master mix of compounds with a buffer with 4.5 µl compound DMSO solution was mixed with 245.5 µl media for dilution, and 100 µl of the mix was dispensed into the cells and incubated for 4 hours. After that, the transfection process was prepared following the vendor's instructions using Lipofectamine 2000. The final ht-DNA concentration was 2 µg/ml, and the plates were incubated overnight. The next day, the cells were harvested for the ELISA. The level of cGAMP in cell lysates was measured by a 2,3-cGAMP ELISA kit (Arbor). For sample preparation, cells were lysed in RIPA buffer (Thermo Fisher Scientific, 89901) supplemented with Halt phosphatase inhibitor cocktail (Thermo Fisher Scientific, 78420) and complete protease inhibitor cocktail (Roche, 11697498001). The cells and media were homogenized for the lysis method and transferred into an Eppendorf for each sample. Centrifuge for a pellet, remove the media and wash 2 times with PBS. The washed pellet was treated with 60 µl of the Ripa buffer master mix, mixed well, and incubated for 15 minutes at room temperature. After incubation, the samples were centrifuged at 1,500 RPM at 4°C for 15 minutes. Finally, 50 µl supernatant from the 60 µl RIPA lysed tube was used for the ELISA experiment. For the ELISA step, we are following the vendor protocol.

**Preparation of MDCKII-MDR1 Cells**

50 µL and 25 mL of cell culture medium were added to each well of the Transwell insert and reservoir, respectively. And then the HTS transwell plates were incubated at 37 °C, 5% $CO_2$ for 1 hour before cell seeding. MDCKII-MDR1 cells were diluted to 1.56 x $10^6$ cells/mL with culture medium and 50 µL of cell suspension were dispensed into the filter well of the 96-well HTS Transwell plate. Cells were cultivated for 3-8 days in a cell culture incubator at 37 °C, 5% $CO_2$, 95% relative humidity. Cell culture medium was replaced every other day, beginning no later than 24 hours after initial plating.

**Assessment of Cell Monolayer Integrity**

Medium was removed from the reservoir and each Transwell insert and replaced with prewarmed fresh culture medium. Transepithelial electrical resistance (TEER) across the monolayer was measured using Millicell Epithelial Volt-Ohm measuring system (Millipore, USA). The Plate was returned to the incubator once the measurement was done. The TEER value was calculated according to the following equation:

TEER measurement (ohms) x Area of membrane ($cm^2$) = TEER value (ohm•$cm^2$)

TEER value should be greater than 42 ohm•$cm^2$, which indicates the well-qualified MDCKII-MDR1 monolayer.

**Bidirectional Permeability Assay Using MDCKII-MDR1 Cells**

MDCKII-MDR1 plates were washed twice with pre-warmed HBSS (10 mM HEPES, pH 7.4) and incubated at 37 °C for 30 min. Test and control compounds were prepared at 1 µM in HBSS containing 0.5% DMSO.

For **apical-to-basolateral (A→B)** transport, 125 µL of the compound solution was added to the apical (donor) side, and 50 µL was immediately sampled into 200 µL acetonitrile containing internal standards (100 nM ketoprofen, 200 nM labetalol, and 100 nM tolbutamide). The basolateral (receiver) compartment was filled with 235 µL of transport buffer.

For **basolateral-to-apical (B→A)** transport, 285 µL of compound was added to the basolateral side, and 50 µL was sampled immediately into acetonitrile + IS. The apical compartment was filled with 75 µL of transport buffer. Both directions were assayed in parallel.

The plates were incubated at 37°C for 2 hours. Final 50 µL samples were collected from donor and receiver compartments and quenched with 200 µL of acetonitrile + IS. Samples were vortexed (10 min), centrifuged (30 min, 4,000 rpm, 4 °C), and the supernatants diluted with water before LC-MS/MS analysis.

**Lucifer Yellow Leakage Assay**

To assess monolayer integrity, Lucifer Yellow was added at 100 µM (in HBSS) to the apical side (100 µL), and 300 µL HBSS was added to the basolateral wells. After 30 min of incubation at 37°C, 80 µL from both compartments was collected, and fluorescence was measured (Ex: 480 nm, Em: 530 nm).

**Data Analysis**

*Apparent permeability ($P_{app}$)* can be calculated for drug transport assays using the following equation:

$$P_{app} = \frac{V_A}{area \times time} \times \frac{[drug]_{acceptor}}{[drug]_{initial,\ donor}}$$

Where: $P_{app}$ is apparent permeability (cm/s x $10^{-6}$). $V_A$ is the volume (in mL) in the acceptor well Area is the surface area of the membrane (0.143 $cm^2$ for Transwell-96 Well Permeable Supports) time is the total transport time in seconds.

*Efflux ratio* can be determined using the following equation:

$$efflux\ ratio = \frac{P_{app\ (B-A)}}{P_{app\ (A-B)}}$$

Where $P_{app(B-A)}$ indicates the apparent permeability coefficient in basolateral to apical direction, and $P_{app(A-B)}$ indicates the apparent permeability coefficient in apical to basolateral direction.

Mass balance (% recovery) can be determined using the following equation:

$$recovery\,\% = \frac{[drug]_{acceptor} \times V_A + [drug]_{donor} \times V_D}{[drug]_{initial,donor} \times V_D} \times 100$$

Where $V_A$ is the volume (in mL) in the acceptor well (0.235 mL for A→B flux, and 0.075 mL for B→A), $V_D$ is the volume (in mL) in the donor well (0.075 mL for A→B flux, and 0.235 mL for B→A).

Lucifer yellow leakage of monolayer can be calculated using the following equation:

$$LY\ leakage = \left(\frac{I_{acceptor} \times 0.3}{I_{acceptor} \times 0.3 + I_{donor} \times 0.1}\right) \times 100\%$$

Where $I_{acceptor}$ is the fluorescence intensity in the acceptor well (0.3 mL), and $I_{donor}$ is the fluorescence intensity in the donor well (0.1 mL) and expressed as % leakage. Lucifer yellow percentage amount transported values should be less than 1.5 %. However, if the $P_{app}$ determined in that transwell is qualitatively similar to that determined in the replicate transwells, based upon the scientific judgement of the responsible scientist, then the monolayer is considered acceptable.

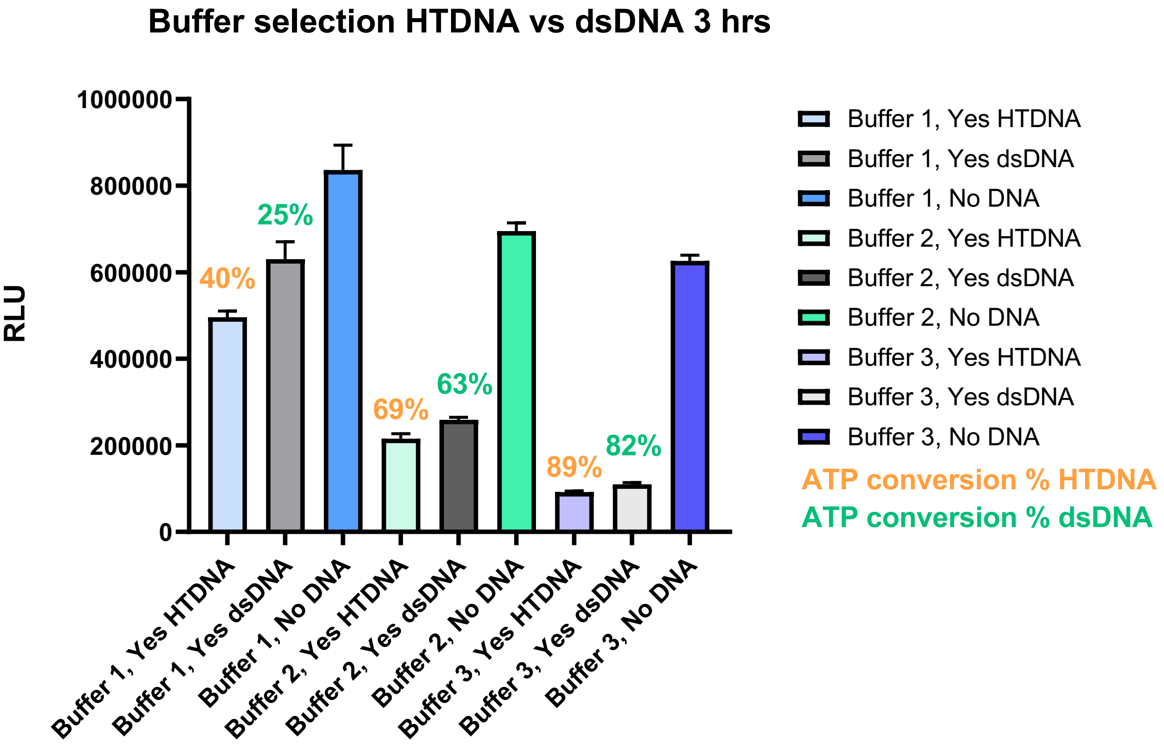


**Plot S1.** Buffer optimization using HTDNA (5 µg/ml) vs. dsDNA (25 nM) after 3 h incubation: Summary of ATP conversion efficiency (raw data in RLU).

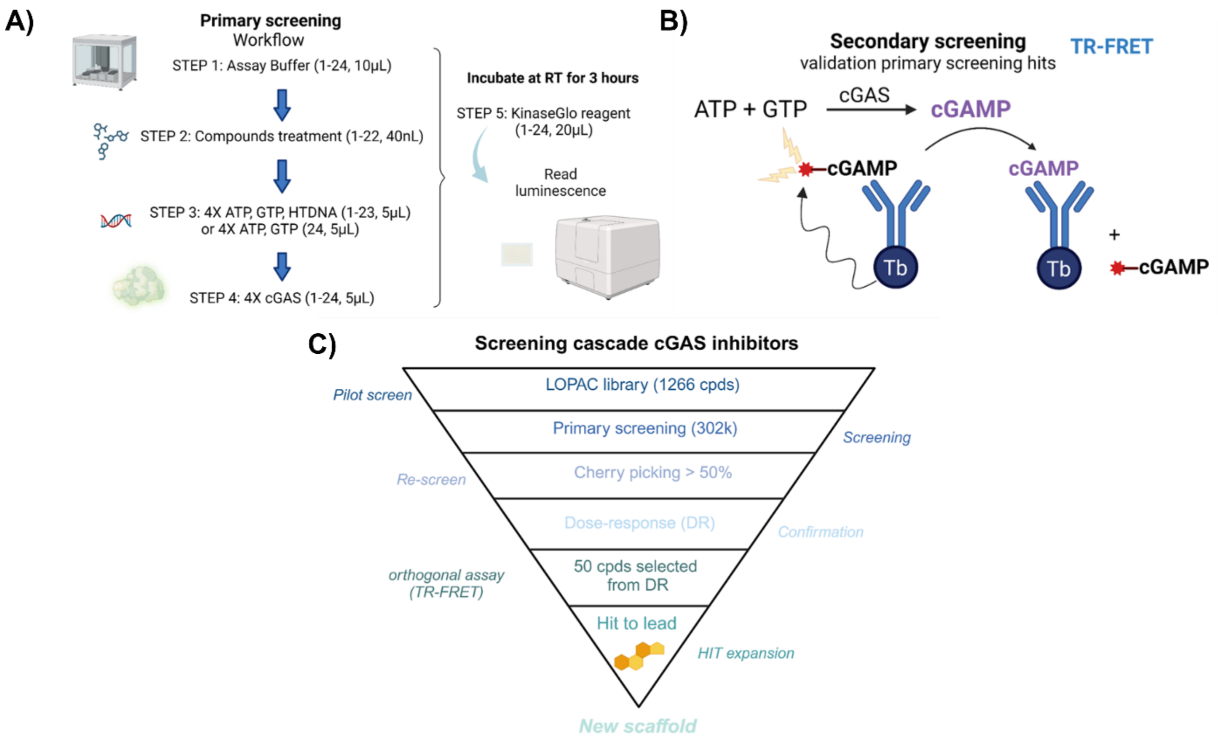


**Supplementary Figure S1A.** Primary screening workflow used in our HTS for the identification of human cGAS inhibitors. Numbers in parentheses indicate the column position in the 384-well plate and the volume dispensed in μL. **S1B.** Secondary screening. Direct method to measure cGAMP and enzyme activity by TR-FRET assay (adapted from Bellbrook protocol). **S1C**. Screening cascade cGAS inhibitors.

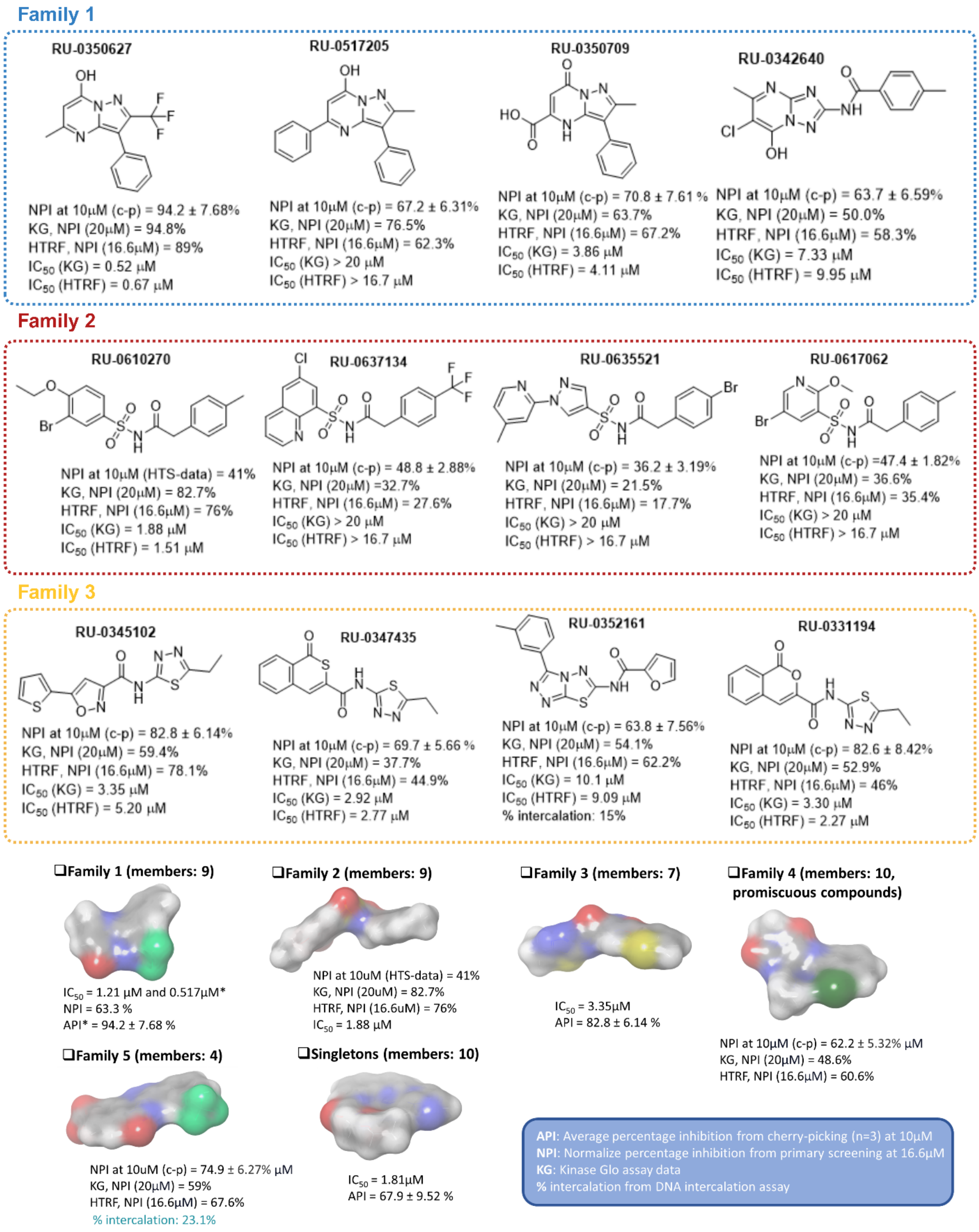


**Supplementary Figure S2.** Representative and selected hits grouped by chemical families: molecular surface and in vitro properties. Molecular surfaces were generated for selected hits using Schrödinger software. Atoms are color-coded as follows: Hydrogen (H), white; Carbon (C), gray; Oxygen (O), network; Nitrogen (N), blue; Sulfur (S), yellow; Chlorine (Cl), dark green; Fluorine (F), light green.

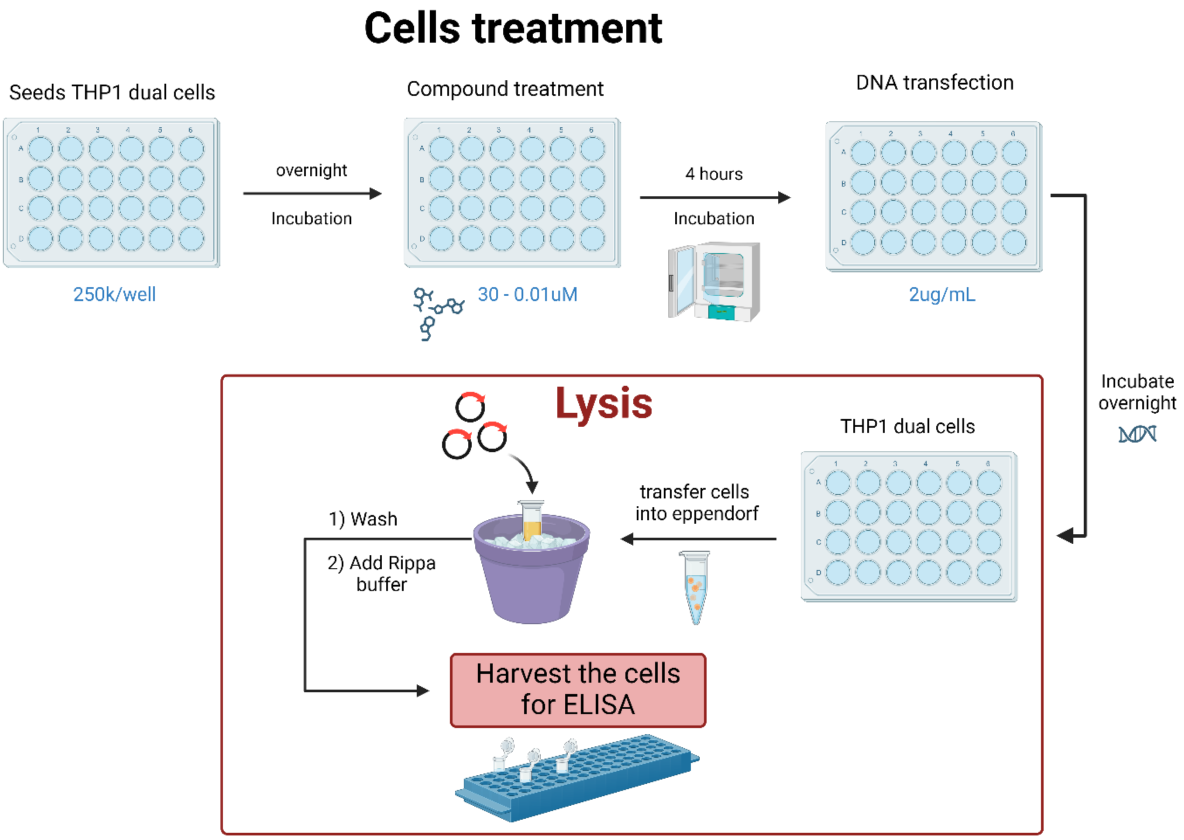


**Supplementary Figure S3.** Sample preparation for cGAMP ELISA in THP1 dual cells.

**Supplementary Table S1.** Activity data from primary screening and cherry picking against Kinase-Glo, TR-FRET, and cytotoxicity assays. Results expressed as normalized percentage of inhibition (NPI, %).

| | | **Family** | **Kinase Glo** | **Confirmation Kinase Glo** | **Secondary readout TR-FRET** | **Cytotoxicity** |
|---|---|---|---|---|---|---|
| **Entry** | **RU ID** | **(F)** | **(NPI, %) 10 µM** | **(NPI, %) 20 µM** | **(NPI, %) 16.6 µM** | **CellGlo MRC5 NPCD** (%)** |
| 1 | **RU-0350627 (JA-1)** | *F1* | 63.3 | 94.8 | 89 | -2.65 |
| 2 | **RU-0350709** | *F1* | 70.8 | 63.7 | 67.2 | -0.18 |
| 3 | **RU-0517205** | *F1* | 67.2 | 76.5 | 62.3 | -19.16 |
| 4 | **RU-0350924** | *F1* | 50 | 75.6 | 66.5 | - |
| 5 | **RU-0342640** | *F1* | 63.7 | 50 | 58.3 | -19.78 |
| 6 | **RU-0342684** | *F1* | 70.5 | 62.1 | 61.7 | -35.39 |
| 7 | **RU-0330955** | *F1* | 38.3 | 36 | 23.4 | 3.88 |
| 8 | **RU-0624225** | *F1* | 53.6 | 42.7 | 41.9 | -13.31 |
| 9 | **RU-0335783** | *F1* | 32.7 | 34.8 | 7.34 | -0.24 |
| 10 | **RU-0610270 (JA-2)** | *F2* | 41 | 82.7 | 76 | - |
| 11 | **RU-0635513** | *F2* | 58 | 58.9 | 51.9 | - |
| 12 | **RU-0615385** | *F2* | 51.4 | 52 | 50.5 | - |
| 13 | **RU-0618980** | *F2* | 45.8 | 58.8 | 54.9 | - |
| 14 | **RU-0637134** | *F2* | 48.8 | 32.7 | 27.6 | 5.28 |
| 15 | **RU-0617062** | *F2* | 47.4 | 36.6 | 35.4 | -6.10 |
| 16 | **RU-0619159** | *F2* | 32.9 | 32.6 | 41 | - |
| 17 | **RU-0619161** | *F2* | 29 | 42.7 | 38.6 | - |
| 18 | **RU-0635521** | *F2* | 36.2 | 21.5 | 17.7 | -0.17 |
| 19 | **RU-0345102** | *F3* | 82.8 | 59.4 | 78.1 | -15.35 |
| 20 | **RU-0331194** | *F3* | 82.6 | 52.9 | 46 | -4.32 |
| 21 | **RU-0347435** | *F3* | 69.7 | 37.7 | 44.9 | -1.32 |
| 22 | **RU-0352161** | *F3* | 63.8 | 54.1 | 62.2 | -17.54 |
| 23 | **RU-0359021** | *F3* | 30.9 | 35.9 | 36.4 | 97.87 |
| 24 | **RU-0612339** | *F3* | 28.7 | 37.6 | 28 | - |
| 25 | **RU-0418316** | *F3* | 35.6 | 73 | 65 | - |
| 26 | **RU-0341285** | *F4* | 80.6 | 75.1 | 70.4 | -2.17 |

| | | | | | | |
|---|---|---|---|---|---|---|
| 27 | **RU-0341329** | *F4* | 81.6 | 66.6 | 50.6 | -6.96 |
| 28 | **RU-0341241** | *F4* | 61.6 | 66.3 | 54.7 | 5.46 |
| 29 | **RU-0341373** | *F4* | 57.9 | 53.2 | 45.6 | 4.80 |
| 30 | **RU-0341417** | *F4* | 73.9 | 60.4 | 28.4 | 5.04 |
| 31 | **RU-0334691** | *F4* | 64.1 | 44.9 | 6.93 | -7.44 |
| 32 | **RU-0341197** | *F4* | 59.2 | 66 | 43.1 | 18.33 |
| 33 | **RU-0422690** | *F4* | 44.9 | 39 | 36.8 | -13.95 |
| 34 | **RU-0331765** | *F4* | 43.9 | 39.2 | 25.6 | 11.81 |
| 35 | **RU-0154302** | *F4* | 29.5 | 75.3 | 51.1 | - |
| 36 | **RU-0430362** | *F5* | 74.9 | 59 | 67.6 | -14.62 |
| 37 | **RU-0332110** | *F5* | 45.3 | 11.6 | 24.5 | 5.94 |
| 38 | **RU-0333746** | *F5* | 43 | 48.6 | 34.7 | -3.39 |
| 39 | **RU-0350434** | *F5* | 62.2 | 48.6 | 60.6 | -8.24 |
| 40 | **RU-0423129** | *F6* | 55.1 | 37.5 | 51.7 | 100.64 |
| 41 | **RU-0330756** | *F6* | 42.8 | 42.1 | 35.6 | 96.99 |
| 42 | **RU-0519290** | *F6* | 67.9 | 61.4 | 43.9 | 1.32 |
| 43 | **RU-0556267** | *F6* | 30.4 | 32.1 | 40.5 | - |
| 44 | **RU-0651275** | *F6* | 32.8 | 15.8 | 25.2 | 1.89 |
| 45 | **RU-0609020** | *F6* | 28.4 | 91.4 | 80.2 | - |
| 46 | **RU-0323352** | *F6* | 34.4 | 29.9 | 30.6 | 2.52 |
| 47 | **RU-0331411** | *F6* | 28.1 | 32.1 | 38.7 | -1.93 |
| 48 | **RU-0485429** | *F6* | 19.4 | 21.4 | 18.4 | -3.24 |
| 49 | **RU-0378298** | *F6* | 49.6 | 88.3 | 47.7 | - |
| 50 | **RU-0328460** | *F6* | 15.52 | - | 18.58 | 1.83 |
| 51 | **RU-0334044** | *F6* | 5.31 | - | 16.0 | 8.22 |

NPCD**: Normalized percentage cells die.

**Supplementary Table S1 (continued).** SMILES representations of selected compounds from the screening campaign.

| | RU ID | SMILE |
|---|---|---|
| 1 | **RU-0350627 (JA-1)** | CC1=NC2=C(C3=CC=CC=C3)C(C(F)(F)F)=NN2C(O)=C1 |

| | | |
|---|---|---|
| 2 | **RU-0350709** | CC1=NN2C(=C1C1=CC=CC=C1)NC(C(O)=O)=CC2=O |
| 3 | **RU-0517205** | OC1=CC(C2=CC=CC=C2)=NC2=C(C3=CC=CC=C3)C(C)=NN12 |
| 4 | **RU-0350924** | CCOC(C1=CNC2=C(C(C)=NN2C1=O)C3=CC=CC=C3)=O |
| 5 | **RU-0342640** | CC1=CC=C(C(=O)NC2=NN3C(=N2)N=C(C)C(Cl)=C3O)C=C1 |
| 6 | **RU-0342684** | CC1=NC2=NC(NC(=O)C3=CC=CO3)=NN2C(O)=C1Cl |
| 7 | **RU-0330955** | CSC1=NN2C(=N1)NC(C1=CC=CC=C1)=CC2=O |
| 8 | **RU-0624225** | FC1=CC=C([C@H]2[C@H](NC(=O)C3=NC=CN4C3=NC=C4)C2)C=C1 |
| 9 | **RU-0335783** | OC1=C2CCCCC2=NC2=NC(C3=CC=CC=C3)=NN12 |
| 10 | **RU-0610270 (JA-2)** | BrC1=C(C=CC(S(=O)(NC(CC2=CC=C(C=C2)C)=O)=O)=C1)OCC |
| 11 | **RU-0635513** | BrC1=C(C=CC(S(=O)(NC(CC2=CC=C(C=C2)CC)=O)=O)=C1)OCC |
| 12 | **RU-0615385** | BrC1=CC(S(=O)(NC(CC2=CC=C(C=C2)Cl)=O)=O)=C(C=C1)Cl |
| 13 | **RU-0618980** | O=S(NC(CC1=CC=C(C=C1)C)=O)(C2=C(C=C(C=C2)C)C(NCC3=CC=C (C=C3)C)=O)=O |
| 14 | **RU-0637134** | ClC1=CC2=CC=CN=C2C(S(=O)(=O)NC(=O)CC2=CC=C(C(F)(F)F)C=C2)=C1 |
| 15 | **RU-0617062** | BrC1=CN=C(OC)C(S(=O)(=O)NC(=O)CC2=CC=C(C)C=C2)=C1 |
| 16 | **RU-0619159** | BrC1=CC=C(C=C1)CC(NS(=O)(C2=C(N=CC(Cl)=C2)OC)=O)=O |
| 17 | **RU-0619161** | BrC1=CC=C(C=C1)CC(NS(=O)(C2=C(C=C(C(OC)=C2)OC)Cl)=O)=O |
| 18 | **RU-0635521** | BrC1=CC=C(CC(=O)NS(=O)(=O)C2=CN(C3=NC=CC(C)=C3)N=C2)C=C1 |
| 19 | **RU-0345102** | CCC1=NN=C(NC(=O)C2=NOC(C3=CC=CS3)=C2)S1 |
| 20 | **RU-0331194** | CCC1=NN=C(NC(=O)C2=CC3=CC=CC=C3C(=O)O2)S1 |
| 21 | **RU-0347435** | CCC1=NN=C(NC(=O)C2=CC3=C(C=CC=C3)C(=O)S2)S1 |
| 22 | **RU-0352161** | CC1=CC=CC(C2=NN=C3SC(NC(=O)C4=CC=CO4)=NN23)=C1 |
| 23 | **RU-0359021** | COC1=CC(OC)=C(CN2CC(=O)C(C3=NC4=C(S3)C=CC=C4)=C2N)C=C1 |
| 24 | **RU-0612339** | O=C(CN1C(C2=C(C=CC=C2)N=C1)=O)NC3=NN=C(C4=C(C=CC(F)=C4)C)S3 |
| 25 | **RU-0418316** | CSC1=NN=C(S1)NC(C2=NOC(C3=CC=CS3)=C2)=O |
| 26 | **RU-0341285** | CN1C(=O)NC2=NN=C(/C=C/C3=CC=CC=C3)N=C2C1=O |
| 27 | **RU-0341329** | CN1C(=O)NC2=NN=C(C3=CC=CC=N3)N=C2C1=O |
| 28 | **RU-0341241** | COC1=CC=C(C2=NN=C3NC(=O)N(C)C(=O)C3=N2)C=C1OC |
| 29 | **RU-0341373** | CCN1N=C(C2=C(Cl)C=CC=C2)N=C2C(=O)N(C)C(=O)N=C12 |
| 30 | **RU-0341417** | CN1C(=O)NC2=NN=C(C3=CC=C(C)C=C3)N=C2C1=O |
| 31 | **RU-0334691** | CCC(=O)NC1=CC2=C(C=C1C)N=C1C(=O)N(C)C(=O)NC1=N2 |
| 32 | **RU-0341197** | CN1C(=O)NC2=NN=C(C3=CC=C(F)C=C3)N=C2C1=O |
| 33 | **RU-0422690** | OC(=O)CNC(=O)C1=C(O)C2=CC=CC=C2C(Cl)=N1 |
| 34 | **RU-0331765** | CCCCSC1=NN2C(=O)C3=CC=CC=C3N=C2C(=O)N1 |
| 35 | **RU-0154302** | O=C(NC1=NC=CC=N1)C2=C(C3=CC=CC(C(F)(F)F)=C3N=C2)O |
| 36 | **RU-0430362** | CC1=NN2C(=C1C#N)NC(=O)C1=CC=CN=C21 |
| 37 | **RU-0332110** | COC1=CC=C(C2=CC(=O)N3N=C(SC)N=C3N2)C=C1OC |
| 38 | **RU-0333746** | OC(=O)C1=CC=C2C(=O)NC3=C(C4=CC(C(F)(F)F)=CC=C4)N=NN3C2=C1 |

| | | |
|---|---|---|
| 39 | **RU-0350434** | CC1=C(C(O)=O)C=NC2=NC3=CC=CC=C3N12 |
| 40 | **RU-0423129** | [H][C@]12NC3=C(C=CC=C3)[C@@]1([C@]13C[C@]45SS[C@](CO)(N(C)C4=O)C(=O)N5[C@@]1([H])NC1=C3C=CC=C1)C[C@]13SS[C@](CO)(N(C)C1=O)C(=O)N23 |
| 41 | **RU-0330756** | CCCN1C(NC2=NC3=C(C=CC=C3)N2C)=NC2=C1C=CC=C2 |
| 42 | **RU-0519290** | O=C(NC1=CC=CC2=CC=C(N3C=NC=C3)N=C12)COCC |
| 43 | **RU-0556267** | CC(C=C1)=CN=C1CN(CCNCC2)CCN2CC3=NC=C(C)C=C3 |
| 44 | **RU-0651275** | S(=O)(=O)(NC)C1=CC=C(NC(=O)C2=CSC3=C2C(=O)NC=N3)C=C1 |
| 45 | **RU-0609020** | O=C(C1=NN(C2=C1C=CC=C2)C)NC3=CC(C)=C(C=C3)NC(C4=C5N=CC(CCN)=CN5N=C4)=O |
| 46 | **RU-0323352** | OC(=O)C1=CC(C2=CC=CS2)=NC2=CC=CC=C12 |
| 47 | **RU-0331411** | CSC1=C2C3=C(SC2=NC(N)=N1)C(O)=CC(=O)N3 |
| 48 | **RU-0485429** | S(C1=CC(C)=C(C#N)C2=NC3=C(N12)C=CC=C3)CC(=O)CC(=O)OCC |
| 49 | **RU-0378298** | CC(OC1=C(C=C2C(C)=CC(OC2=C1)=O)Cl)=O |
| 50 | **RU-0328460** | CC1(C)CC2=C(SC(NC(=O)C3=CC(C4=CC=CC=C4)=NC4=CC=CC=C34)=N2)C(=O)C1 |
| 51 | **RU-0334044** | COC1=CC=C2N=CC3=C(SC(C(O)=O)=C3C3=CC=CC=C3)C2=C1 |

**Supplementary Table S2.** Confirmation of selected hits through LC-MS.

| RU-ID | MW (g/mol) | Found? | Purity | Family | $IC_{50}$ (μM) Kinase Glo | $IC_{50}$ (μM) TR-FRET |
|---|---|---|---|---|---|---|
| **RU-0350627 (JA-1)** | 293.248 | yes | 100 | *F1* | 0.517 | 0.96 |
| **RU-0350709** | 269.26 | yes | 100 | *F1* | 3.86 | 4.19 |
| **RU-0342640** | 317.736 | yes | 100 | *F1* | 7.33 | 10.18 |
| **RU-0342684** | 293.67 | yes | 100 | *F1* | 7.39 | 8.04 |
| **RU-0610270 (JA-2)** | 412.305 | yes | 100 | *F2* | 1.88 | 1.55 |
| **RU-0619159** | 419.684 | yes | 100 | *F2* | > 20 | > 16.6 |
| **RU-0635513** | 426.332 | yes | 100 | *F2* | 11.3 | > 16.6 |
| **RU-0615385** | 423.115 | yes | 100 | *F2* | 13.7 | > 16.6 |
| **RU-0333746** | 374.278 | yes | 93 | *F5* | 2.05 | > 16.6 |
| **RU-0430362** | 225.211 | yes | 100 | *F5* | 4.26 | 4.34 |
| **RU-0350434** | 227.223 | yes | 100 | *F5* | 6.59 | 14.22 |
| **RU-0519290** | 296.33 | yes | 100 | *F6* | 11.18 | > 16.6 |

**Supplementary Table S3.** DNA Intercalation Assay: Intercalator at 33.3 μM, Expressed as % Intercalation

| RU-ID | % Intercalation |
|---|---|
| **RU-0331765** | 10.7 |
| **RU-0332120** | 18.1 |
| **RU-0334691** | 5.68 |
| **RU-0342684** | 14.4 |
| **RU-0352161** | 13.1 |
| **RU-0422690** | 4.02 |
| **RU-0430362** | 23.1 |
| **RU-0485429** | 44.2 |
| **RU-0519290** | 5.39 |
| **RU-0651275** | 19.5 |

**Supplementary Table S4.** Library information screening.

| **Category** | **Parameter** | **Description** |
|---|---|---|
| Library | Library size | 302,332 pure, low molecular weight compounds from the Rockefeller University Compound Library. |
| | Source | Chembridge, ChemDiv, Enamine, LifeChem, Selleck |

**Supplementary Table S5.** Library composition screen in Fisher Drug Discovery Resource Center at the Rockefeller University.

| **Number of Compounds** | **Vendor** |
|---|---|
| 50,240 | Chembridge |
| 99,912 | ChemDiv |
| 100,477 | Enamine |
| 50,240 | LifeChem |
| 1,867 | Selleck |

**Supplementary Table S6.** Activity data in Kinase Glo and TR-FRET assay from analogs of sulfonamide selected hit. Percentage inhibition at 12.5 μM and 10.4 μM respectively.

**Structure 1**

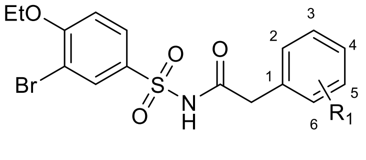


**Structure 2**

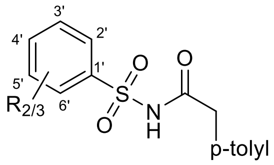


**Structure 3**

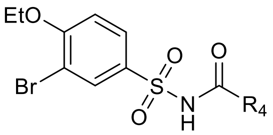


**Structure 4**

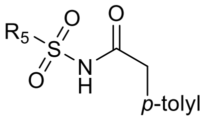

| | | Kinase Glo[1] | Kinase Glo[2] | TR-FRET[3] |
|---|---|---|---|---|
| **Cpd** | **Structure 1 $R_1$** | **% inhibition at 12.5 µM ± SD (n = 3)** | **% inhibition at 12.5 µM ± SD (n = 3)** | **% inhibition at 10.4 µM ± SD (n = 2)** |
| **1** | 4-Me | 74.45 ± 2.93 | 79.41 ± 4.40 | 96.92 ± 3.14 |
| **2** | 4-CF3 | 31.15 ± 3.52 | 39.97 ± 3.15 | 57.87 ± 8.15 |
| **3** | 4-F | 10.82 ± 2.28 | 14.22 ± 7.07 | 25.35 ± 3.90 |
| **4** | 4-Cl | 82.85 ± 4.81 | 68.47 ± 2.79 | 96.06 ± 4.79 |
| **5** | 4-Br | 82.08 ± 3.13 | 72.79 ± 1.94 | 102.53 ± 4.39 |
| **6** | 4-I | 97.85 ± 1.89 | 84.22 ± 1.48 | 98.85 ± 1.51 |
| **7** | 4-OMe | 23.26 ± 2.09 | 32.73 ± 0.60 | 58.02 ± 9.44 |
| **8** | 4-O$(CH_3)_2$ | 7.53 ± 1.61 | -1.03 ± 2.87 | 15.46 ± 5.07 |
| **9** | 4-OPh | 19.70 ± 4.66 | 15.29 ± 1.46 | 27.00 ± 4.14 |
| **10** | 4-SMe | 69.68 ± 4.20 | 63.90 ± 5.59 | 83.46 ± 2.93 |
| **11** | 4-$SCF_3$ | 6.65 ± 3.76 | 1.72 ± 0.93 | 12.35 ± 14.98 |
| **12** | 4-CN | 5.79 ± 2.02 | 3.75 ± 2.75 | 8.07 ± 10.27 |
| **13** | 2-Me | 2.75 ± 1.51 | 5.33 ± 2.60 | 12.18 ± 1.66 |
| **14** | 2-$CF_3$ | 0.06 ± 0.91 | -5.97 ± 1.38 | 5.18 ± 5.39 |
| **15** | 2-Cl | 3.79 ± 2.60 | -0.48 ± 0.50 | 10.15 ± 2.65 |
| **16** | 2-I | 1.24 ± 0.99 | -3.14 ± 1.42 | 2.77 ± 2.69 |
| **17** | 2-OMe | -1.35 ± 3.09 | -2.54 ± 2.26 | 9.82 ± 6.31 |
| **18** | 2-$OCF_3$ | 11.46 ± 3.15 | 12.33 ± 1.21 | 30.21 ± 2.32 |
| **19** | 6-F | 9.56 ± 2.37 | 2.73 ± 4.06 | 16.53 ± 6.26 |
| **20** | 6-$OCF_3$ | 1.62 ± 2.05 | -6.72 ± 0.97 | -0.95 ± 8.78 |
| **21** | 5-Me | 3.39 ± 0.40 | 1.73 ± 3.94 | 4.74 ± 15.19 |
| **22** | 5-$CH_2CH_2Cl$ | 2.51 ± 2.16 | 1.85 ± 2.28 | 13.68 ± 5.12 |
| **23** | 5-Cl | 5.41 ± 2.82 | 2.71 ± 1.35 | 19.38 ± 2.56 |
| **24** | 5-Br | 67.16 ± 6.57 | 3.77 ± 1.27 | 20.87 ± 3.49 |
| | **Structure 2 $R_{2/3}$** | | | |
| **25** | H | -0.27 ± 1.39 | -6.69 ± 1.69 | n.d. |
| **26** | $R_2$ = 4’-OEt | 5.24 ± 1.87 | 6.36 ± 2.23 | n.d. |
| **27** | $R_2$ = 4’-OMe | 0.35 ± 2.73 | -1.69 ± 0.71 | n.d. |
| **28** | $R_2$ = 4’-$OCF_3$ | 2.15 ± 2.27 | -5.59 ± 0.74 | n.d. |
| **29** | $R_2$ = 4’-$C(CH_3)_3$ | 4.76 ± 0.26 | -2.90 ± 0.46 | n.d. |
| **30** | $R_2$ = 4’-$CF_3$ | 49.07 ± 15.39 | -4.97 ± 1.86 | n.d. |
| **31** | $R_2$ = 5’-Br | 2.20 ± 1.15 | -0.60 ± 2.48 | n.d. |
| **32** | $R_2$ = 4’-Me | 22.06 ± 3.90 | 19.35 ± 0.49 | n.d. |

| | | | | |
|---|---|---|---|---|
| | $R_3$ = 5'-Br | | | |
| **33** | $R_2$ = 4'-Br<br>$R_3$ = 5'-F | 12.56 ± 3.72 | 14.73 ± 2.21 | n.d. |
| **34** | $R_2$ = 4'-Oet<br>$R_3$ = 5'-<br>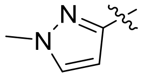 | 5.81 ± 1.65 | 2.06 ± 2.12 | 13.25 ± 8.70 |
| **35** | $R_2$ = 3'-Br<br>$R_3$ = 6'-F | 24.05 ± 5.72 | 26.76 ± 0.67 | n.d. |
| **36** | $R_2$ = 3'-Cl<br>$R_3$ = 6'-Cl | 35.39 ± 2.12 | 35.97 ± 2.08 | n.d. |
| **37** | $R_2$ = 3'-$CF_3$<br>$R_3$ = 6'-F | 19.66 ± 1.94 | 22.53 ± 1.21 | n.d. |
| **38** | $R_2$ = 2'-Me<br>$R_3$ = 3'-Cl | 16.53 ± 2.85 | 19.89 ± 0.61 | n.d. |
| | **Structure 3**<br>**$R_4$** | | | |
| **39** | 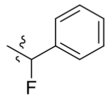 | 15.70 ± 2.85 | 20.99 ± 1.46 | 45.07 ± 8.13 |
| **40** | 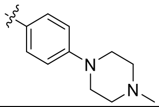 | -0.71 ± 0.99 | -4.95 ± 1.17 | 5.94 ± 0.49 |
| **41** | 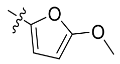 | -0.01 ± 1.52 | -5.96 ± 1.98 | 3.00 ± 3.97 |
| **42** | 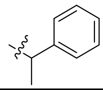 | 5.96 ± 4.80 | -3.48 ± 2.73 | -2.92 ± 3.79 |
| **43** | 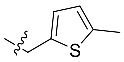 | 23.10 ± 2.37 | 40.40 ± 0.63 | n.d. |
| **44** | 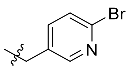 | 8.23 ± 2.02 | 19.87 ± 1.65 | n.d. |
| **45** | 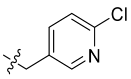 | 6.96 ± 3.06 | 13.68 ± 0.03 | n.d. |
| **46** | 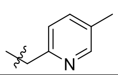 | 0.37 ± 1.84 | -2.21 ± 0.00 | n.d. |
| | **Structure 4**<br>**$R_5$** | | | |
| **47** | 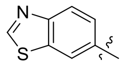 | 4.13 ± 2.17 | 6.37 ± 0.06 | n.d. |

| | | | | |
|---|---|---|---|---|
| **48** | 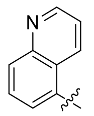 | 5.36 ± 1.07 | 5.57 ± 1.36 | n.d. |
| **49** | 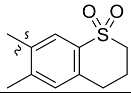 | 7.71 ± 4.19 | 4.90 ± 2.46 | n.d. |

Experimental condition[1]: 25nM enzyme, 37ºC.

Experimental condition[2]: 100nM enzyme, 25ºC.

Experimental condition[3]: 30 nM enzyme, 25ºC.

### $^1$H-NMR data and LC-MS data of sulfonamide derivatives

**Compound 1:** *N*-((3-bromo-4-ethoxyphenyl)sulfonyl)-2-(*p*-tolyl)acetamide:

$^1$H NMR (600 MHz, DMSO-d$_6$) δ 12.28 (s, 1H), 7.96 (d, *J* = 2.4 Hz, 1H), 7.84 (dd, *J* = 8.8, 2.4 Hz, 1H), 7.28 (d, *J* = 8.8 Hz, 1H), 7.08 (d, *J* = 7.7 Hz, 2H), 7.03 (d, *J* = 7.6 Hz, 2H), 4.22 (q, *J* = 6.9 Hz, 2H), 3.47 (s, 2H), 2.26 (s, 3H), 1.38 (t, *J* = 6.9 Hz, 3H).

MS (ESI) m/z calculated for $[C_{17}H_{18}BrNO_4S]^+$ ($[M+H]^+$): 412.01, observed: 412.2.

**Compound 3:** *N*-((3-bromo-4-ethoxyphenyl)sulfonyl)-2-(4-fluorophenyl)acetamide:

$^1$H NMR (600 MHz, DMSO-d$_6$) δ 12.33 (s, 1H), 7.96 (d, *J* = 2.3 Hz, 1H), 7.84 (dd, *J* = 8.8, 2.3 Hz, 1H), 7.28 (d, *J* = 8.8 Hz, 1H), 7.20 (dd, *J* = 8.3, 5.6 Hz, 2H), 7.10 (t, *J* = 8.3 Hz, 2H), 4.22 (q, *J* = 7.0 Hz, 2H), 3.53 (s, 2H), 1.38 (t, *J* = 7.0 Hz, 3H).

MS (ESI) m/z calculated for $[C_{16}H_{15}BrFNO_4S]^+$ ($[M+H]^+$): 415.99, observed: 416.2.

**Compound 4**: *N*-((3-bromo-4-ethoxyphenyl)sulfonyl)-2-(4-chlorophenyl)acetamide:

$^1$H NMR (600 MHz, DMSO-d$_6$) δ 12.34 (s, 1H), 7.9 – 7.94 (m, 1H), 7.87 – 7.83 (m, 1H), 7.34 (d, *J* = 8.1 Hz, 2H), 7.27 (d, *J* = 9.0 Hz, 1H), 7.19 (d, *J* = 8.1 Hz, 2H), 4.23 (t, *J* = 7.0 Hz, 2H), 3.55 (s, 2H), 1.38 (t, *J* = 6.9 Hz, 3H).

$^{13}$C NMR (150 MHz, DMSO-d$_6$): δ 170.0, 159.0, 133.5, 132.5, 132.1, 131.7, 129.7, 128.7, 128.6, 113.7, 111.1, 65.7, 41.8, 14.8.

LC-MS m/z calculated for $[C_{16}H_{15}BrClNO_4S]^+$ ($[M+H]^+$): 431.96, observed: 432.1.

**Compound 5:** *N*-((3-bromo-4-ethoxyphenyl)sulfonyl)-2-(4-bromophenyl)acetamide:

$^1$H NMR (600 MHz, DMSO-d$_6$) δ 7.97 (t, *J* = 2.3 Hz, 1H), 7.85 (dd, *J* = 8.8, 2.3 Hz, 1H), 7.47 (d, *J* = 8.0 Hz, 2H), 7.28 (d, *J* = 8.8 Hz, 1H), 7.12 (d, *J* = 7.0 Hz, 2H), 3.54 (s, 2H), 1.38 (t, *J* = 7.0 Hz, 3H).

MS (ESI) m/z calculated for $[C_{16}H_{15}Br_2NO_4S]^+$ ($[M+H]^+$): 475.91, observed: 416.1.

**Compound 6**: *N*-((3-bromo-4-ethoxyphenyl)sulfonyl)-2-(4-iodophenyl)acetamide:

$^1$H NMR (600 MHz, DMSO-d$_6$): δ 12.34 (s, 1H), 7.97 (d, *J* = 2.3, Hz, 1H), 7.85 (dd, *J* = 8.8, 2.3 Hz, 1H), 7.64 (d, *J* = 8.2 Hz, 2H), 7.28 (d, *J* = 8.8 Hz, 2H), 6.97 (d, *J* = 8.2 Hz, 2H), 4.23 (q, *J* = 7.0 Hz, 2H), 3.52

(s, 2H), 1.39 (t, *J* = 7.0 Hz, 2H).

$^{13}$C NMR (150 MHz, DMSO-$d_6$): δ 169.7, 159.2, 137.5, 134.1, 132.5, 132.2, 131.9, 129.8, 113.7, 111.1, 93.4, 65.8, 41.9, 14.8.

MS (ESI) m/z calculated for $[C_{16}H_{15}BrINO_4S]^+$ ($[M+H]^+$): 523.8950, observed: 524.1.

**Compound 7:** N-((3-bromo-4-ethoxyphenyl)sulfonyl)-2-(4-methoxyphenyl)acetamide:

$^1$H NMR (600 MHz, DMSO-$d_6$) δ 12.26 (s, 1H), 7.97 (d, *J* = 2.3 Hz, 1H), 7.85 (dd, *J* = 8.8, 2.3 Hz, 1H), 7.28 (d, *J* = 8.8 Hz, 1H), 7.07 (d, *J* = 8.6 Hz, 2H), 6.84 (d, *J* = 8.6 Hz, 2H), 4.22 (q, *J* = 7.0 Hz, 2H), 3.72 (s, 3H), 3.46 (s, 2H), 1.38 (t, *J* = 6.9 Hz, 3H).

MS (ESI) m/z calculated for $[C_{17}H_{18}BrNO_5S]^+$ ($[M+H]^+$): 428.01, observed: 428.2.

**Compound 8:** *N*-((3-bromo-4-ethoxyphenyl)sulfonyl)-2-(4-isopropoxyphenyl)acetamide:

$^1$H NMR (600 MHz, DMSO-$d_6$): δ 12.26 (s, 1H), 7.98 (d, *J* = 2.2, Hz, 1H), 7.85 (dd, *J* = 8.9, 2.2 Hz, 1H), 7.28 (d, *J* = 8.9, Hz, 1H), 7.04 (d, *J* = 8.5 Hz, 2H), 6.81 (d, *J* = 8.5 Hz, 2H), 4.55 (h, *J* = 6.0 Hz, 1H), 4.22 (q, *J* = 7.0 Hz, 2H), 3.45 (s, 2H), 1.38 (t, *J* = 7.0 Hz, 3H), 1.24 (d, *J* = 6.0 Hz, 6H).

MS (ESI) m/z calculated for $[C_{19}H_{22}BrNO_5S]^+$ ($[M+H]^+$): 456.0402, observed: 456.2.

**Compound 9:** *N*-((3-bromo-4-ethoxyphenyl)sulfonyl)-2-(4-phenoxyphenyl)acetamide:

$^1$H NMR (600 MHz, DMSO-$d_6$): δ 12.32 (s, 1H), 7.99 (d, *J* = 2.2, Hz, 1H), 7.87 (dd, *J* = 9.0, 2.2 Hz, 1H), 7.40 – 7.37 (m, 2H), 7.29 (d, *J* = 9.0 Hz, 1H), 7.17 (d, *J* = 8.5 Hz, 1H), 7.14 (t, *J* = 7.4 Hz, 1H), 7.00–6.97 (m, 2H), 6.92 (d, *J* = 8.5 Hz, 1H), 4.22 (q, *J* = 7.0 Hz, 2H), 3.54 (s, 2H), 1.38 (t, *J* = 7.0 Hz, 3H).

MS (ESI) m/z calculated for $[C_{22}H_{20}BrNO_5S]^+$ ($[M+H]^+$): 490.0246, observed: 490.2.

**Compound 10:** *N*-((3-bromo-4-ethoxyphenyl)sulfonyl)-2-(4-(methylthio)phenyl)acetamide:

$^1$H NMR (600 MHz, DMSO-$d_6$): δ 12.30 (s, 1H), 7.97 (d, *J* = 2.2, Hz, 1H), 7.85 (dd, *J* = 8.9, 2.2 Hz, 1H), 7.28 (d, *J* = 8.9 Hz, 1H), 7.18 (d, *J* = 8.2 Hz, 2H), 7.12 (d, *J* = 8.2, Hz, 2H), 4.22 (q, *J* = 7.0 Hz, 2H), 3.50 (s, 2H), 2.45 (s, 3H), 1.38 (t, *J* = 7.0 Hz, 2H).

$^{13}$C NMR (150 MHz, DMSO-$d_6$): δ 159.1, 137.1, 132.6, 132.0, 130.9, 130.3, 129.8, 126.5, 113.7, 111.1, 65.8, 41.9, 15.2,14.8.

MS (ESI) m/z calculated for $[C_{17}H_{18}BrNO_4S_2]^+$ ($[M+H]^+$): 443.9861, observed: 444.2.

**Compound 11:** *N*-((3-bromo-4-ethoxyphenyl)sulfonyl)-2-(4-((trifluoromethyl)thio)phenyl)ace- tamide:

$^1$H NMR (600 MHz, DMSO-$d_6$): δ 12.40 (s, 1H), 7.98 (d, *J* = 2.2, Hz, 1H), 7.86 (dd, *J* = 8.8, 2.2 Hz, 1H), 7.64 (d, *J* = 8.0 Hz, 2H), 7.33 (d, *J* = 8.0 Hz, 2H), 7.29 (d, *J* = 8.8, Hz, 1H), 4.22 (q, *J* = 7.0 Hz, 2H), 3.66 (s, 2H), 1.38 (t, *J* = 7.0 Hz, 2H).

MS (ESI) m/z calculated for $[C_{17}H_{15}BrF_3NO_4S_2]^+$ ($[M+H]^+$): 497.9578, observed: 498.1.

**Compound 12:** *N*-((3-bromo-4-ethoxyphenyl)sulfonyl)-2-(4-cyanophenyl)acetamide:

$^1H$ NMR (600 MHz, DMSO-d$_6$) δ 12.41 (s, 1H), 7.97 (d, *J* = 2.3 Hz, 1H), 7.86 (dd, *J* = 8.8, 2.3 Hz, 1H), 7.76 (d, *J* = 8.0 Hz, 2H), 7.37 (d, *J* = 8.0 Hz, 2H), 7.29 (d, *J* = 8.8 Hz, 1H), 4.23 (q, *J* = 6.9 Hz, 2H), 3.69 (s, 2H), 1.39 (t, *J* = 6.9 Hz, 3H).

MS (ESI) m/z calculated for $[C_{17}H_{15}BrN_2O_4S]^+$ ($[M+H]^+$): 422.99, observed: 423.1.

**<u>Compound 13</u>:** *N*-((3-bromo-4-ethoxyphenyl)sulfonyl)-2-(o-tolyl)acetamide:

$^1H$ NMR (600 MHz, DMSO-d$_6$): δ 12.33 (s, 1H), 8.0 (d, *J* = 2.1, Hz, 1H), 7.88 (dd, *J* = 8.8, 2.1 Hz, 1H), 7.31 (d, *J* = 8.8 Hz, 1H), 7.61 – 7.11 (m, 2H), 7.10 – 7.06 (m, 2H), 4.23 (q, *J* = 7.0 Hz, 2H), 3.59 (s, 2H), 2.05 (s, 3H), 1.39 (t, *J* = 7.0 Hz, 2H).

MS (ESI) m/z calculated for $[C_{17}H_{18}BrNO_4S]^+$ ($[M+H]^+$): 412.0140, observed: 412.2.

**<u>Compound 14</u>:** *N*-((3-bromo-4-ethoxyphenyl)sulfonyl)-2-(2-(trifluoromethyl)phenyl)acetami- de:

$^1H$ NMR (600 MHz, DMSO-d$_6$) δ 7.98 (d, *J* = 2.3 Hz, 1H), 7.86 (dd, *J* = 8.8, 2.3 Hz, 1H), 7.65 (d, *J* = 7.9 Hz, 1H), 7.60 (t, *J* = 7.6 Hz, 1H), 7.47 (t, *J* = 7.7 Hz, 1H), 7.40 (d, *J* = 7.7 Hz, 1H), 7.30 (d, *J* = 8.8 Hz, 1H), 4.22 (q, *J* = 6.9 Hz, 2H), 3.84 (s, 2H), 1.38 (t, *J* = 6.9 Hz, 3H).

MS (ESI) m/z calculated for $[C_{17}H_{15}BrF_3NO_4S]^+$ ($[M+H]^+$): 465.9857, observed: 466.2

**<u>Compound 16</u>:** *N*-((3-bromo-4-ethoxyphenyl)sulfonyl)-2-(2-iodophenyl)acetamide:

$^1H$ NMR (600 MHz, DMSO-d$_6$) δ 12.42 (s, 1H), 8.01 (d, *J* = 2.3 Hz, 1H), 7.88 (dd, *J* = 8.0, 2.3 Hz, 1H), 7.78 (d, *J* = 8.0 Hz, 1H), 7.31 (q, *J* = 8.1 Hz, 2H), 7.25 (dd, *J* = 7.5, 1.7 Hz, 1H), 7.02 – 6.98 (m, 1H), 4.23 (q, *J* = 6.9 Hz, 2H), 3.75 (s, 2H), 1.39 (t, *J* = 6.9 Hz, 3H).

MS (ESI) m/z calculated for $[C_{16}H_{15}BrINO_4S]^+$ ($[M+H]^+$): 523.89, observed: 524.1.

**<u>Compound 19</u>:** *N*-((3-bromo-4-ethoxyphenyl)sulfonyl)-2-(2-fluorophenyl)acetamide:

$^1H$ NMR (600 MHz, DMSO-d$_6$): δ 12.39 (s, 1H), 8.0 (d, *J* = 2.0, Hz, 1H), 7.87 (dd, *J* = 8.8, 2.0 Hz, 1H), 7.32 – 7.28 (m, 1H), 7.30 (d, *J* = 8.8 Hz, 1H), 7.24 (t, *J* = 7.6 Hz, 1H), 7.15 – 7.11 (m, 2H), 4.23 (q, *J* = 7.0 Hz, 2H), 3.64 (s, 2H), 1.39 (t, *J* = 7.0 Hz, 2H).

MS (ESI) m/z calculated for $[C_{16}H_{15}BrFNO_4S]^+$ ($[M+H]^+$): 415.9889, observed: 416.2.

**<u>Compound 20</u>:** *N*-((3-bromo-4-ethoxyphenyl)sulfonyl)-2-(2(trifluoromethoxy)phenyl)aceta-mide:

$^1H$ NMR (600 MHz, DMSO-d$_6$): δ 12.41 (s, 1H), 7.99 (d, *J* = 2.2, Hz, 1H), 7.87 (dd, *J* = 8.6, 2.2 Hz, 1H), 7.39 (ddd, *J* = 7.6, 7.4, 2.2 Hz, 1H), 7.35 (dd, *J* = 6.5, 2.2 Hz, 1H), 7.32 (d, *J* = 7.4, Hz, 1H), 7.30 (d, *J* = 8.6 Hz, 1H), 7.28 (d, *J* = 7.4, Hz, 1H), 4.22 (q, *J* = 7.0 Hz, 2H), 3.70 (s, 2H), 1.39 (t, *J* = 7.0 Hz, 2H).

MS (ESI) m/z calculated for $[C_{17}H_{15}BrF_3NO_5S]^+$ ($[M+H]^+$): 481.98, observed: 482.2.

**<u>Compound 21</u>:** *N*-((3-bromo-4-ethoxyphenyl)sulfonyl)-2-(*m*-tolyl)acetamide:

$^1H$ NMR (600 MHz, DMSO-d$_6$) δ 12.3 (s, 1H), 7.98 (d, *J* = 2.2 Hz, 1H), 7.86 (dd, *J* = 8.8, 2.2 Hz, 1H), 7.29 (d, *J* = 8.8 Hz, 1H), 7.16 (t, *J* = 7.5 Hz, 1H), 7.04 (d, *J* = 7.6 Hz, 1H), 6.94 (d, *J* = 7.5 Hz, 1H), 6.93 (d, *J* = 7.6 Hz, 1H), 4.23 (q, *J* = 6.9 Hz, 2H), 3.50 (s, 2H), 2.25 (s, 3H), 1.39 (t, *J* = 6.9 Hz, 3H).

MS (ESI) m/z calculated for $[C_{17}H_{18}BrNO_4S]^+$ ($[M+H]^+$): 412.01, 412.2.

**Compound 22:** *N*-((3-bromo-4-ethoxyphenyl)sulfonyl)-4-(2-chloroethyl)benzamide:

$^1$H NMR (600 MHz, DMSO-$d_6$) δ 12.43 (s, 1H), 8.10 (d, *J* = 2.2 Hz, 1H), 7.95 (dd, *J* = 8.8, 2.2 Hz, 1H), 7.82 (d, *J* = 8.0 Hz, 2H), 7.40 (d, *J* = 8.0 Hz, 2H), 7.31 (d, *J* = 8.8 Hz, 1H), 4.23 (q, *J* = 7.0 Hz, 2H), 3.88 (t, *J* = 6.8 Hz, 2H), 3.09 (t, *J* = 6.8 Hz, 2H), 1.38 (t, *J* = 7.0 Hz, 3H).

MS (ESI) m/z calculated for $[C_{17}H_{17}BrClNO_4S]^+$ ($[M+H]^+$): 445.98, observed: 446.1.

**Compound 23:** *N*-((3-bromo-4-ethoxyphenyl)sulfonyl)-2-(3-chlorophenyl)acetamide:

$^1$H NMR (600 MHz, DMSO-$d_6$) δ 12.36 (s, 1H), 7.98 (d, *J* = 2.3 Hz, 1H), 7.86 (dd, *J* = 8.8, 2.3 Hz, 1H), 7.31 (dd, *J* = 4.5 Hz, 1H), 7.29 (d, *J* = 8.8 Hz, 1H), 7.22 (s, 1H), 7.13 (t, *J* = 4.1 Hz, 1H), 4.23 (q, *J* = 6.9 Hz, 2H), 3.59 (s, 2H), 1.39 (t, *J* = 6.9 Hz, 3H).

MS (ESI) m/z calculated for $[C_{16}H_{15}BrClNO_4S]^+$ ($[M+H]^+$): 431.96, observed: 432.1.

**Compound 24:** *N*-((3-bromo-4-ethoxyphenyl)sulfonyl)-2-(3-bromophenyl)acetamide:

$^1$H NMR (600 MHz, DMSO-$d_6$) δ 12.36 (s, 1H), 7.98 (d, *J* = 2.3 Hz, 1H), 7.86 (dd, *J* = 8.8, 2.3 Hz, 1H), 7.44 (dd, *J* = 7.6, 2.0 Hz, 1H), 7.37 (d, *J* = 2.0 Hz, 1H), 7.29 (d, *J* = 8.8 Hz, 1H), 7.25 (t, *J* = 7.8 Hz, 1H), 7.17 (d, *J* = 7.6 Hz, 1H), 4.23 (q, *J* = 6.9 Hz, 2H), 3.58 (s, 2H), 1.39 (t, *J* = 6.9 Hz, 3H).

MS (ESI) m/z calculated for $[C_{16}H_{15}Br_2NO_4S]^+$ ($[M+H]^+$): 475.91, observed: 476.1.

**Compound 25:** *N*-(phenylsulfonyl)-2-(*p*-tolyl)acetamide:

$^1$H NMR (600 MHz, DMSO-$d_6$): δ 12.31 (s, 1H), 7.89 (d, *J* = 7.9 Hz, 2H), 7.7 (t, *J* = 7.3 Hz, 1H), 7.61 (dd, *J* = 7.9, 7.3 Hz, 2H), 7.08 (d, *J* = 7.7 Hz, 2H), 7.02 (d, *J* = 7.7 Hz, 2H), 3.49 (s, 3H), 2.25 (s, 3H).

MS (ESI) m/z calculated for $[C_{16}H_{17}NO_4S]^+$ ($[M+H]^+$): 290.08, observed: 290.1

**Compound 26:** *N*-((3-bromo-4-ethoxyphenyl)sulfonyl)-2-(*p*-tolyl)acetamide:

$^1$H NMR (600 MHz, DMSO-$d_6$): δ 12.14 (s, 1H), 7.80 (d, *J* = 8.6 Hz, 2H), 7,10 – 7.06 (m, 4H), 7.02 (d, *J* = 7.7 Hz, 2H), 4.12 (q, *J* = 7.0 Hz, 2H), 3.46 (s, 2H), 2.25 (s, 3H), 1.35 (t, *J* = 7.0 Hz, 3H).

MS (ESI) m/z calculated for $[C_{17}H_{19}NO_4S]^+$ ($[M+H]^+$): 334.103, observed: 334.2.

**Compound 27:** *N*-((4-methoxyphenyl)sulfonyl)-2-(*p*-tolyl)acetamide:

$^1$H NMR (600 MHz, DMSO-$d_6$): δ 12.15 (s, 1H), 7.82 (d, *J* = 8.8 Hz, 2H), 7.11 (d, *J* = 8.8 Hz, 2H), 7.08 (d, *J* = 7.7 Hz, 2H), 7.02 (d, *J* = 7.7 Hz, 2H), 3.48 (s, 3H), 3.46 (s, 2H), 2.25 (s, 3H).

MS (ESI) m/z calculated for $[C_{16}H_{17}NO_4S]^+$ ($[M+H]^+$): 320.0878, observed: 320.2.

**Compound 28:** 2-(*p*-tolyl)-*N*-((4-(trifluoromethoxy)phenyl)sulfonyl)acetamide:

$^1$H NMR (600 MHz, DMSO-$d_6$): δ 12.46 (s, 1H), 8.02 (d, *J* = 8.6 Hz, 2H), 7.60 (d, *J* = 8.6 Hz, 2H), 7.07 (d, *J* = 7.8 Hz, 2H), 7.02 (d, *J* = 7.8 Hz, 2H), 3.50 (s, 2H), 2.25 (s, 3H).

MS (ESI) m/z calculated for $[C_{16}H_{14}F_3NO_4S]^+$ ($[M+H]^+$): 374.0596, observed: 374.2.

**Compound 29:** N-((4-(tert-butyl)phenyl)sulfonyl)-2-(p-tolyl)acetamide:

$^1$H NMR (600 MHz, DMSO-d$_6$): δ 12.23 (s, 1H), 7.81 (d, *J* = 8.4 Hz, 2H), 7.62 (d, *J* = 8.4 Hz, 2H), 7.07 (d, *J* = 7.8 Hz, 2H), 7.02 (d, *J* = 7.8 Hz, 2H), 3.48 (s, 2H), 2.25 (s, 3H), 1.30 (s, 9H).

MS (ESI) m/z calculated for $[C_{19}H_{23}NO_3S]^+$ ($[M+H]^+$): 346.1399, observed: 346.2.

**Compound 30:** 2-(*p*-tolyl)-*N*-((4-(trifluoromethyl)phenyl)sulfonyl)acetamide:

$^1$H NMR (600 MHz, DMSO-d$_6$): δ 12.57 (s, 1H), 8.09 (d, *J* = 8.2 Hz, 2H), 8.0 (d, *J* = 8.2 Hz, 2H), 7.07 (d, *J* = 7.8 Hz, 2H), 7.02 (d, *J* = 7.8 Hz, 2H), 3.51 (s, 2H), 2.25 (s, 3H).

MS (ESI) m/z calculated for $[C_{16}H_{14}F_3NO_3S]^+$ ($[M+H]^+$): 358.0646, observed: 358.2.

**Compound 32:** *N*-((3-bromo-4-methylphenyl)sulfonyl)-2-(*p*-tolyl)acetamide:

$^1$H NMR (600 MHz, DMSO-d$_6$): δ 12.40 (s, 1H), 7.97 (s, 1H), 7.78 (d, *J* = 8.0 Hz, 1H), 7.59 (d, *J* = 8.0 Hz, 1H), 7.08 (d, *J* = 7.7 Hz, 2H), 7.04 (d, *J* = 7.7 Hz, 2H), 3.49 (s, 2H), 2.42 (s, 3H), 2.26 (s, 3H).

MS (ESI) m/z calculated for $[C_{16}H_{16}BrNO_3S]^+$ ($[M+H]^+$): 381.0034, observed: 382.1

**Compound 33:** *N*-((4-bromo-3-fluorophenyl)sulfonyl)-2-(p-tolyl)acetamide:

$^1$H NMR (600 MHz, DMSO-d$_6$): δ 12.57 (s, 1H), 7.98 (dd, *J* = 8.3, 8.0 Hz, 1H), 7.76 (d, *J* = 8.0 Hz, 1H), 7.65 (d, *J* = 83 Hz, 1H), 7.08 (d, *J* = 7.7 Hz, 2H), 7.04 (d, *J* = 7.7 Hz, 2H), 3.49 (s, 2H), 2.26(s, 3H).

MS (ESI) m/z calculated for $[C_{15}H_{13}BrFNO_3S]^+$ ($[M+H]^+$): 385.9784, observed: 386.1.

**Compound 34:** *N*-((4-ethoxy-3-(1-methyl-1*H*-pyrazol-3-yl)phenyl)sulfonyl)-2-(*p*-tolyl) aceta-mide:

$^1$H NMR (600 MHz, DMSO-d$_6$) δ 12.14 (s, 1H), 8.47 (d, *J* = 2.5 Hz, 1H), 7.77 (dd, *J* = 8.8, 2.5 Hz, 2H), 7.25 (d, *J* = 8.8 Hz, 1H), 7.05 (d, *J* = 7.9 Hz, 2H), 7.02 (d, *J* = 7.9 Hz, 2H), 6.79 (d, *J* = 2.2 Hz, 1H), 4.23 (q, *J* = 6.9 Hz, 2H), 3.92 (s, 3H), 3.47 (s, 2H), 2.23 (s, 3H), 1.44 (t, *J* = 6.9 Hz, 3H).

MS (ESI) m/z calculated for $[C_{21}H_{23}N_3O_4S]^+$ ($[M+H]^+$): 413.14, observed: 414.4.

**Compound 35:** *N*-((5-bromo-2-fluorophenyl)sulfonyl)-2-(*p*-tolyl)acetamide:

$^1$H NMR (600 MHz, DMSO-d$_6$): δ 12.90 (s, 1H), 7.98 – 7.94 (m, 1H), 7.93 – 7.90 (m, 1H), 7.46 (t, *J* = 9.4 Hz, 1H), 7.09 (d, *J* = 7.9 Hz, 2H), 7.04 (d, *J* = 7.9 Hz, 2H), 3.52 (s, 2H), 2.26 (s, 3H).

MS (ESI) m/z calculated for $[C_{15}H_{13}BrFNO_3S]^+$ ($[M+H]^+$): 385.9784, observed: 386.1.

**Compound 36:** *N*-((2,5-dichlorophenyl)sulfonyl)-2-(*p*-tolyl)acetamide:

$^1$H NMR (600 MHz, DMSO-d$_6$): δ 12.92 (s, 1H), 7.99 (d, *J* = 2.4 Hz, 1H), 7.80 (dd, *J* = 8.6, 2.4 Hz, 1H), 7.73 (d, *J* = 8.6 Hz, 1H), 7.09 (d, *J* = 7.9 Hz, 2H), 7.05 (d, *J* = 7.9 Hz, 2H), 3.56 (s, 2H), 2.26 (s, 3H).

MS (ESI) m/z calculated for $[C_{15}H_{13}Cl_2NO_3S]^+$ ($[M+H]^+$): 357.9993, observed: 358.1.

**Compound 37:** *N*-((2-fluoro-5-(trifluoromethyl)phenyl)sulfonyl)-2-(*p*-tolyl)acetamide:

$^1$H NMR (600 MHz, DMSO-d$_6$): δ 13.01 (s, 1H), 8.18 (d, *J* = 7.7 Hz, 1H), 8.09 (d, *J* = 5.2 Hz, 1H), 7.72 (t, *J* = 9.0 Hz, 1H), 7.08 (d, *J* = 7.8, Hz, 2H), 7.04 (d, *J* = 7.8 Hz, 2H), 3.51 (s, 2H), 2.25(s, 3H).

MS (ESI) m/z calculated for [$C_{16}H_{13}F_4NO_3S$]$^+$ ([M+H]$^+$): 376.0552, observed: 376.2

**<u>Compound 38</u>:** N-((3-chloro-2-methylphenyl)sulfonyl)-2-(p-tolyl)acetamide:

$^1$H NMR (600 MHz, DMSO-d$_6$): δ 12.57 (s, 1H), 7.93 (d, *J* = 8.0 Hz, 1H), 7.76 (d, *J* = 8.0 Hz, 1H), 7.44 (t, *J* = 8.0 Hz, 1H), 7.08 (d, *J* = 7.7 Hz, 2H), 7.02 (d, *J* = 7.7 Hz, 2H), 3.51 (s, 2H), 2.49 (s, 3H), 2.26 (s, 3H).

MS (ESI) m/z calculated for [$C_{16}H_{16}ClNO_3S$]$^+$ ([M+H]$^+$): 338.0539, observed: 338.2.

**<u>Compound 41:</u>** *N*-((3-bromo-4-ethoxyphenyl)sulfonyl)-5-methoxyfuran-2-carboxa-mide:

$^1$H NMR (600 MHz, DMSO-d$_6$) δ 12.18 (s, 1H), 8.07 (d, *J* = 2.3 Hz, 1H), 7.92 (dd, *J* = 8.8, 2.3 Hz, 1H), 7.49 (d, *J* = 3.8 Hz, 1H), 7.31 (d, *J* = 8.8 Hz, 1H), 5.66 (d, *J* = 3.8 Hz, 1H), 4.23 (q, *J* = 7.0 Hz, 2H), 3.92 (s, 3H), 1.39 (t, *J* = 7.0 Hz, 3H).

MS (ESI) m/z calculated for [$C_{14}H_{14}BrNO_6S$]$^+$ ([M+H]$^+$): 403.97, observed: 404.2.

**<u>Compound 42</u>:** *N*-((3-bromo-4-ethoxyphenyl)sulfonyl)-2-phenylpropanamide:

$^1$H NMR (600 MHz, DMSO-d$_6$): δ 12.23 (s, 1H), 7.91 (d, *J* = 2.0, Hz, 1H), 7.80 (dd, *J* = 8.6, 2.0 Hz, 1H), 7.30 – 7.23 (m, 3H), 7.27 (d, *J* = 8.6, Hz, 1H), 7.14 (d, *J* = 7.5 Hz, 2H), 4.23 (q, *J* = 7.0 Hz, 2H), 3.70 (q, *J* = 7.0 Hz, 2H), 1.38 (t, *J* = 7.0 Hz, 3H), 1.24 (d, *J* = 7.0 Hz, 3H).

MS (ESI) m/z calculated for [$C_{17}H_{18}BrNO_4S$]$^+$ ([M+H]$^+$): 412.0140, observed: 412.2.

**<u>Compound 47</u>:** *N*-(benzo[*d*]thiazol-6-ylsulfonyl)-2-(*p*-tolyl)acetamide:

$^1$H NMR (600 MHz, DMSO-d$_6$): δ 12.44 (s, 1H), 9.66 (s, 1H), 8.81 (s, 1H), 8.26 (d, *J* = 8.7 Hz, 1H), 7.99 (d, *J* = 8.7 Hz, 1H), 7.05 (d, *J* = 7.8 Hz, 2H), 7.02 (d, *J* = 7.8 Hz, 2H), 3.49 (s, 2H), 2.23 (s, 3H).

MS (ESI) m/z calculated for [$C_{16}H_{14}N_2O_3S_2$]$^+$ ([M+H]$^+$): 347.0446, observed: 347.2.

**<u>Compound 48:</u>** *N*-(quinolin-5-ylsulfonyl)-2-(*p*-tolyl)acetamide:

$^1$H NMR (600 MHz, DMSO-d$_6$): δ 12.70 (s, 1H), 9.04 (d, *J* = 3.9 Hz, 1H), 8.92 (d, *J* = 8.8 Hz, 1H), 8.33 (dd, *J* = 7.8, 7.0 Hz, 2H), 7.92 (t, *J* = 7.8 Hz, 1H), 7.71 (dd, *J* = 8.8, 3.9 Hz, 1H), 6.93 (d, *J* = 7.7 Hz, 1H), 6.88 (d, *J* = 7.7 Hz, 2H), 3.43 (s, 2H), 2.20 (s, 3H).

MS (ESI) m/z calculated for [$C_{16}H_{14}N_2O_3S_2$]$^+$ ([M+H]$^+$): 341.0882, observed: 341.2.

**Note:** The identities of compounds **2**, **15**, **17**, **18**, **31**, **39**, **40**, **43**, **44**, **45**, **46**, and **49** were determined by LC-MS.

$^{1}$H NMR of **Compound 1:** *N*-((3-bromo-4-ethoxyphenyl)sulfonyl)-2-(*p*-tolyl)acetamide:

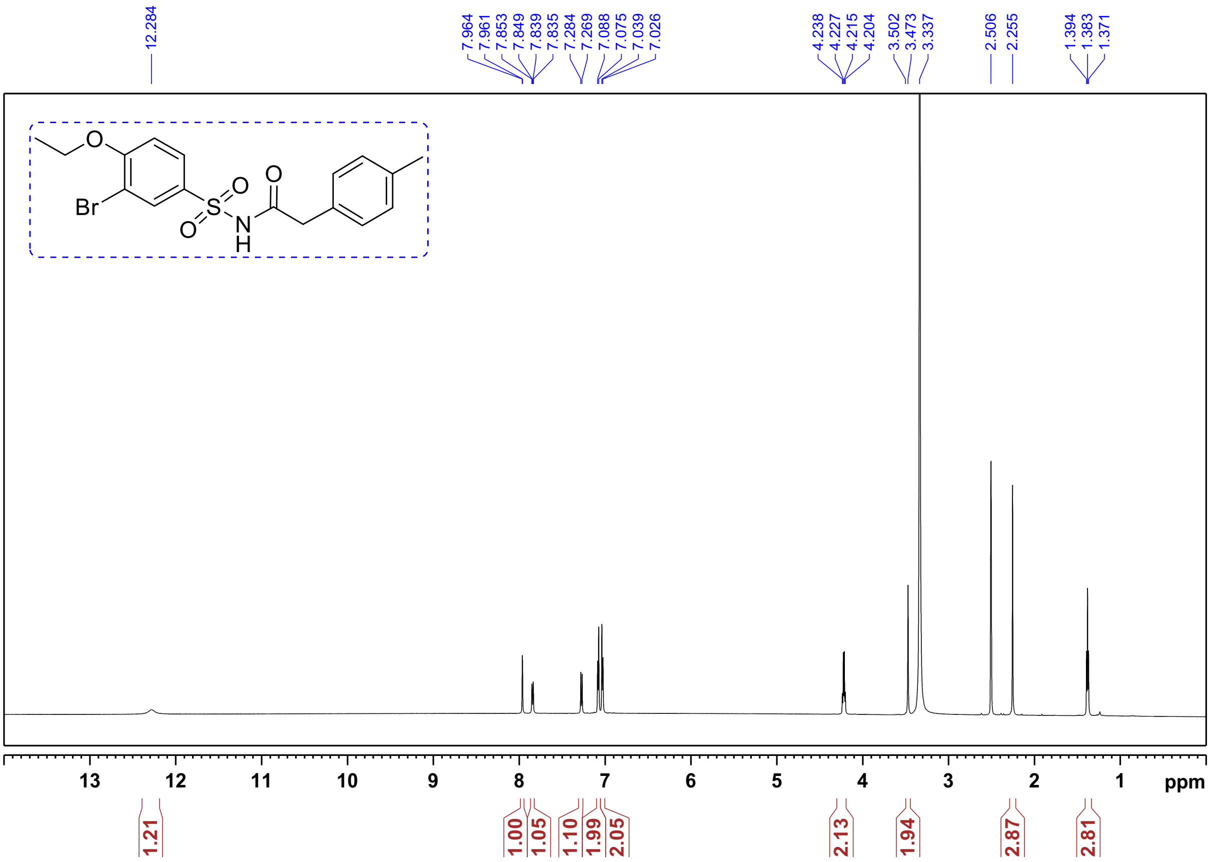


$^{1}$H NMR of **Compound 3:** *N*-((3-bromo-4-ethoxyphenyl)sulfonyl)-2-(4-fluorophenyl)acetamide:

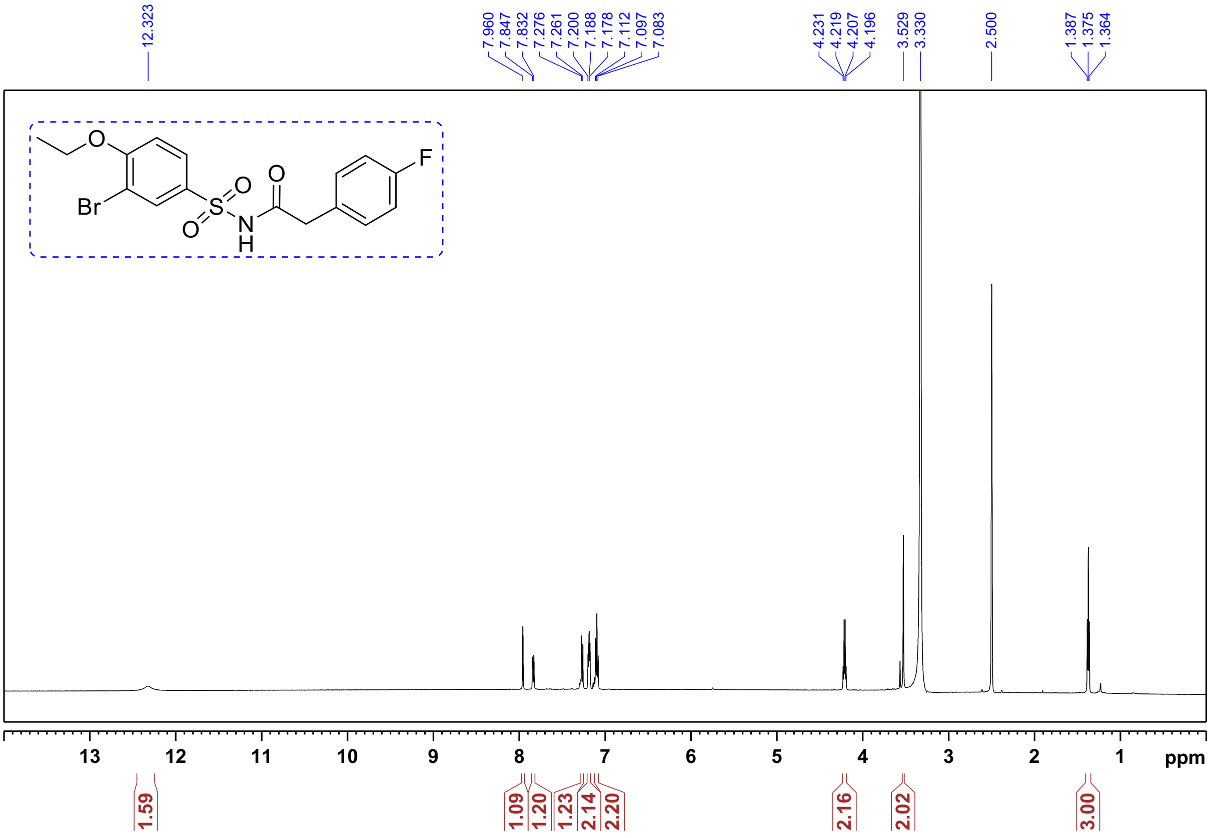

$^{1}$H NMR of **Compound 4**: *N*-((3-bromo-4-ethoxyphenyl)sulfonyl)-2-(4-chlorophenyl)acetamide:

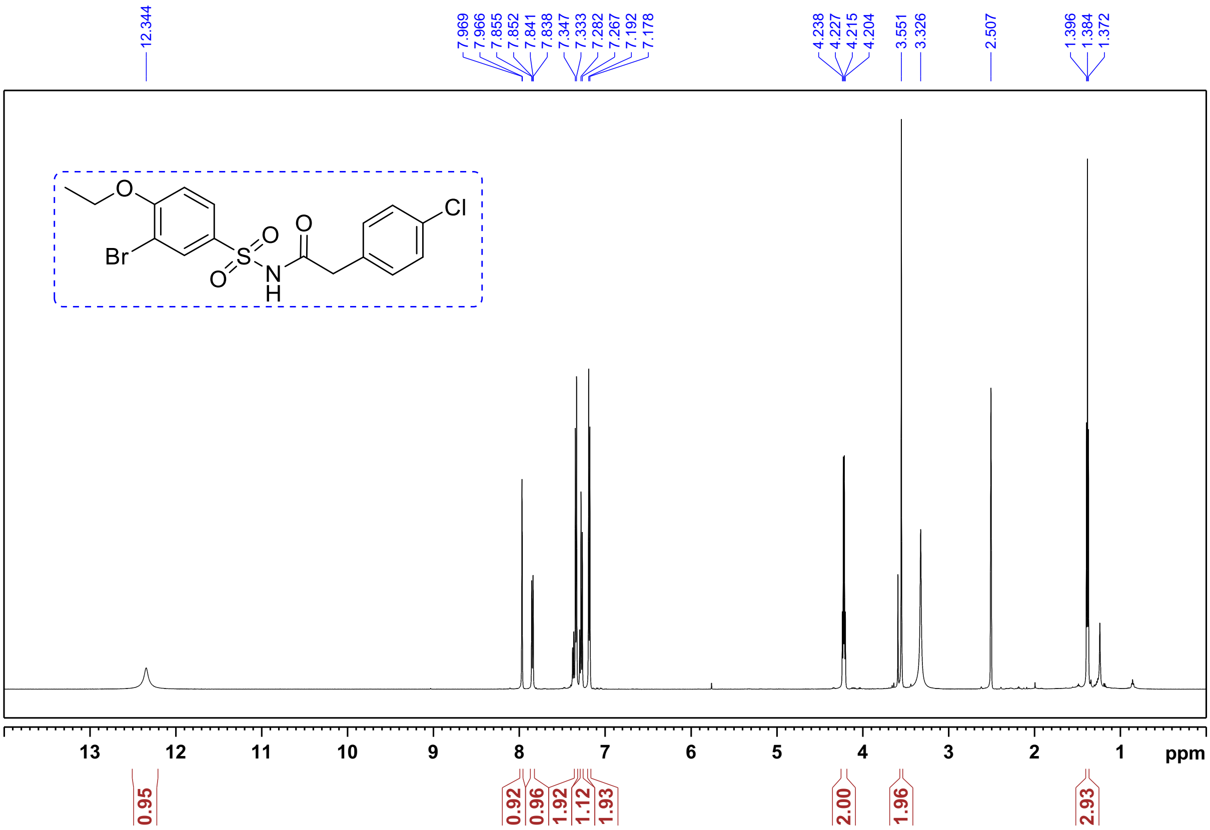


$^{13}$C NMR of **Compound 4:** *N*-((3-bromo-4-ethoxyphenyl)sulfonyl)-2-(4-chlorophenyl)acetamide:

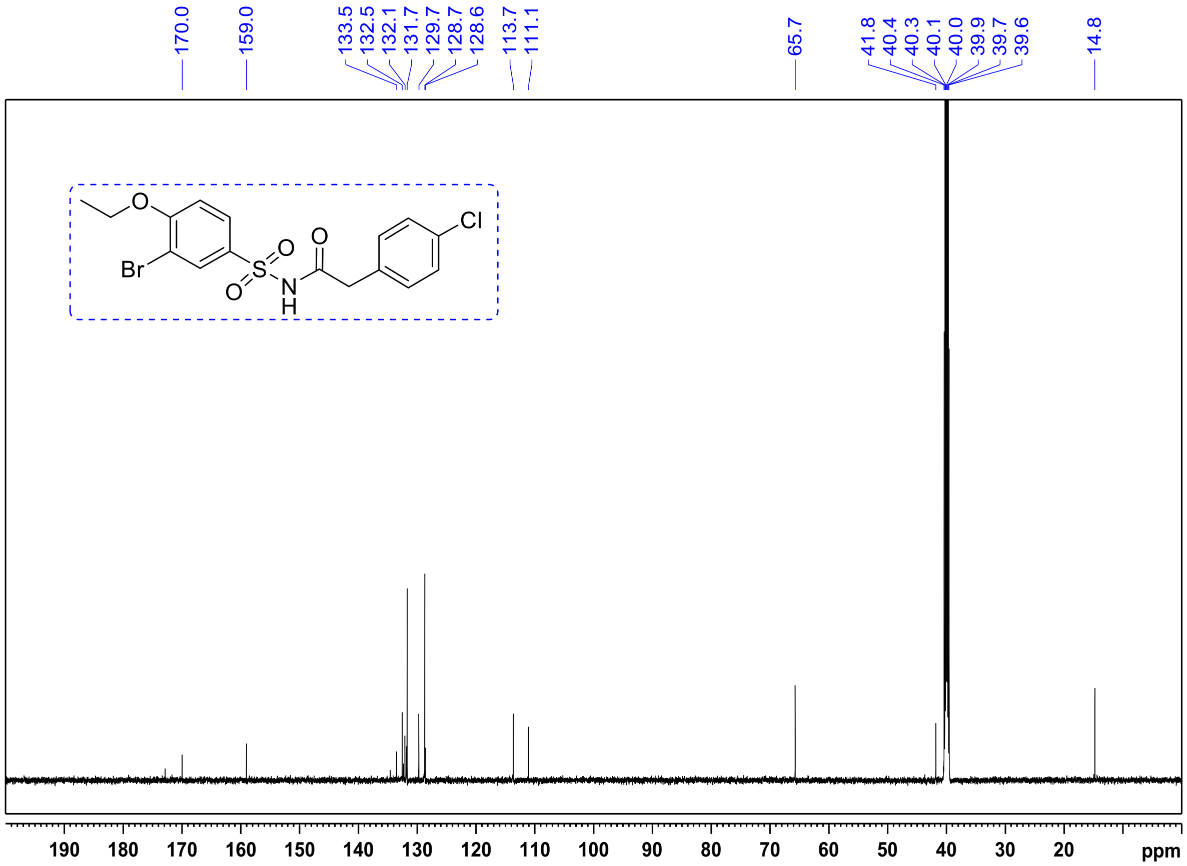

[1]H NMR of **<u>Compound 5:</u>** *N*-((3-bromo-4-ethoxyphenyl)sulfonyl)-2-(4-bromophenyl)acetamide.

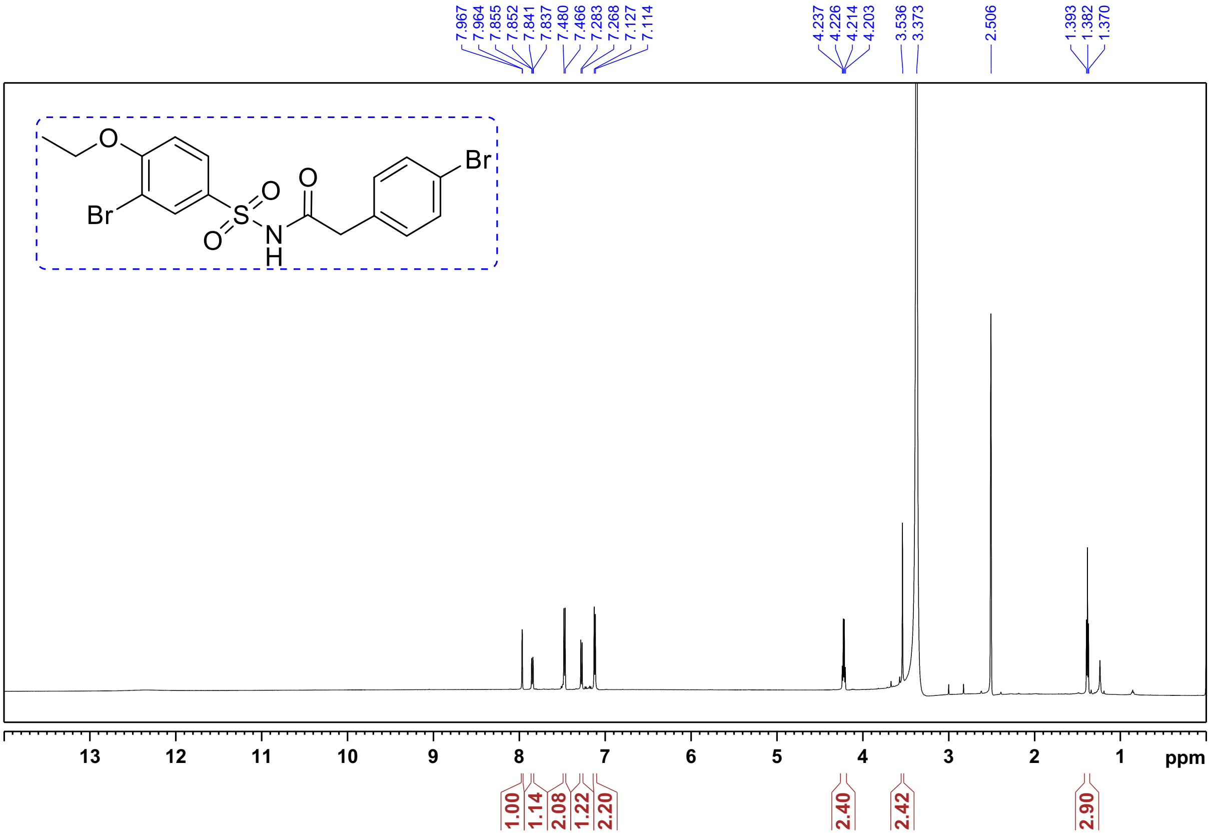


[1]H NMR of **<u>Compound 6</u>:** *N*-((3-bromo-4-ethoxyphenyl)sulfonyl)-2-(4-iodophenyl)acetamide.

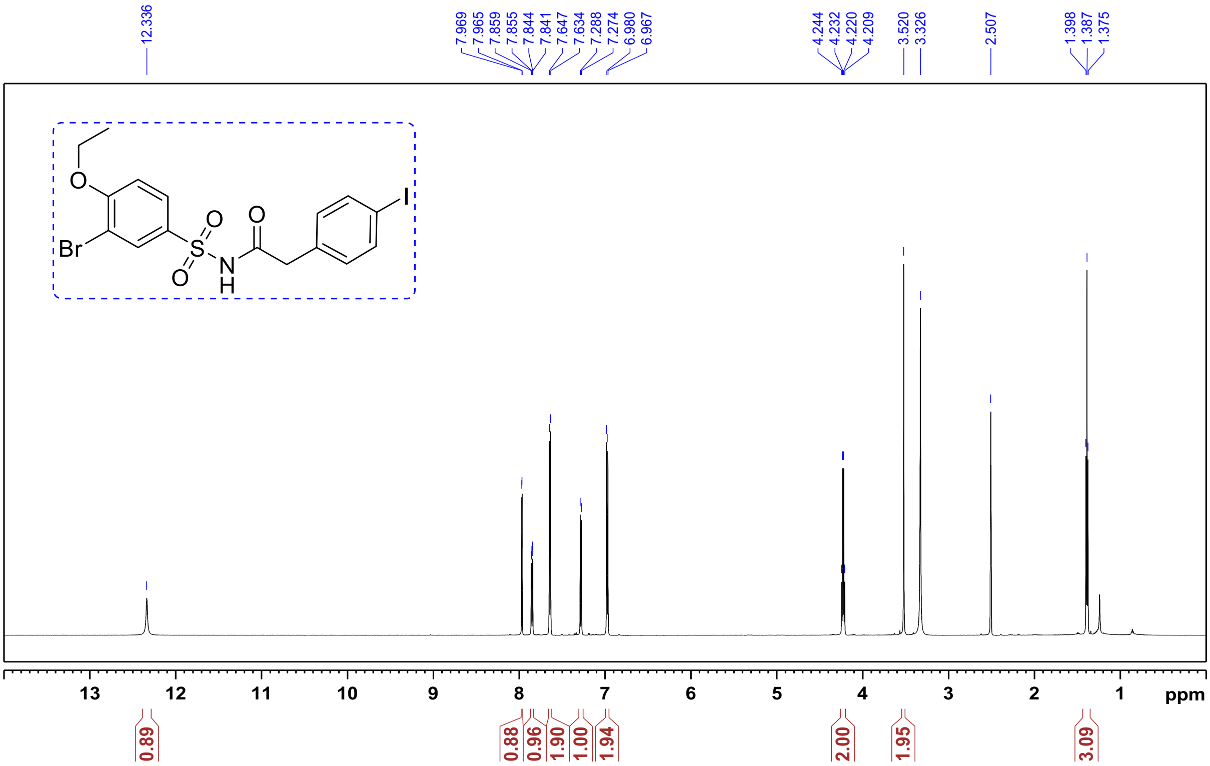

$^{13}C$ NMR of **Compound 6**: *N*-((3-bromo-4-ethoxyphenyl)sulfonyl)-2-(4-iodophenyl)acetamide.

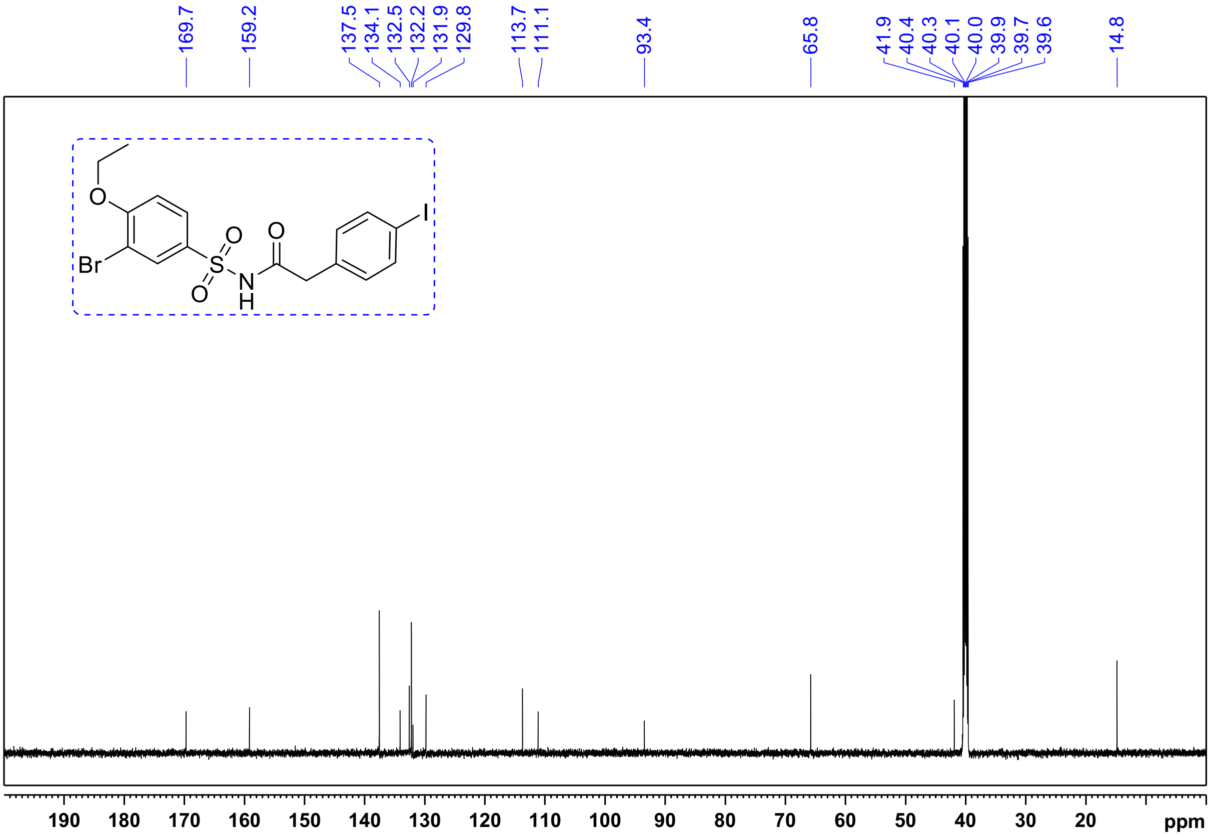


$^{1}H$ NMR of **Compound 7:** N-((3-bromo-4-ethoxyphenyl)sulfonyl)-2-(4-methoxyphenyl)acetamide.

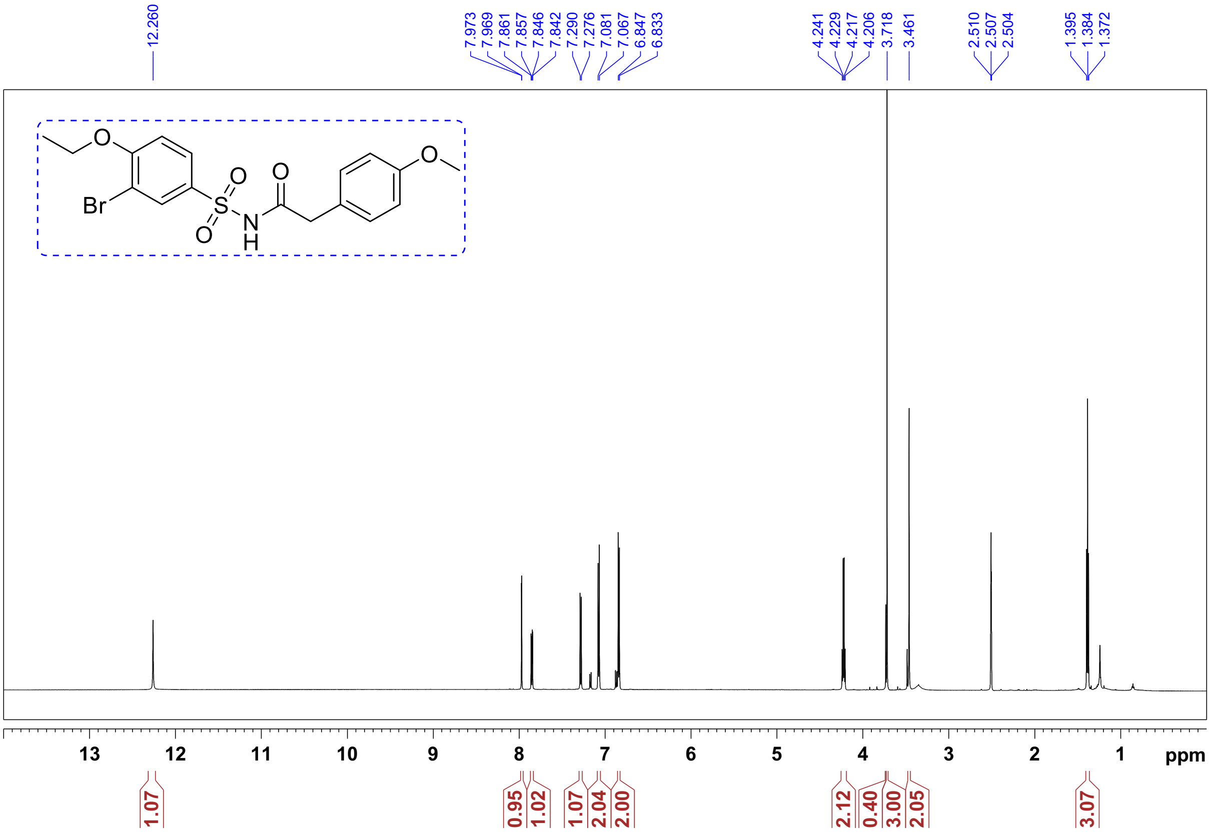

[1]H NMR of **Compound 8**: N-((3-bromo-4-ethoxyphenyl)sulfonyl)-2-(4-isopropoxyphenyl)acetamide.

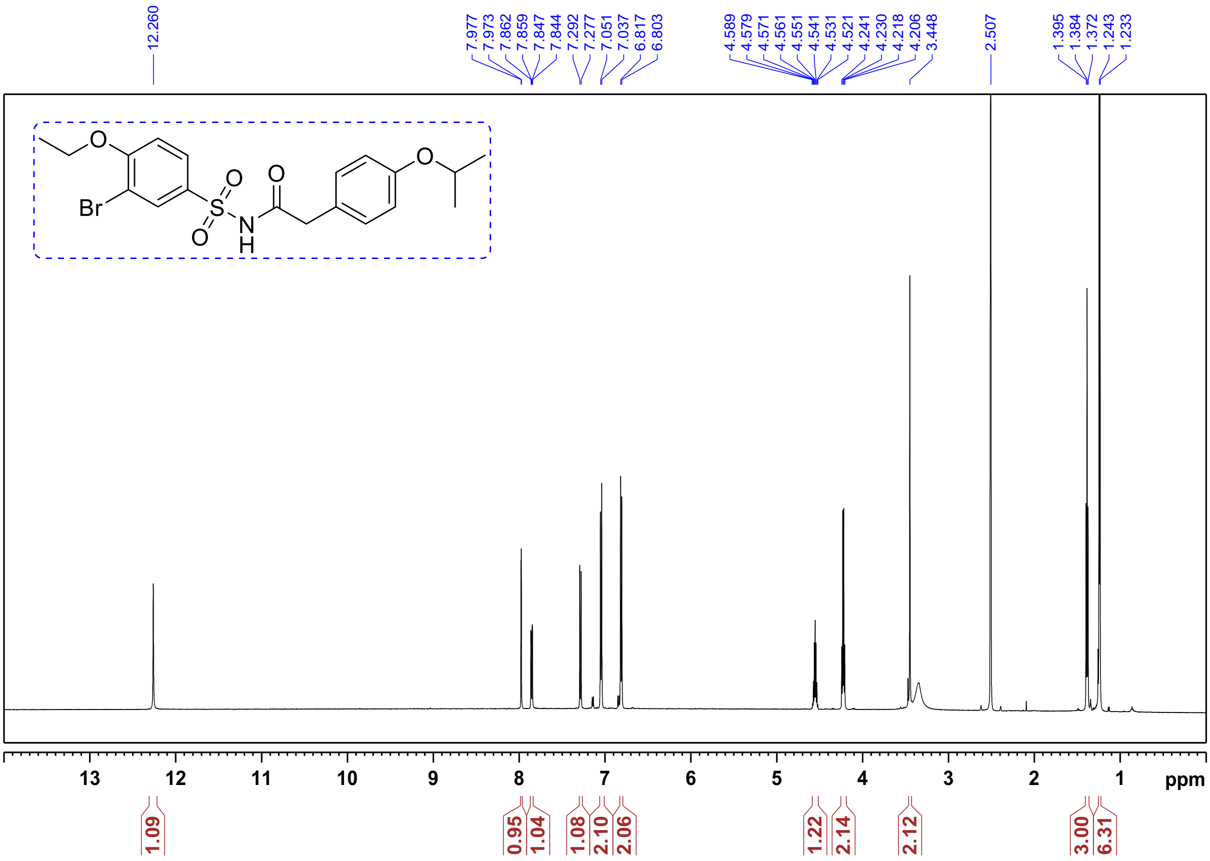

[1]H NMR of **Compound 9**: *N*-((3-bromo-4-ethoxyphenyl)sulfonyl)-2-(4-phenoxyphenyl)acetamide.

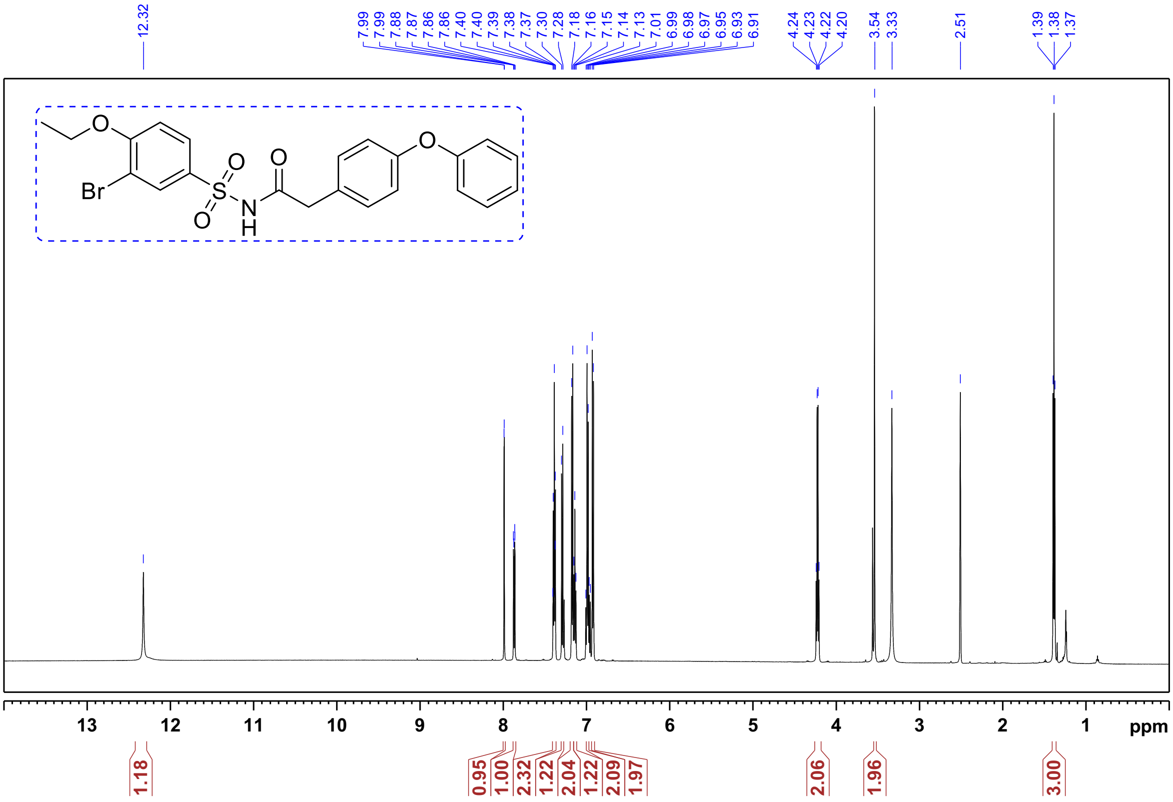

$^1$H NMR of **Compound 10**: *N*-((3-bromo-4-ethoxyphenyl)sulfonyl)-2-(4-(methylthio)phenyl)acetamide:

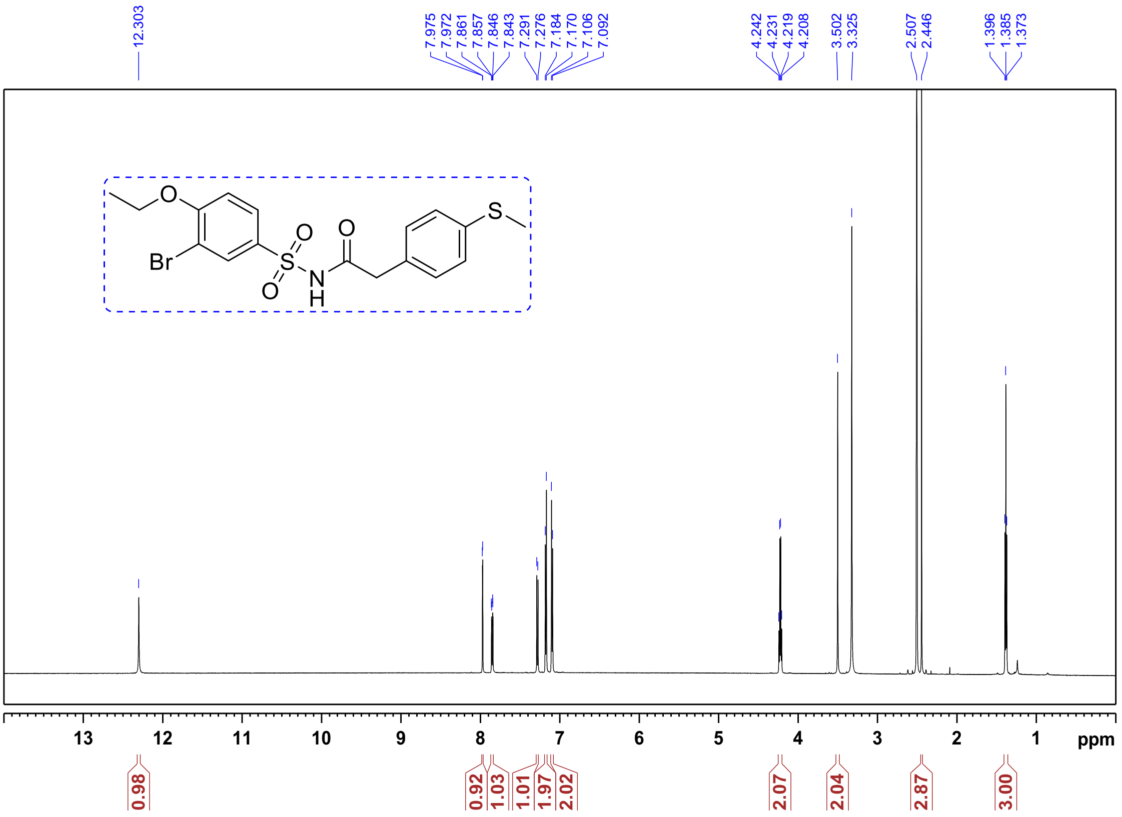


$^{13}$C NMR of **Compound 10**: *N*-((3-bromo-4-ethoxyphenyl)sulfonyl)-2-(4-(methylthio)phenyl)acetamide:

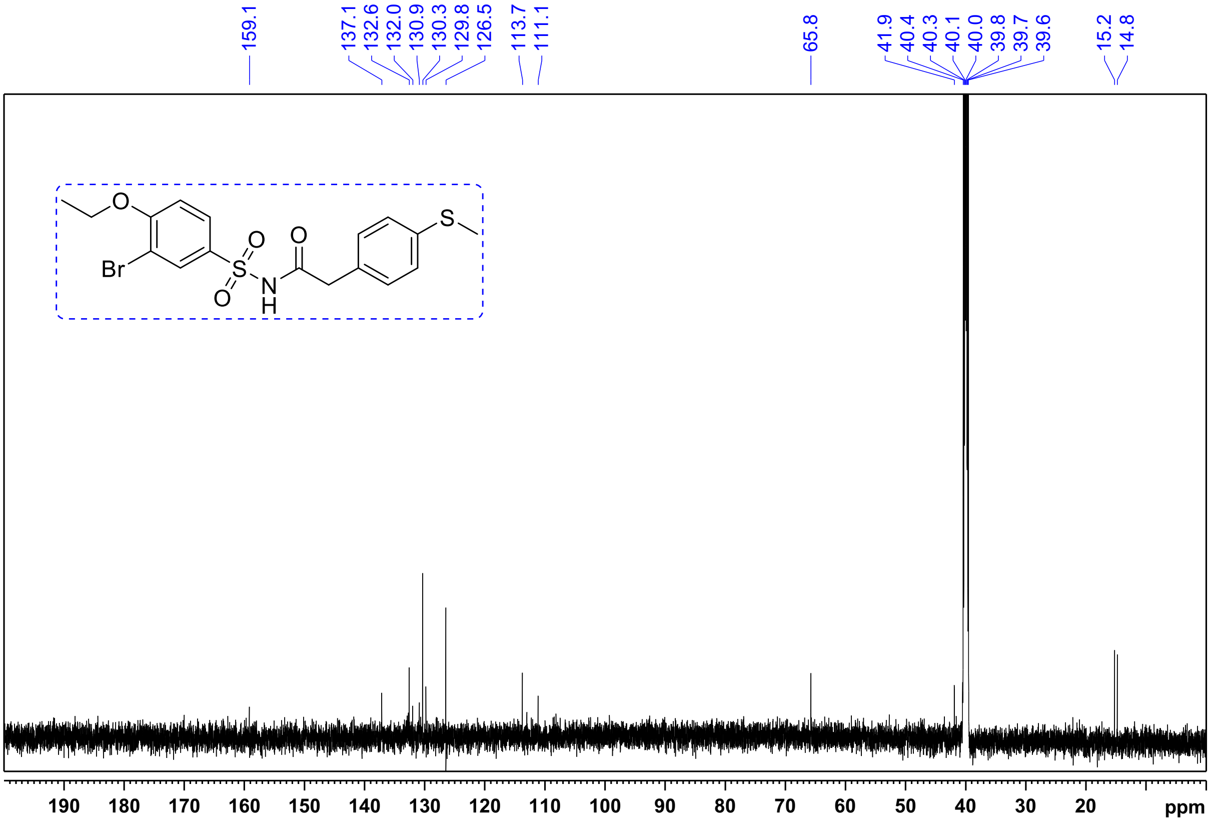

[1]H NMR of **Compound 11:** *N*-((3-bromo-4-ethoxyphenyl)sulfonyl)-2-(4-((trifluoromethyl)thio)phenyl) acetamide:

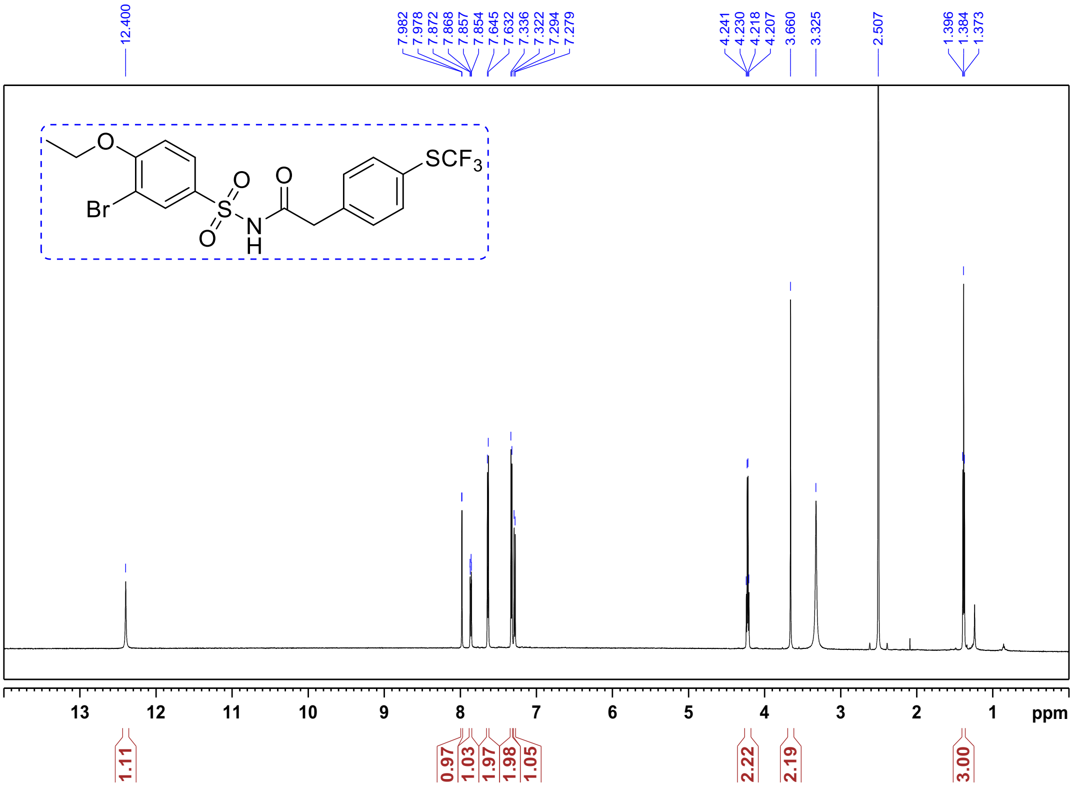


[1]H NMR of **Compound 12:** *N*-((3-bromo-4-ethoxyphenyl)sulfonyl)-2-(4-cyanophenyl)acetamide.

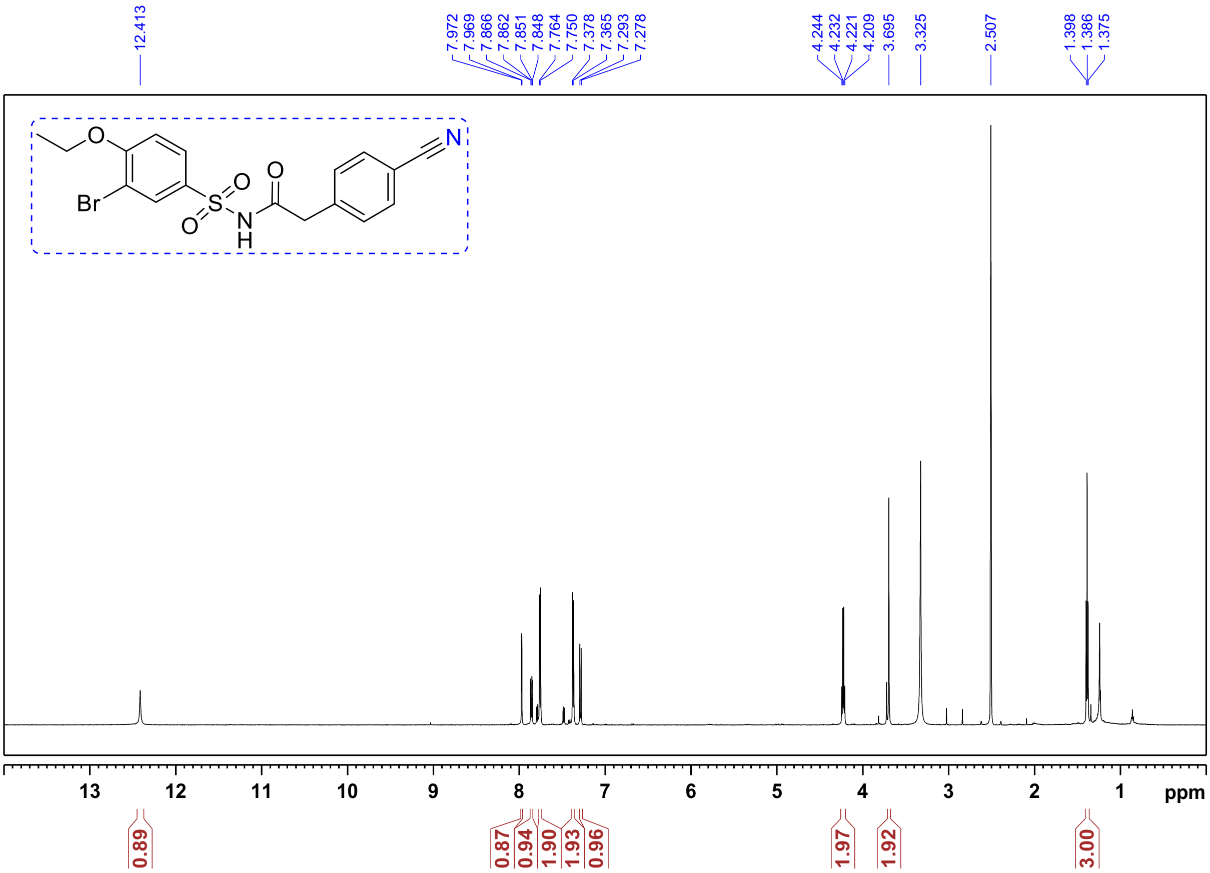

[1]H NMR of **Compound 13:** *N*-((3-bromo-4-ethoxyphenyl)sulfonyl)-2-(o-tolyl)acetamide:

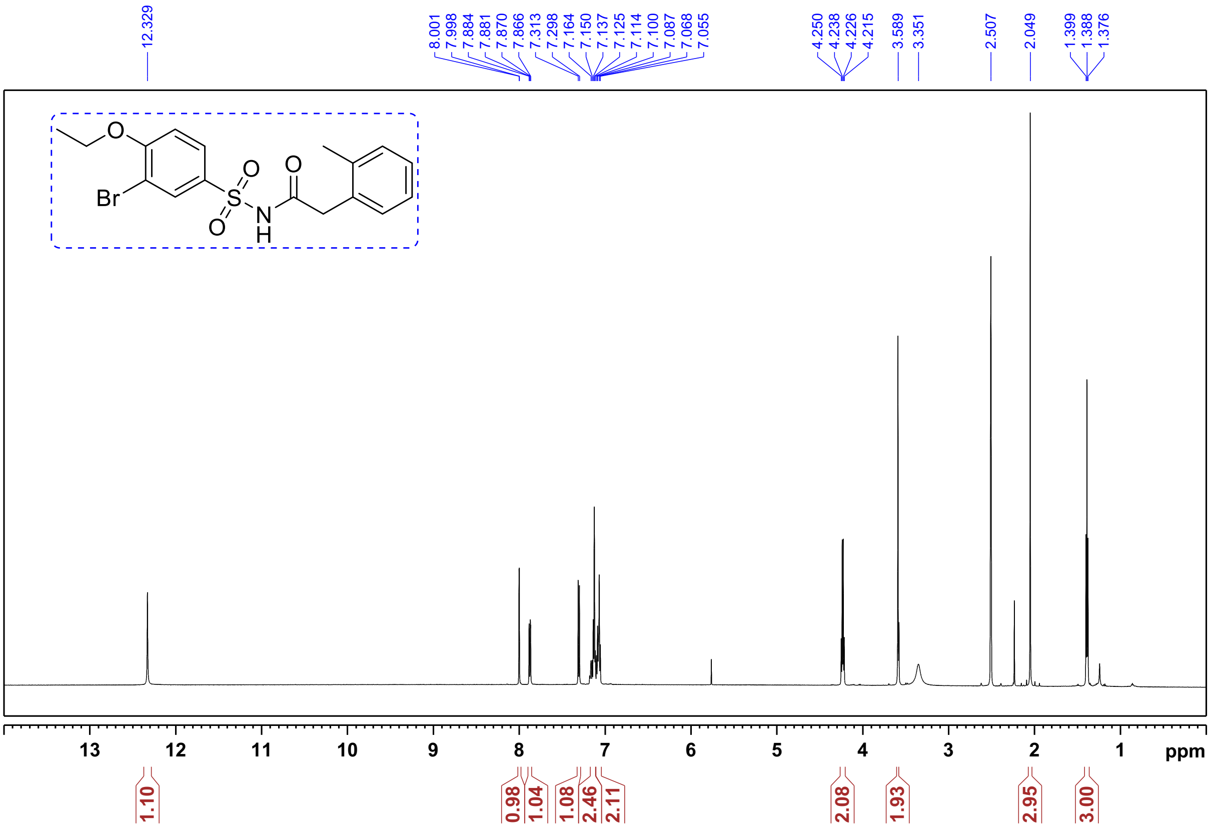

[1]H NMR of **Compound 14:** N-((3-bromo-4-ethoxyphenyl)sulfonyl)-2-(2-(trifluoromethyl)phenyl) acetamide

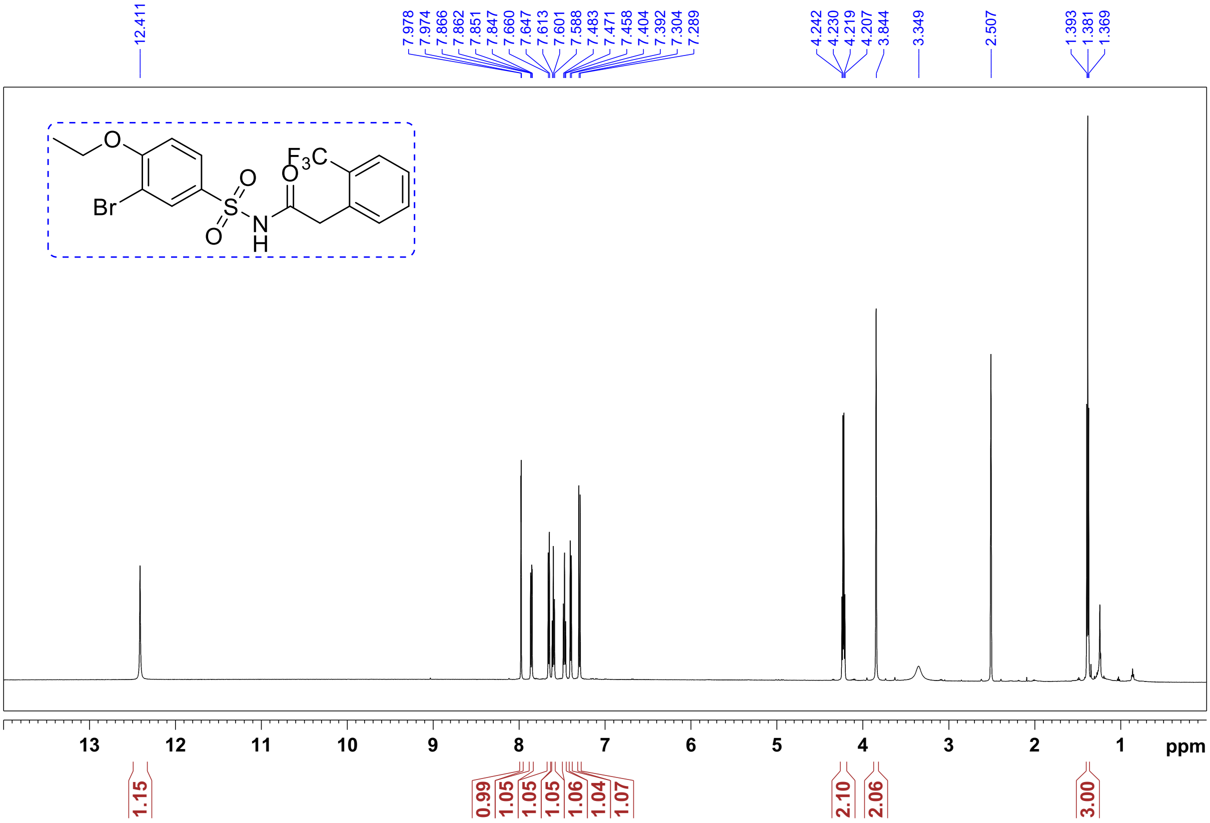

[1]H NMR of **Compound 16:** *N*-((3-bromo-4-ethoxyphenyl)sulfonyl)-2-(2-iodophenyl)acetamide:

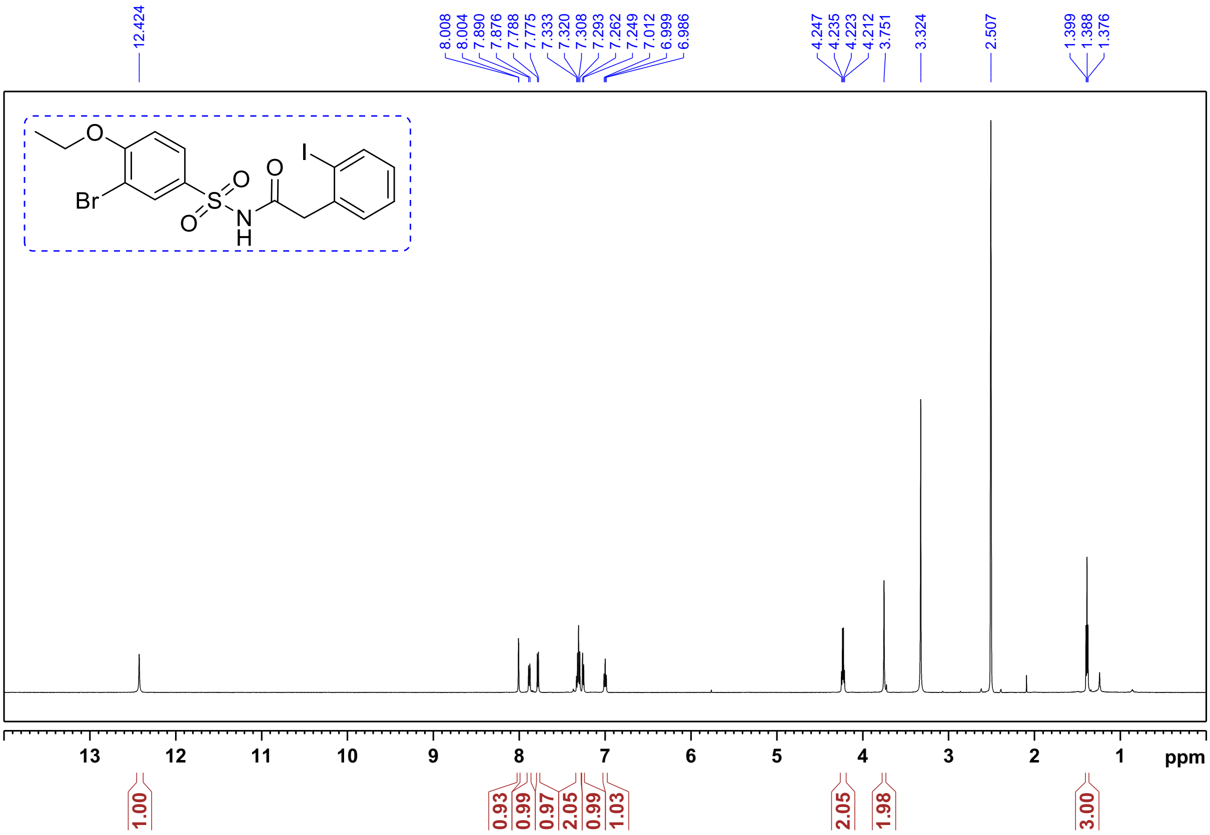

[1]H NMR of **Compound 19:** N-((3-bromo-4-ethoxyphenyl)sulfonyl)-2-(2-fluorophenyl)acetamide:

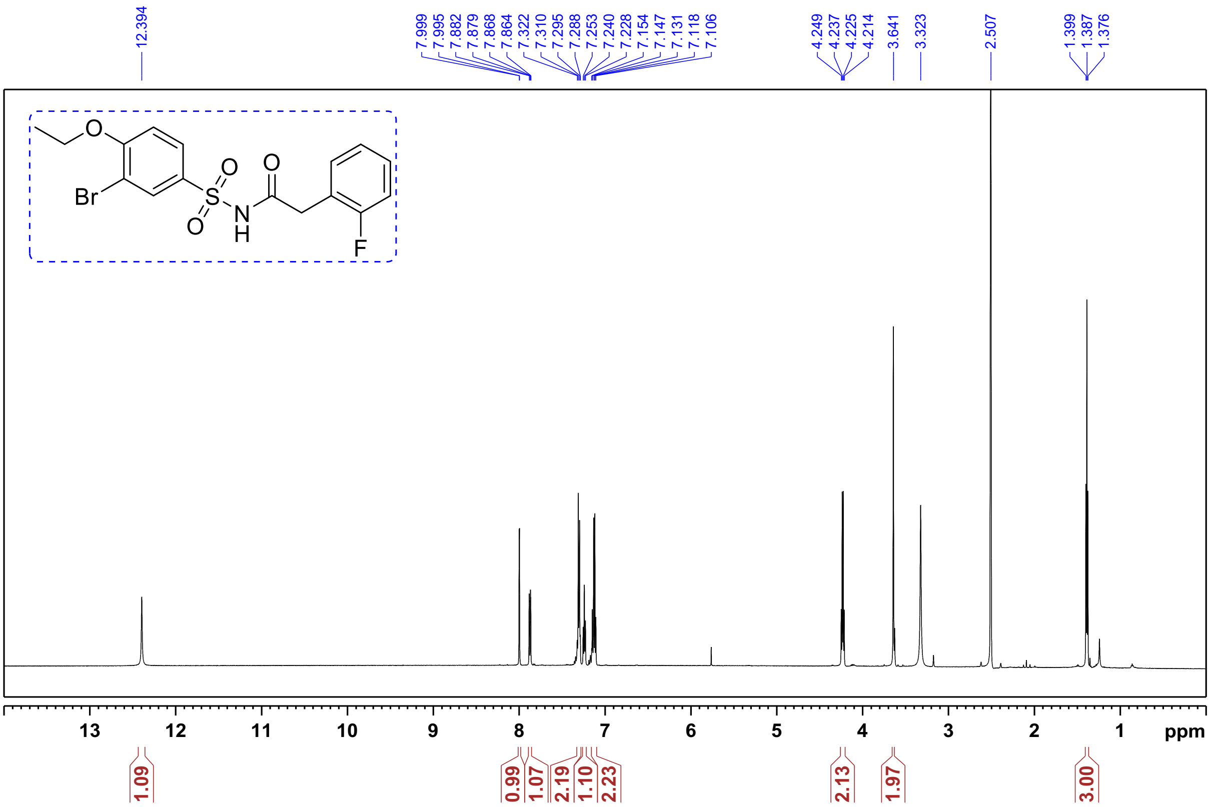

[1]H NMR of **Compound 20**: *N*-((3-bromo-4-ethoxyphenyl)sulfonyl)-2-(2 (trifluoromethoxy)phenyl)

acetamide.

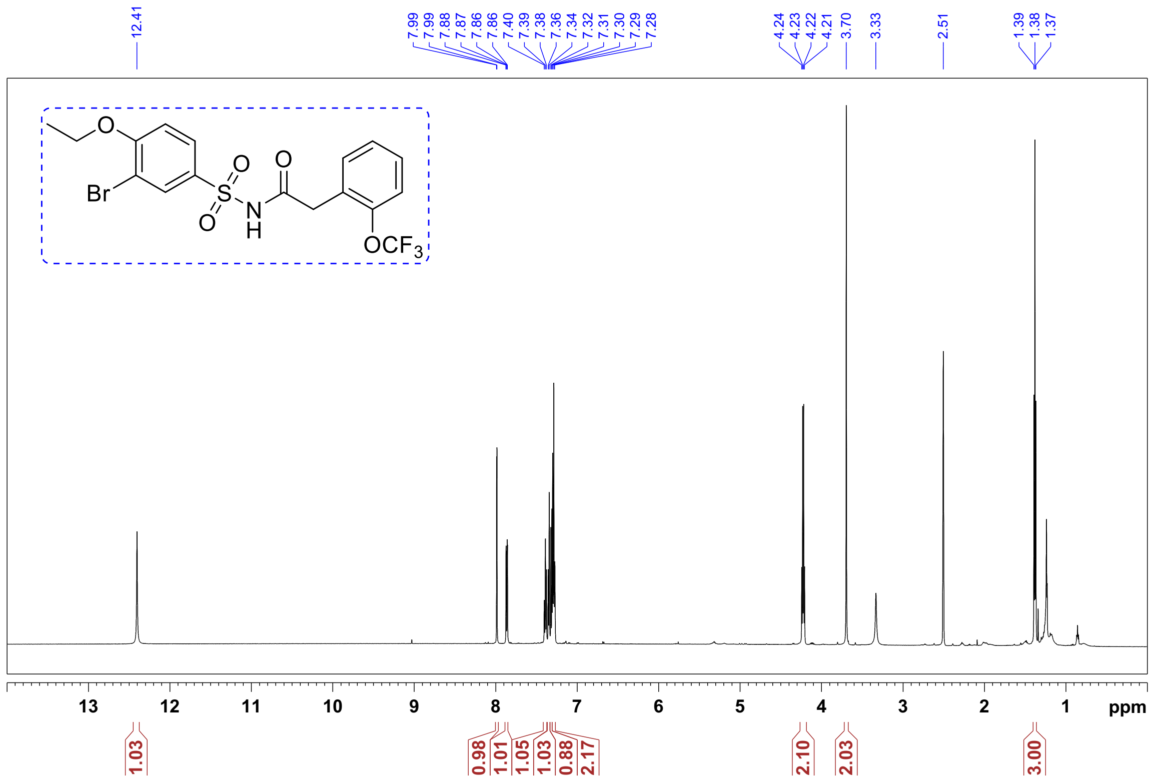


[1]H NMR of **Compound 21**: *N*-((3-bromo-4-ethoxyphenyl)sulfonyl)-2-(*m*-tolyl)acetamide.

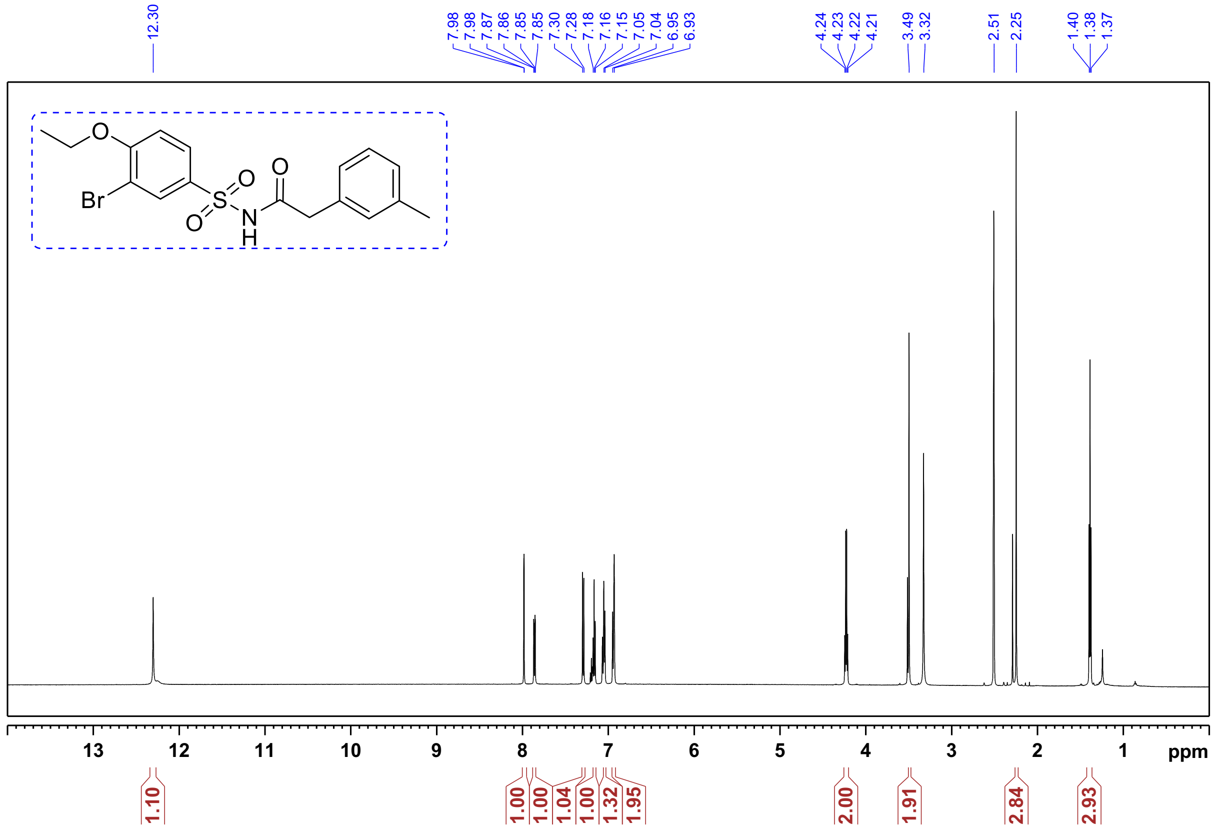

[1]H NMR of **Compound 22:** *N*-((3-bromo-4-ethoxyphenyl)sulfonyl)-4-(2-chloroethyl)benzamide.

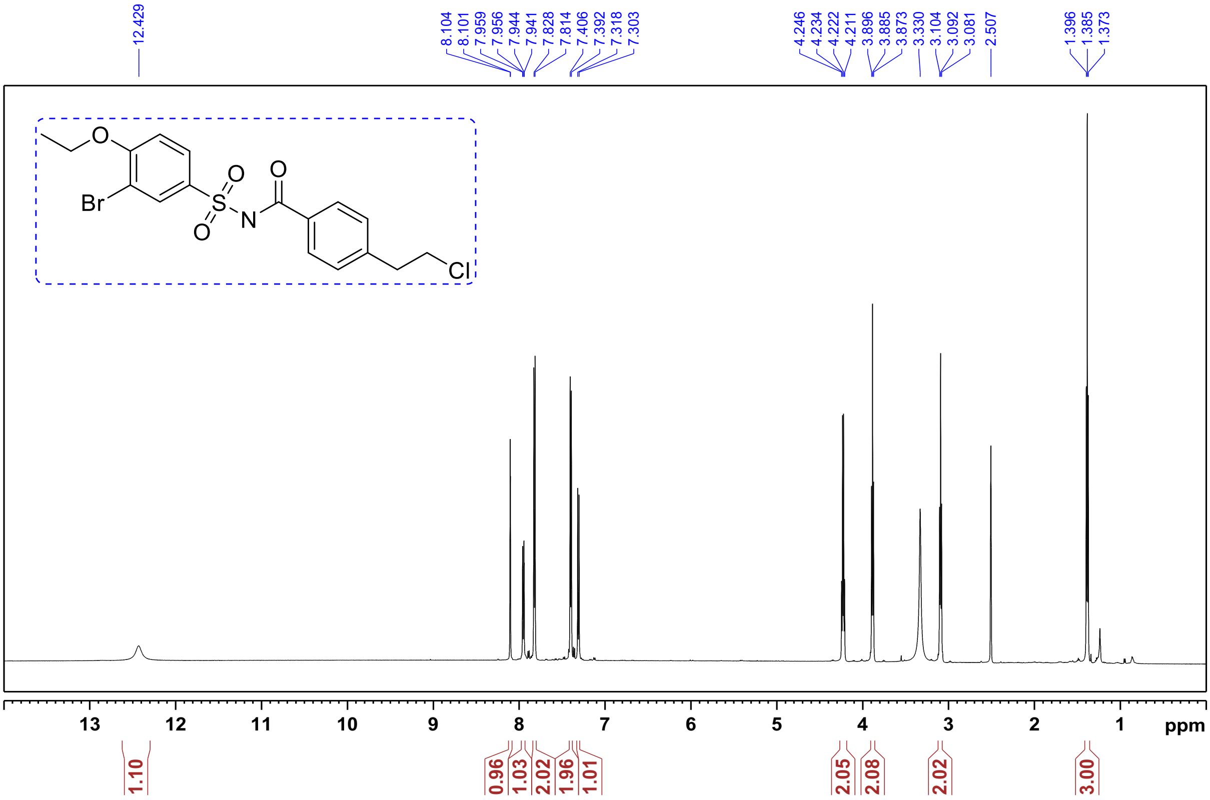

[1]H NMR of **Compound 24:** *N*-((3-bromo-4-ethoxyphenyl)sulfonyl)-2-(3-bromophenyl)acetamide.

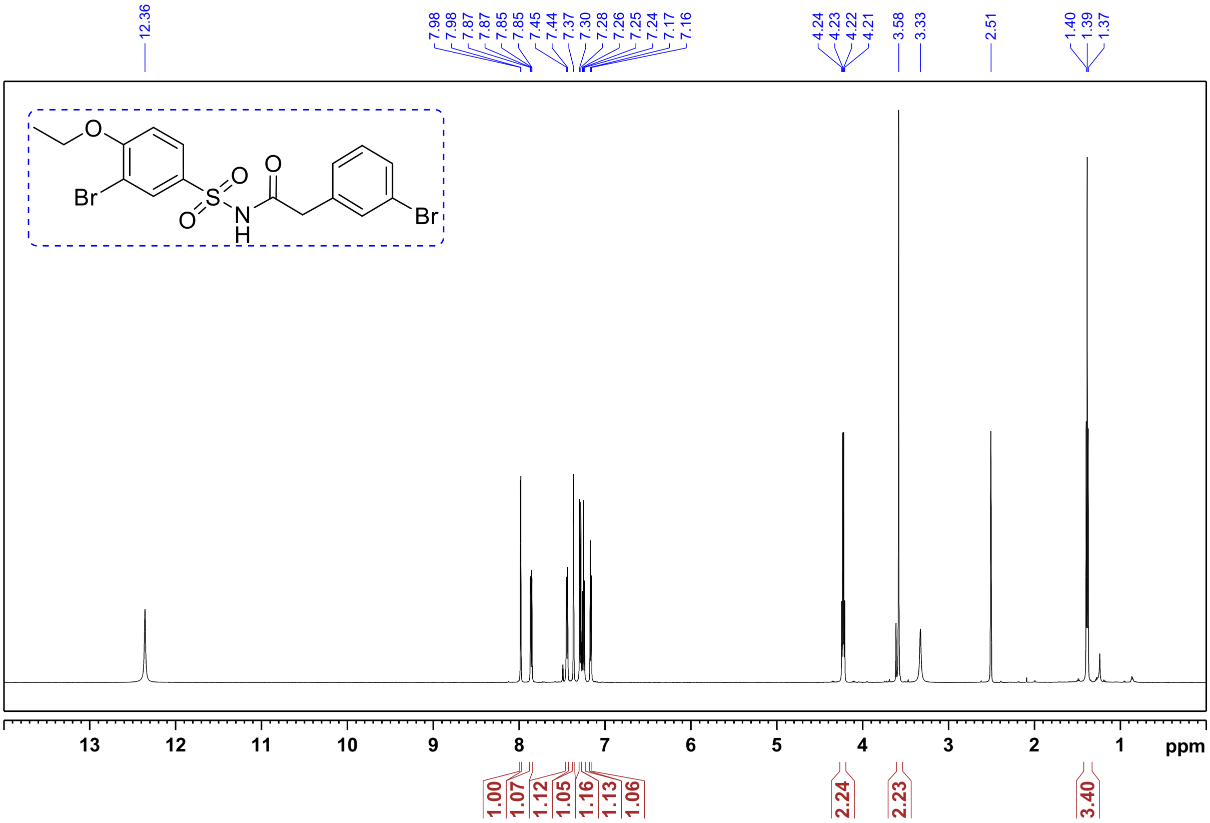

$^1$H NMR of **Compound 23:** *N*-((3-bromo-4-ethoxyphenyl)sulfonyl)-2-(3-chlorophenyl)acetamide.

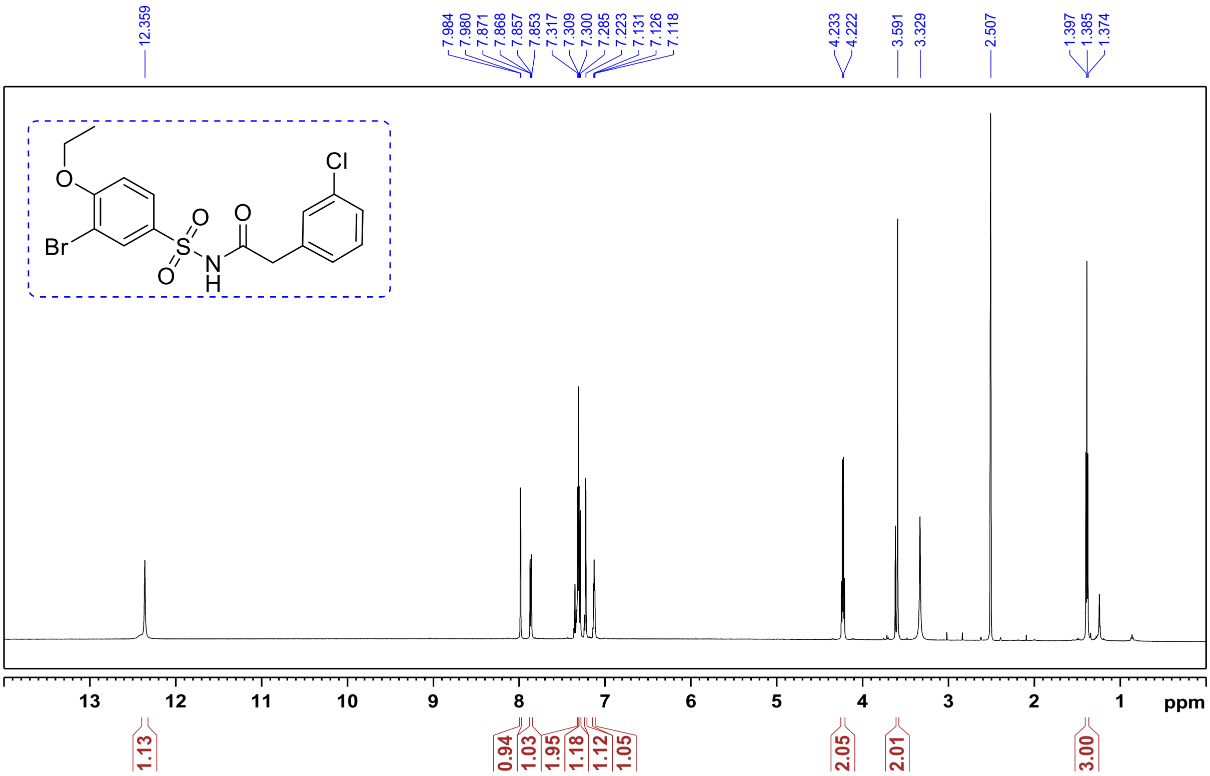


$^1$H NMR of **Compound 25:** *N-(phenylsulfonyl)-2-(p-tolyl)acetamide*

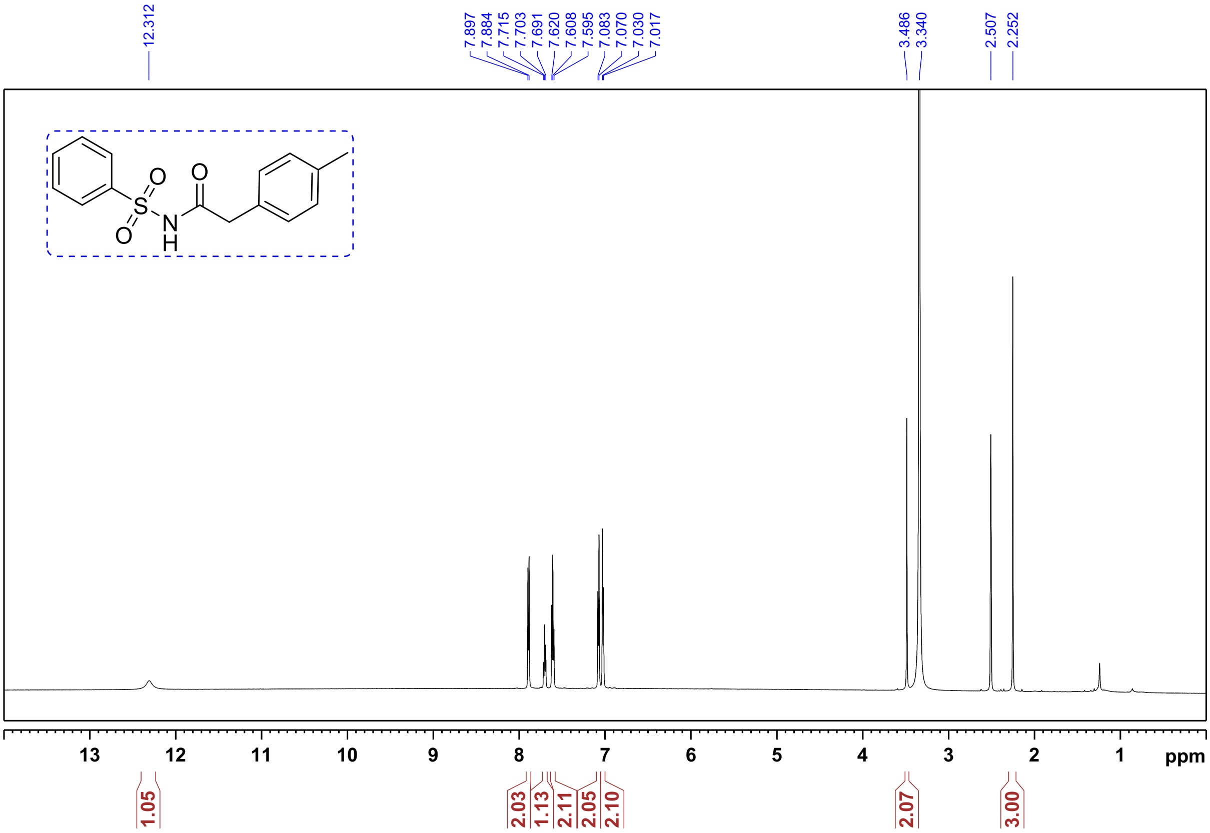

[1]H NMR of **Compound 26:** *(N-((3-bromo-4-ethoxyphenyl)sulfonyl)-2-(p-tolyl)acetamide)*

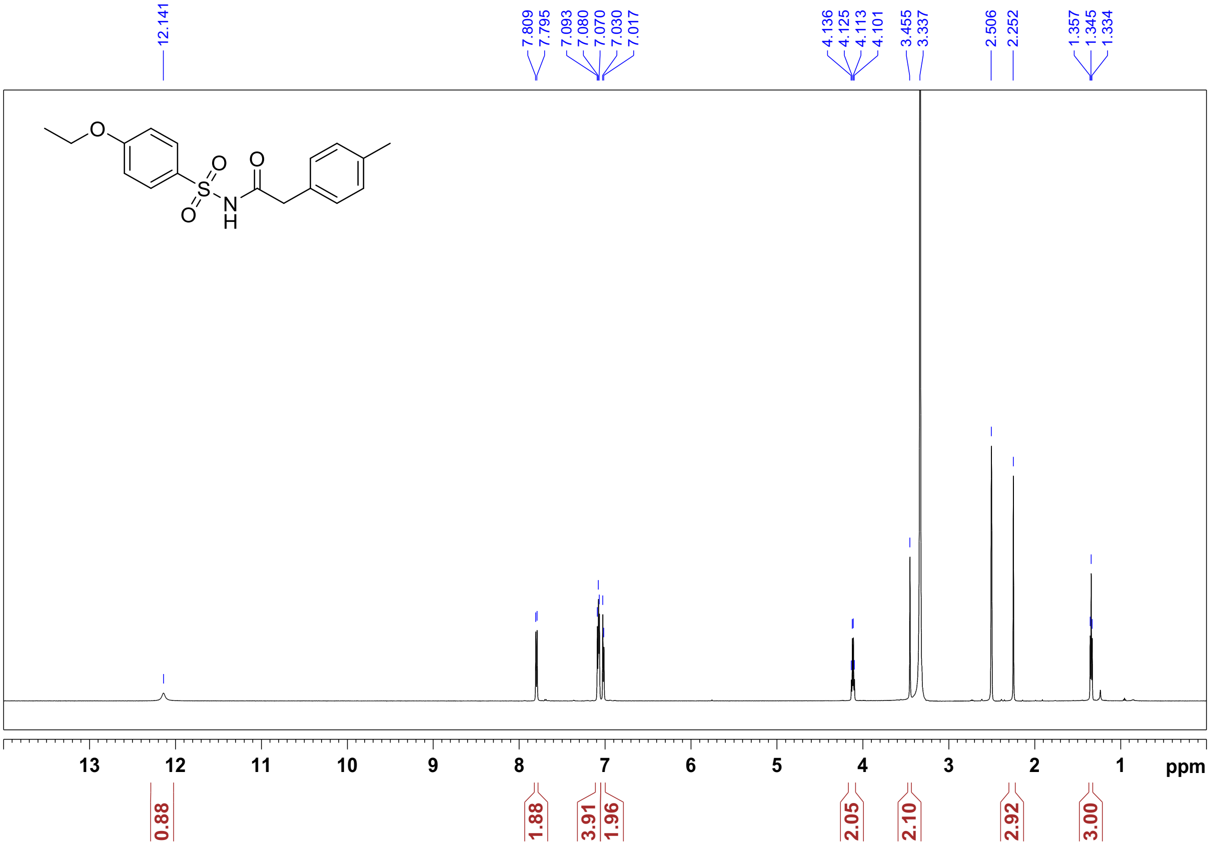


[1]H NMR of **Compound 27:** *N-((4-methoxyphenyl)sulfonyl)-2-(p-tolyl)acetamide*

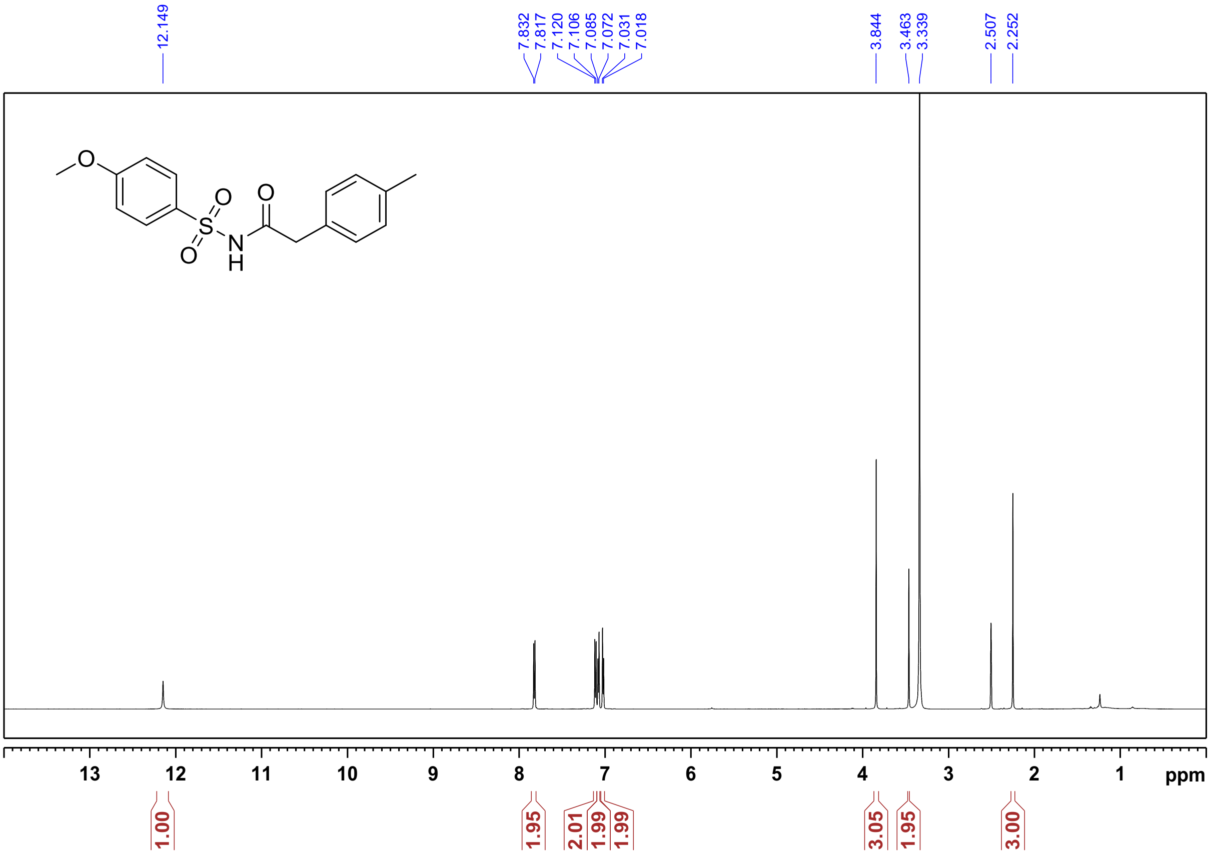

$^1$H NMR of **Compound 28:** *2-(p-tolyl)-N-((4-(trifluoromethoxy)phenyl)sulfonyl)acetamide*

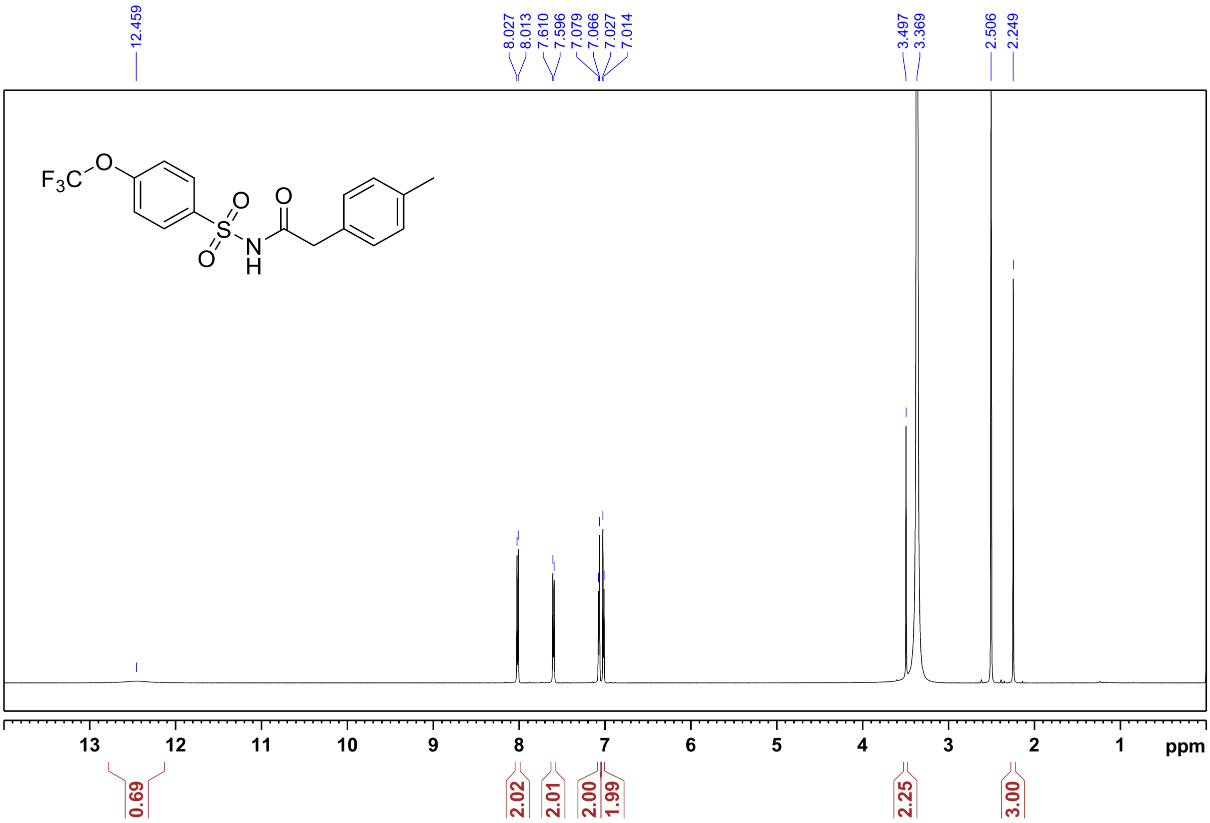


$^1$H NMR of **Compound 29:** *N-((4-(tert-butyl)phenyl)sulfonyl)-2-(p-tolyl)acetamide*

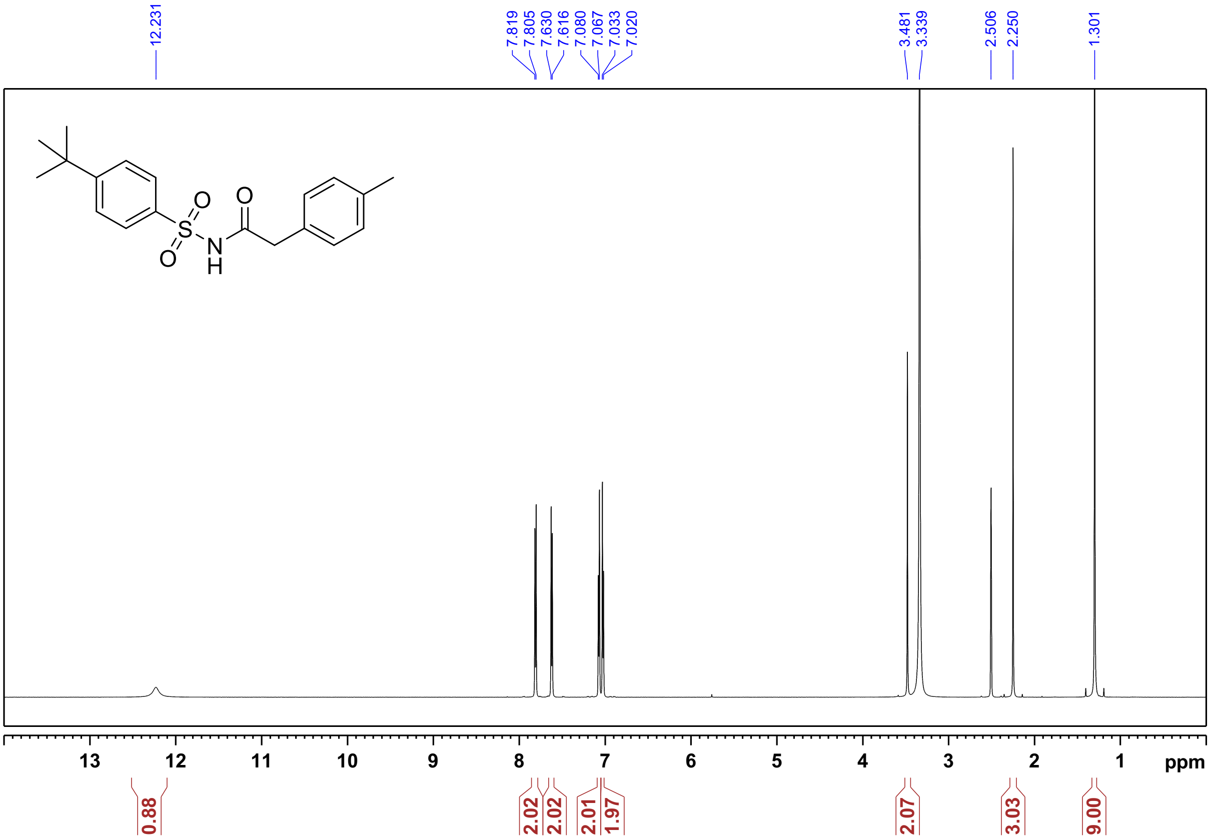

$^1$H NMR of **Compound 30**: *2-(p-tolyl)-N-((4-(trifluoromethyl)phenyl)sulfonyl)acetamide*

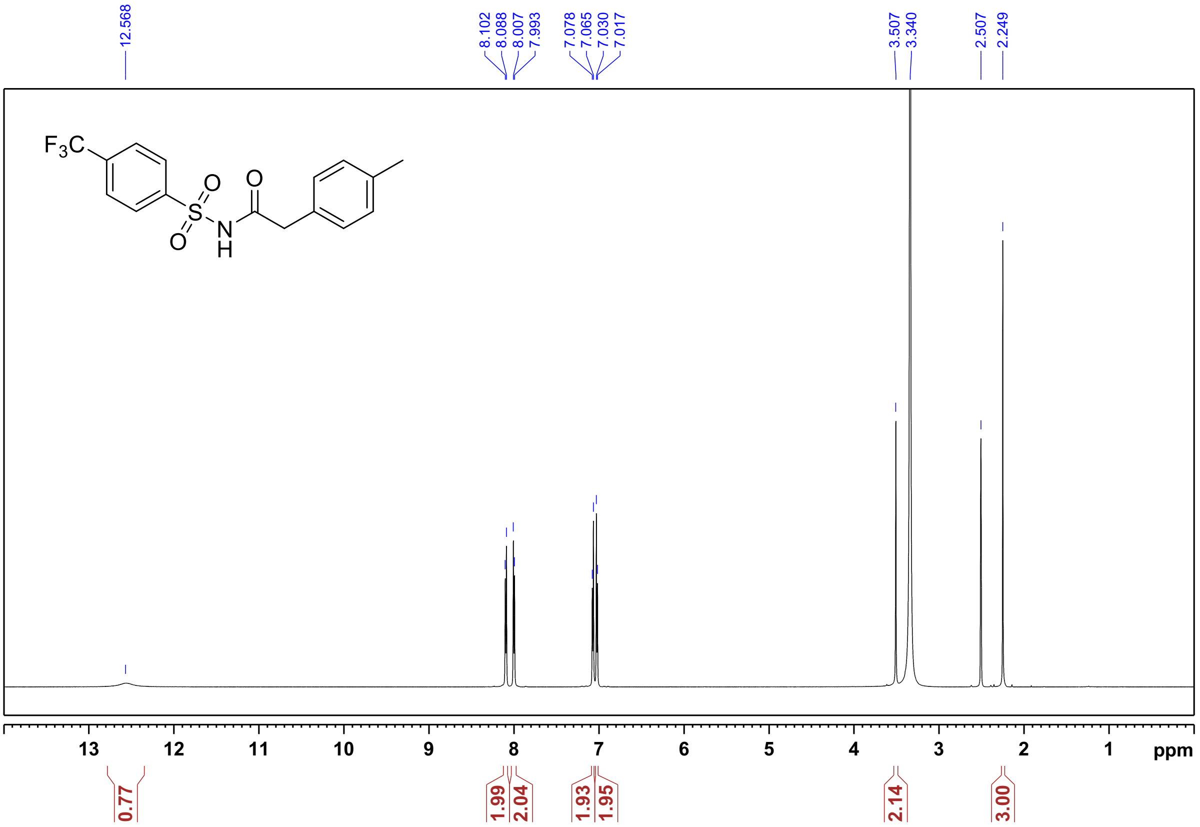


$^1$H NMR of **Compound 32:** *N-((3-bromo-4-methylphenyl)sulfonyl)-2-(p-tolyl)acetamide*

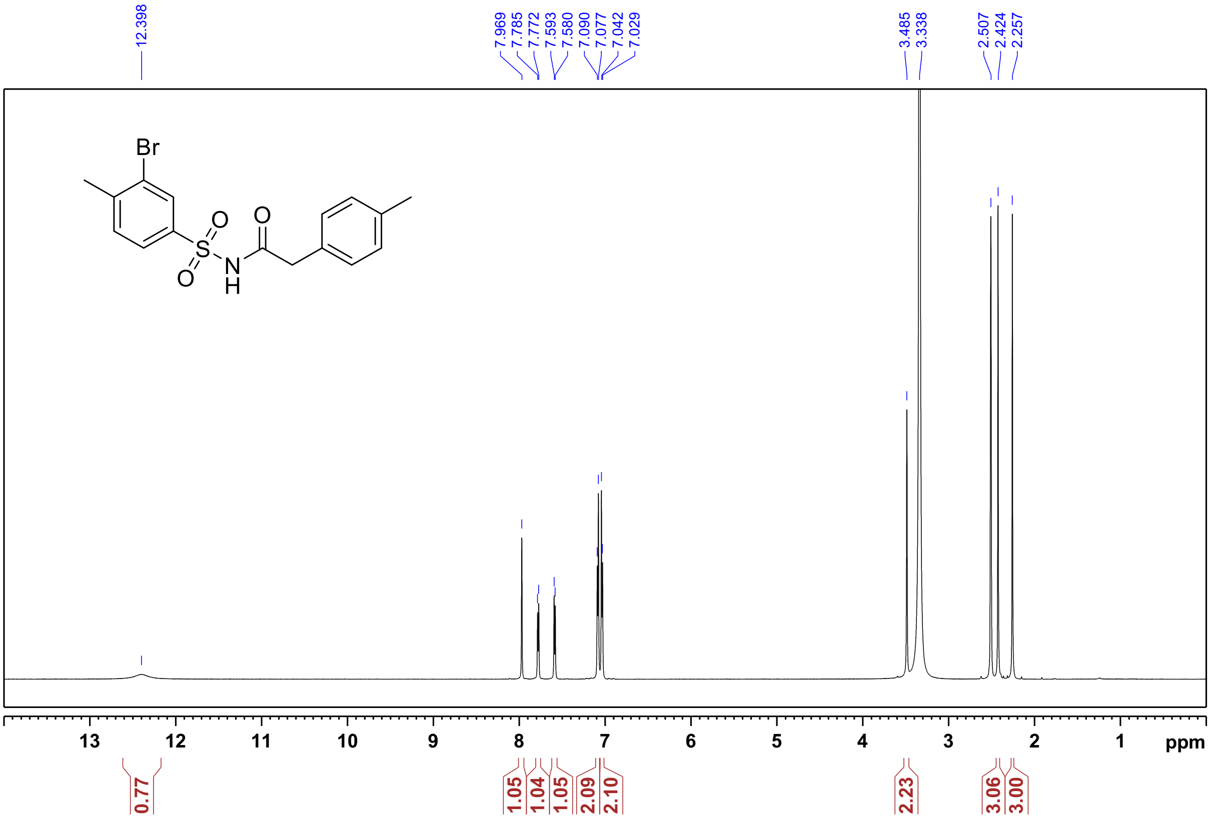

$^{1}$H NMR of **Compound 33**: *N*-((4-bromo-3-fluorophenyl)sulfonyl)-2-(p-tolyl)acetamide:

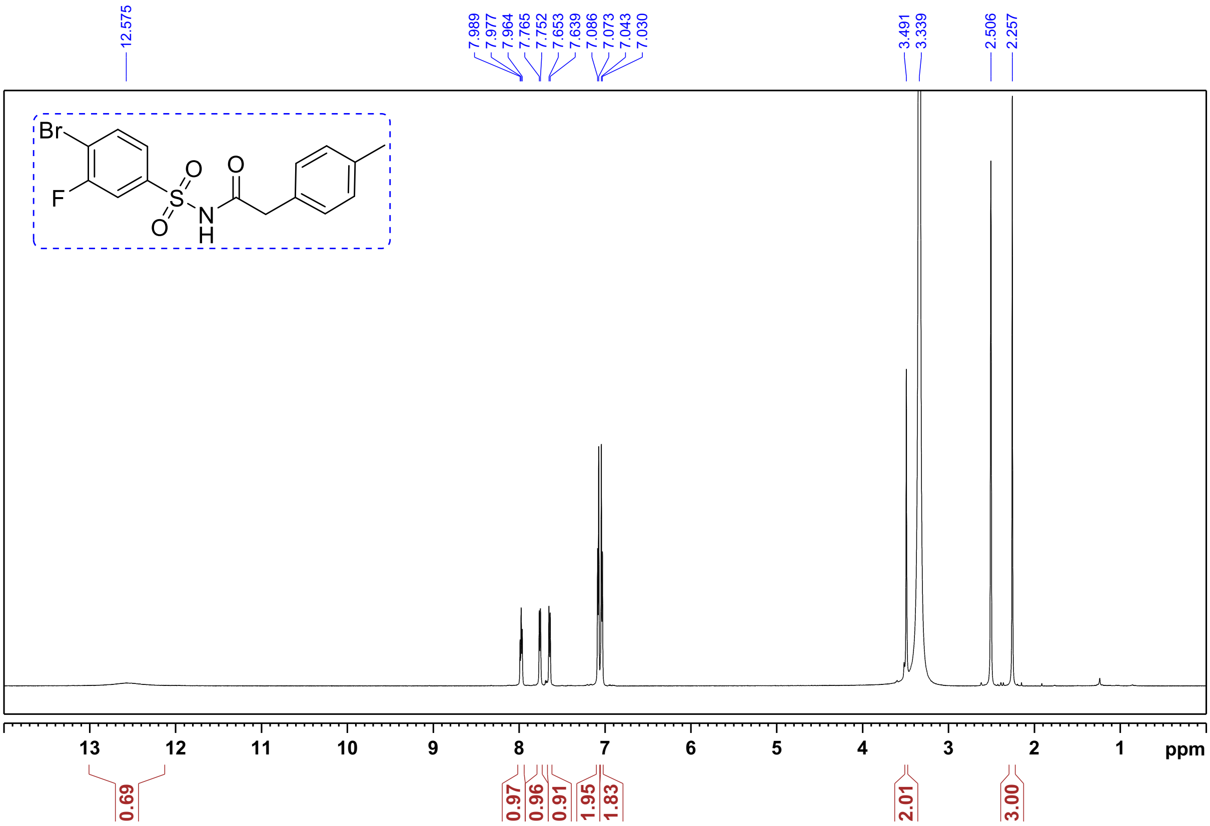

$^{1}$H NMR of **Compound 34:** *N*-((4-ethoxy-3-(1-methyl-1*H*-pyrazol-3-yl)phenyl)sulfonyl)-2-(*p*-tolyl) acetamide.

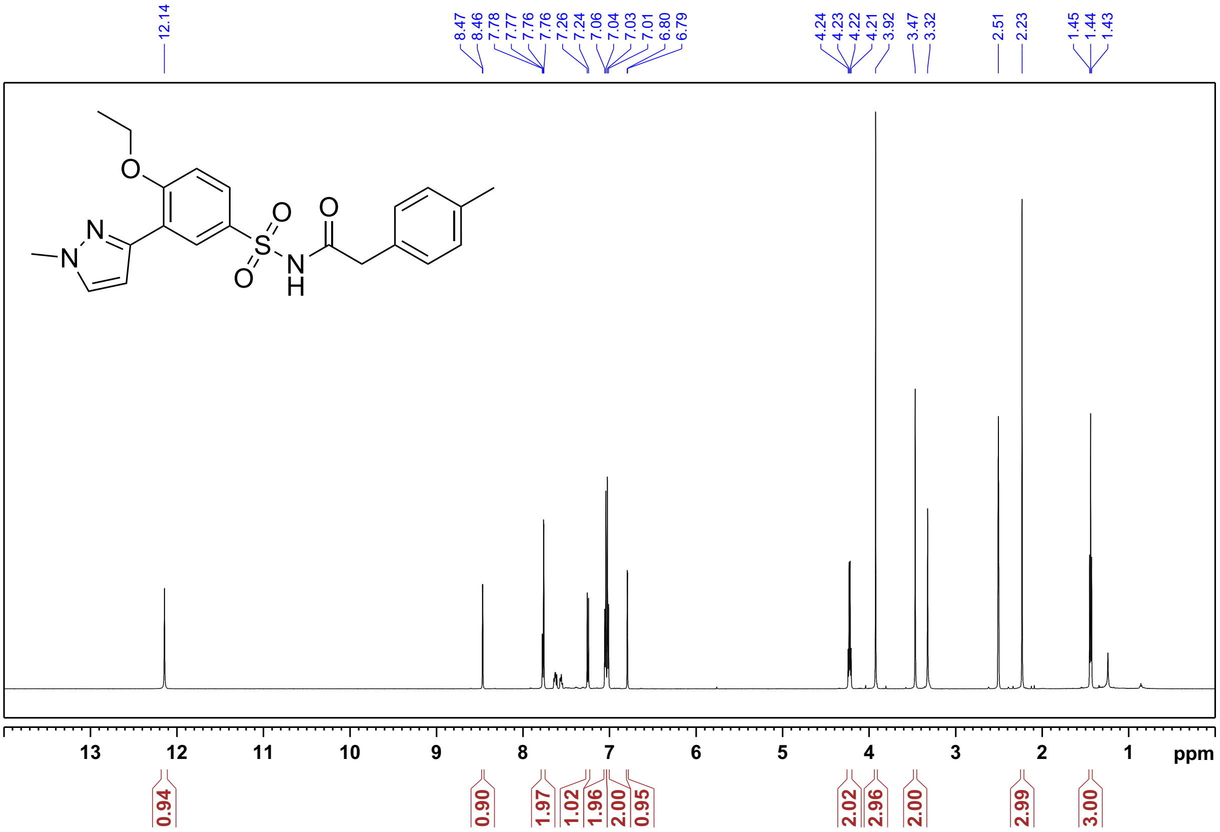

$^{1}$H NMR of **Compound 35**: *N-((5-bromo-2-fluorophenyl)sulfonyl)-2-(p-tolyl)acetamide*

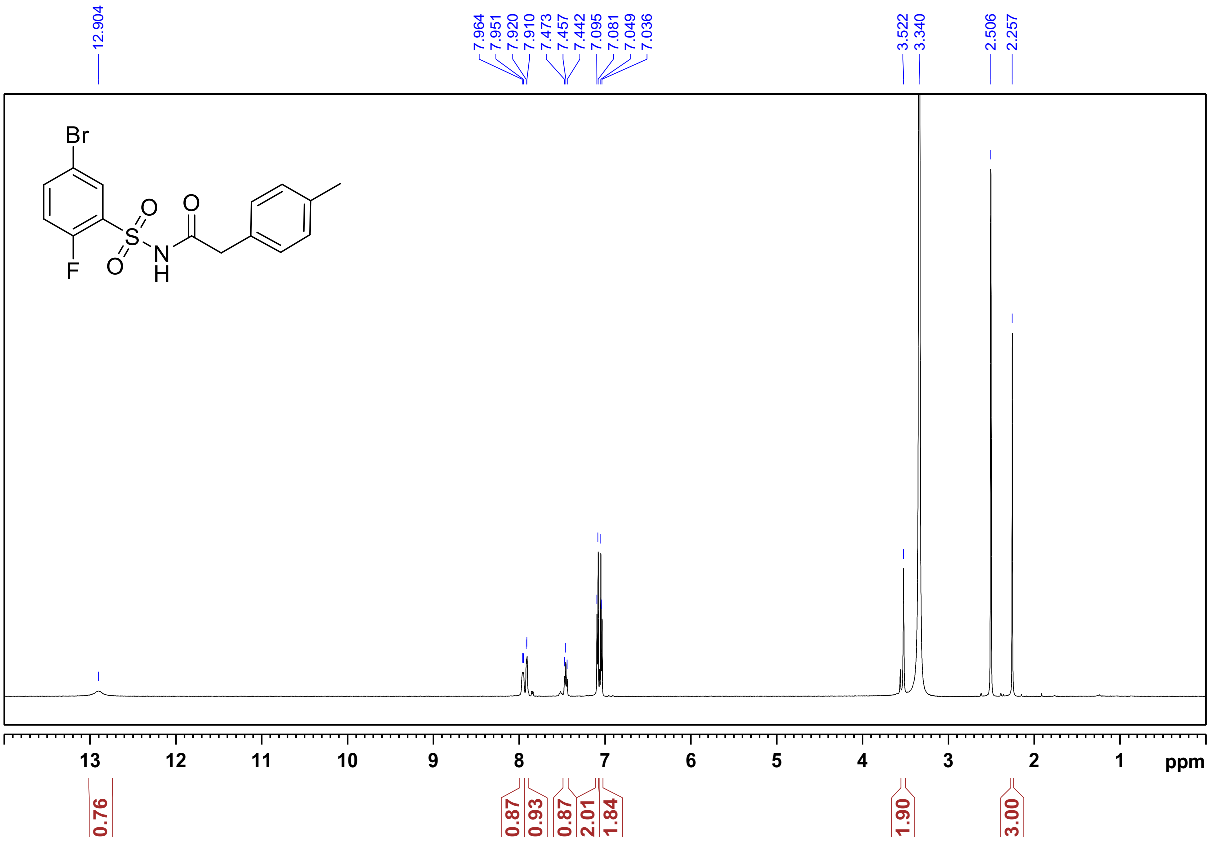


$^{1}$H NMR of **Compound 36**: *N-((2,5-dichlorophenyl)sulfonyl)-2-(p-tolyl)acetamide*

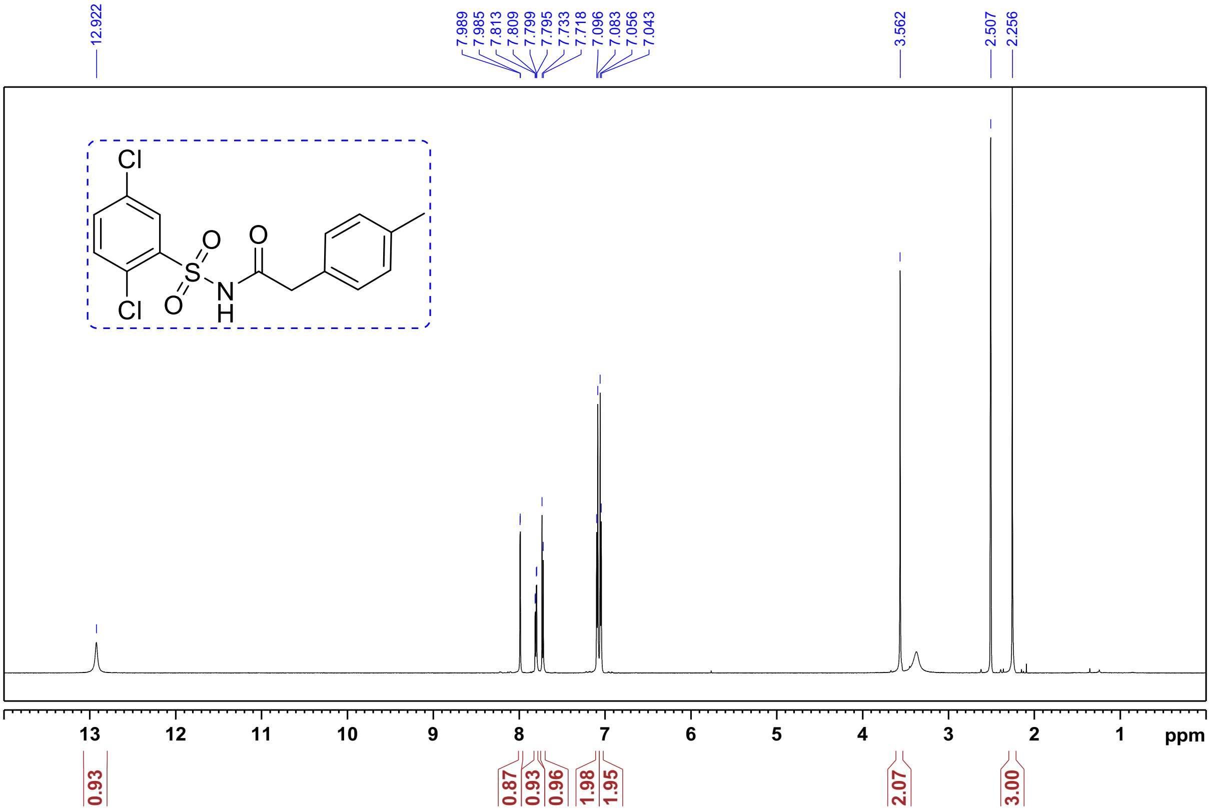

$^{1}H$ NMR of **Compound 37**: *N*-((2-fluoro-5-(trifluoromethyl)phenyl)sulfonyl)-2-(*p*-tolyl)acetamide:

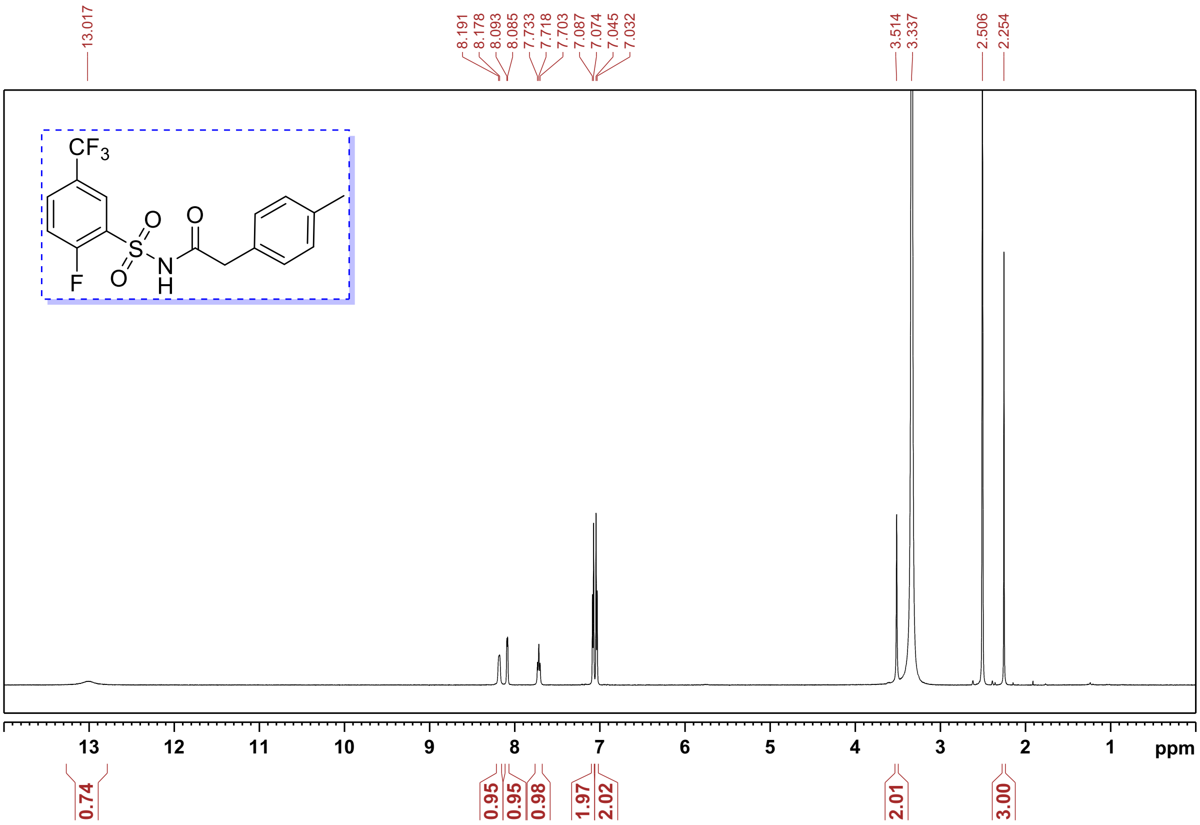


$^{1}H$ NMR of **Compound 38**: *N*-((3-chloro-2-methylphenyl)sulfonyl)-2-(*p*-tolyl)acetamide

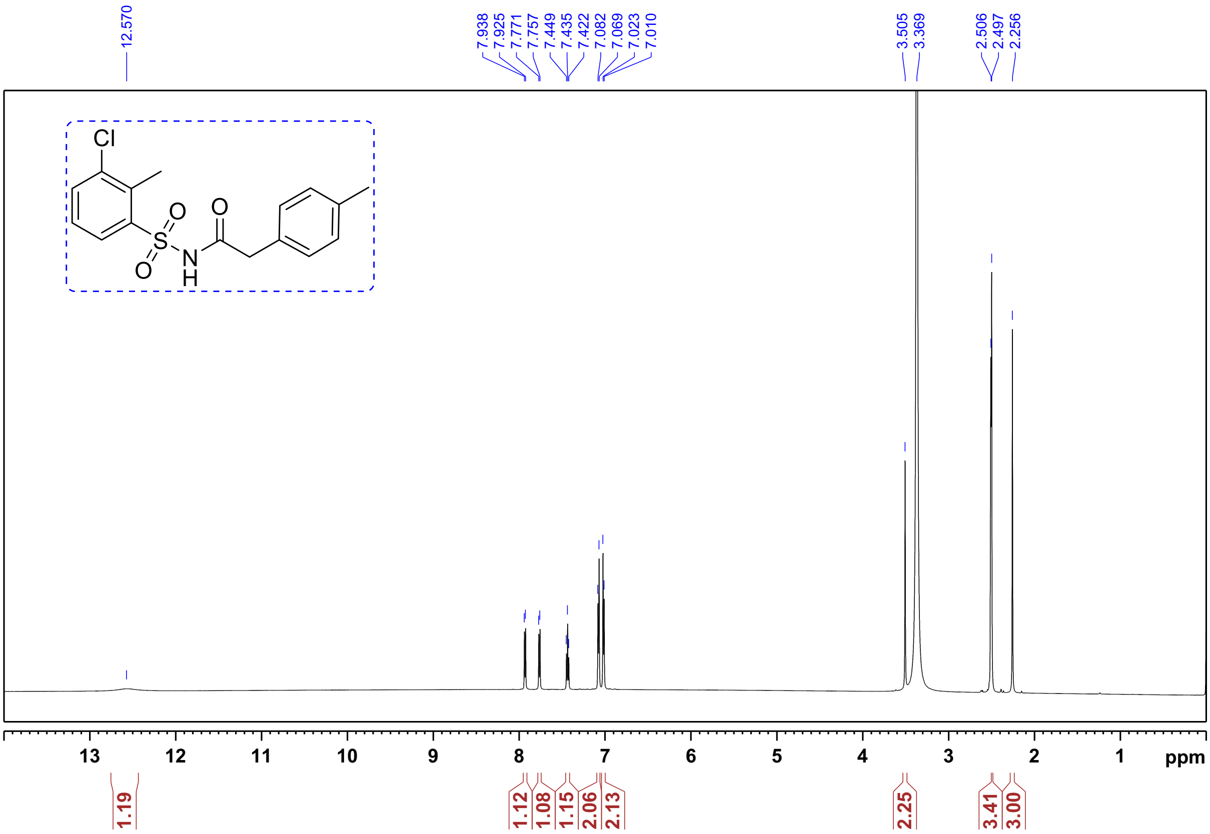

$^1$H NMR of **Compound 41**: *N*-((3-bromo-4-ethoxyphenyl)sulfonyl)-5-methoxyfuran-2-carboxamide:

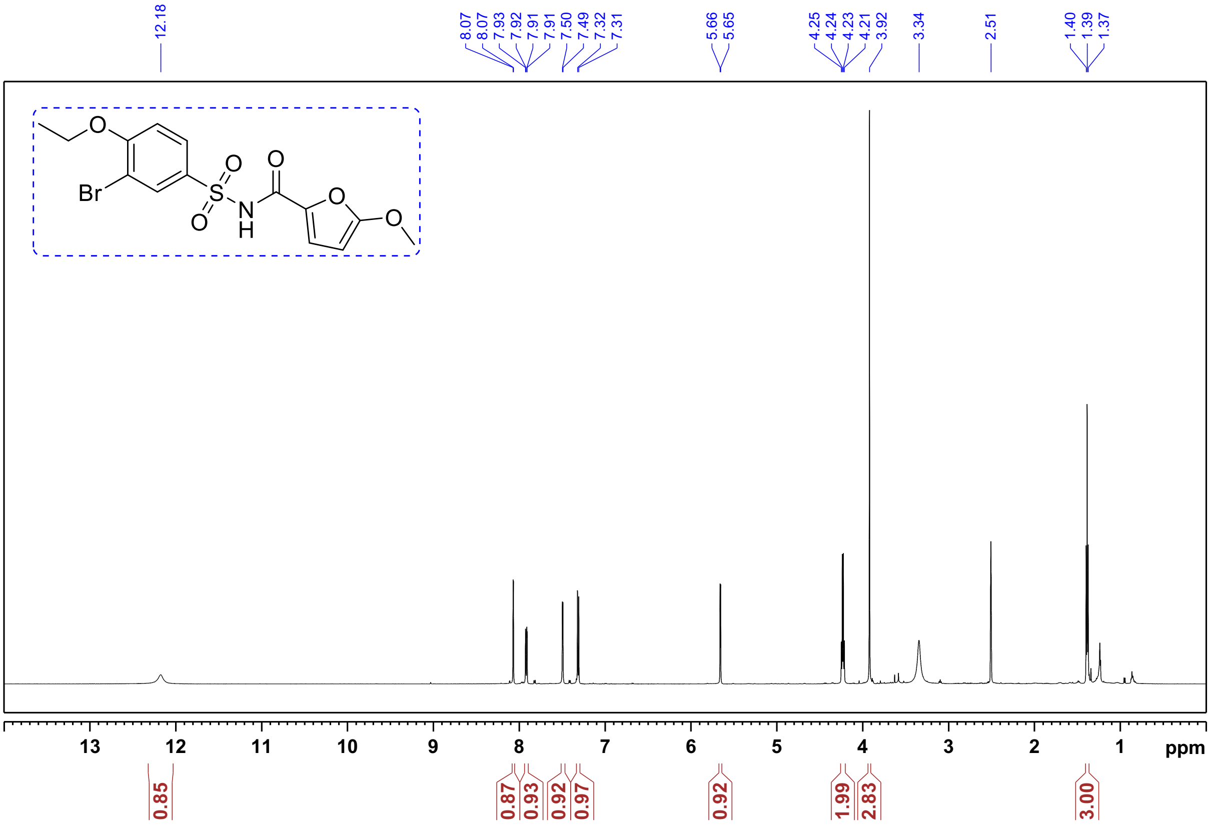

$^1$H NMR of **Compound 42:** *N*-((3-bromo-4-ethoxyphenyl)sulfonyl)-2-phenylpropanamide.

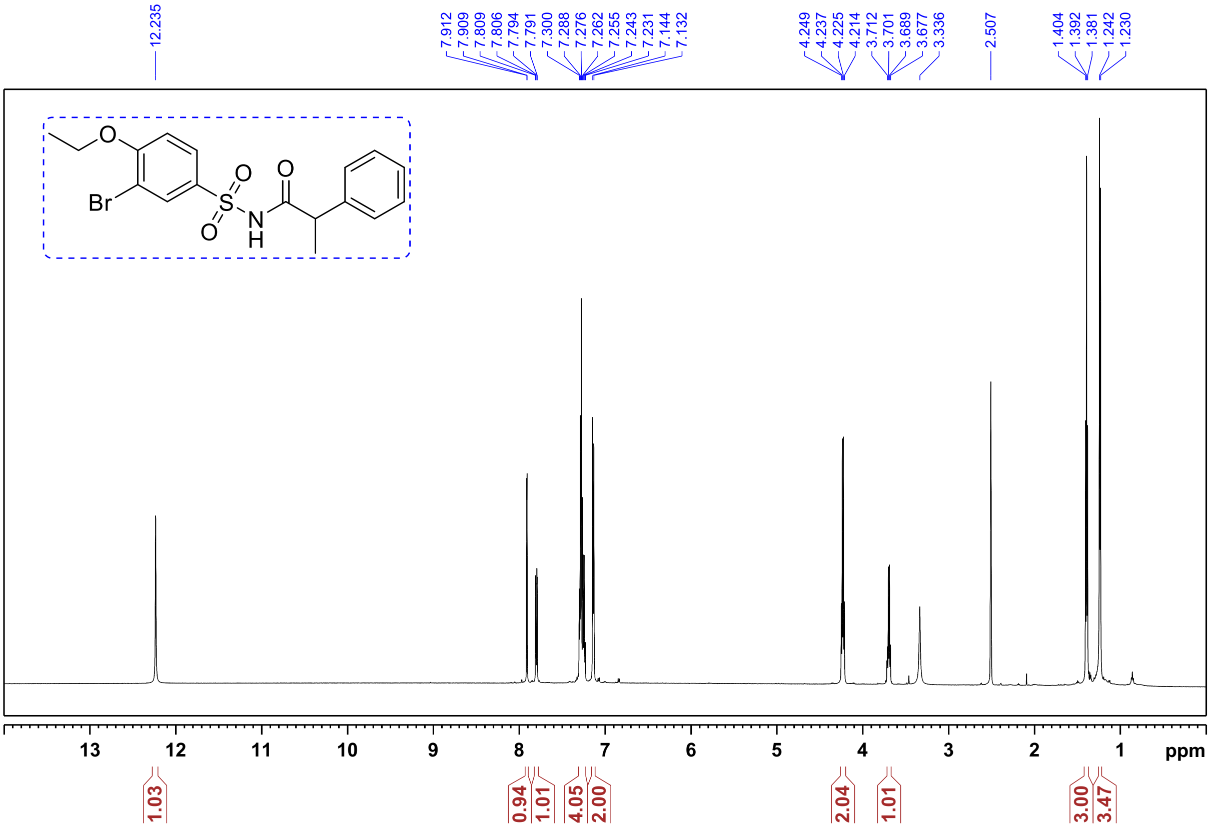

$^1$H NMR of **Compound 47:** *N*-(benzo[*d*]thiazol-6-ylsulfonyl)-2-(*p*-tolyl)acetamide

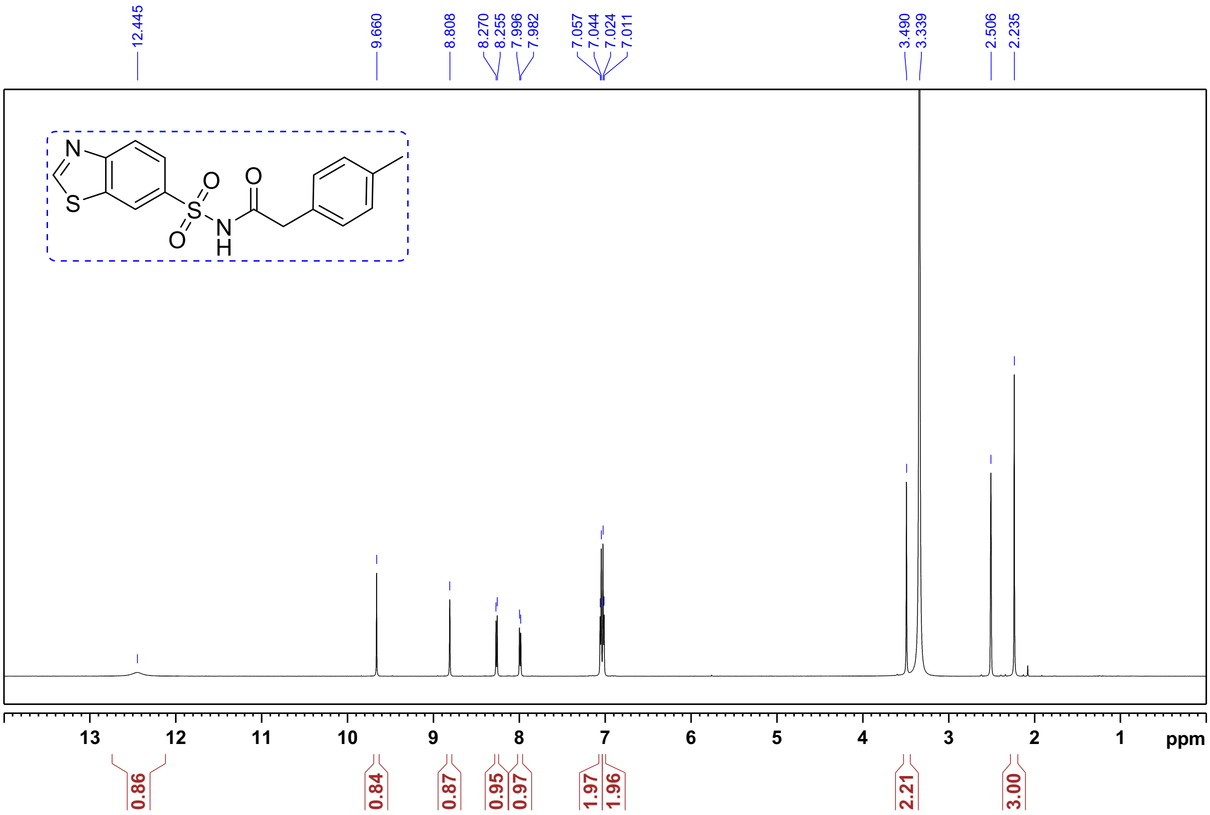

$^1$H NMR of **Compound 48:** *N*-(quinolin-5-ylsulfonyl)-2-(*p*-tolyl)acetamide

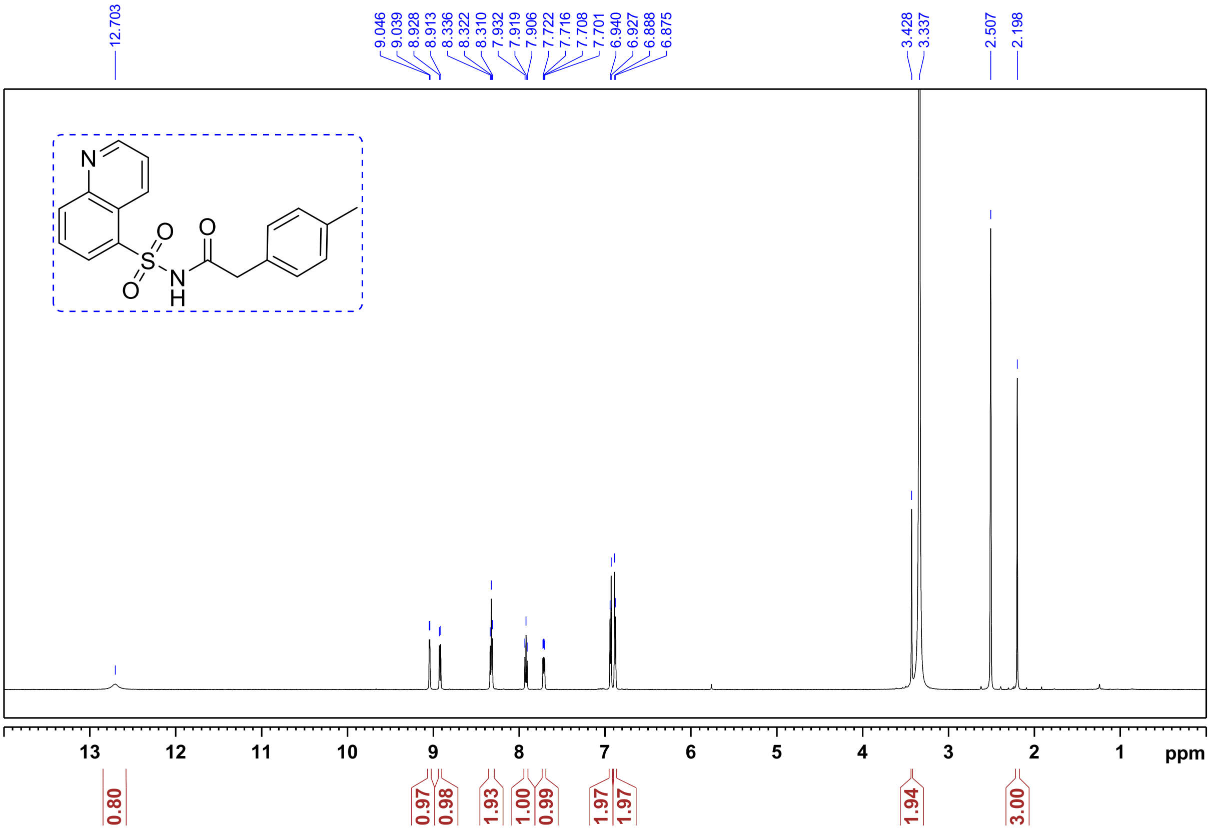

### LC-MS data

**Compound 1:**

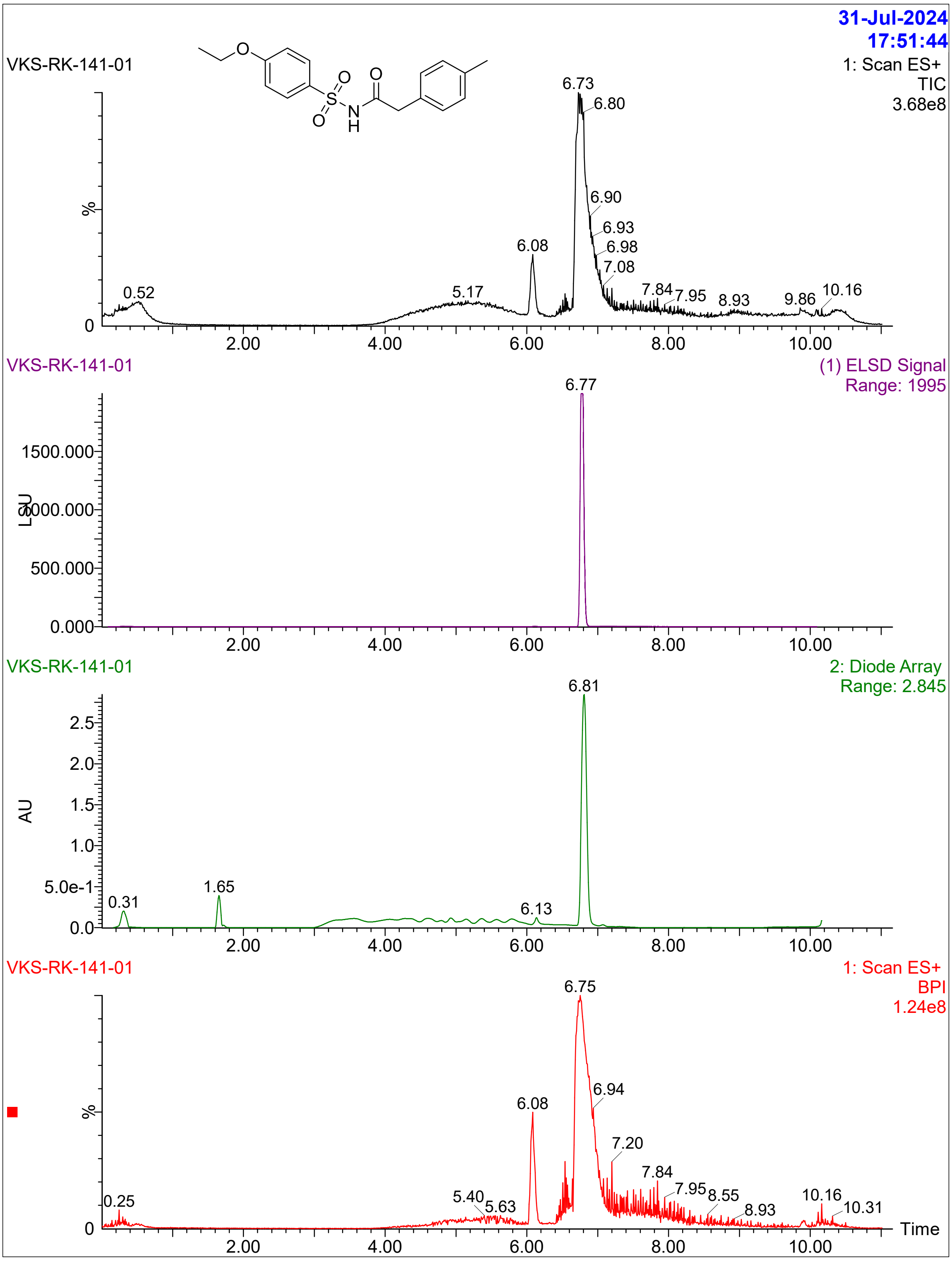

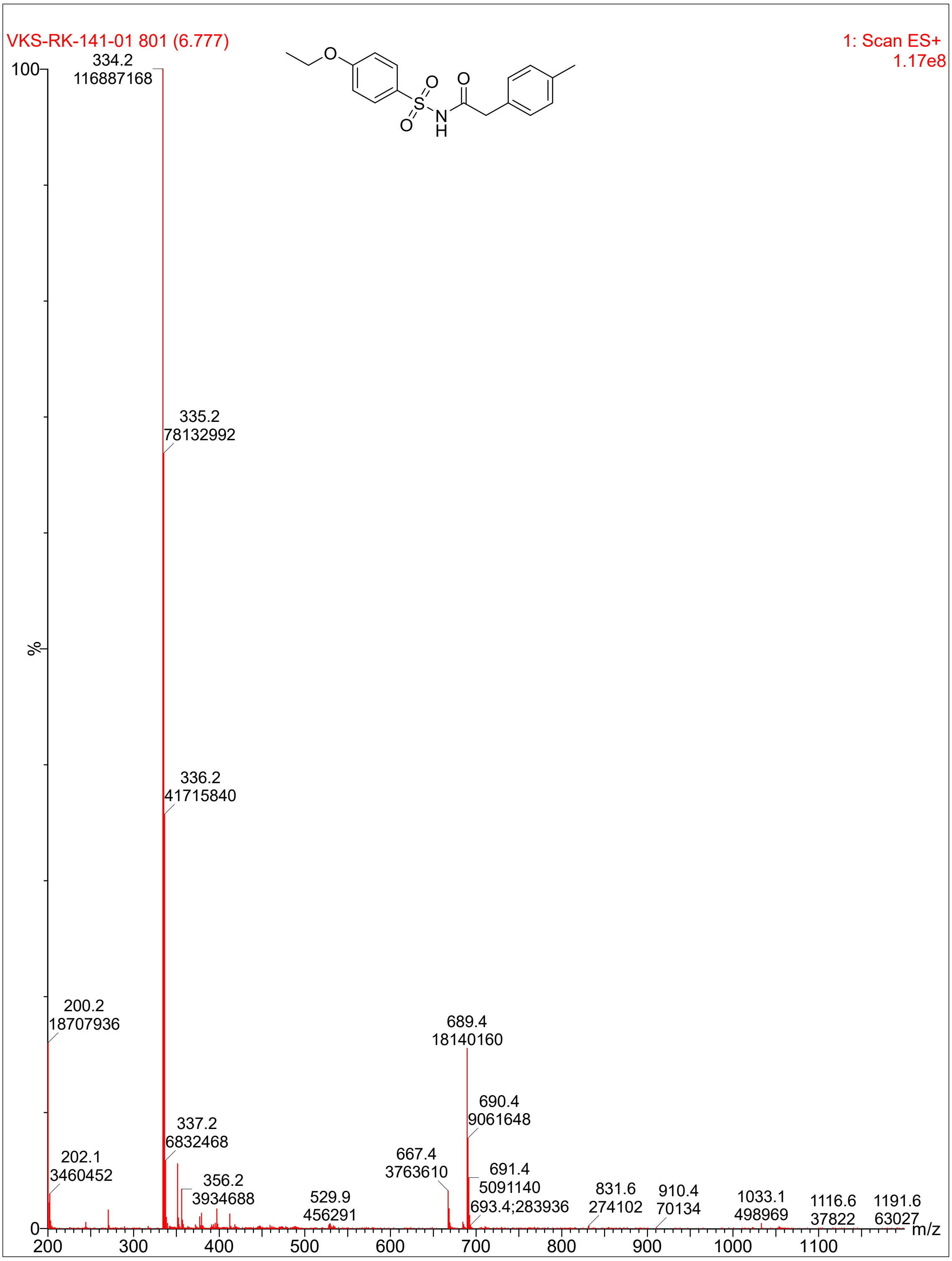

VKS-RK-141-01 801 (6.777)
1: Scan ES+
1.17e8
100
%
0
334.2
116887168
335.2
78132992
336.2
41715840
200.2
18707936
689.4
18140160
690.4
9061648
337.2
6832468
202.1
3460452
667.4
3763610
691.4
5091140
356.2
3934688
693.4;283936
529.9
456291
831.6
274102
910.4
70134
1033.1
498969
1116.6
37822
1191.6
63027
200
300
400
500
600
700
800
900
1000
1100
m/z

**Compound 27:**

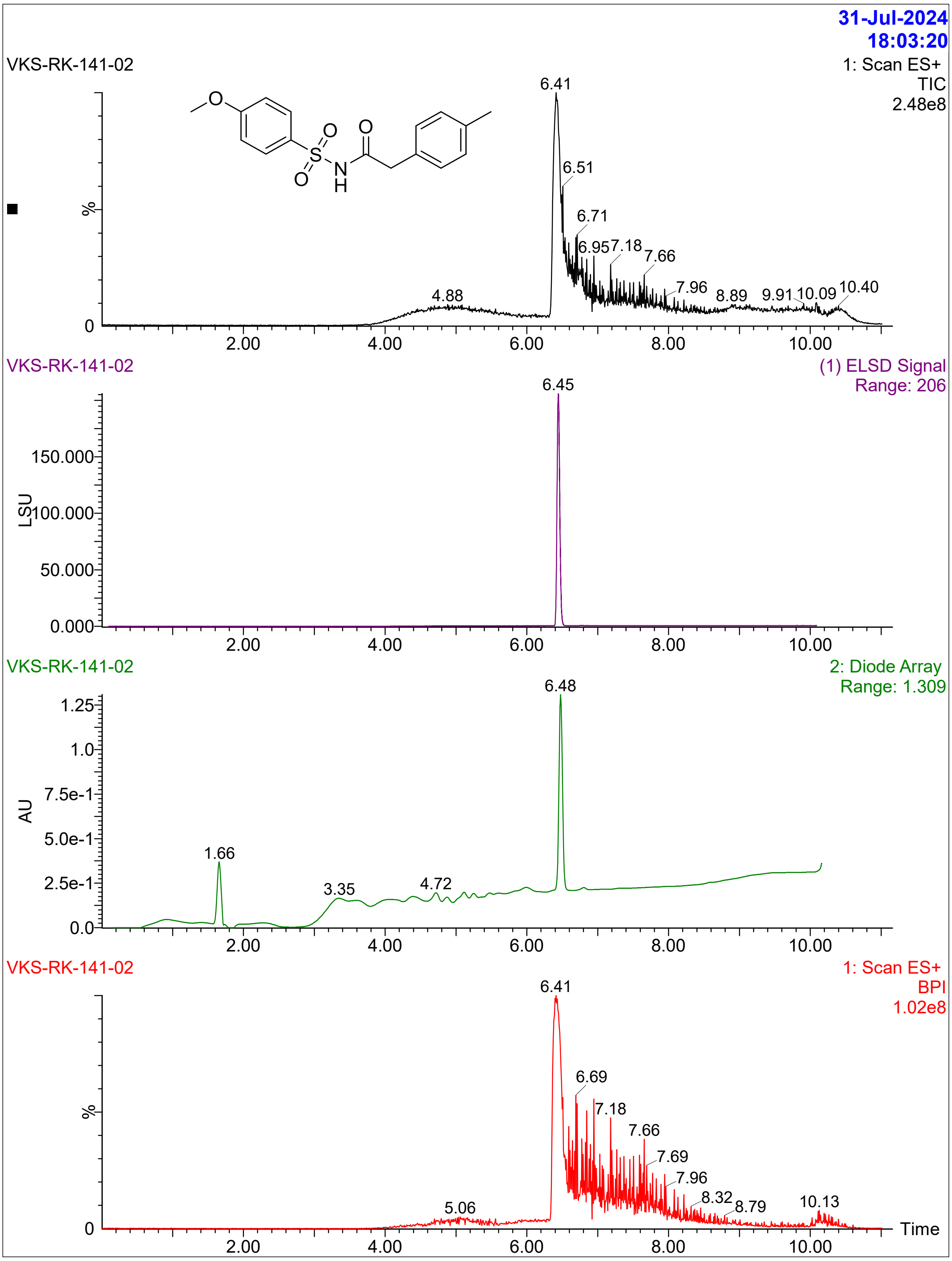
31-Jul-2024
18:03:20
VKS-RK-141-02
1: Scan ES+
TIC
2.48e8
6.41
6.51
6.71
6.95
7.18
7.66
7.96
8.89
9.91
10.09
10.40
4.88
%
0
2.00
4.00
6.00
8.00
10.00
VKS-RK-141-02
(1) ELSD Signal
Range: 206
6.45
150.000
100.000
50.000
0.000
LSU
VKS-RK-141-02
2: Diode Array
Range: 1.309
6.48
1.25
1.0
7.5e-1
5.0e-1
2.5e-1
0.0
AU
1.66
3.35
4.72
VKS-RK-141-02
1: Scan ES+
BPI
1.02e8
6.41
6.69
7.18
7.66
7.69
7.96
8.32
8.79
10.13
5.06
Time

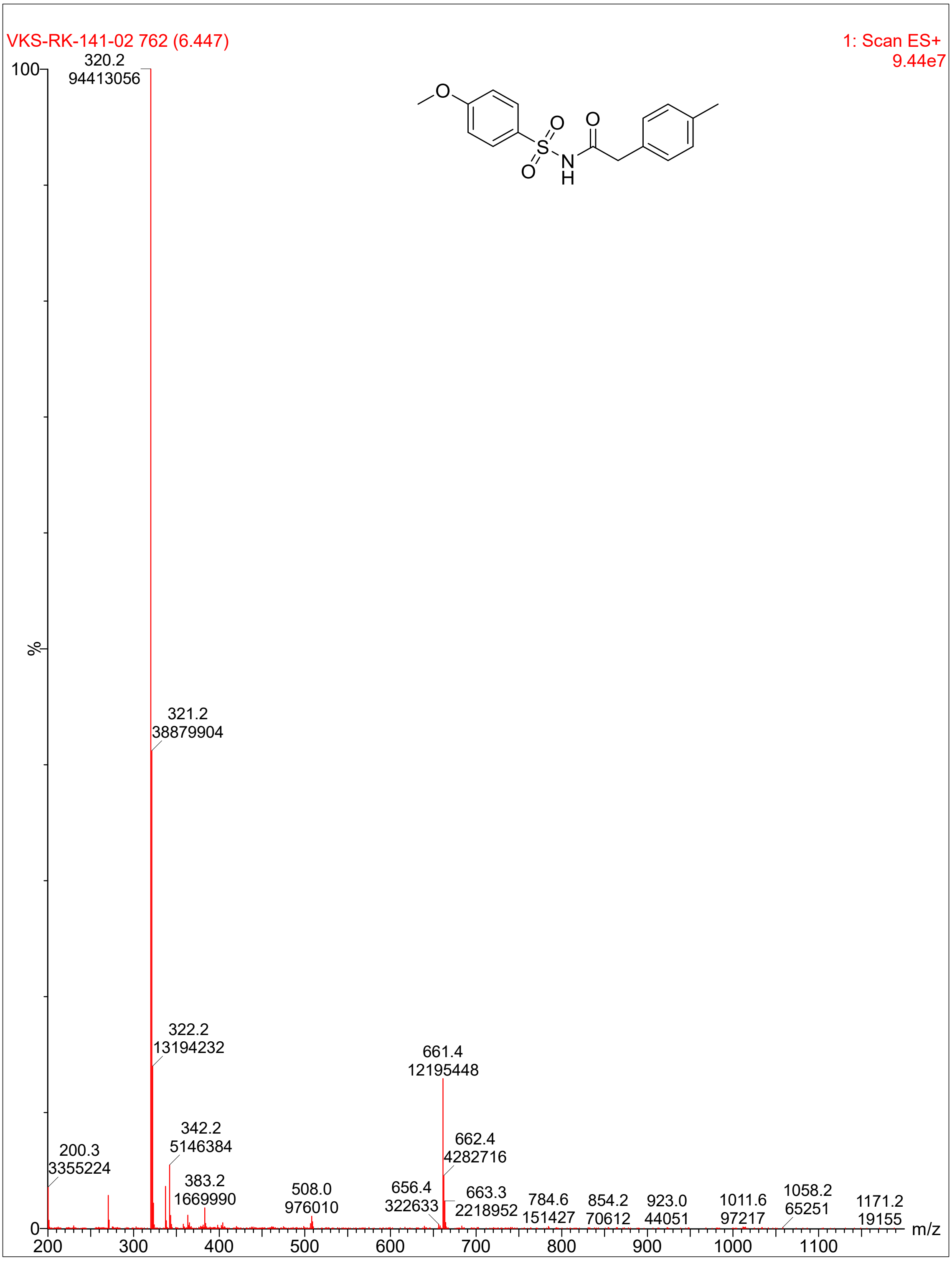
VKS-RK-141-02 762 (6.447)
1: Scan ES+
9.44e7
320.2
94413056
321.2
38879904
322.2
13194232
661.4
12195448
342.2
5146384
662.4
4282716
200.3
3355224
383.2
1669990
508.0
976010
656.4
322633
663.3
2218952
784.6
151427
854.2
70612
923.0
44051
1011.6
97217
1058.2
65251
1171.2
19155
%
m/z
100
0
200
300
400
500
600
700
800
900
1000
1100

**Compound 25:**

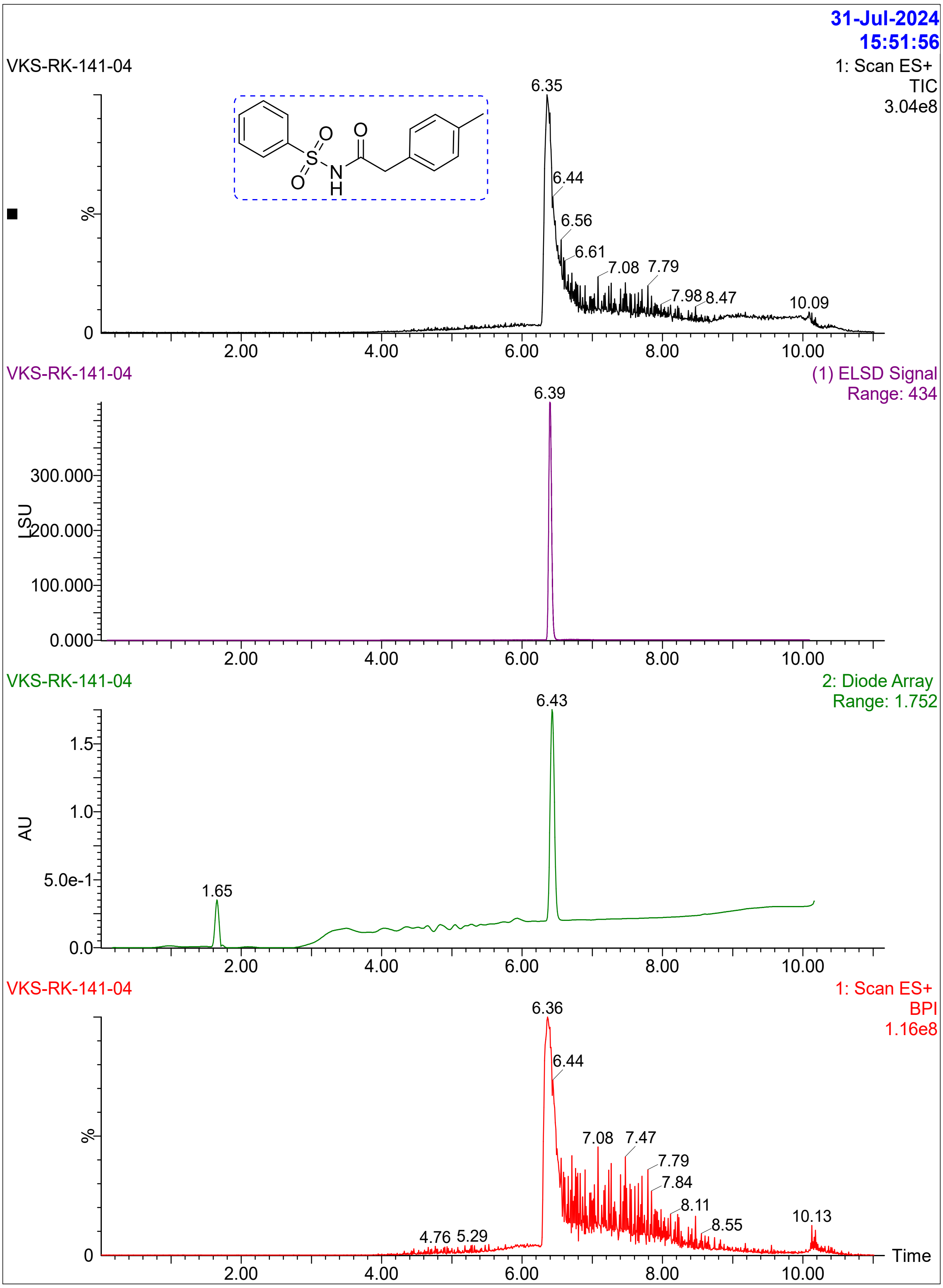

31-Jul-2024
15:51:56
VKS-RK-141-04
1: Scan ES+
TIC
3.04e8
6.35
6.44
6.56
6.61
7.08
7.79
7.98
8.47
10.09
%
0
2.00
4.00
6.00
8.00
10.00
VKS-RK-141-04
(1) ELSD Signal
Range: 434
6.39
300.000
200.000
100.000
0.000
LSU
VKS-RK-141-04
2: Diode Array
Range: 1.752
6.43
1.65
1.5
1.0
5.0e-1
0.0
AU
VKS-RK-141-04
1: Scan ES+
BPI
1.16e8
6.36
6.44
7.08
7.47
7.79
7.84
8.11
8.55
10.13
4.76
5.29
Time

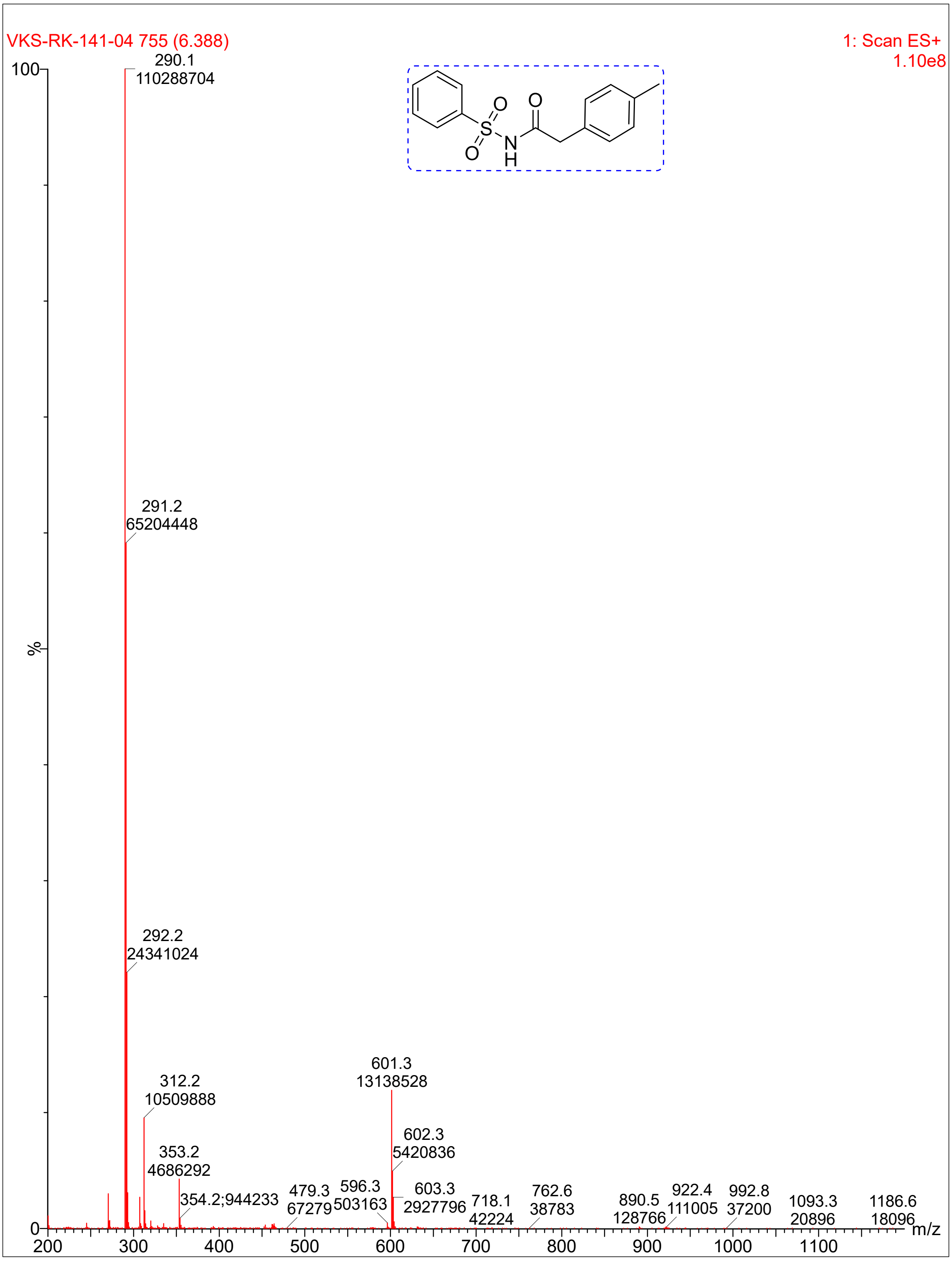

VKS-RK-141-04 755 (6.388)
1: Scan ES+
1.10e8
290.1
110288704
291.2
65204448
292.2
24341024
312.2
10509888
353.2
4686292
354.2;944233
479.3
67279
596.3
503163
601.3
13138528
602.3
5420836
603.3
2927796
718.1
42224
762.6
38783
890.5
128766
922.4
111005
992.8
37200
1093.3
20896
1186.6
18096
%
m/z
100
0
200
300
400
500
600
700
800
900
1000
1100
O
S
N
H

**Compound 28:**

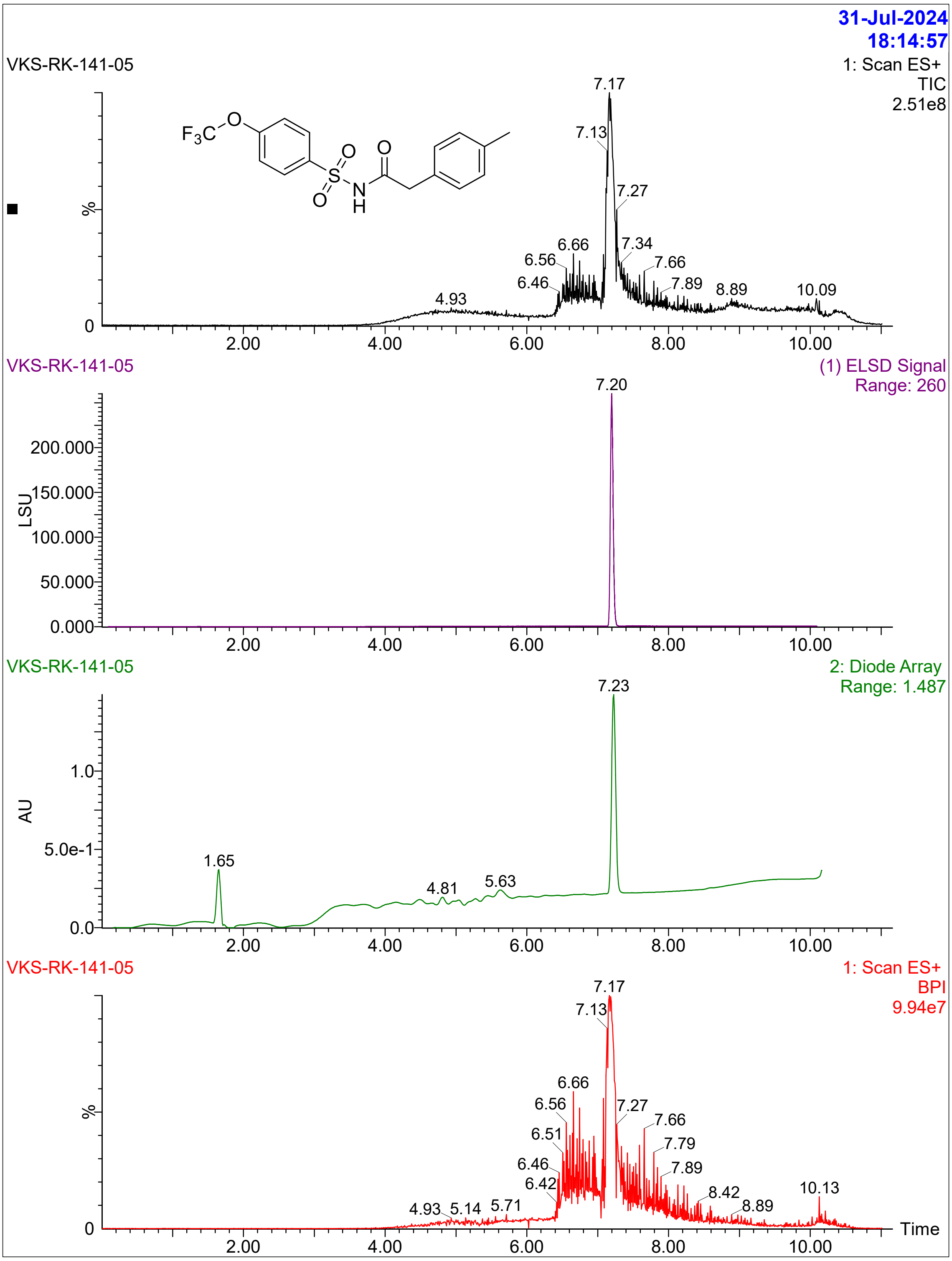

31-Jul-2024
18:14:57
VKS-RK-141-05
1: Scan ES+
TIC
2.51e8
%
7.17
7.13
7.27
6.66
6.56
7.34
7.66
6.46
7.89
8.89
10.09
4.93
2.00
4.00
6.00
8.00
10.00
F3C
O
O
S
O
N
H
O
VKS-RK-141-05
(1) ELSD Signal
Range: 260
7.20
LSU
200.000
150.000
100.000
50.000
0.000
VKS-RK-141-05
2: Diode Array
Range: 1.487
7.23
AU
1.0
5.0e-1
0.0
1.65
4.81
5.63
VKS-RK-141-05
1: Scan ES+
BPI
9.94e7
7.17
7.13
6.66
6.56
7.27
7.66
6.51
7.79
6.46
7.89
6.42
8.42
10.13
8.89
4.93
5.14
5.71
Time

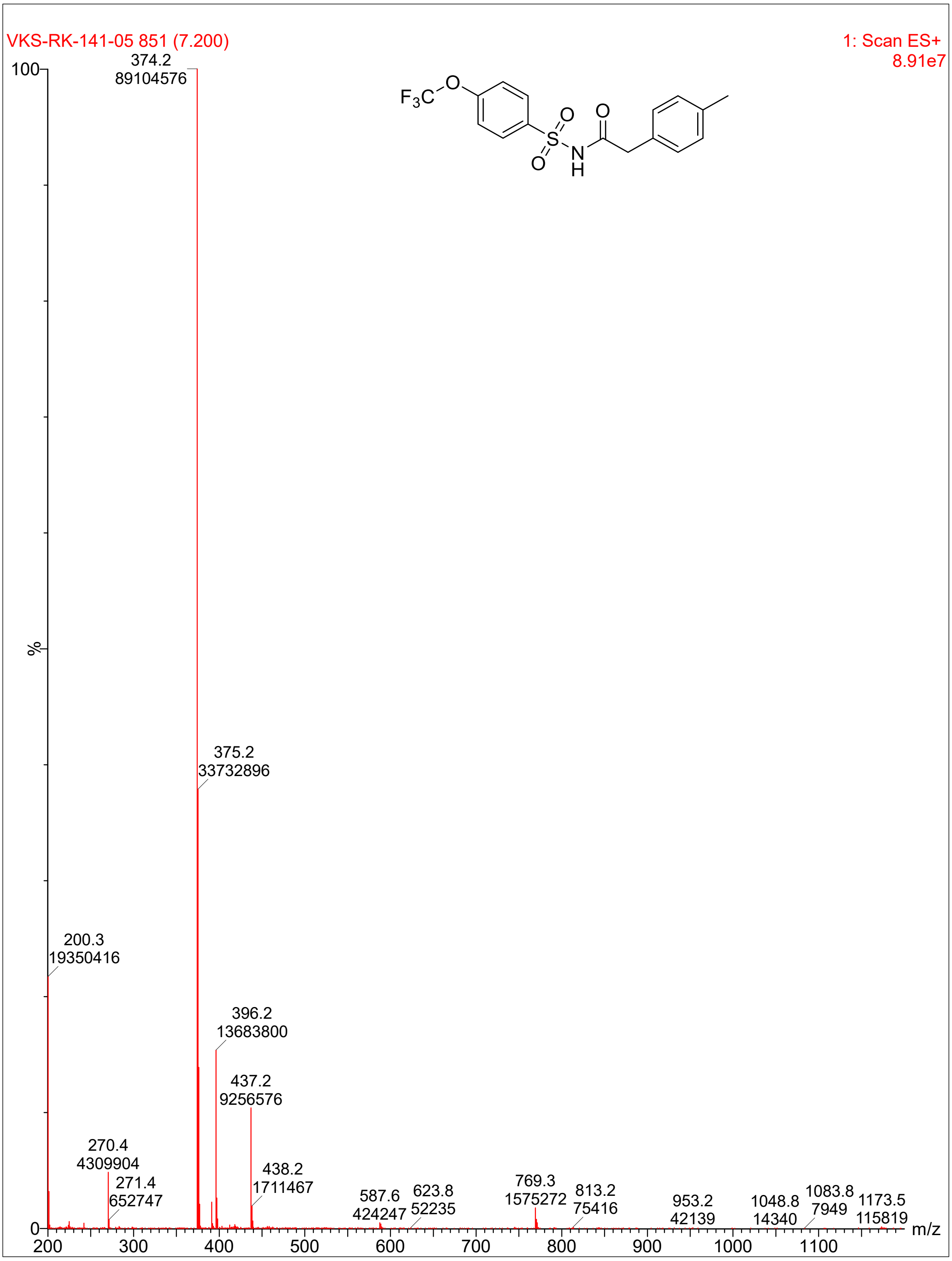
VKS-RK-141-05 851 (7.200)
1: Scan ES+
8.91e7
100
%
0
374.2
89104576
375.2
33732896
200.3
19350416
396.2
13683800
437.2
9256576
270.4
4309904
271.4
652747
438.2
1711467
587.6
424247
623.8
52235
769.3
1575272
813.2
75416
953.2
42139
1048.8
14340
1083.8
7949
1173.5
115819
200
300
400
500
600
700
800
900
1000
1100
m/z
F3C
O
S
O
O
N
H
O

**Compound 29:**

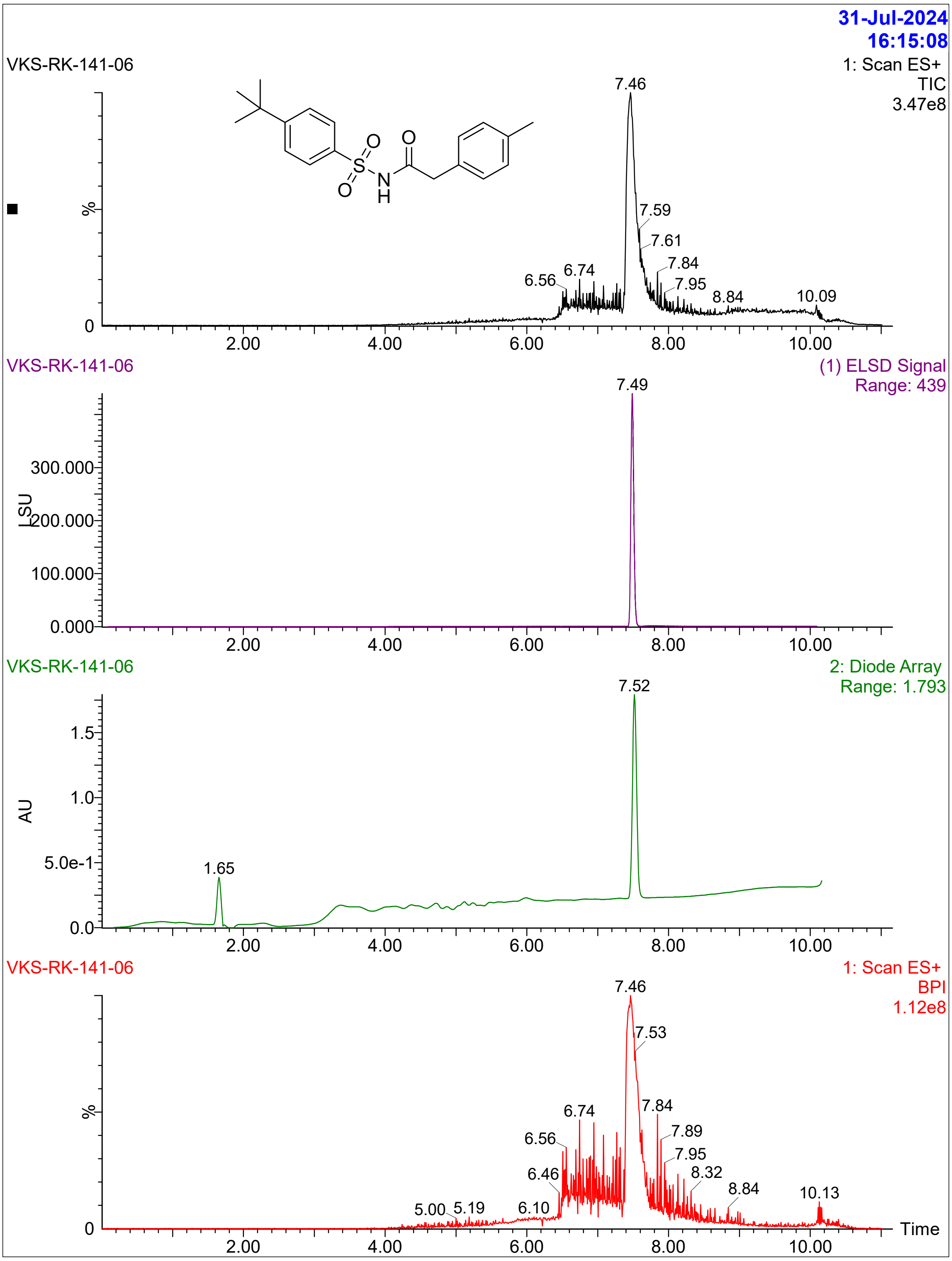

31-Jul-2024
16:15:08
VKS-RK-141-06
1: Scan ES+
TIC
3.47e8
7.46
7.59
7.61
7.84
7.95
6.56
6.74
8.84
10.09
%
0
2.00
4.00
6.00
8.00
10.00
VKS-RK-141-06
(1) ELSD Signal
Range: 439
7.49
300.000
200.000
100.000
0.000
LSU
VKS-RK-141-06
2: Diode Array
Range: 1.793
7.52
1.5
1.0
5.0e-1
0.0
AU
1.65
VKS-RK-141-06
1: Scan ES+
BPI
1.12e8
7.46
7.53
7.84
7.89
7.95
8.32
8.84
10.13
6.74
6.56
6.46
6.10
5.00
5.19
Time

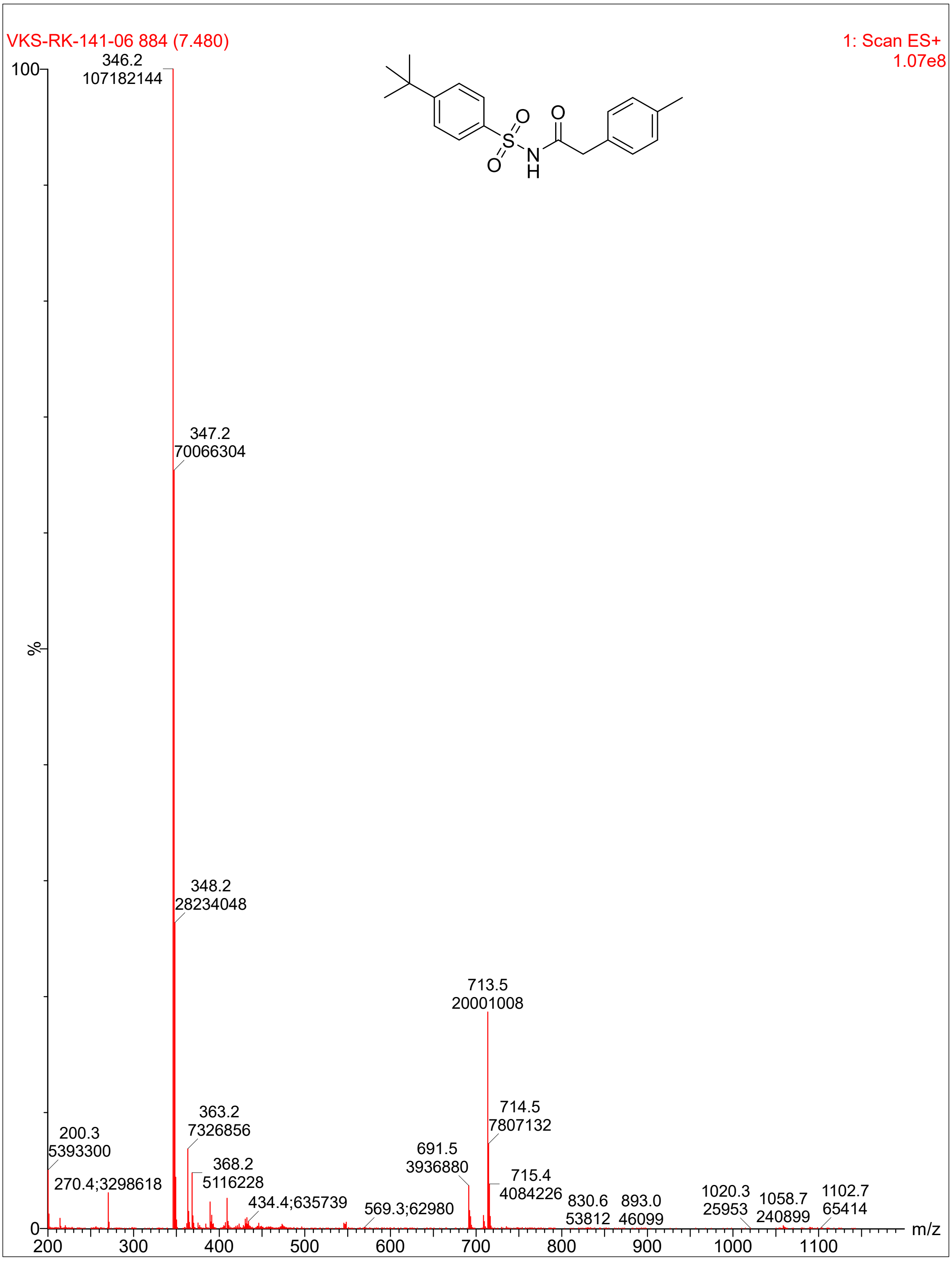

VKS-RK-141-06 884 (7.480)
1: Scan ES+
1.07e8
100
%
0
346.2
107182144
347.2
70066304
348.2
28234048
713.5
20001008
714.5
7807132
715.4
4084226
691.5
3936880
363.2
7326856
368.2
5116228
200.3
5393300
270.4;3298618
434.4;635739
569.3;62980
830.6
53812
893.0
46099
1020.3
25953
1058.7
240899
1102.7
65414
200
300
400
500
600
700
800
900
1000
1100
m/z
O
S
O
N
H
O

**Compound 30**

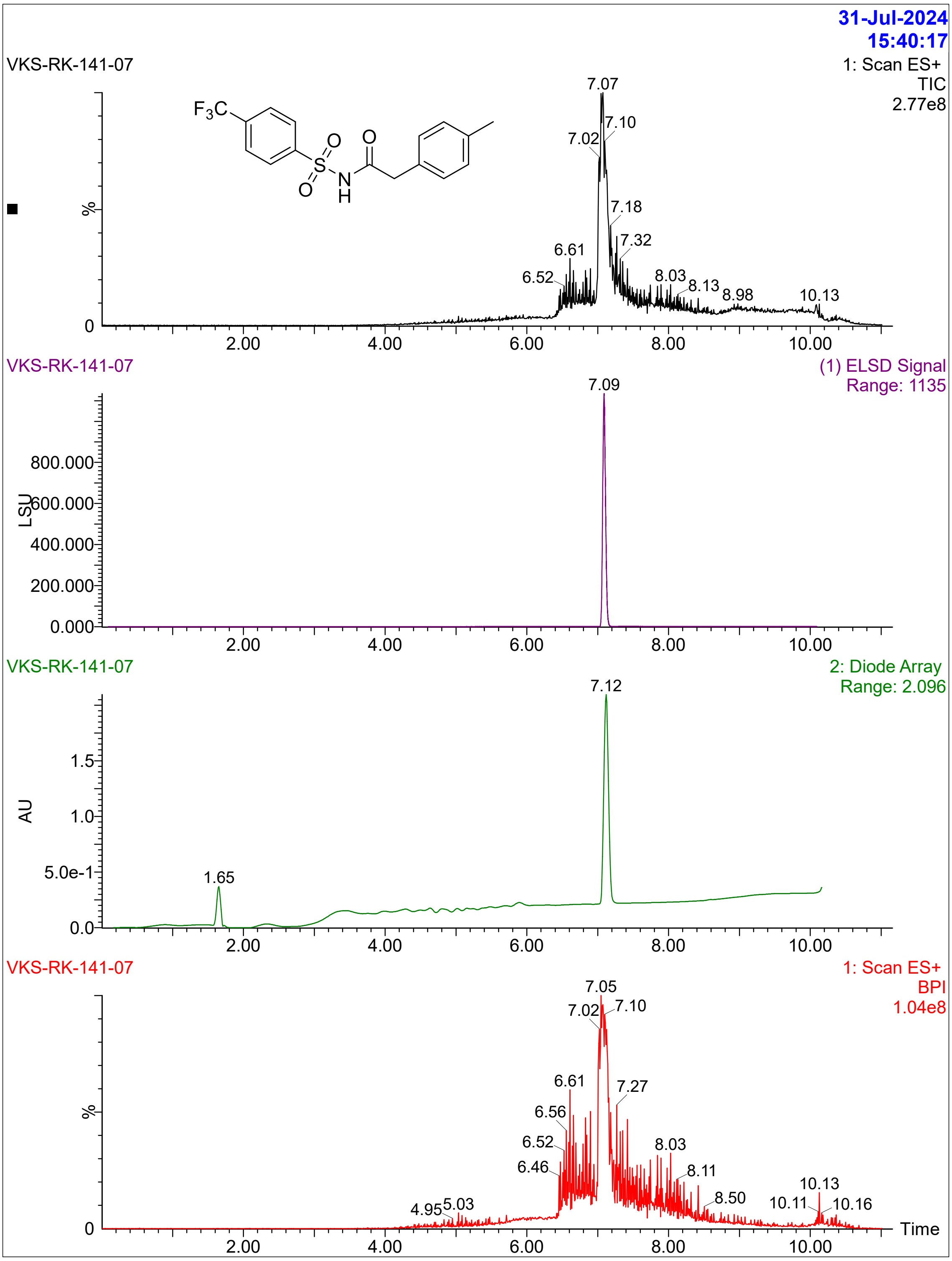

31-Jul-2024
15:40:17
VKS-RK-141-07
1: Scan ES+
TIC
2.77e8
%
7.07
7.10
7.02
7.18
7.32
6.61
6.52
8.03
8.13
8.98
10.13
2.00
4.00
6.00
8.00
10.00
VKS-RK-141-07
(1) ELSD Signal
Range: 1135
7.09
LSU
800.000
600.000
400.000
200.000
0.000
VKS-RK-141-07
2: Diode Array
Range: 2.096
7.12
AU
1.5
1.0
5.0e-1
0.0
1.65
VKS-RK-141-07
1: Scan ES+
BPI
1.04e8
7.05
7.02
7.10
6.61
7.27
6.56
6.52
8.03
6.46
8.11
10.13
4.95
5.03
8.50
10.11
10.16
Time

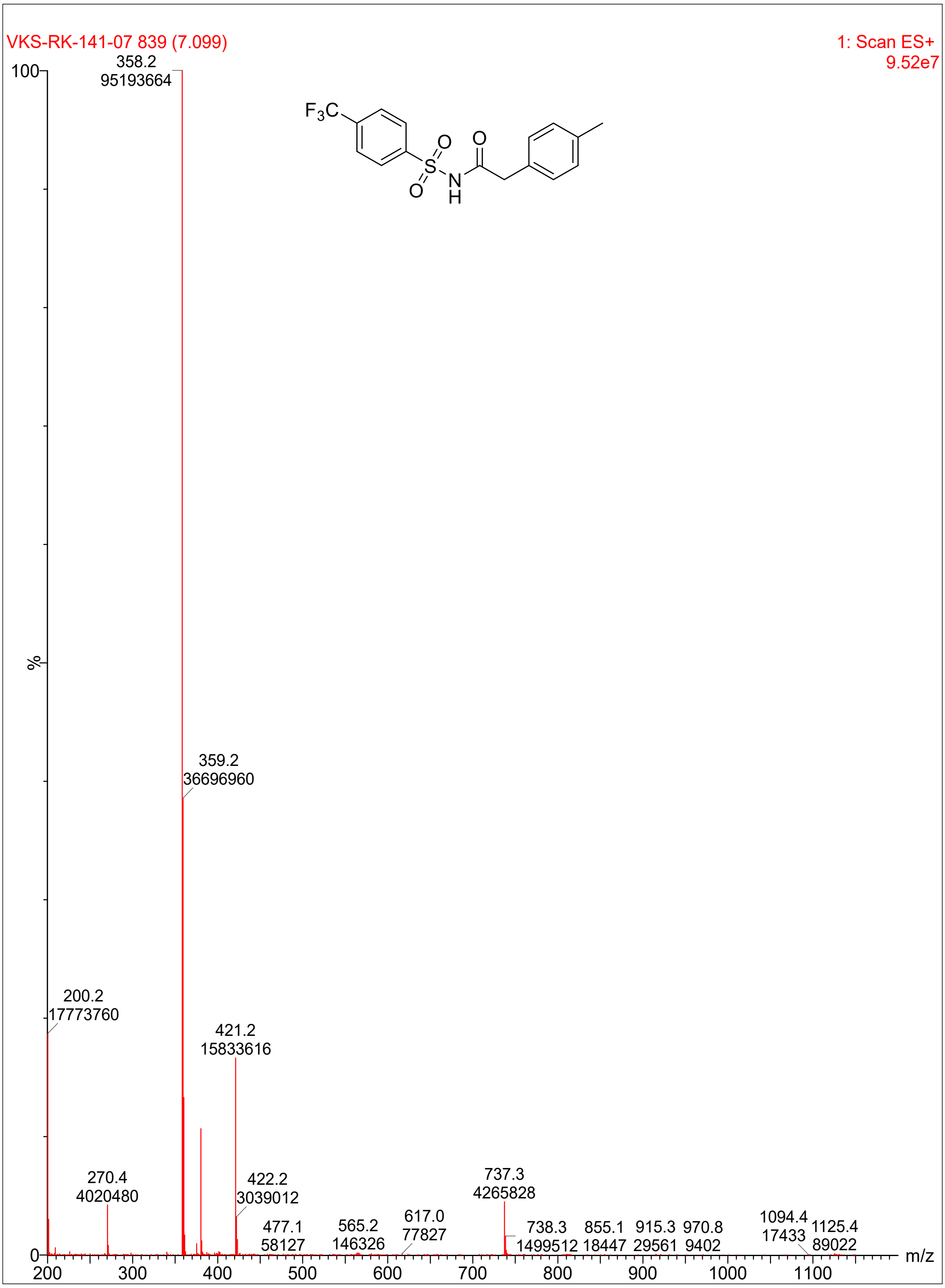

VKS-RK-141-07 839 (7.099)
1: Scan ES+
9.52e7
100
%
0
358.2
95193664
359.2
36696960
200.2
17773760
421.2
15833616
270.4
4020480
422.2
3039012
737.3
4265828
477.1
58127
565.2
146326
617.0
77827
738.3
1499512
855.1
18447
915.3
29561
970.8
9402
1094.4
17433
1125.4
89022
200
300
400
500
600
700
800
900
1000
1100
m/z
F3C
O
S
O
N
H
O

**Compound 32:**

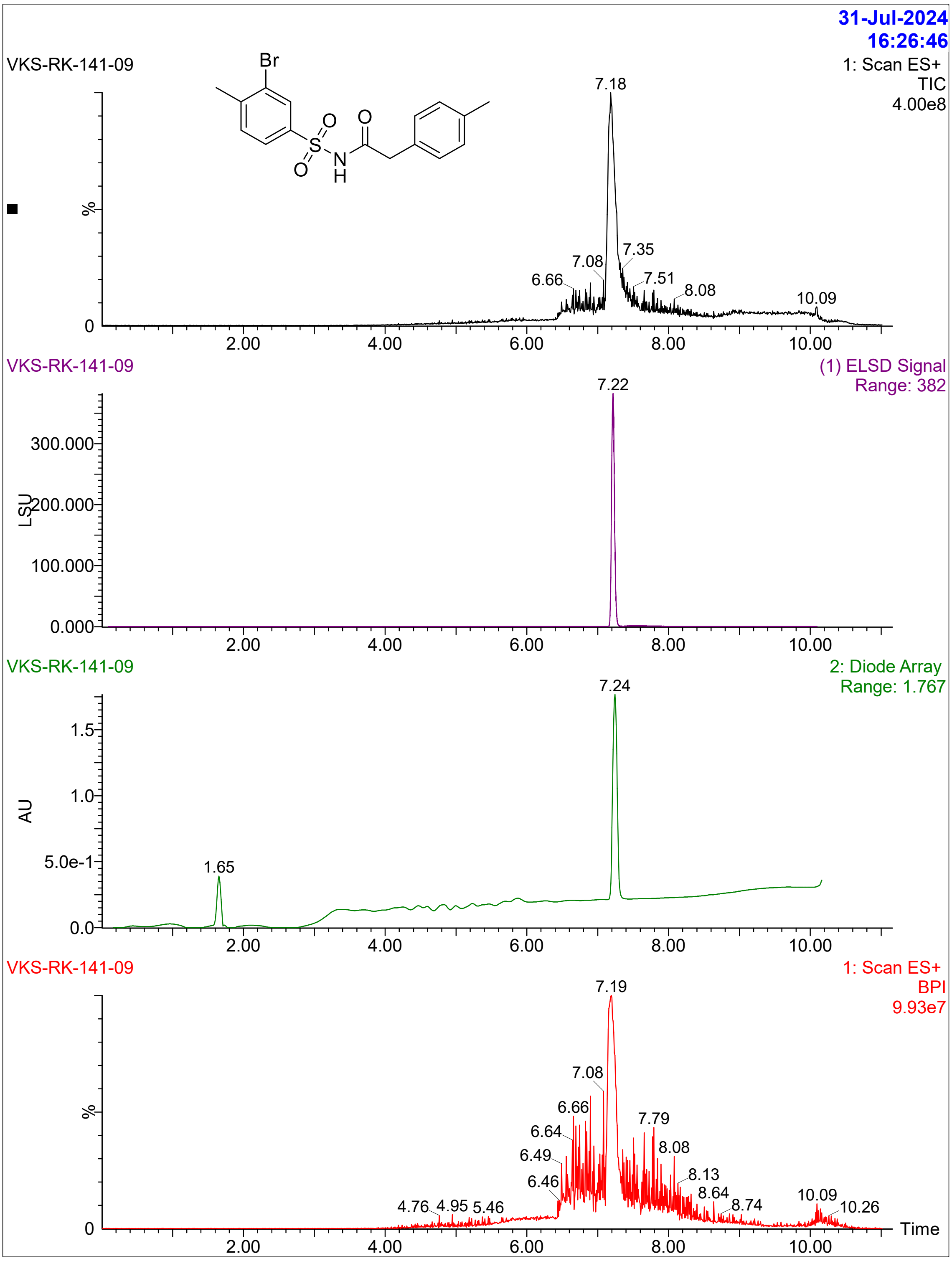

31-Jul-2024
16:26:46
VKS-RK-141-09
Br
O
O
S
O
N
H
1: Scan ES+
TIC
4.00e8
7.18
%
7.08
7.35
6.66
7.51
8.08
10.09
0
2.00
4.00
6.00
8.00
10.00
VKS-RK-141-09
(1) ELSD Signal
Range: 382
7.22
300.000
LSU
200.000
100.000
0.000
VKS-RK-141-09
2: Diode Array
Range: 1.767
7.24
1.5
1.0
AU
5.0e-1
1.65
0.0
VKS-RK-141-09
1: Scan ES+
BPI
9.93e7
7.19
7.08
%
6.66
7.79
6.64
8.08
6.49
8.13
6.46
8.64
4.76
4.95
5.46
8.74
10.09
10.26
Time

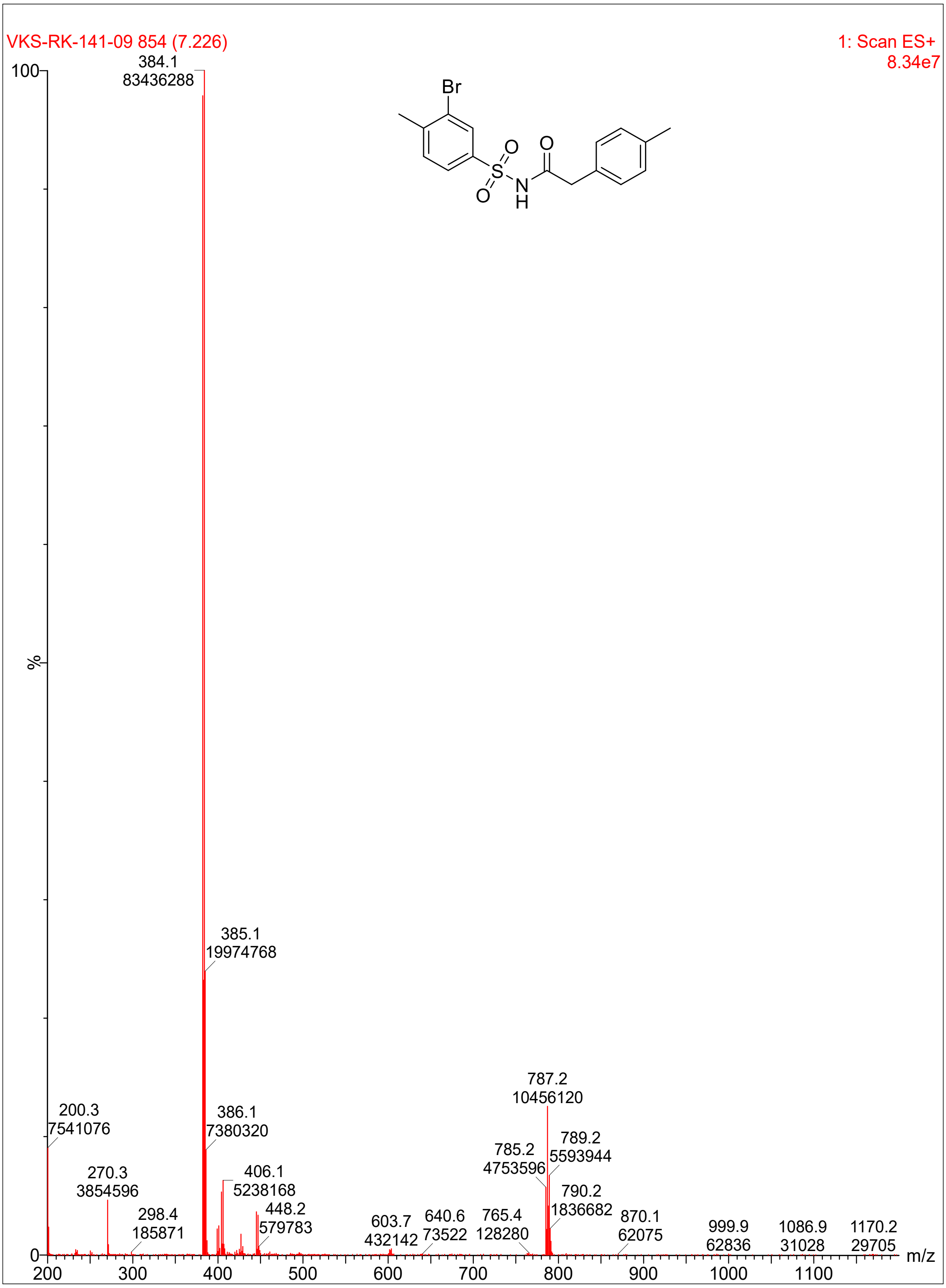

VKS-RK-141-09 854 (7.226)
1: Scan ES+
8.34e7
Br
O
S
O
N
H
O
%
384.1
83436288
385.1
19974768
200.3
7541076
386.1
7380320
270.3
3854596
406.1
5238168
298.4
185871
448.2
579783
603.7
432142
640.6
73522
765.4
128280
785.2
4753596
787.2
10456120
789.2
5593944
790.2
1836682
870.1
62075
999.9
62836
1086.9
31028
1170.2
29705
m/z
100
0
200
300
400
500
600
700
800
900
1000
1100

**Compound 35:**

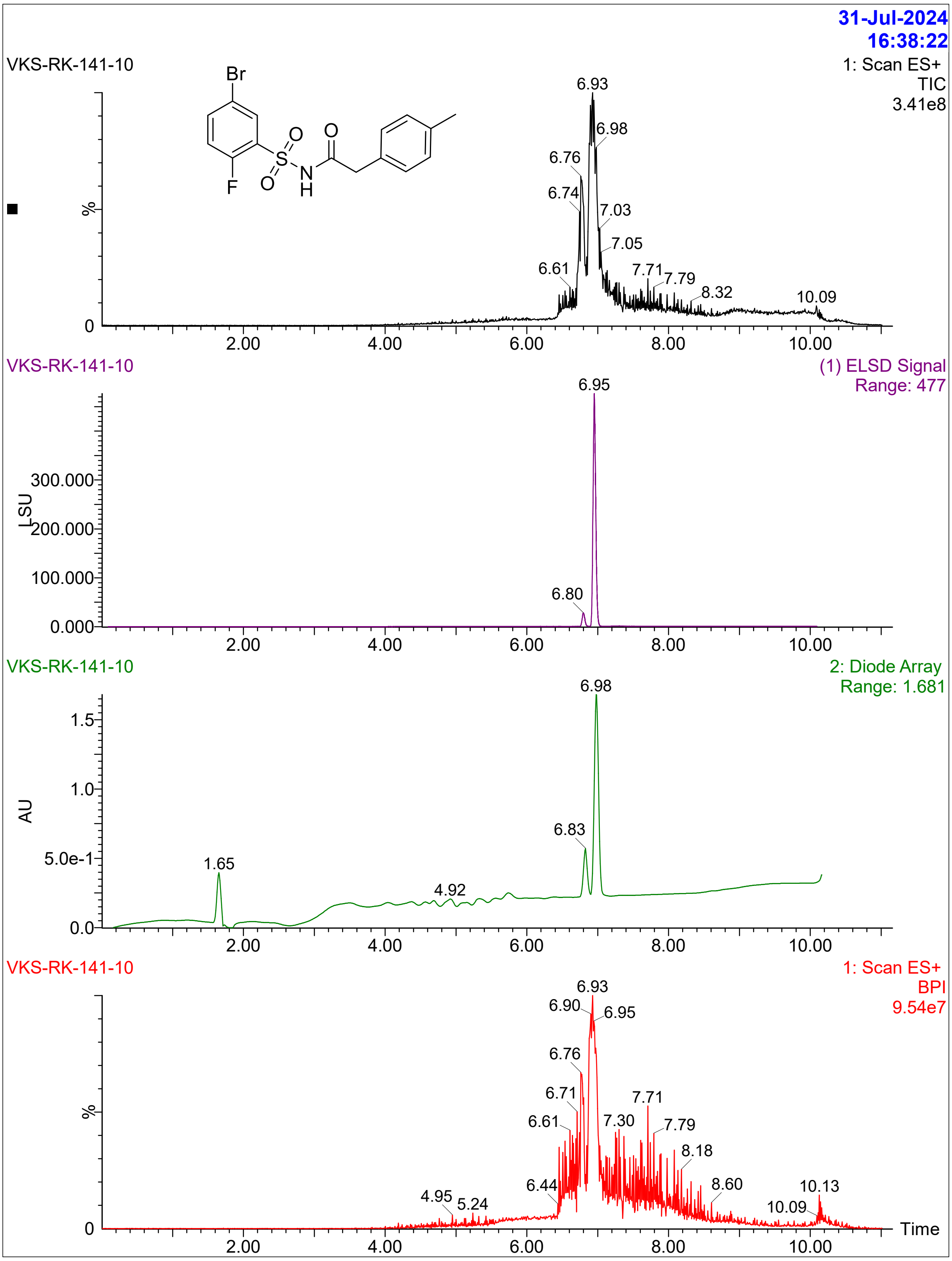

31-Jul-2024
16:38:22
VKS-RK-141-10
1: Scan ES+
TIC
3.41e8
Br
F
O
S
O
N
H
O
%
0
6.93
6.98
6.76
6.74
7.03
7.05
6.61
7.71
7.79
8.32
10.09
2.00
4.00
6.00
8.00
10.00
VKS-RK-141-10
(1) ELSD Signal
Range: 477
6.95
LSU
300.000
200.000
100.000
0.000
6.80
VKS-RK-141-10
2: Diode Array
Range: 1.681
6.98
AU
1.5
1.0
5.0e-1
0.0
6.83
1.65
4.92
VKS-RK-141-10
1: Scan ES+
BPI
9.54e7
6.93
6.90
6.95
6.76
6.71
7.71
6.61
7.30
7.79
8.18
6.44
4.95
5.24
8.60
10.13
10.09
Time

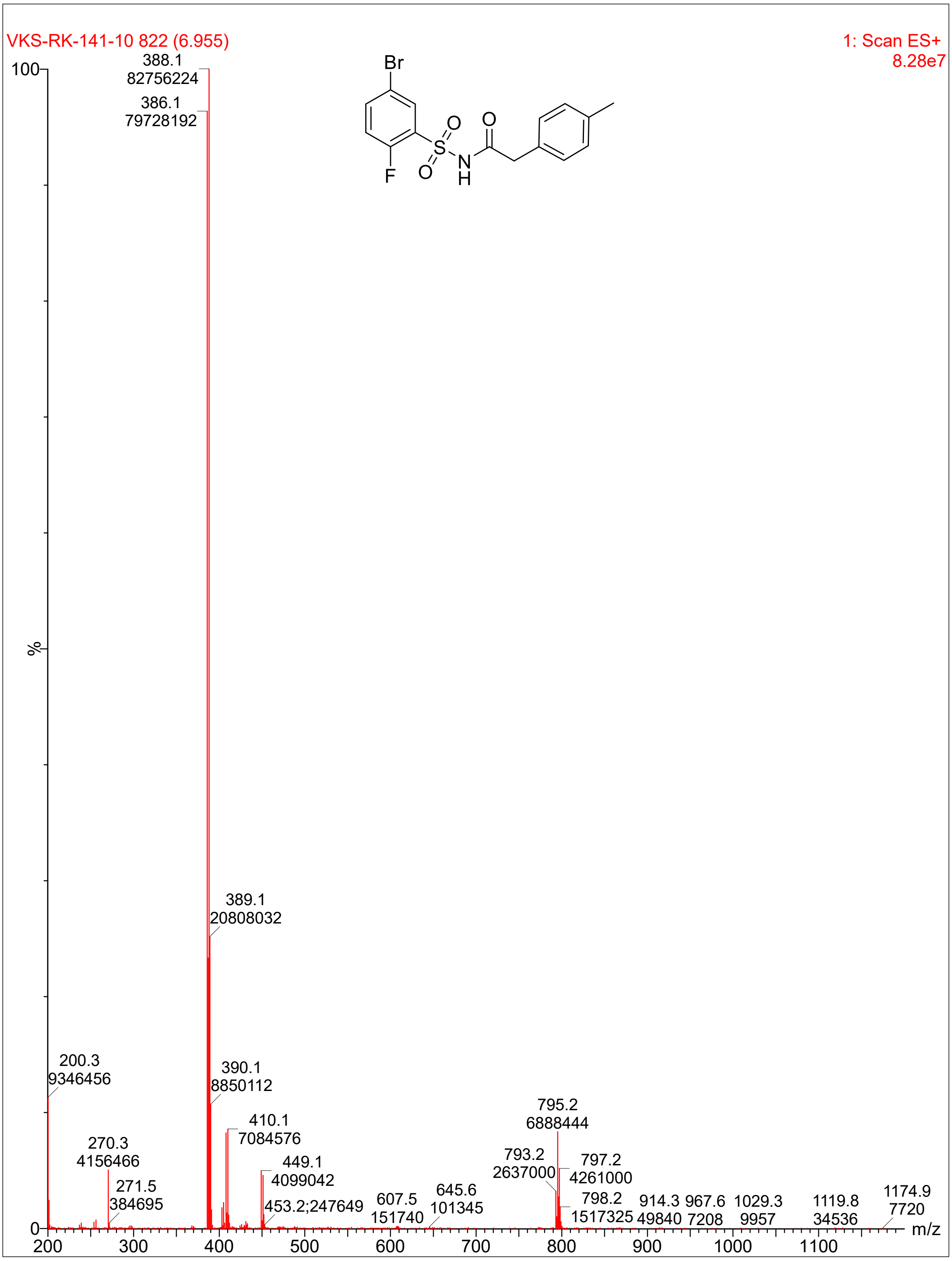
VKS-RK-141-10 822 (6.955)
1: Scan ES+
8.28e7
Br
F
O
S
O
N
H
O
388.1
82756224
386.1
79728192
389.1
20808032
200.3
9346456
390.1
8850112
410.1
7084576
270.3
4156466
449.1
4099042
271.5
384695
453.2;247649
607.5
151740
645.6
101345
795.2
6888444
793.2
2637000
797.2
4261000
798.2
1517325
914.3
49840
967.6
7208
1029.3
9957
1119.8
34536
1174.9
7720
100
%
0
200
300
400
500
600
700
800
900
1000
1100
m/z

**Compound 36**:

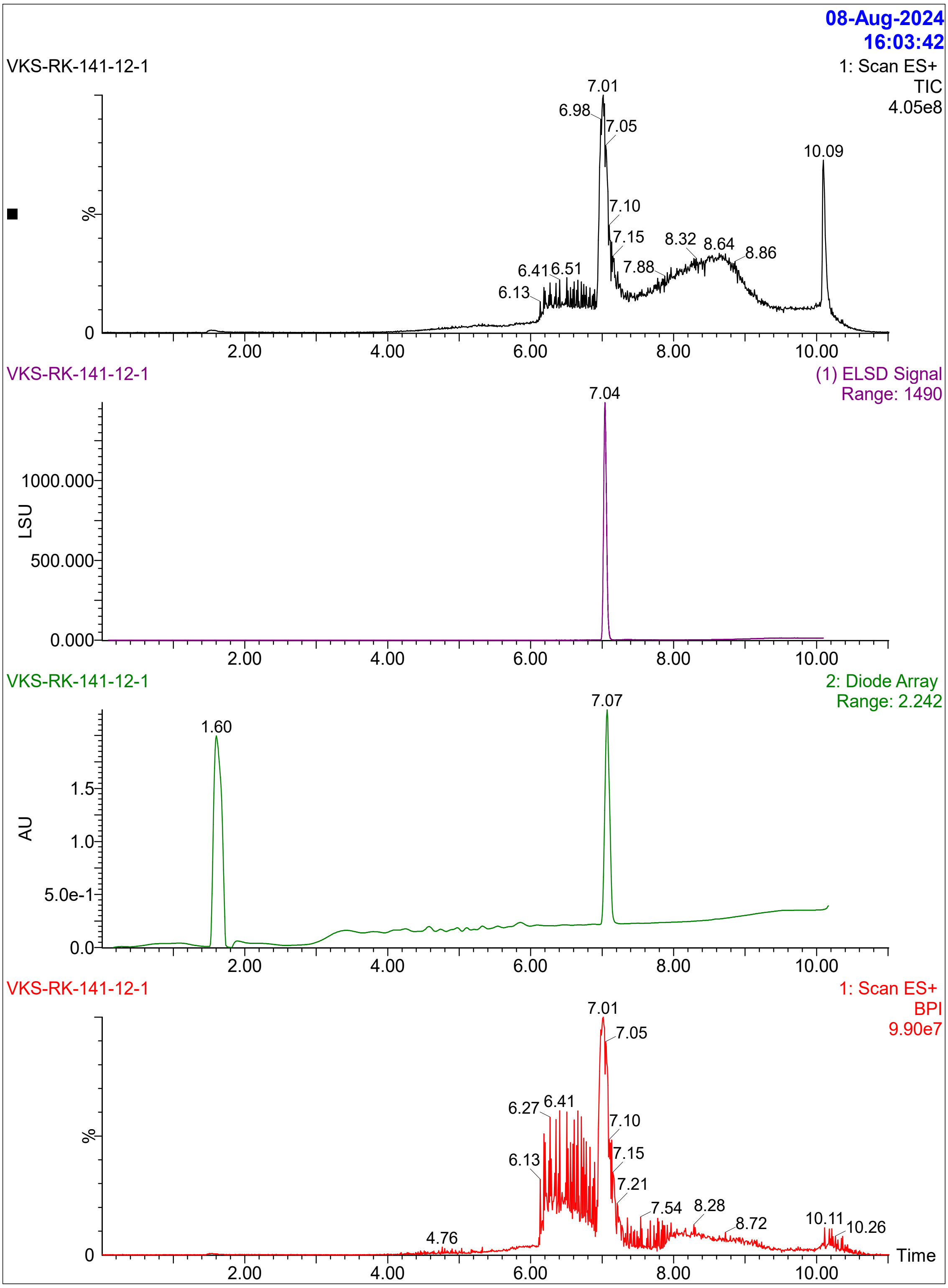

08-Aug-2024
16:03:42
VKS-RK-141-12-1
1: Scan ES+
TIC
4.05e8
%
7.01
6.98
7.05
10.09
7.10
7.15
8.32
8.64
8.86
6.41
6.51
7.88
6.13
0
2.00
4.00
6.00
8.00
10.00
VKS-RK-141-12-1
(1) ELSD Signal
Range: 1490
7.04
1000.000
LSU
500.000
0.000
VKS-RK-141-12-1
2: Diode Array
Range: 2.242
7.07
1.60
1.5
AU
1.0
5.0e-1
0.0
VKS-RK-141-12-1
1: Scan ES+
BPI
9.90e7
7.01
7.05
6.27
6.41
7.10
%
7.15
6.13
7.21
7.54
8.28
4.76
8.72
10.11
10.26
Time

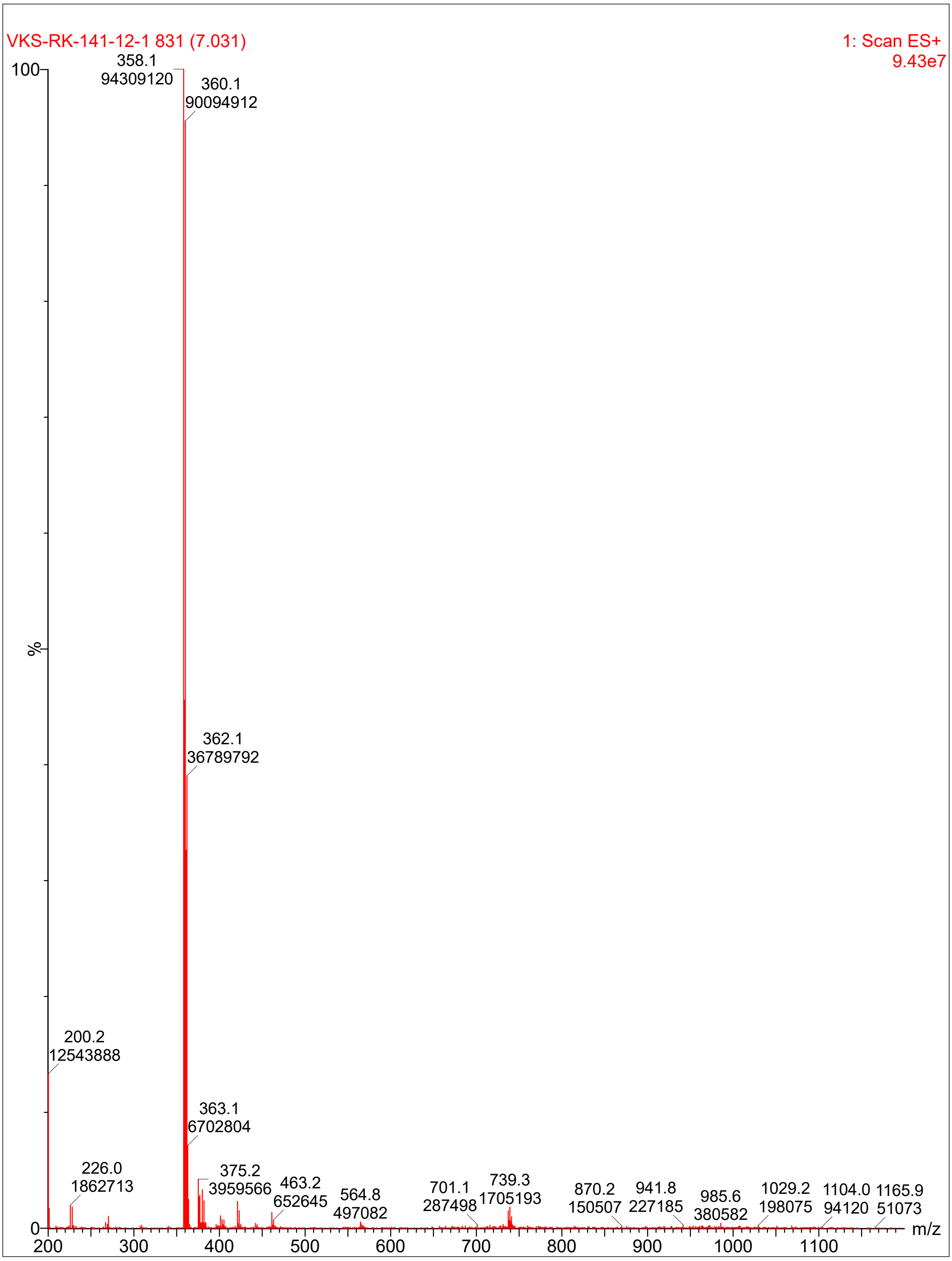
VKS-RK-141-12-1 831 (7.031)
1: Scan ES+
9.43e7
100
%
0
358.1
94309120
360.1
90094912
362.1
36789792
200.2
12543888
363.1
6702804
226.0
1862713
375.2
3959566
463.2
652645
564.8
497082
701.1
287498
739.3
1705193
870.2
150507
941.8
227185
985.6
380582
1029.2
198075
1104.0
94120
1165.9
51073
200
300
400
500
600
700
800
900
1000
1100
m/z

**Compound 38**:

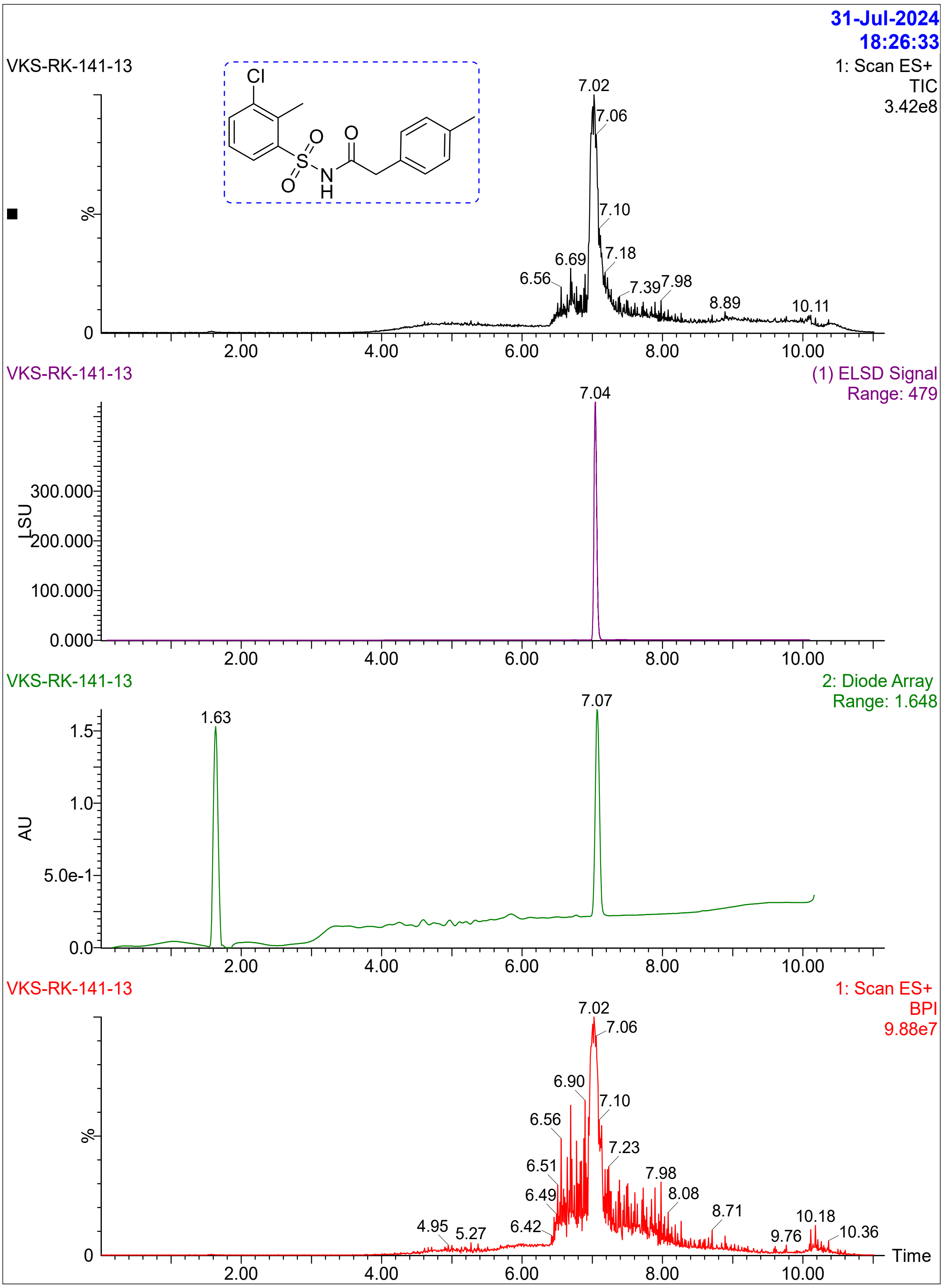
31-Jul-2024
18:26:33
VKS-RK-141-13
1: Scan ES+
TIC
3.42e8
Cl
O
O
S
O
N
H
7.02
7.06
7.10
7.18
6.69
6.56
7.39
7.98
8.89
10.11
%
0
2.00
4.00
6.00
8.00
10.00
VKS-RK-141-13
(1) ELSD Signal
Range: 479
7.04
LSU
300.000
200.000
100.000
0.000
VKS-RK-141-13
2: Diode Array
Range: 1.648
1.63
7.07
AU
1.5
1.0
5.0e-1
0.0
VKS-RK-141-13
1: Scan ES+
BPI
9.88e7
7.02
7.06
6.90
7.10
6.56
7.23
6.51
7.98
6.49
8.08
8.71
4.95
5.27
6.42
10.18
9.76
10.36
Time

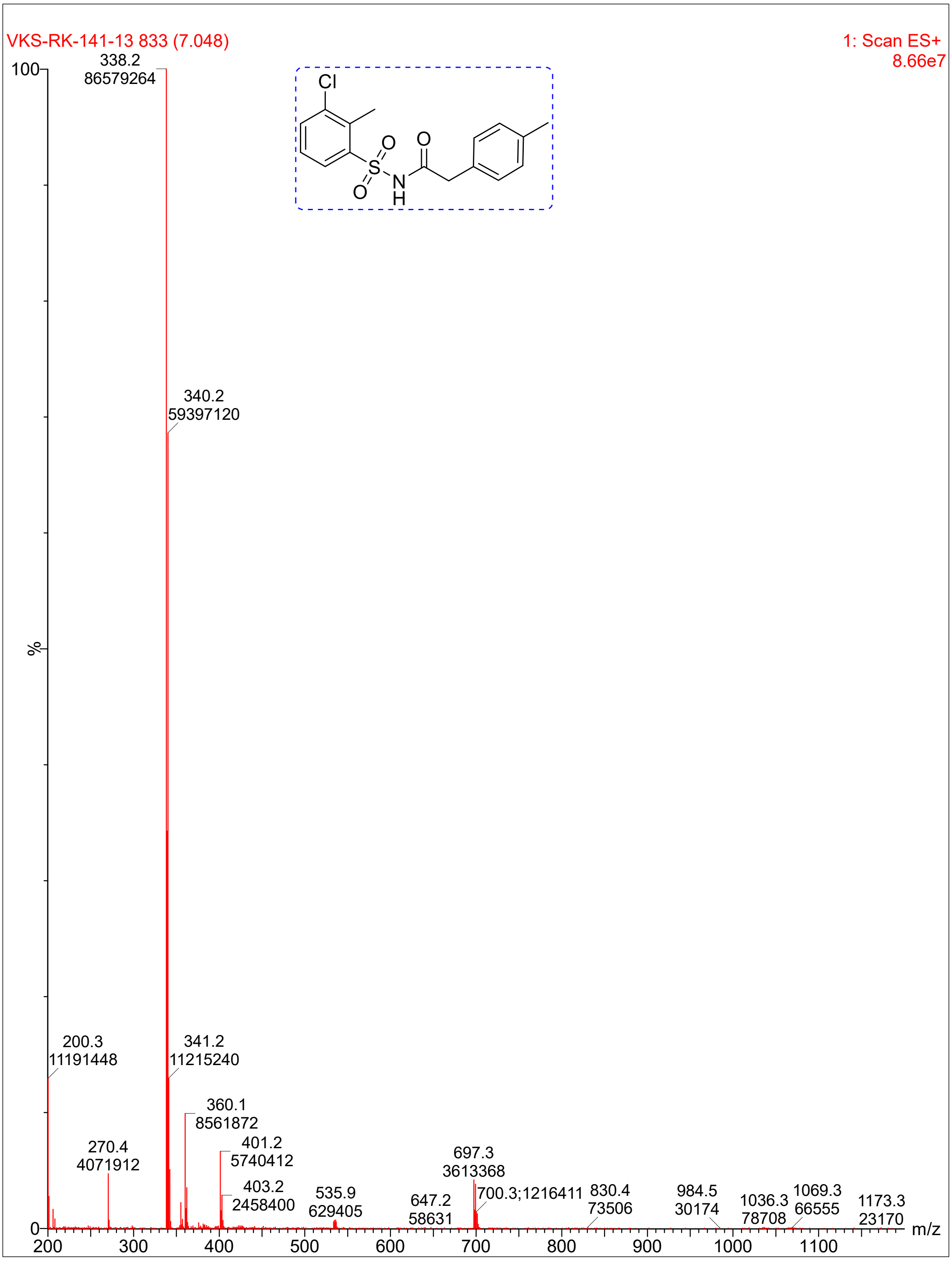

VKS-RK-141-13 833 (7.048)
1: Scan ES+
8.66e7
Cl
O
O
O
N
H
S
338.2
86579264
340.2
59397120
200.3
11191448
341.2
11215240
360.1
8561872
270.4
4071912
401.2
5740412
403.2
2458400
535.9
629405
647.2
58631
697.3
3613368
700.3;1216411
830.4
73506
984.5
30174
1036.3
78708
1069.3
66555
1173.3
23170
%
m/z

**Compound 47:**

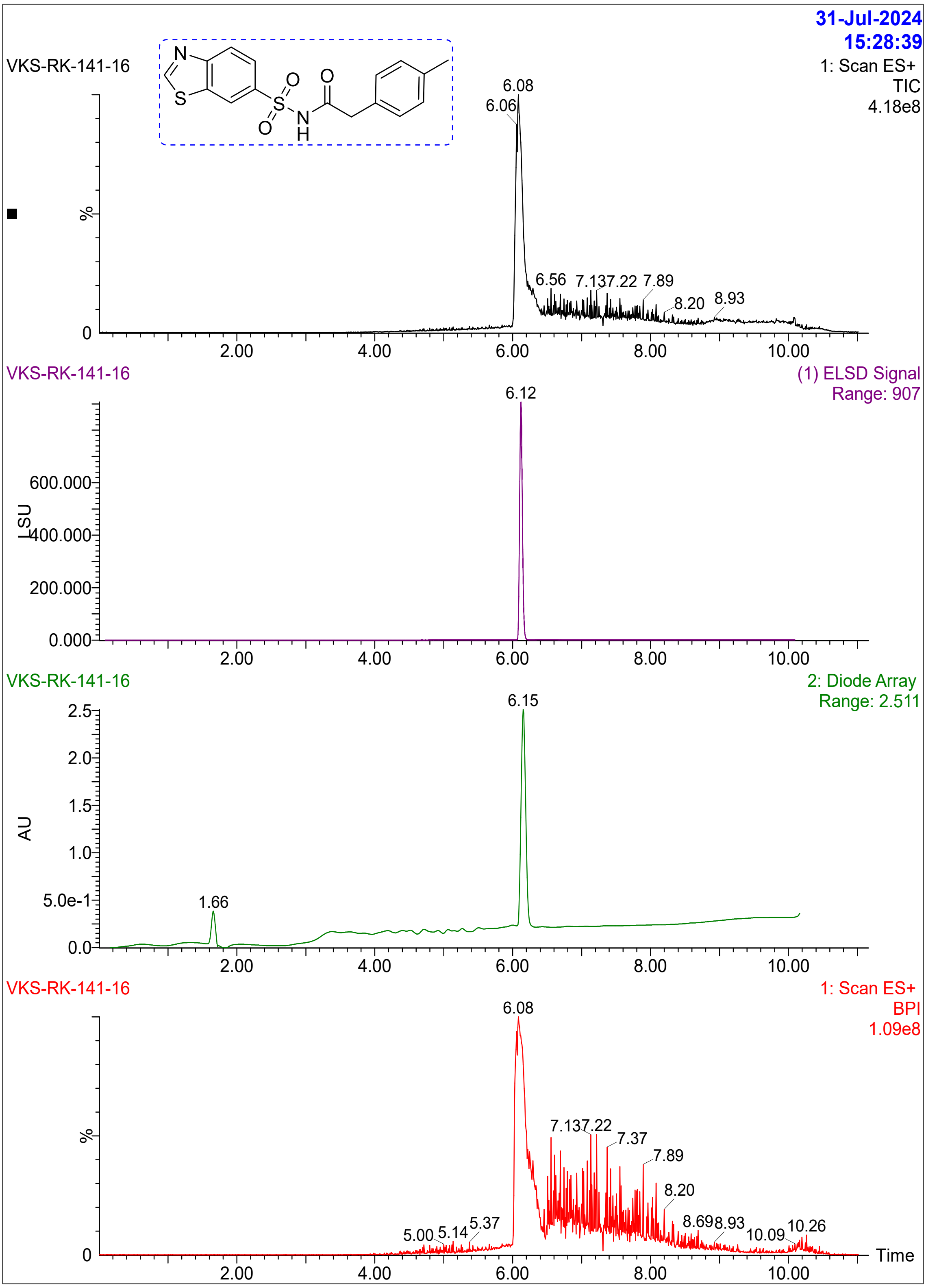

31-Jul-2024
15:28:39
VKS-RK-141-16
1: Scan ES+
TIC
4.18e8
6.08
6.06
%
6.56 7.137.22 7.89
8.20 8.93
0
2.00 4.00 6.00 8.00 10.00
VKS-RK-141-16
(1) ELSD Signal
Range: 907
6.12
600.000
LSU
400.000
200.000
0.000
2.00 4.00 6.00 8.00 10.00
VKS-RK-141-16
2: Diode Array
Range: 2.511
6.15
2.5
2.0
1.5
AU
1.0
5.0e-1
1.66
0.0
2.00 4.00 6.00 8.00 10.00
VKS-RK-141-16
1: Scan ES+
BPI
1.09e8
6.08
%
7.137.22
7.37
7.89
8.20
5.00 5.14 5.37
8.698.93
10.09 10.26
0
Time
2.00 4.00 6.00 8.00 10.00

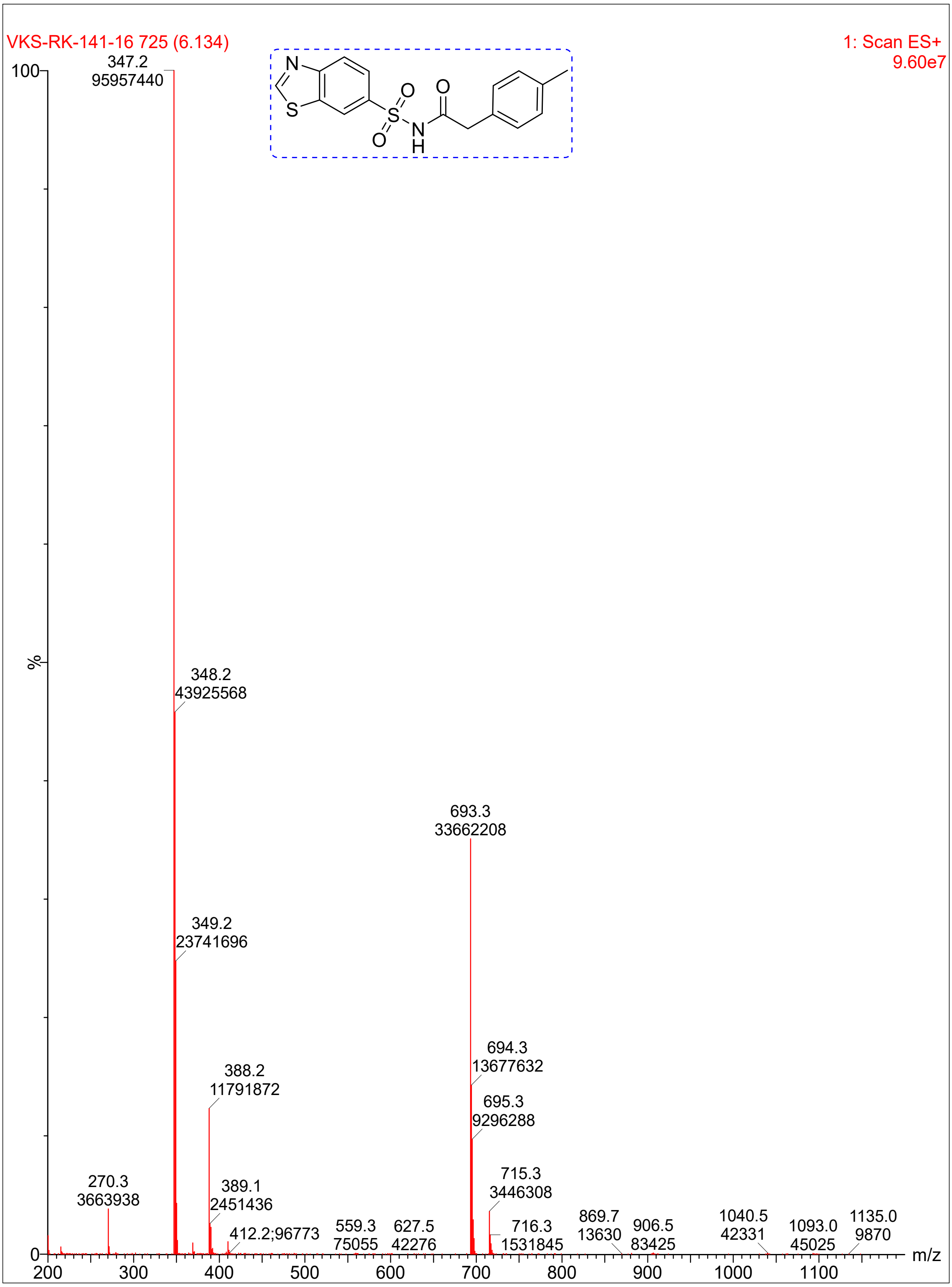

VKS-RK-141-16 725 (6.134)
1: Scan ES+
9.60e7
347.2
95957440
348.2
43925568
349.2
23741696
388.2
11791872
389.1
2451436
270.3
3663938
412.2;96773
559.3
75055
627.5
42276
693.3
33662208
694.3
13677632
695.3
9296288
715.3
3446308
716.3
1531845
869.7
13630
906.5
83425
1040.5
42331
1093.0
45025
1135.0
9870
%
m/z

### Compound 48:

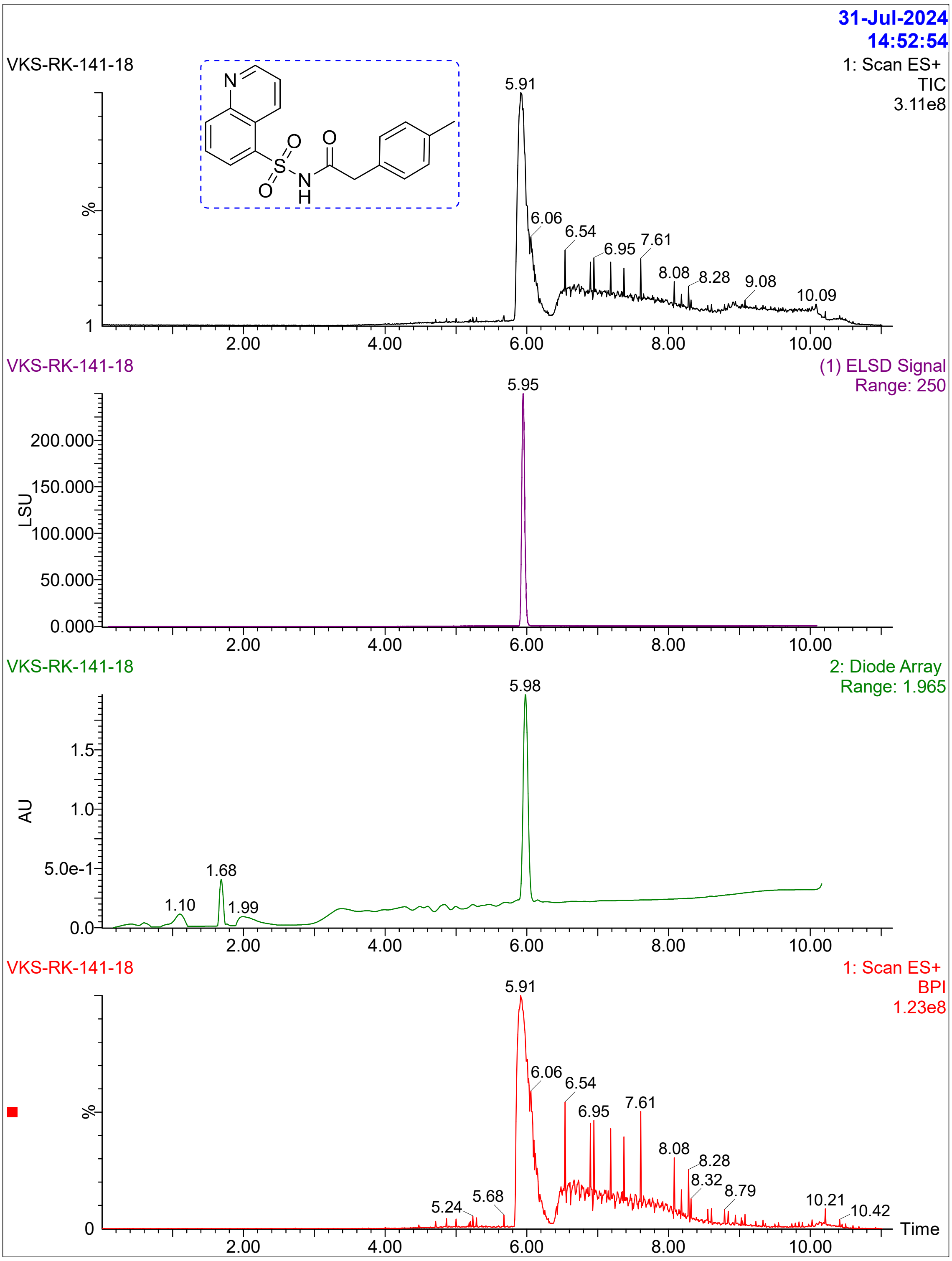
31-Jul-2024
14:52:54
VKS-RK-141-18
1: Scan ES+
TIC
3.11e8
5.91
6.06
6.54
6.95
7.61
8.08
8.28
9.08
10.09
VKS-RK-141-18
(1) ELSD Signal
Range: 250
5.95
LSU
200.000
150.000
100.000
50.000
0.000
VKS-RK-141-18
2: Diode Array
Range: 1.965
5.98
AU
1.5
1.0
5.0e-1
0.0
1.10
1.68
1.99
VKS-RK-141-18
1: Scan ES+
BPI
1.23e8
5.91
6.06
6.54
6.95
7.61
8.08
8.28
8.32
8.79
5.24
5.68
10.21
10.42
2.00
4.00
6.00
8.00
10.00
Time

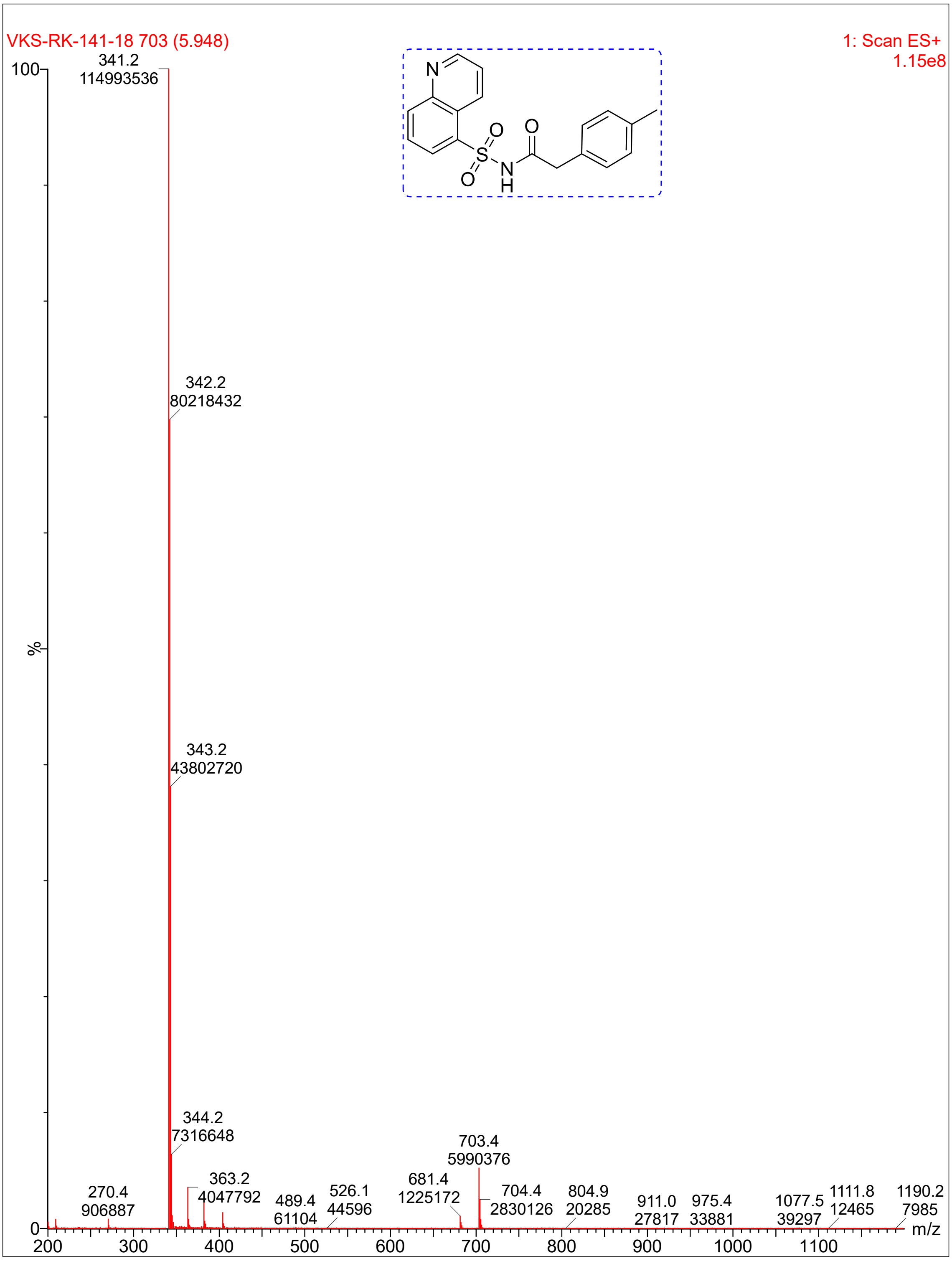
VKS-RK-141-18 703 (5.948)
1: Scan ES+
1.15e8
341.2
114993536
342.2
80218432
343.2
43802720
344.2
7316648
363.2
4047792
270.4
906887
489.4
61104
526.1
44596
681.4
1225172
703.4
5990376
704.4
2830126
804.9
20285
911.0
27817
975.4
33881
1077.5
39297
1111.8
12465
1190.2
7985
%
m/z
N
O
S
N
H

**Compound 37**:

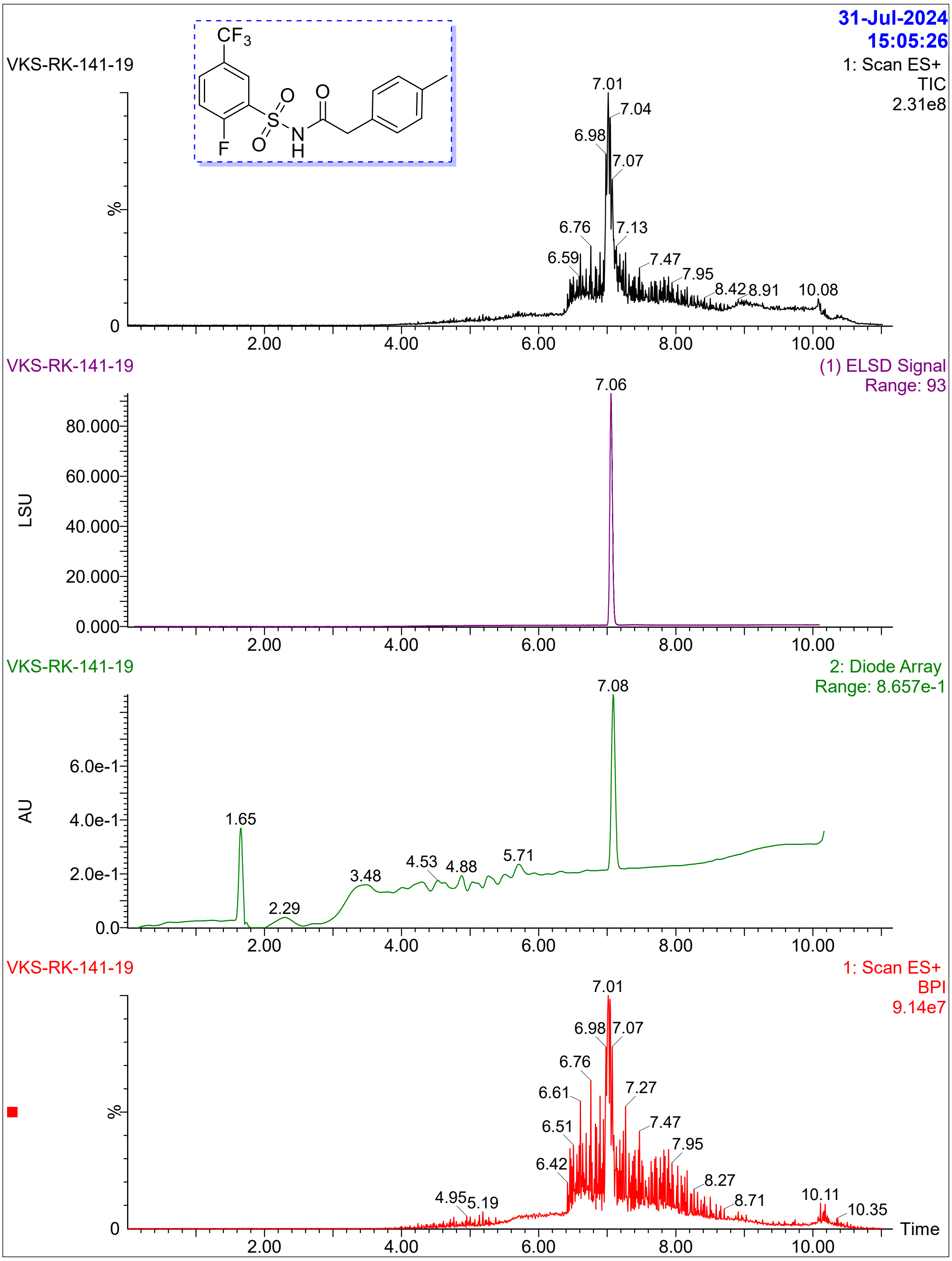
VKS-RK-141-19
31-Jul-2024
15:05:26
1: Scan ES+
TIC
2.31e8
CF3
O
S
N
H
F
%
7.01
7.04
6.98
7.07
6.76
7.13
6.59
7.47
7.95
8.42
8.91
10.08
0
2.00
4.00
6.00
8.00
10.00
VKS-RK-141-19
(1) ELSD Signal
Range: 93
7.06
LSU
80.000
60.000
40.000
20.000
0.000
VKS-RK-141-19
2: Diode Array
Range: 8.657e-1
7.08
AU
6.0e-1
4.0e-1
2.0e-1
0.0
1.65
2.29
3.48
4.53
4.88
5.71
VKS-RK-141-19
1: Scan ES+
BPI
9.14e7
7.01
6.98
7.07
6.76
6.61
7.27
6.51
7.47
7.95
6.42
8.27
8.71
4.95
5.19
10.11
10.35
Time

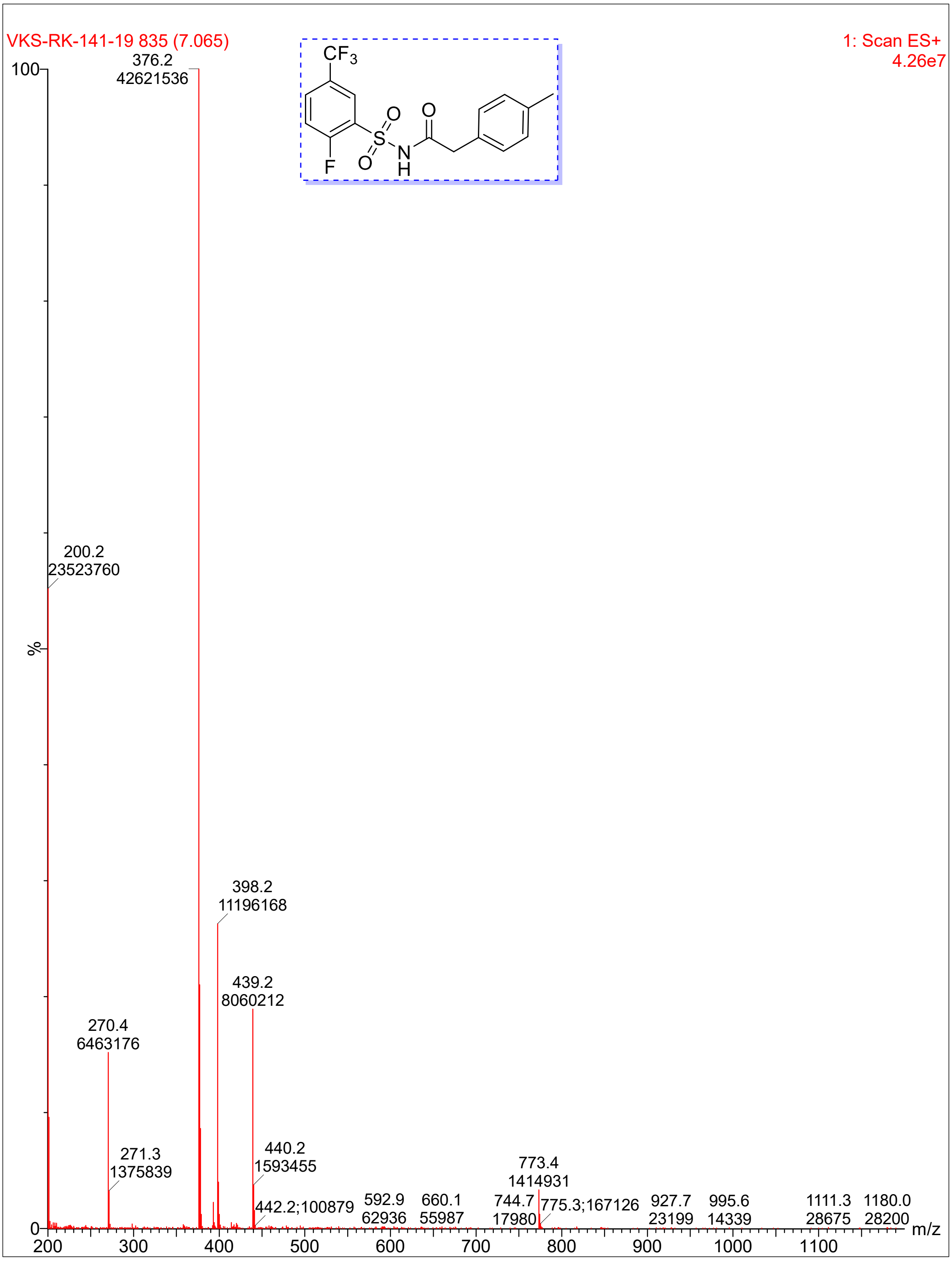

VKS-RK-141-19 835 (7.065)
1: Scan ES+
4.26e7
376.2
42621536
200.2
23523760
398.2
11196168
439.2
8060212
270.4
6463176
271.3
1375839
440.2
1593455
773.4
1414931
442.2;100879
592.9
62936
660.1
55987
744.7
17980
775.3;167126
927.7
23199
995.6
14339
1111.3
28675
1180.0
28200
%
m/z

**Compound 33**:

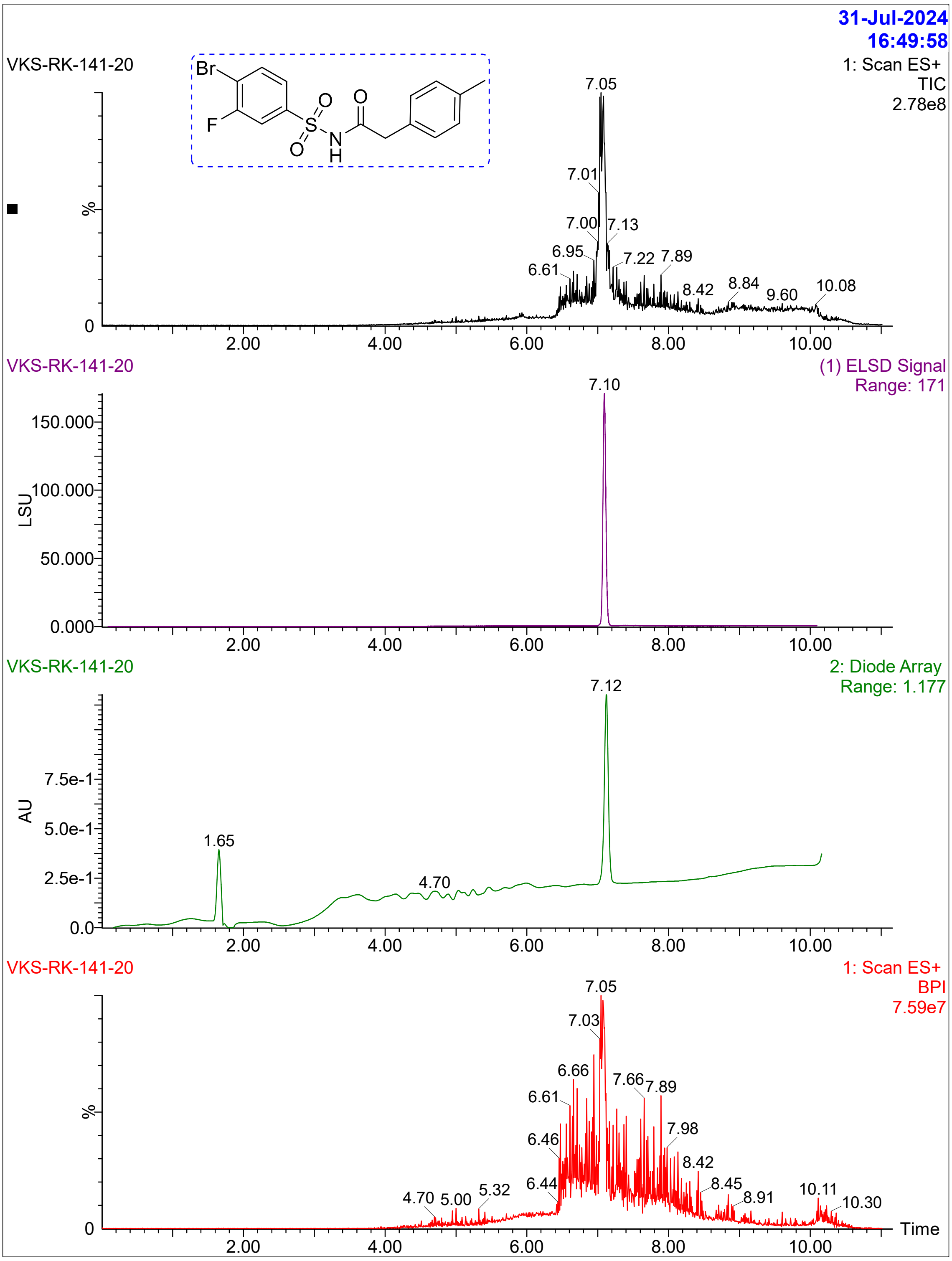

31-Jul-2024
16:49:58
VKS-RK-141-20
1: Scan ES+
TIC
2.78e8
7.05
7.01
7.00
7.13
6.95
6.61
7.22
7.89
8.42
8.84
9.60
10.08
%
0
2.00
4.00
6.00
8.00
10.00
VKS-RK-141-20
(1) ELSD Signal
Range: 171
7.10
150.000
100.000
50.000
0.000
LSU
VKS-RK-141-20
2: Diode Array
Range: 1.177
7.12
7.5e-1
5.0e-1
2.5e-1
0.0
AU
1.65
4.70
VKS-RK-141-20
1: Scan ES+
BPI
7.59e7
7.05
7.03
6.66
6.61
7.66
7.89
6.46
7.98
8.42
6.44
8.45
8.91
10.11
10.30
4.70
5.00
5.32
Time

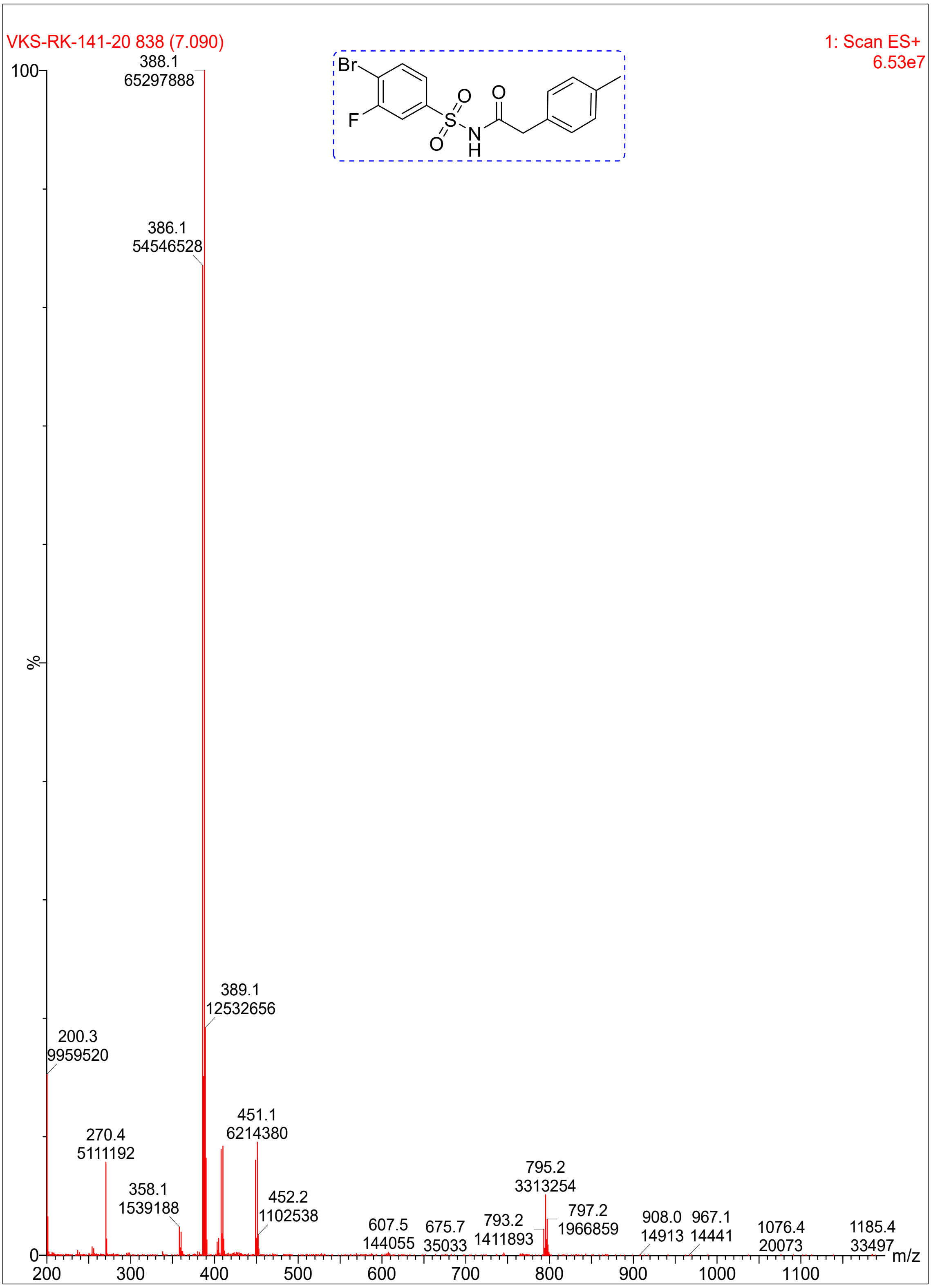
VKS-RK-141-20 838 (7.090)
1: Scan ES+
6.53e7
100
%
0
388.1
65297888
386.1
54546528
389.1
12532656
200.3
9959520
270.4
5111192
358.1
1539188
451.1
6214380
452.2
1102538
607.5
144055
675.7
35033
793.2
1411893
795.2
3313254
797.2
1966859
908.0
14913
967.1
14441
1076.4
20073
1185.4
33497
200
300
400
500
600
700
800
900
1000
1100
m/z
Br
F
O
O
S
N
H
O

**Compound 4**:

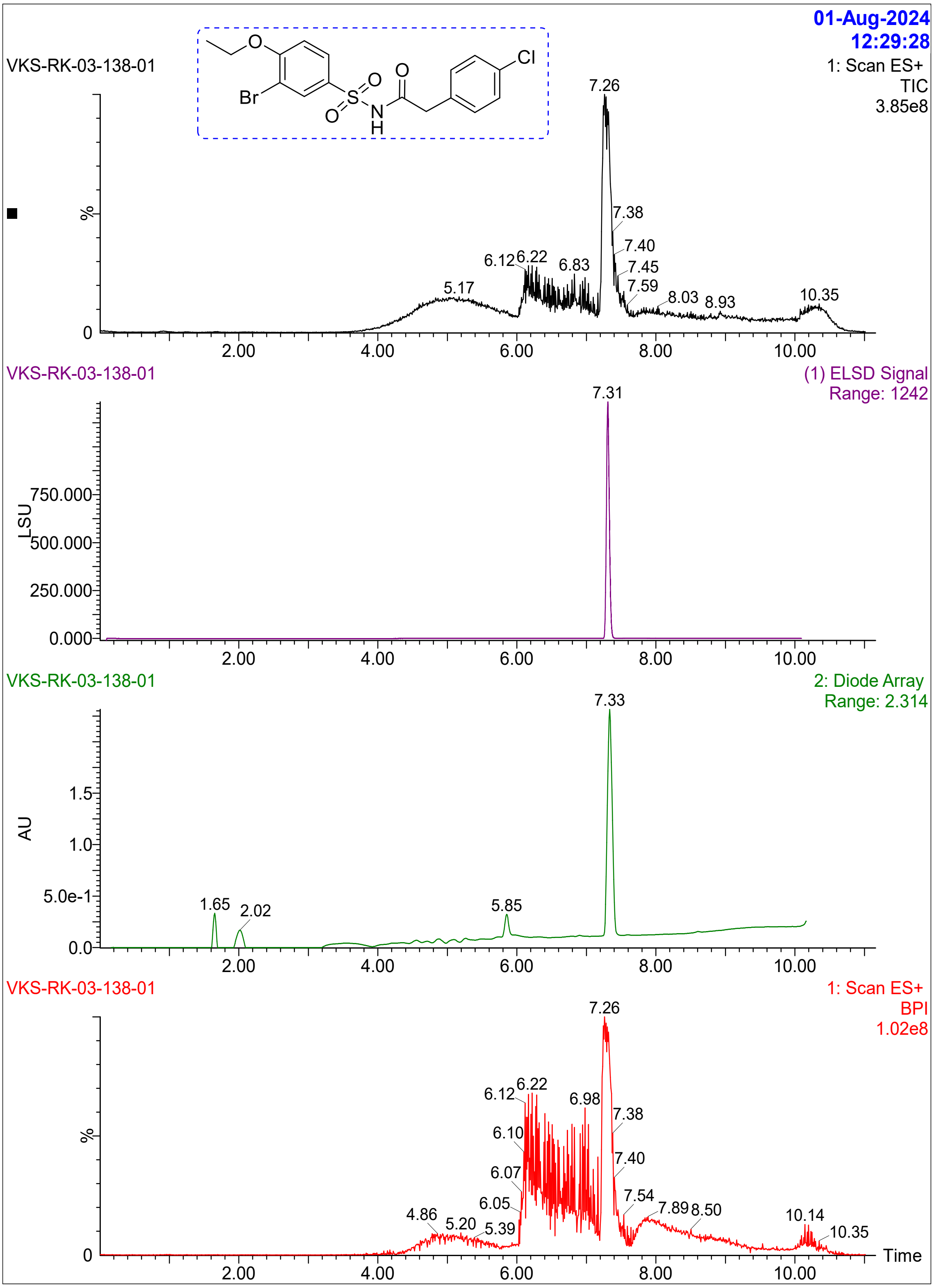

VKS-RK-03-138-01
01-Aug-2024
12:29:28
1: Scan ES+
TIC
3.85e8
Br
O
S
N
H
Cl
7.26
7.38
7.40
7.45
7.59
6.12
6.22
6.83
5.17
8.03
8.93
10.35
%
0
2.00
4.00
6.00
8.00
10.00
VKS-RK-03-138-01
(1) ELSD Signal
Range: 1242
7.31
LSU
750.000
500.000
250.000
0.000
VKS-RK-03-138-01
2: Diode Array
Range: 2.314
7.33
AU
1.5
1.0
5.0e-1
0.0
1.65
2.02
5.85
VKS-RK-03-138-01
1: Scan ES+
BPI
1.02e8
7.26
6.12
6.22
6.98
7.38
6.10
7.40
6.07
7.54
6.05
7.89
8.50
4.86
5.20
5.39
10.14
10.35
Time

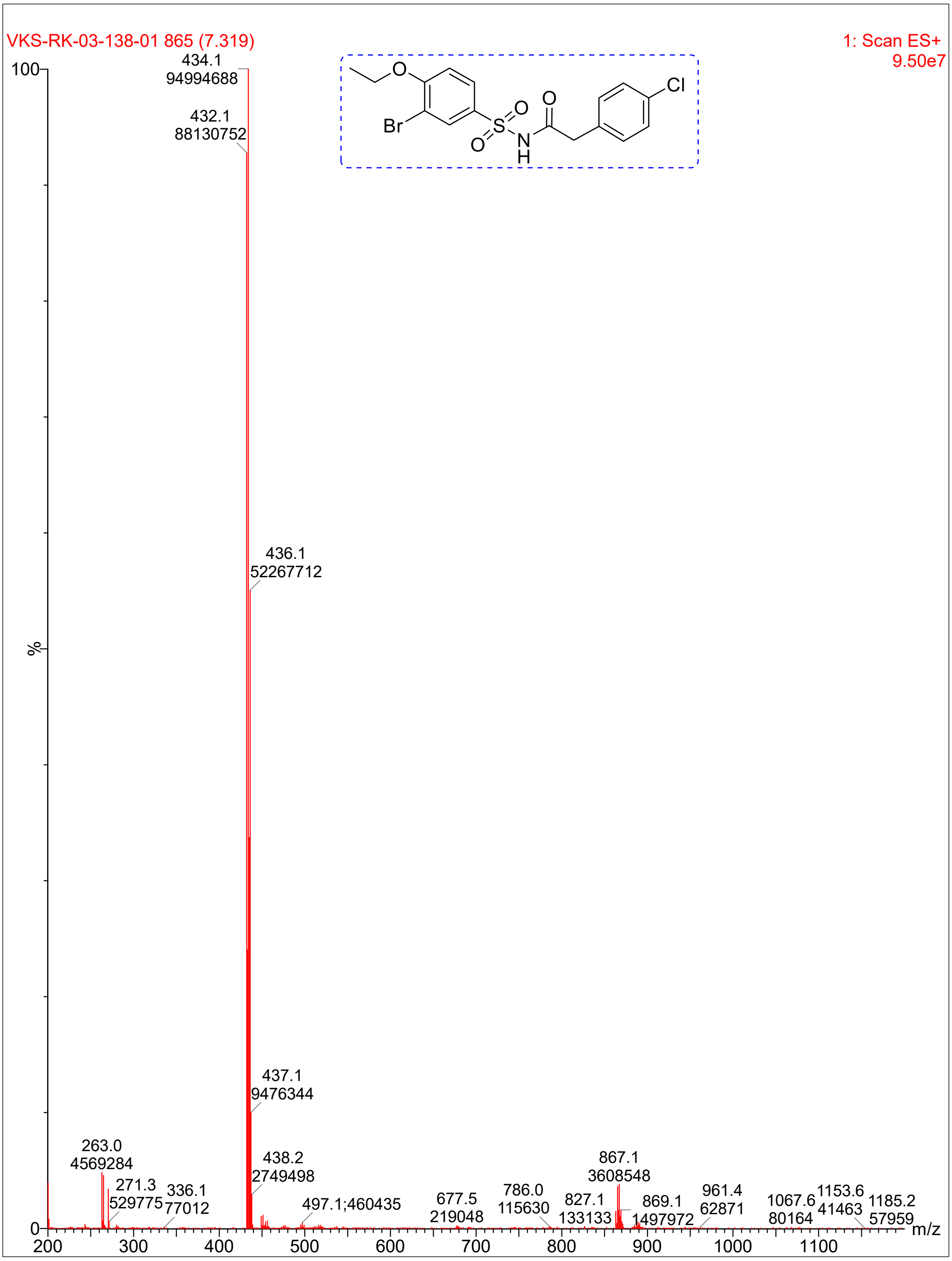
VKS-RK-03-138-01 865 (7.319)
1: Scan ES+
9.50e7
100
%
0
434.1
94994688
432.1
88130752
436.1
52267712
437.1
9476344
263.0
4569284
438.2
2749498
271.3
529775
336.1
77012
497.1;460435
677.5
219048
786.0
115630
827.1
133133
867.1
3608548
869.1
1497972
961.4
62871
1067.6
80164
1153.6
41463
1185.2
57959
m/z
200
300
400
500
600
700
800
900
1000
1100
O
Br
S
O
O
N
H
O
Cl

**Compound 23:**

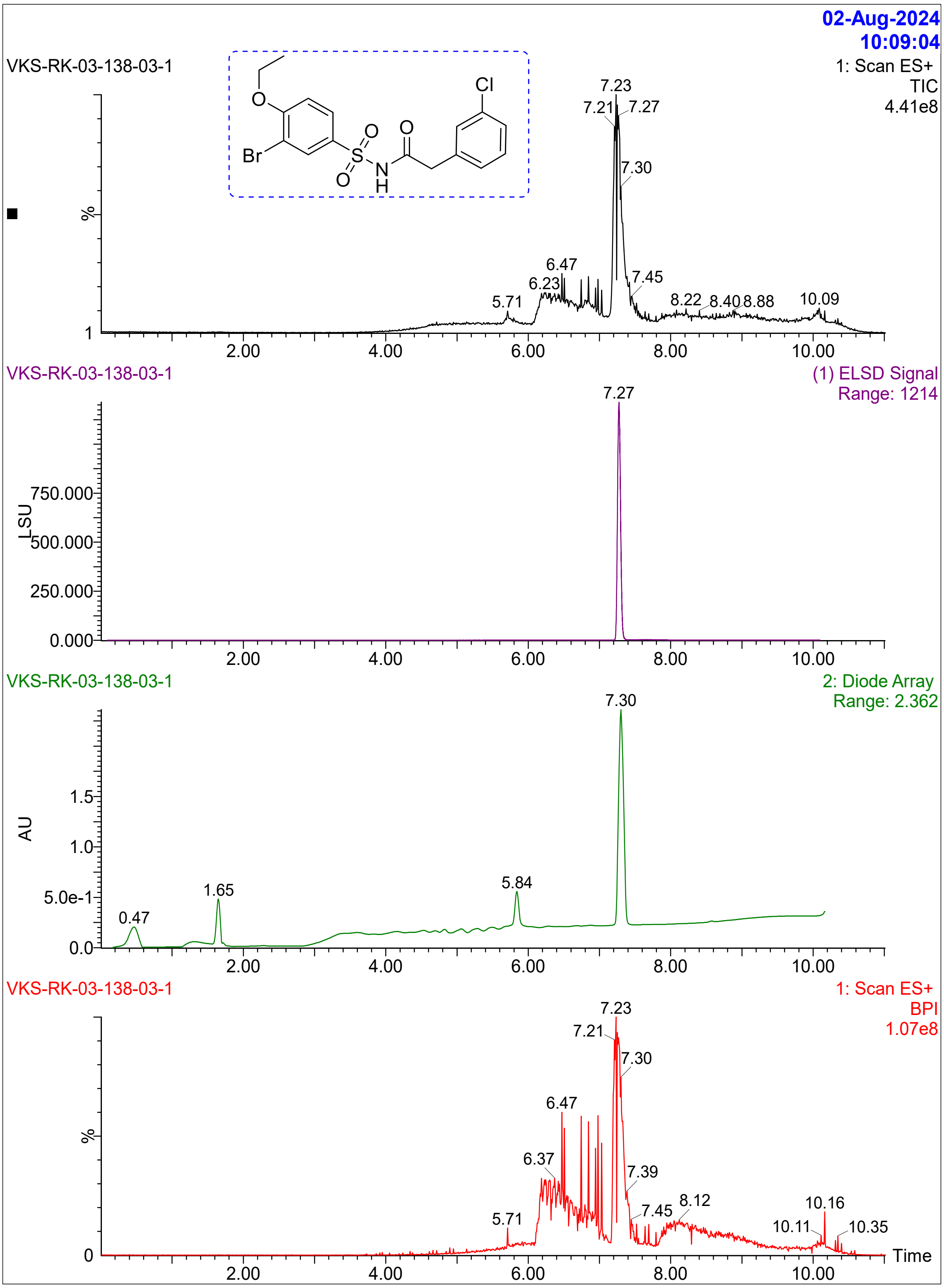

02-Aug-2024
10:09:04
VKS-RK-03-138-03-1
1: Scan ES+
TIC
4.41e8
Cl
O
Br
O
S
O
N
H
O
%
7.23
7.21
7.27
7.30
6.47
6.23
7.45
5.71
8.22
8.40
8.88
10.09
1
2.00
4.00
6.00
8.00
10.00
VKS-RK-03-138-03-1
(1) ELSD Signal
Range: 1214
7.27
750.000
LSU
500.000
250.000
0.000
2.00
4.00
6.00
8.00
10.00
VKS-RK-03-138-03-1
2: Diode Array
Range: 2.362
7.30
1.5
AU
1.0
5.0e-1
0.0
0.47
1.65
5.84
2.00
4.00
6.00
8.00
10.00
VKS-RK-03-138-03-1
1: Scan ES+
BPI
1.07e8
7.23
7.21
7.30
6.47
%
6.37
7.39
7.45
8.12
10.16
10.11
10.35
5.71
0
Time
2.00
4.00
6.00
8.00
10.00

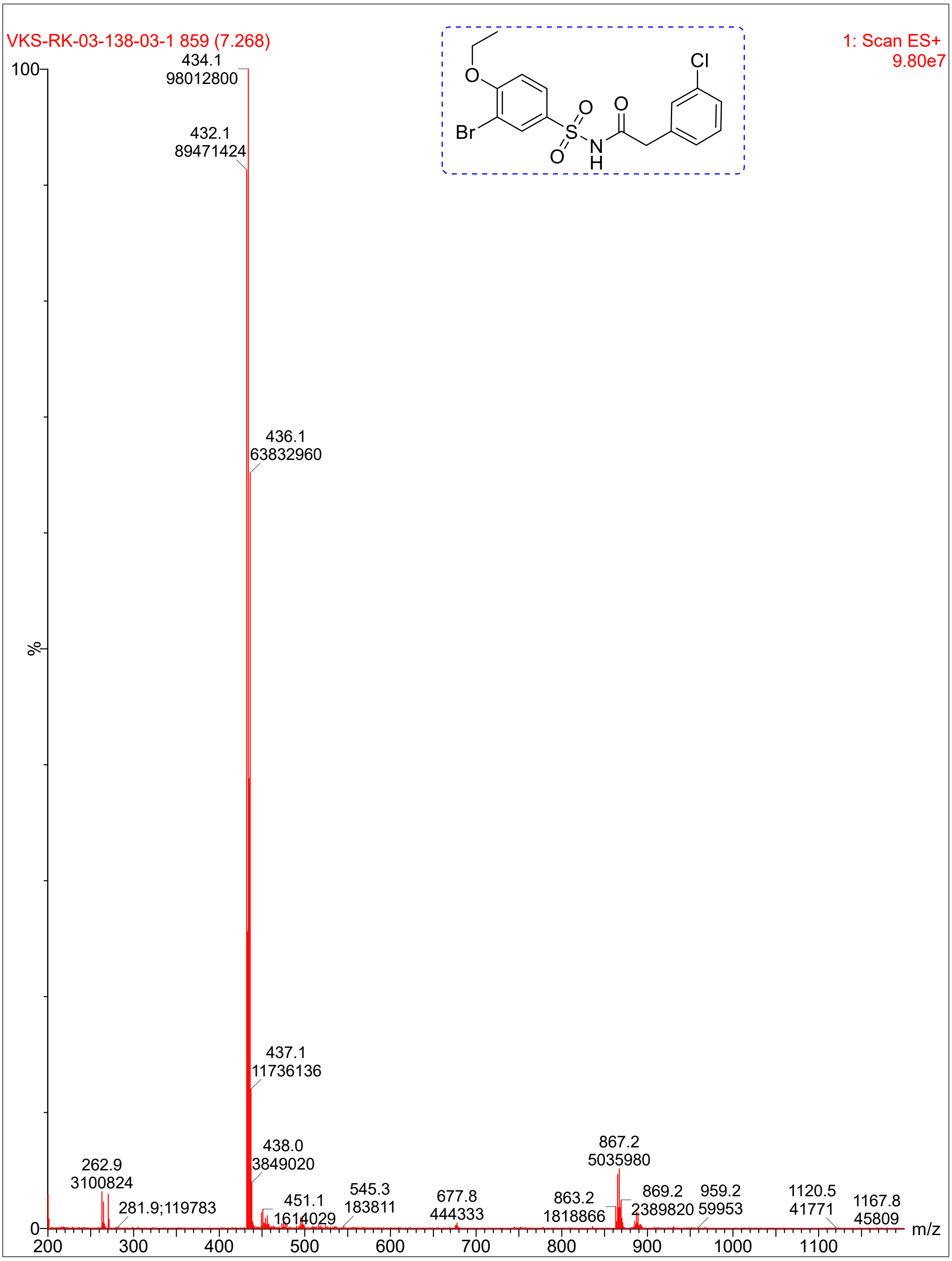
VKS-RK-03-138-03-1 859 (7.268)
1: Scan ES+
9.80e7
434.1
98012800
432.1
89471424
436.1
63832960
437.1
11736136
438.0
3849020
262.9
3100824
281.9;119783
451.1
1614029
545.3
183811
677.8
444333
863.2
1818866
867.2
5035980
869.2
2389820
959.2
59953
1120.5
41771
1167.8
45809
100
%
0
200
300
400
500
600
700
800
900
1000
1100
m/z
O
Br
S
O
O
N
H
O
Cl

**Compound 6**:

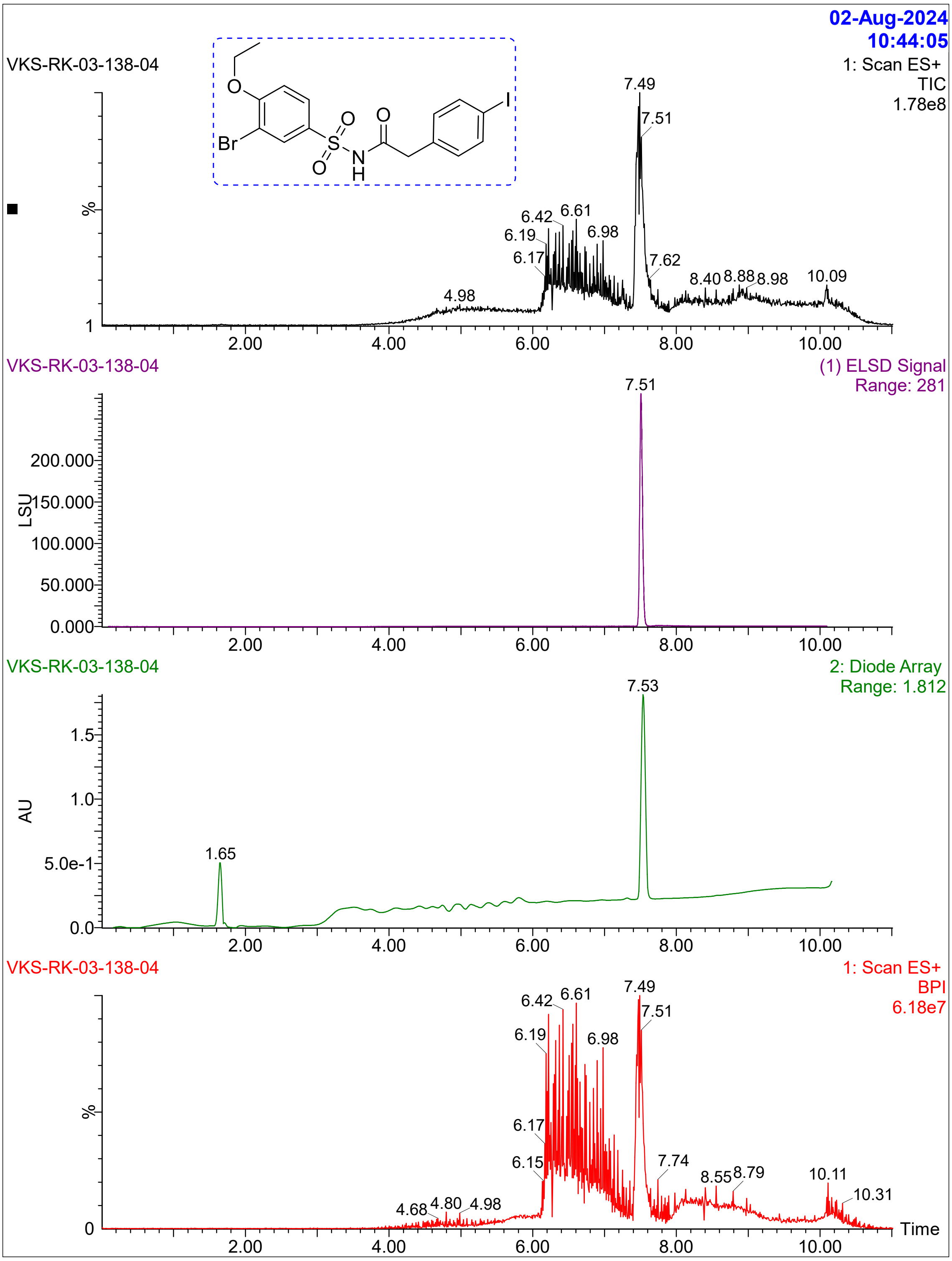

02-Aug-2024
10:44:05
VKS-RK-03-138-04
1: Scan ES+
TIC
1.78e8
Br
O
S
N
H
I
%
7.49
7.51
6.42
6.61
6.19
6.98
6.17
7.62
8.40
8.88
8.98
10.09
4.98
2.00
4.00
6.00
8.00
10.00
VKS-RK-03-138-04
(1) ELSD Signal
Range: 281
7.51
LSU
200.000
150.000
100.000
50.000
0.000
VKS-RK-03-138-04
2: Diode Array
Range: 1.812
7.53
AU
1.5
1.0
5.0e-1
0.0
1.65
VKS-RK-03-138-04
1: Scan ES+
BPI
6.18e7
6.42
6.61
7.49
7.51
6.19
6.98
6.17
6.15
7.74
8.55
8.79
10.11
10.31
4.68
4.80
4.98
Time

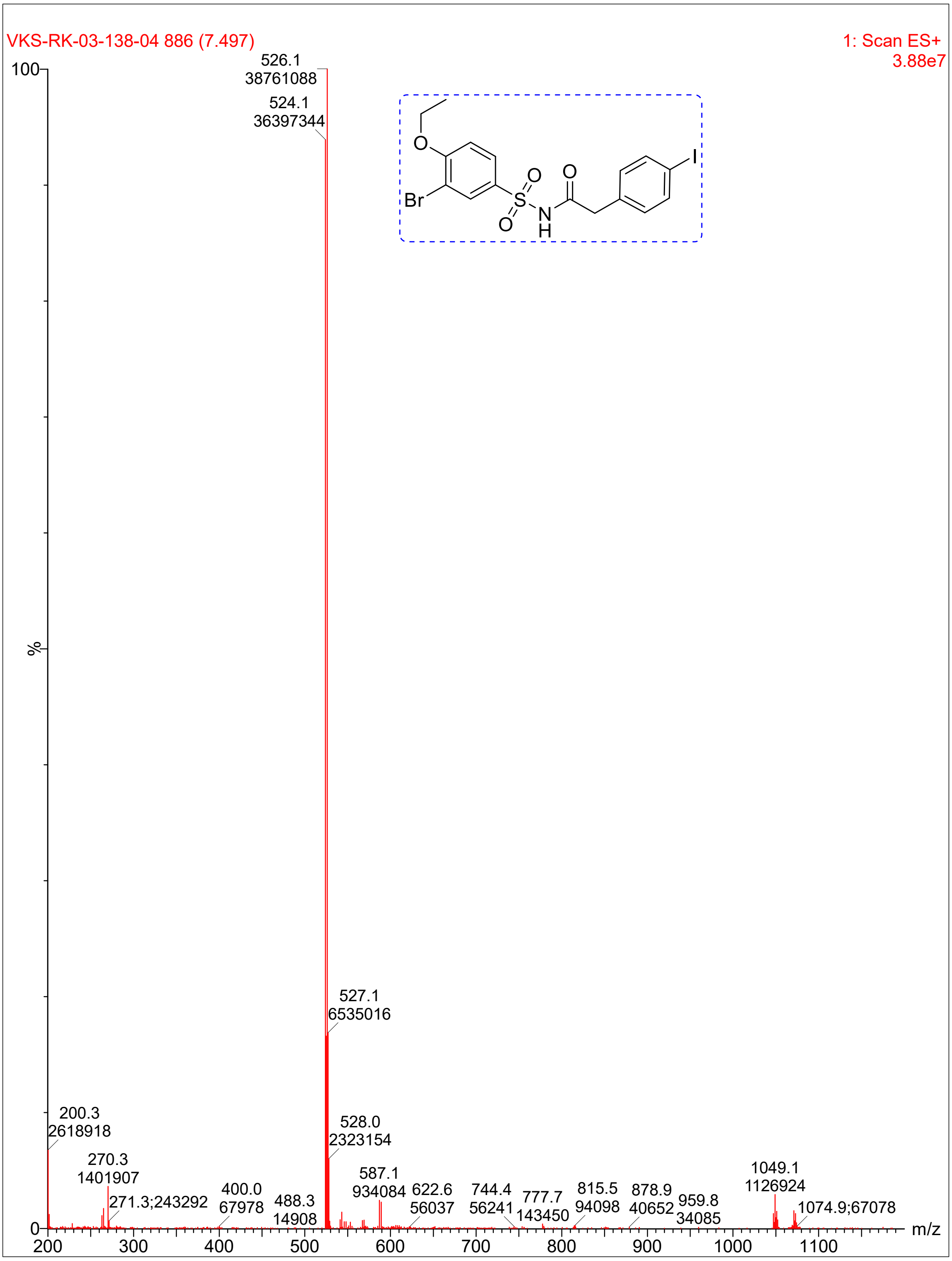
VKS-RK-03-138-04 886 (7.497)
1: Scan ES+
3.88e7
526.1
38761088
524.1
36397344
527.1
6535016
528.0
2323154
200.3
2618918
270.3
1401907
271.3;243292
400.0
67978
488.3
14908
587.1
934084
622.6
56037
744.4
56241
777.7
143450
815.5
94098
878.9
40652
959.8
34085
1049.1
1126924
1074.9;67078
%
m/z
Br
O
S
N
H
I

**Compound 16:**

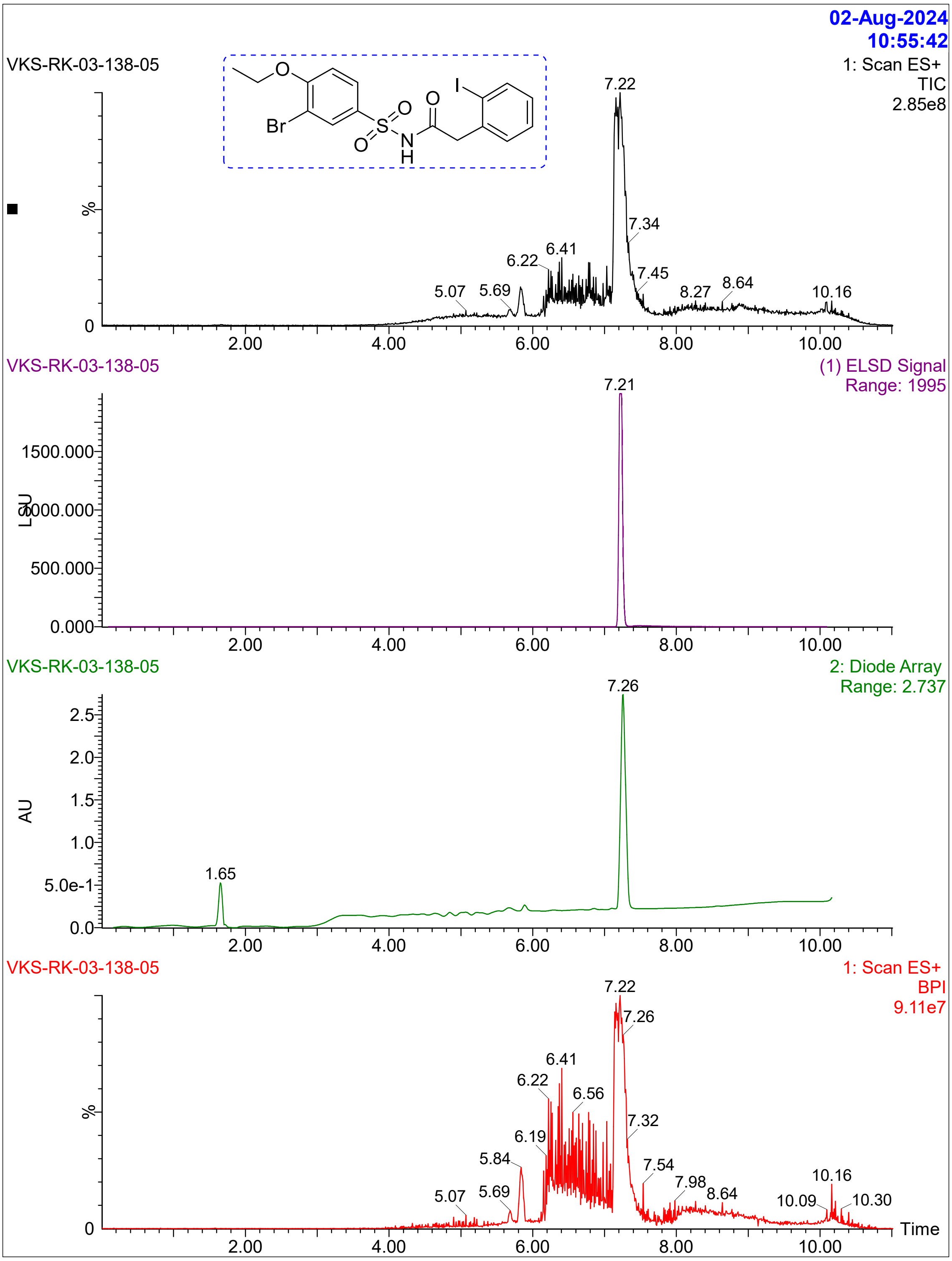
02-Aug-2024
10:55:42
VKS-RK-03-138-05
1: Scan ES+
TIC
2.85e8
Br
O
S
N
H
I
7.22
7.34
7.45
6.41
6.22
5.07
5.69
8.27
8.64
10.16
%
0
2.00
4.00
6.00
8.00
10.00
VKS-RK-03-138-05
(1) ELSD Signal
Range: 1995
7.21
1500.000
1000.000
500.000
0.000
LSU
VKS-RK-03-138-05
2: Diode Array
Range: 2.737
7.26
1.65
2.5
2.0
1.5
1.0
5.0e-1
0.0
AU
VKS-RK-03-138-05
1: Scan ES+
BPI
9.11e7
7.22
7.26
6.41
6.22
6.56
6.19
7.32
5.84
7.54
5.07
5.69
7.98
8.64
10.16
10.09
10.30
Time

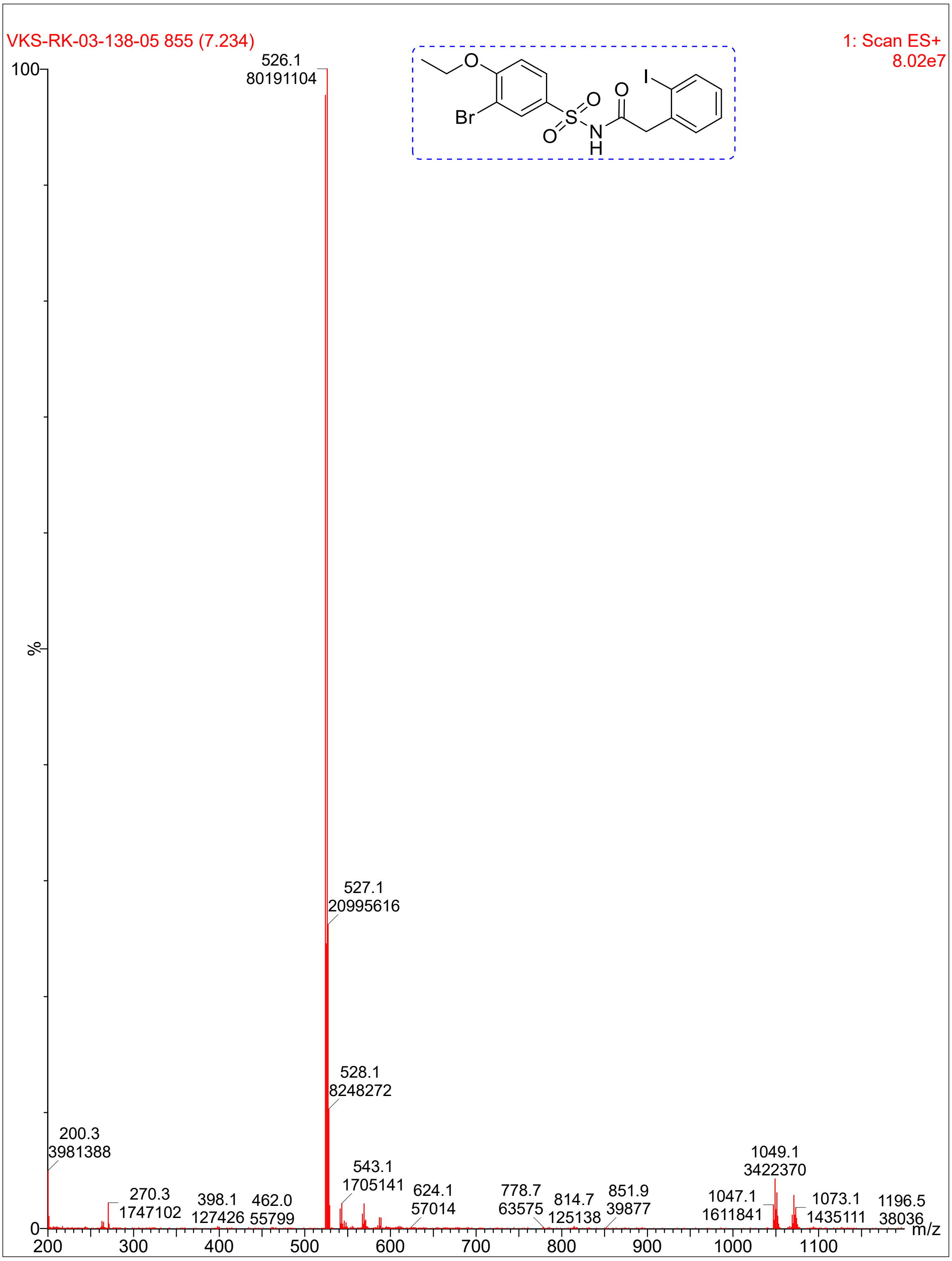

VKS-RK-03-138-05 855 (7.234)
1: Scan ES+
8.02e7
526.1
80191104
527.1
20995616
528.1
8248272
200.3
3981388
270.3
1747102
398.1
127426
462.0
55799
543.1
1705141
624.1
57014
778.7
63575
814.7
125138
851.9
39877
1047.1
1611841
1049.1
3422370
1073.1
1435111
1196.5
38036
%
m/z
Br
O
S
N
H
I

**Compound 7:**

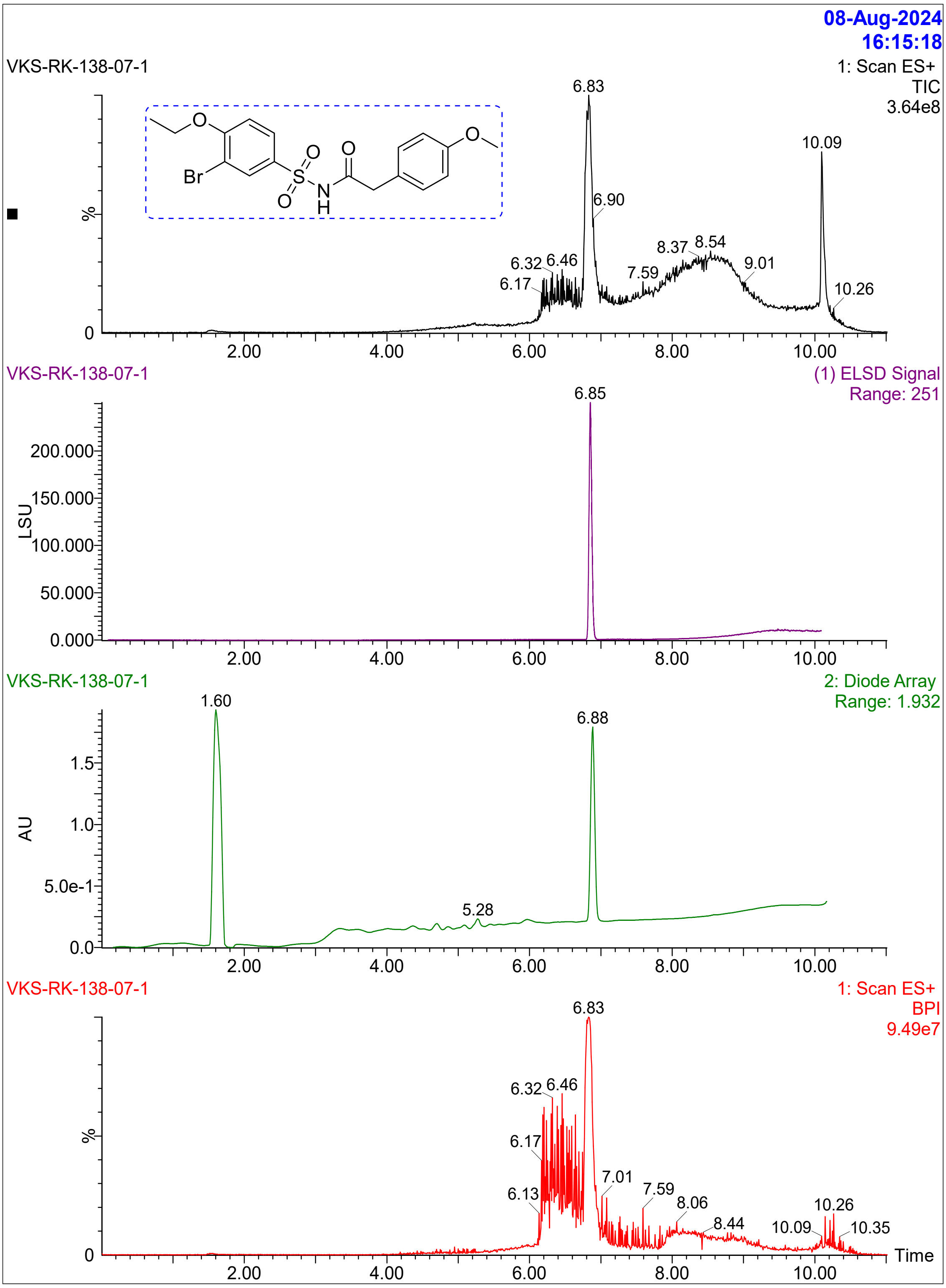

08-Aug-2024
16:15:18
VKS-RK-138-07-1
1: Scan ES+
TIC
3.64e8
Br
O
S
N
H
6.83
10.09
6.90
8.37
8.54
9.01
7.59
6.32
6.46
6.17
10.26
%
0
2.00
4.00
6.00
8.00
10.00
VKS-RK-138-07-1
(1) ELSD Signal
Range: 251
6.85
200.000
150.000
100.000
50.000
0.000
LSU
VKS-RK-138-07-1
2: Diode Array
Range: 1.932
1.60
6.88
5.28
1.5
1.0
5.0e-1
0.0
AU
VKS-RK-138-07-1
1: Scan ES+
BPI
9.49e7
6.83
6.32
6.46
6.17
6.13
7.01
7.59
8.06
8.44
10.09
10.26
10.35
Time

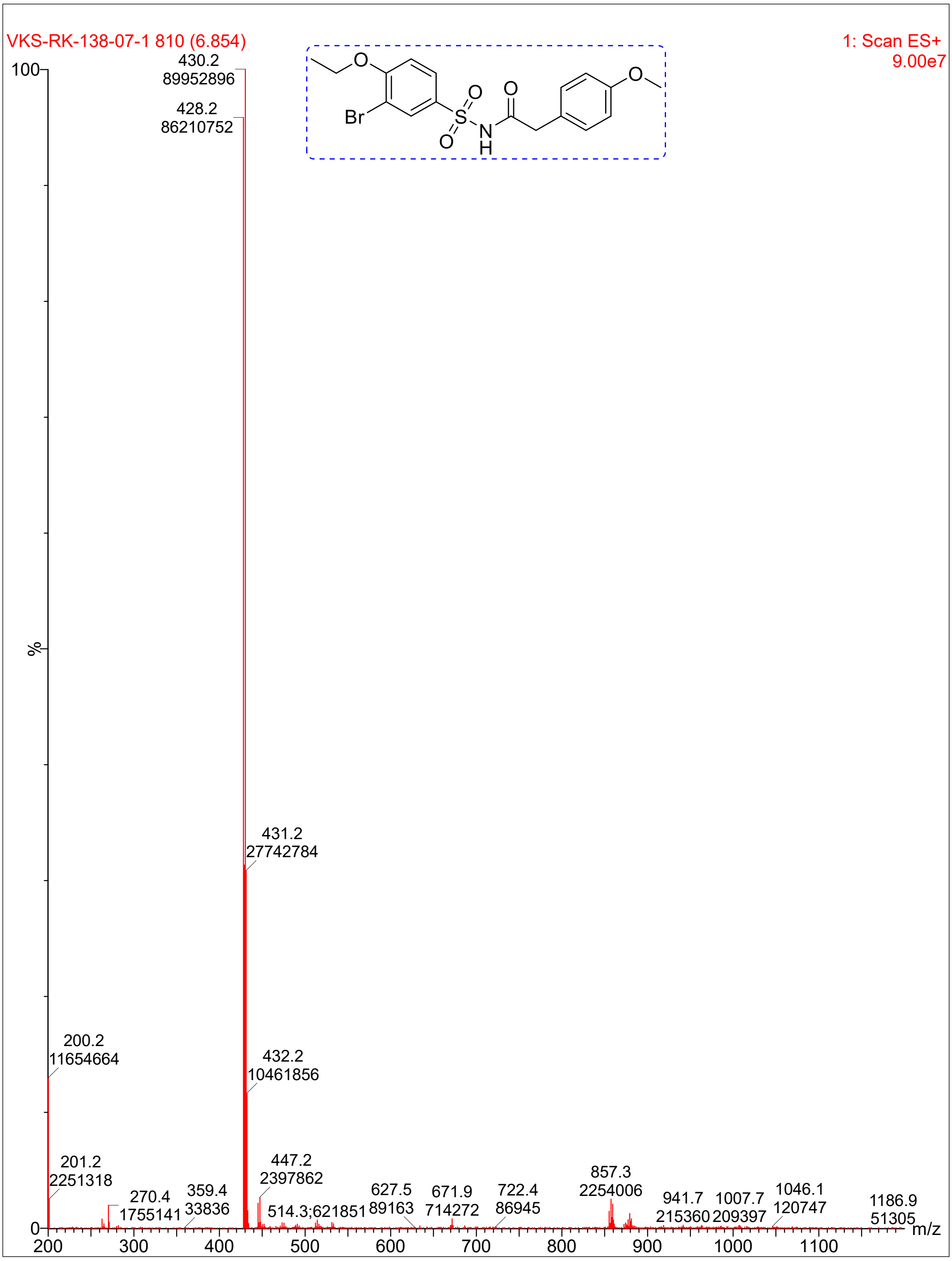

VKS-RK-138-07-1 810 (6.854)
1: Scan ES+
9.00e7
430.2
89952896
428.2
86210752
431.2
27742784
432.2
10461856
200.2
11654664
201.2
2251318
270.4
1755141
359.4
33836
447.2
2397862
514.3;621851
627.5
89163
671.9
714272
722.4
86945
857.3
2254006
941.7
215360
1007.7
209397
1046.1
120747
1186.9
51305
%
m/z
Br

**Compound 13:**

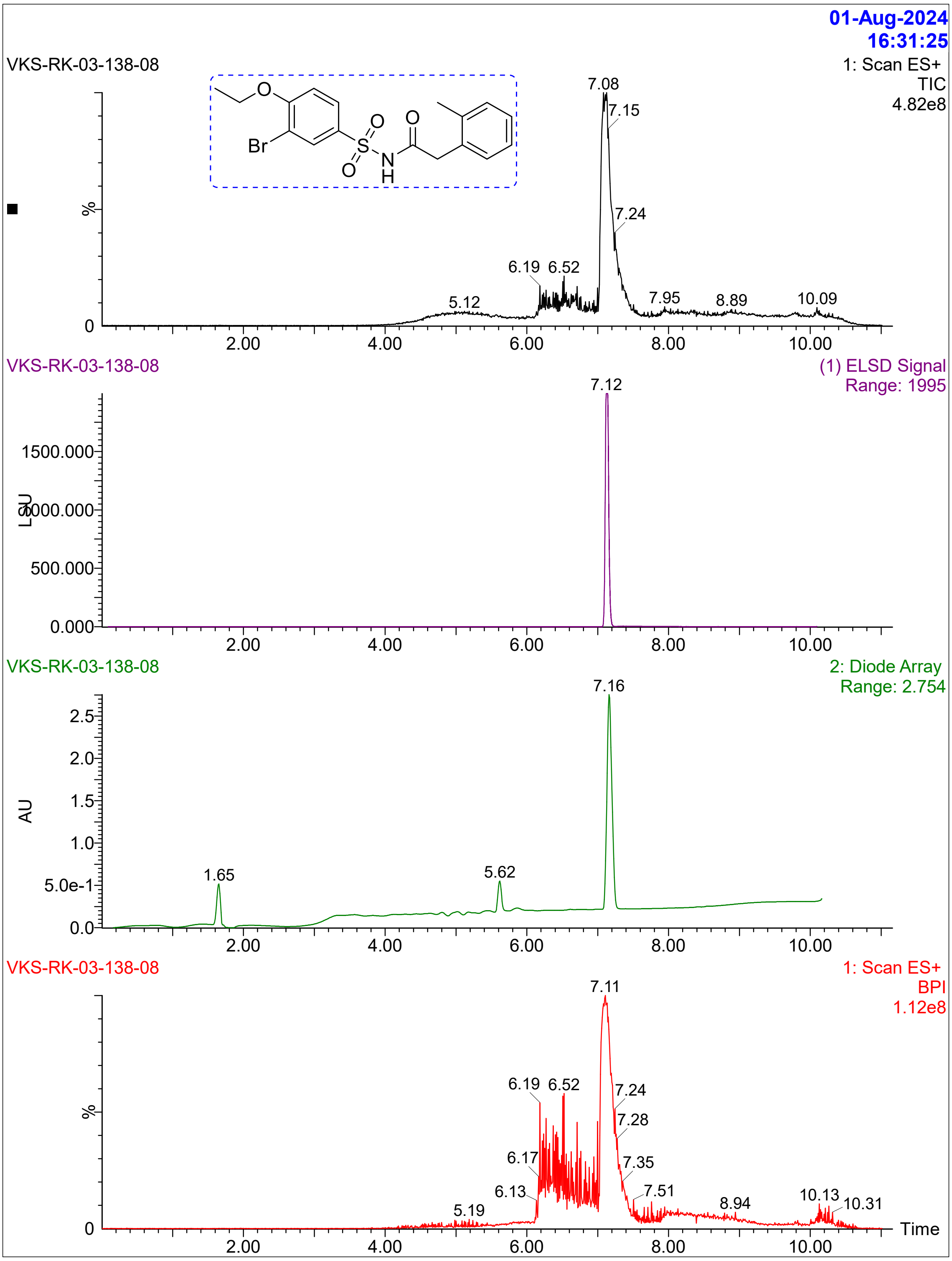

01-Aug-2024
16:31:25
VKS-RK-03-138-08
1: Scan ES+
TIC
4.82e8
Br
O
S
N
H
%
0
7.08
7.15
7.24
6.19
6.52
5.12
7.95
8.89
10.09
2.00
4.00
6.00
8.00
10.00
VKS-RK-03-138-08
(1) ELSD Signal
Range: 1995
7.12
LSU
1500.000
1000.000
500.000
0.000
VKS-RK-03-138-08
2: Diode Array
Range: 2.754
7.16
AU
2.5
2.0
1.5
1.0
5.0e-1
0.0
1.65
5.62
VKS-RK-03-138-08
1: Scan ES+
BPI
1.12e8
7.11
6.19
6.52
7.24
7.28
6.17
7.35
6.13
7.51
5.19
8.94
10.13
10.31
Time

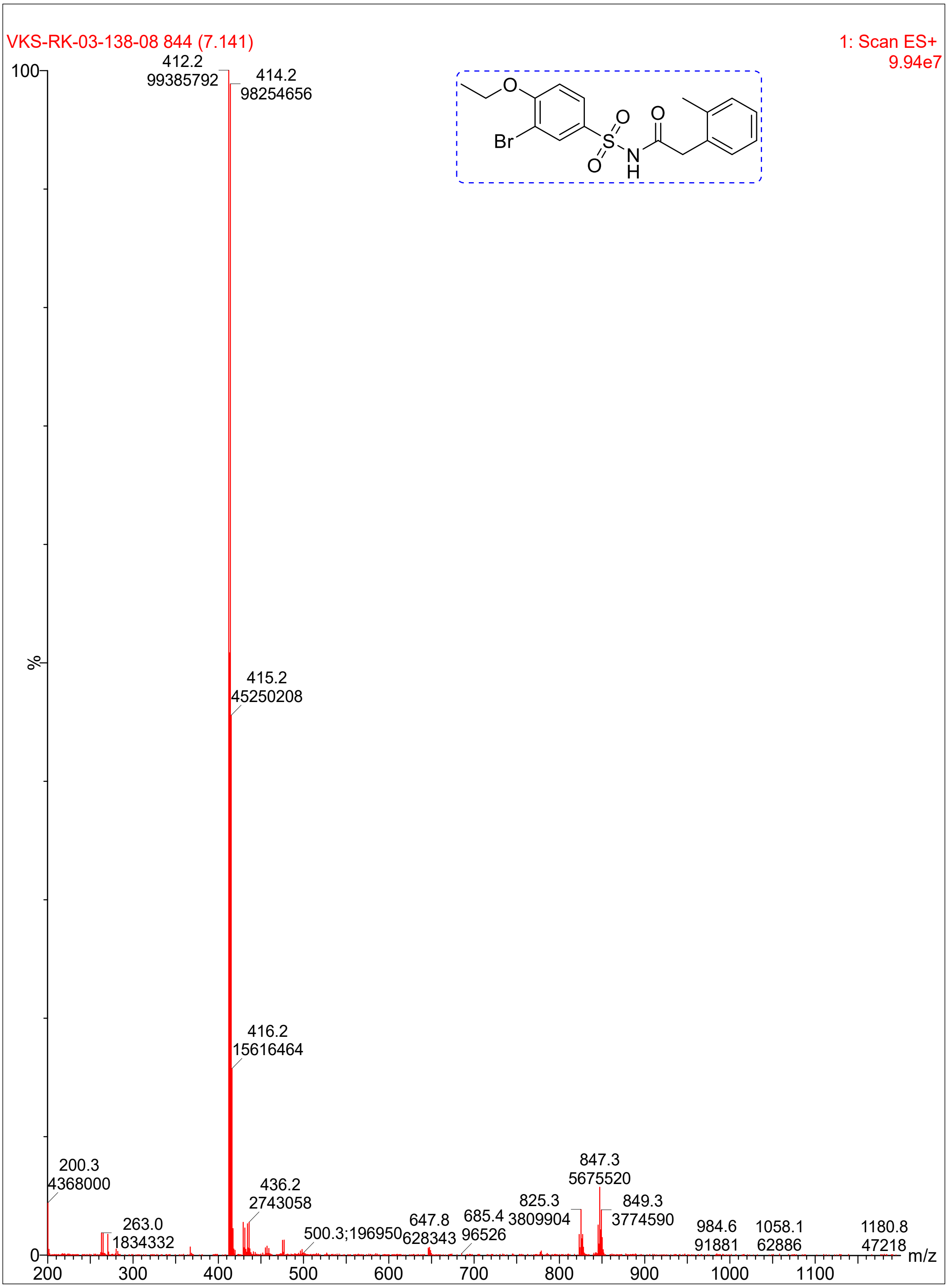

VKS-RK-03-138-08 844 (7.141)
1: Scan ES+
9.94e7
100
%
0
412.2
99385792
414.2
98254656
415.2
45250208
416.2
15616464
200.3
4368000
263.0
1834332
436.2
2743058
500.3;196950
647.8
628343
685.4
96526
825.3
3809904
847.3
5675520
849.3
3774590
984.6
91881
1058.1
62886
1180.8
47218
200
300
400
500
600
700
800
900
1000
1100
m/z
O
Br
S
O
O
N
H
O

**Compound 21**:

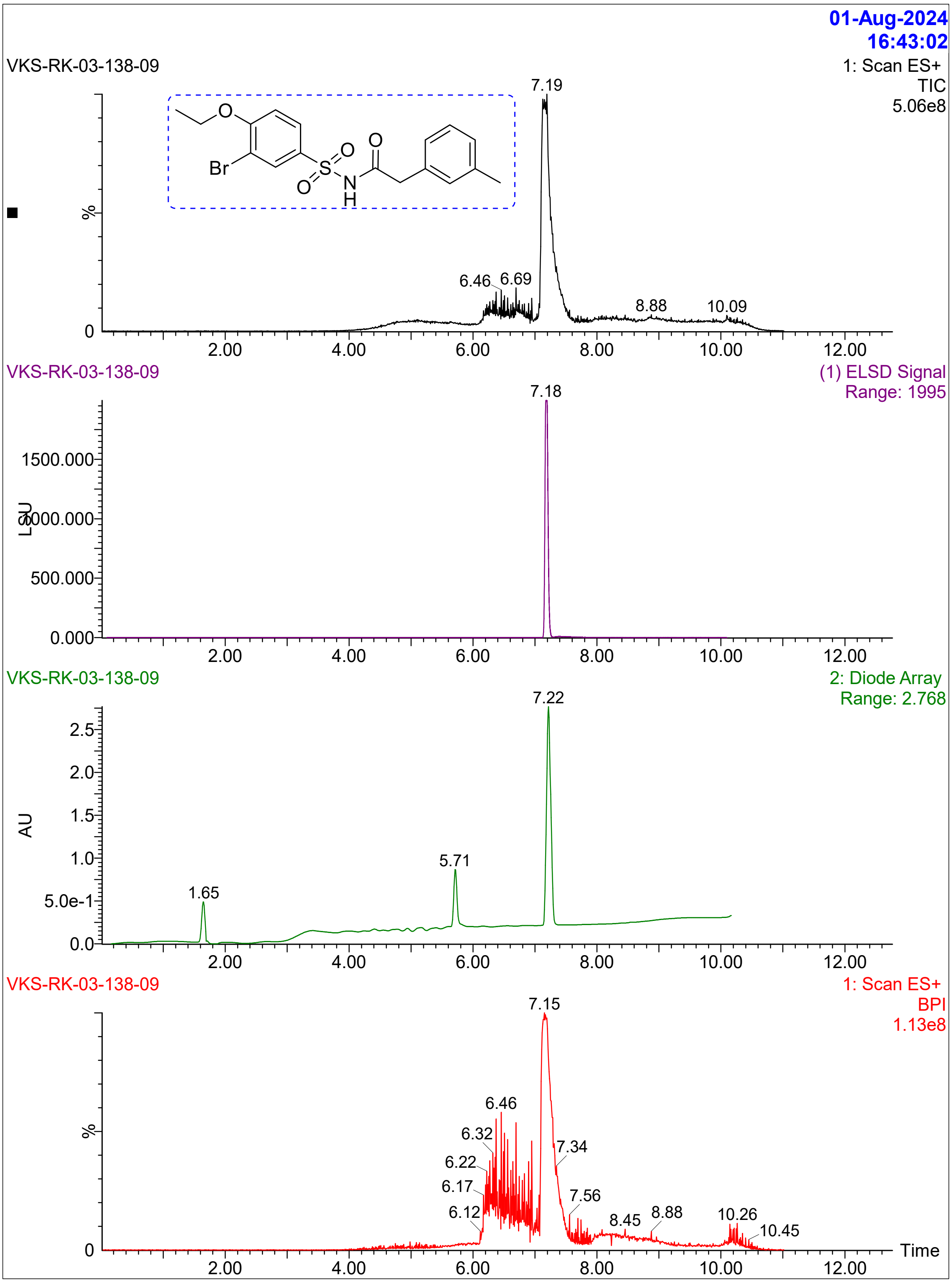

01-Aug-2024
16:43:02
VKS-RK-03-138-09
1: Scan ES+
TIC
5.06e8
7.19
6.46
6.69
8.88
10.09
Br
O
S
N
H
%
0
2.00
4.00
6.00
8.00
10.00
12.00
VKS-RK-03-138-09
(1) ELSD Signal
Range: 1995
7.18
1500.000
1000.000
500.000
0.000
LSU
VKS-RK-03-138-09
2: Diode Array
Range: 2.768
7.22
5.71
1.65
2.5
2.0
1.5
1.0
5.0e-1
0.0
AU
VKS-RK-03-138-09
1: Scan ES+
BPI
1.13e8
7.15
6.46
6.32
6.22
6.17
6.12
7.34
7.56
8.45
8.88
10.26
10.45
Time

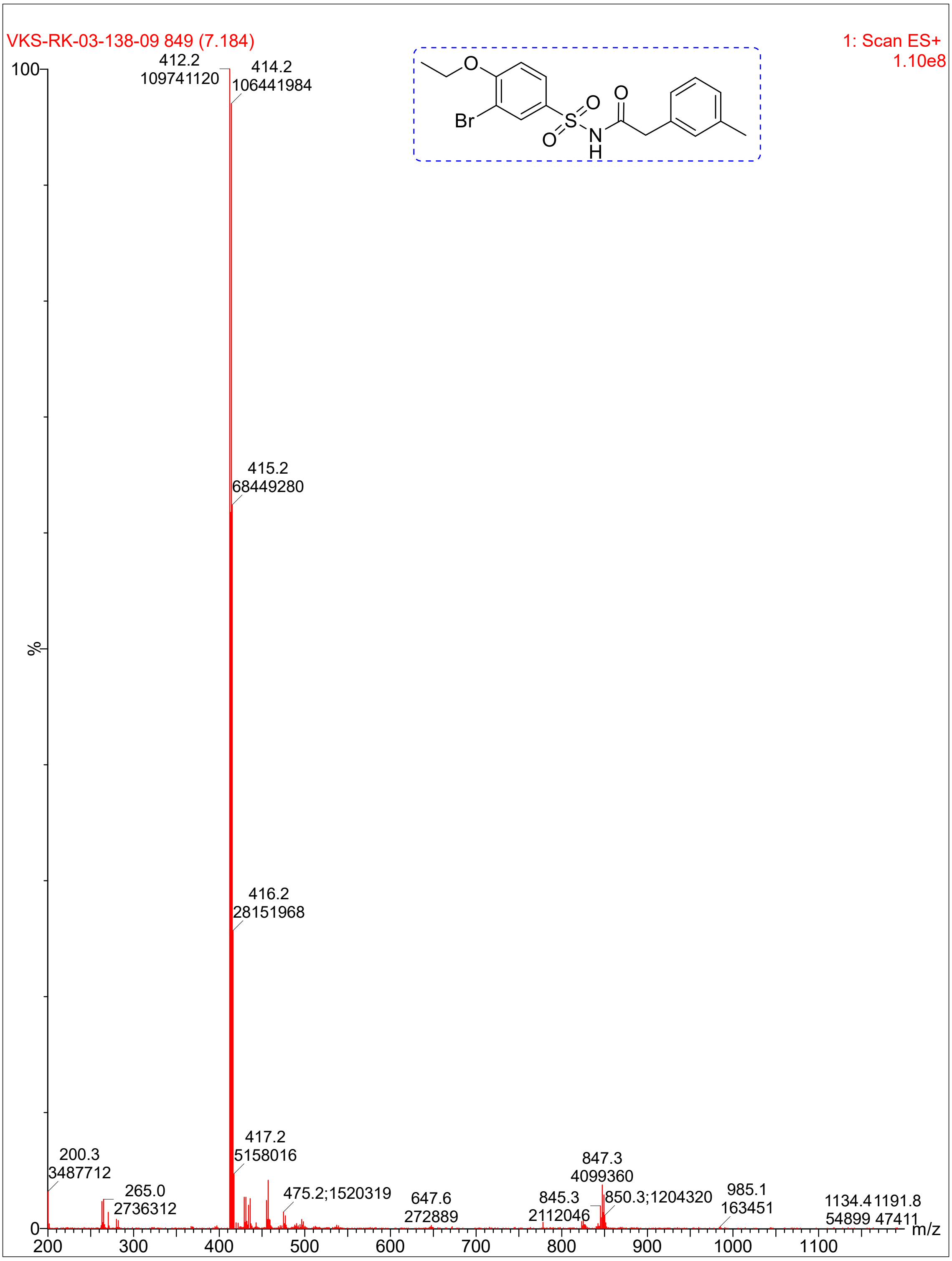

VKS-RK-03-138-09 849 (7.184)
1: Scan ES+
1.10e8
Br
O
S
N
H
412.2
109741120
414.2
106441984
415.2
68449280
416.2
28151968
417.2
5158016
200.3
3487712
265.0
2736312
475.2;1520319
647.6
272889
845.3
2112046
847.3
4099360
850.3;1204320
985.1
163451
1134.4
54899
1191.8
47411
100
%
0
m/z
200
300
400
500
600
700
800
900
1000
1100

**Compound 14**:

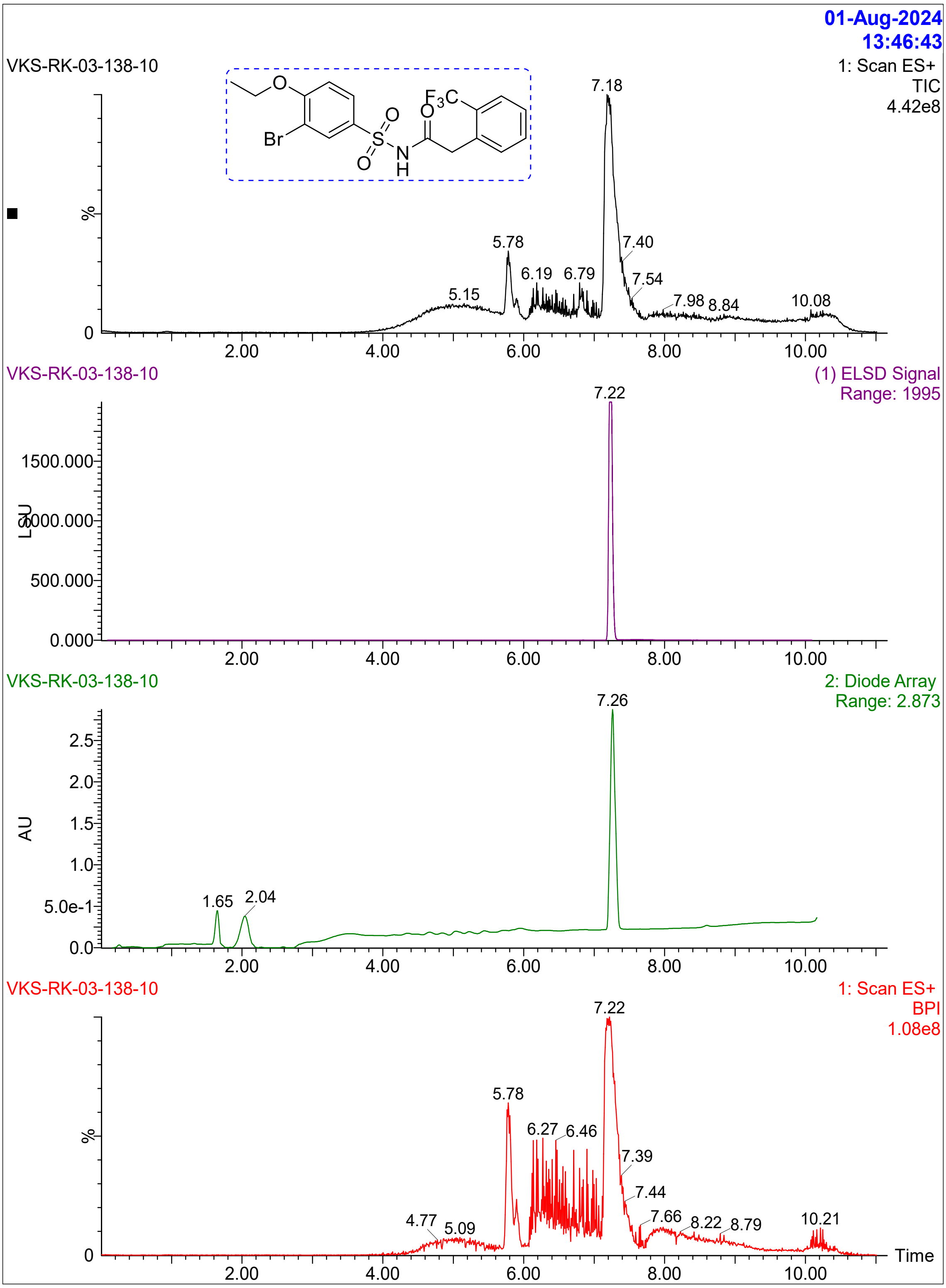

01-Aug-2024
13:46:43
VKS-RK-03-138-10
1: Scan ES+
TIC
4.42e8
Br
O
S
N
H
F3C
7.18
5.78
7.40
6.19
6.79
7.54
5.15
7.98
8.84
10.08
%
0
2.00
4.00
6.00
8.00
10.00
VKS-RK-03-138-10
(1) ELSD Signal
Range: 1995
7.22
1500.000
1000.000
500.000
0.000
LSU
VKS-RK-03-138-10
2: Diode Array
Range: 2.873
7.26
2.5
2.0
1.5
1.0
5.0e-1
0.0
AU
1.65
2.04
VKS-RK-03-138-10
1: Scan ES+
BPI
1.08e8
7.22
5.78
6.27
6.46
7.39
7.44
4.77
5.09
7.66
8.22
8.79
10.21
Time

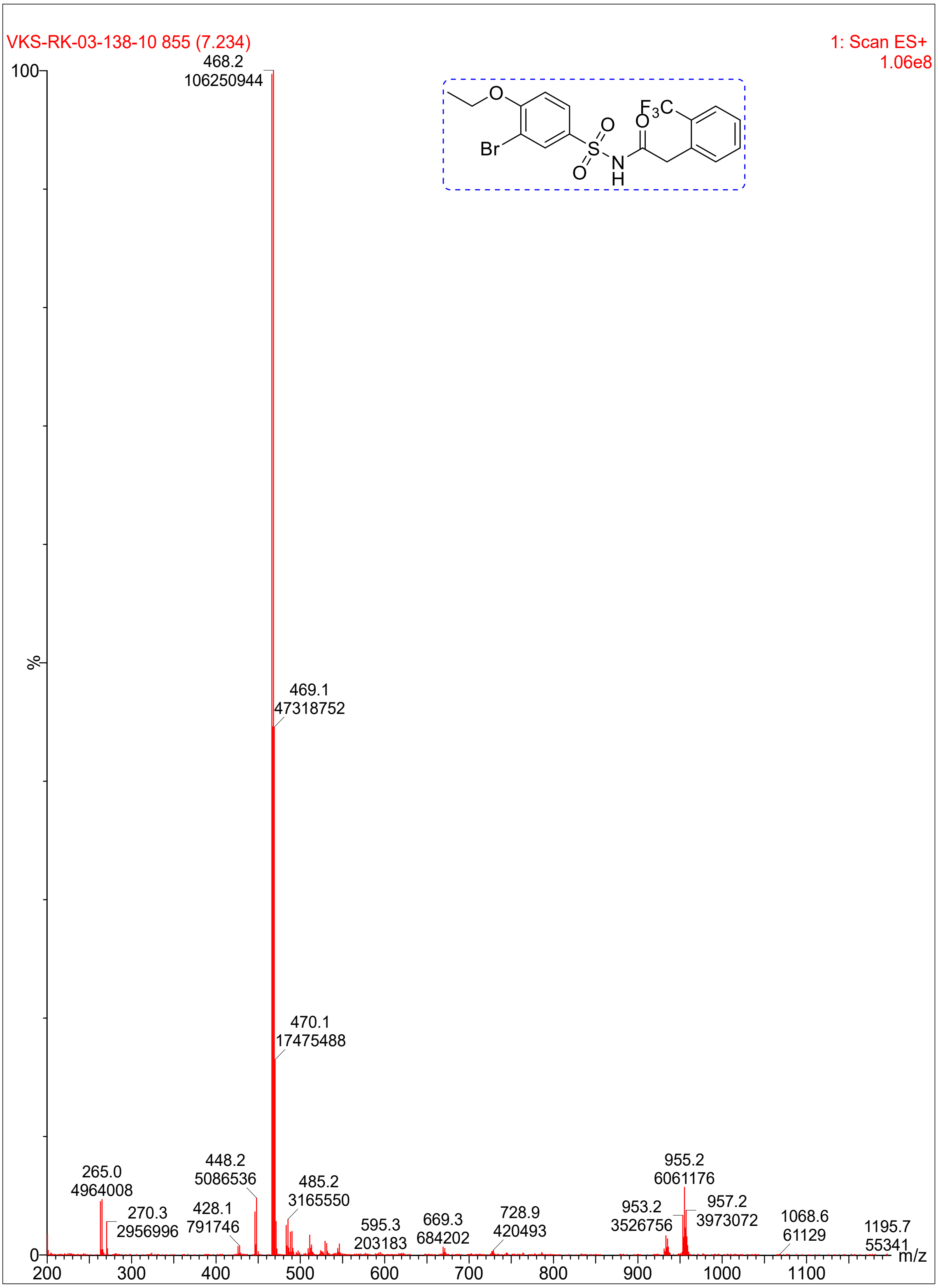

VKS-RK-03-138-10 855 (7.234)
1: Scan ES+
1.06e8
468.2
106250944
469.1
47318752
470.1
17475488
265.0
4964008
270.3
2956996
428.1
791746
448.2
5086536
485.2
3165550
595.3
203183
669.3
684202
728.9
420493
953.2
3526756
955.2
6061176
957.2
3973072
1068.6
61129
1195.7
55341
100
%
0
200
300
400
500
600
700
800
900
1000
1100
m/z
Br
O
S
N
H

**Compound 20:**

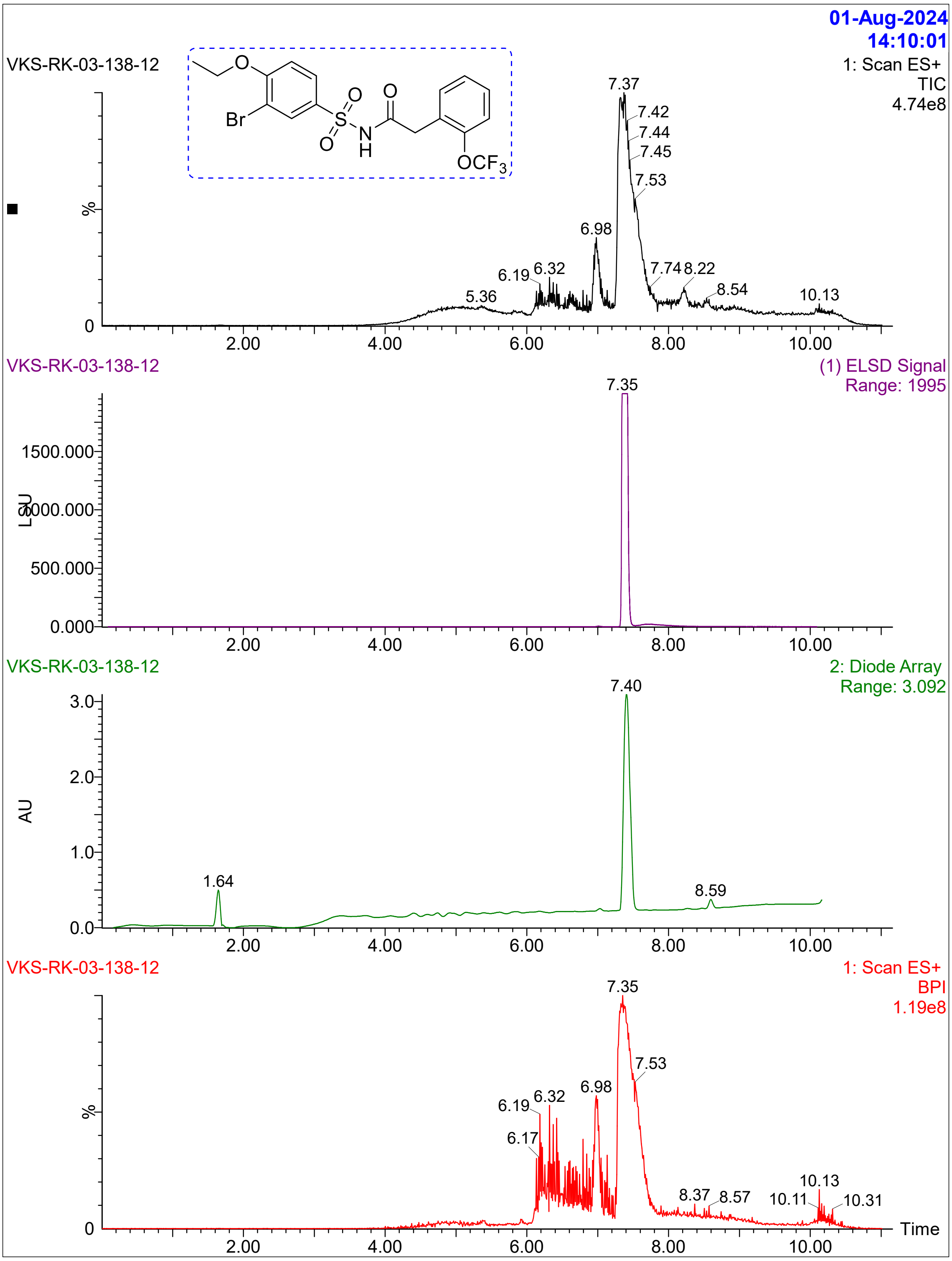

01-Aug-2024
14:10:01
VKS-RK-03-138-12
1: Scan ES+
TIC
4.74e8
Br
O
S
N
H
OCF3
7.37
7.42
7.44
7.45
7.53
6.98
6.19
6.32
5.36
7.74
8.22
8.54
10.13
%
0
2.00
4.00
6.00
8.00
10.00
VKS-RK-03-138-12
(1) ELSD Signal
Range: 1995
7.35
1500.000
1000.000
500.000
0.000
LSU
VKS-RK-03-138-12
2: Diode Array
Range: 3.092
7.40
3.0
2.0
1.0
0.0
AU
1.64
8.59
VKS-RK-03-138-12
1: Scan ES+
BPI
1.19e8
7.35
7.53
6.98
6.19
6.32
6.17
8.37
8.57
10.13
10.11
10.31
Time

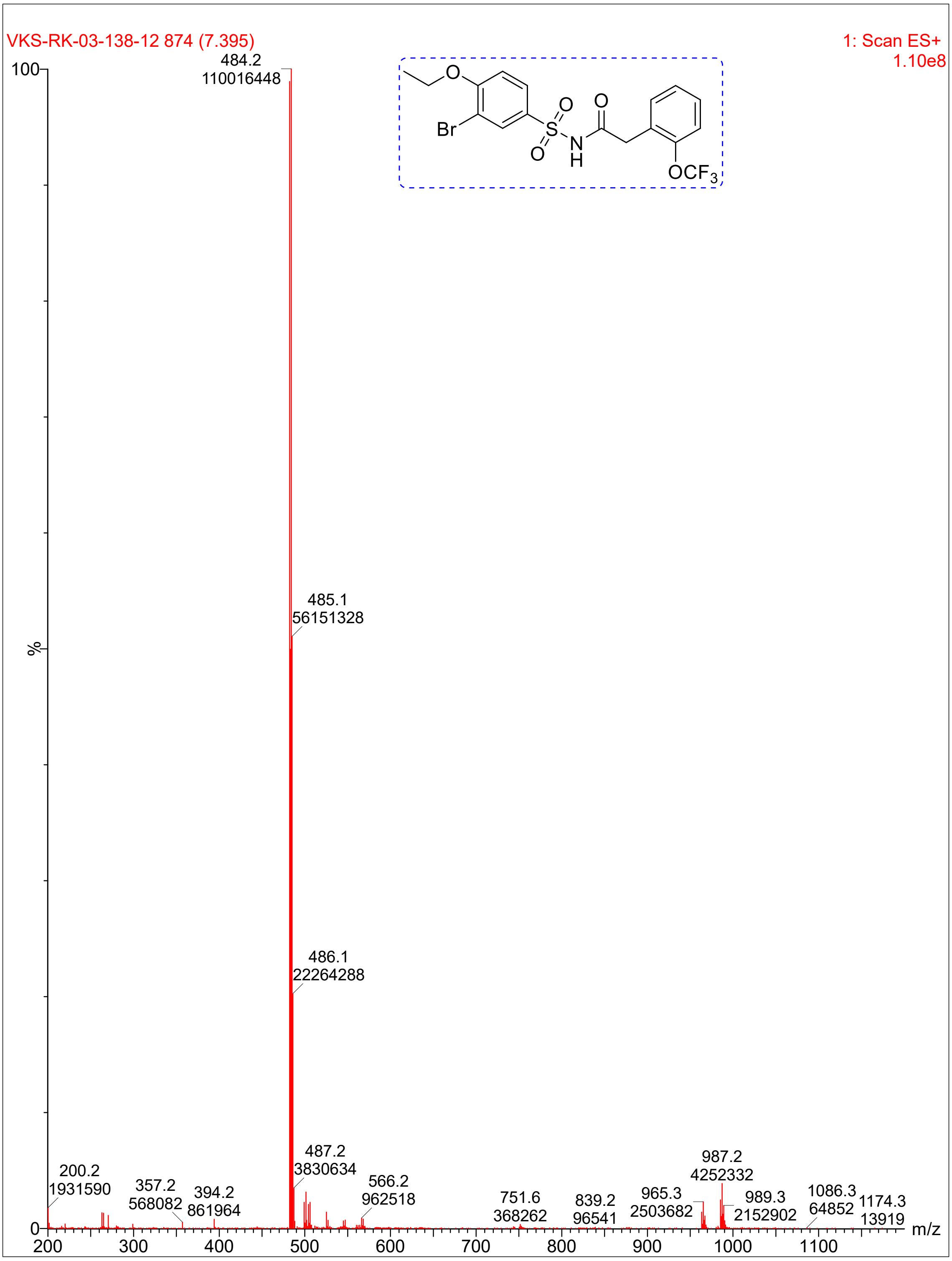

VKS-RK-03-138-12 874 (7.395)
1: Scan ES+
1.10e8
484.2
110016448
485.1
56151328
486.1
22264288
487.2
3830634
200.2
1931590
357.2
568082
394.2
861964
566.2
962518
751.6
368262
839.2
96541
965.3
2503682
987.2
4252332
989.3
2152902
1086.3
64852
1174.3
13919
%
m/z
Br
O
S
N
H
OCF3

**Compound 11**:

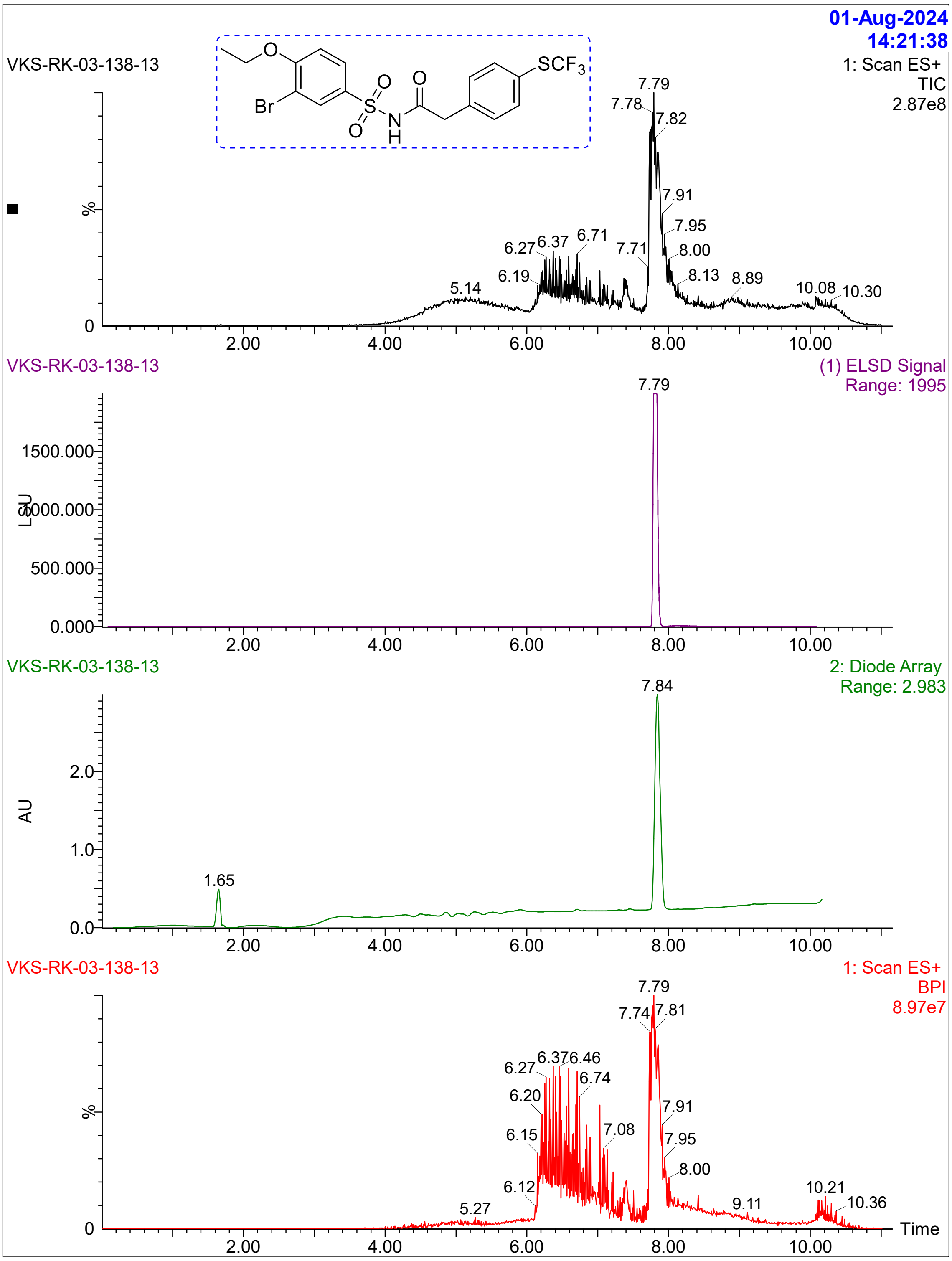

01-Aug-2024
14:21:38
VKS-RK-03-138-13
1: Scan ES+
TIC
2.87e8
(1) ELSD Signal
Range: 1995
2: Diode Array
Range: 2.983
1: Scan ES+
BPI
8.97e7
Time

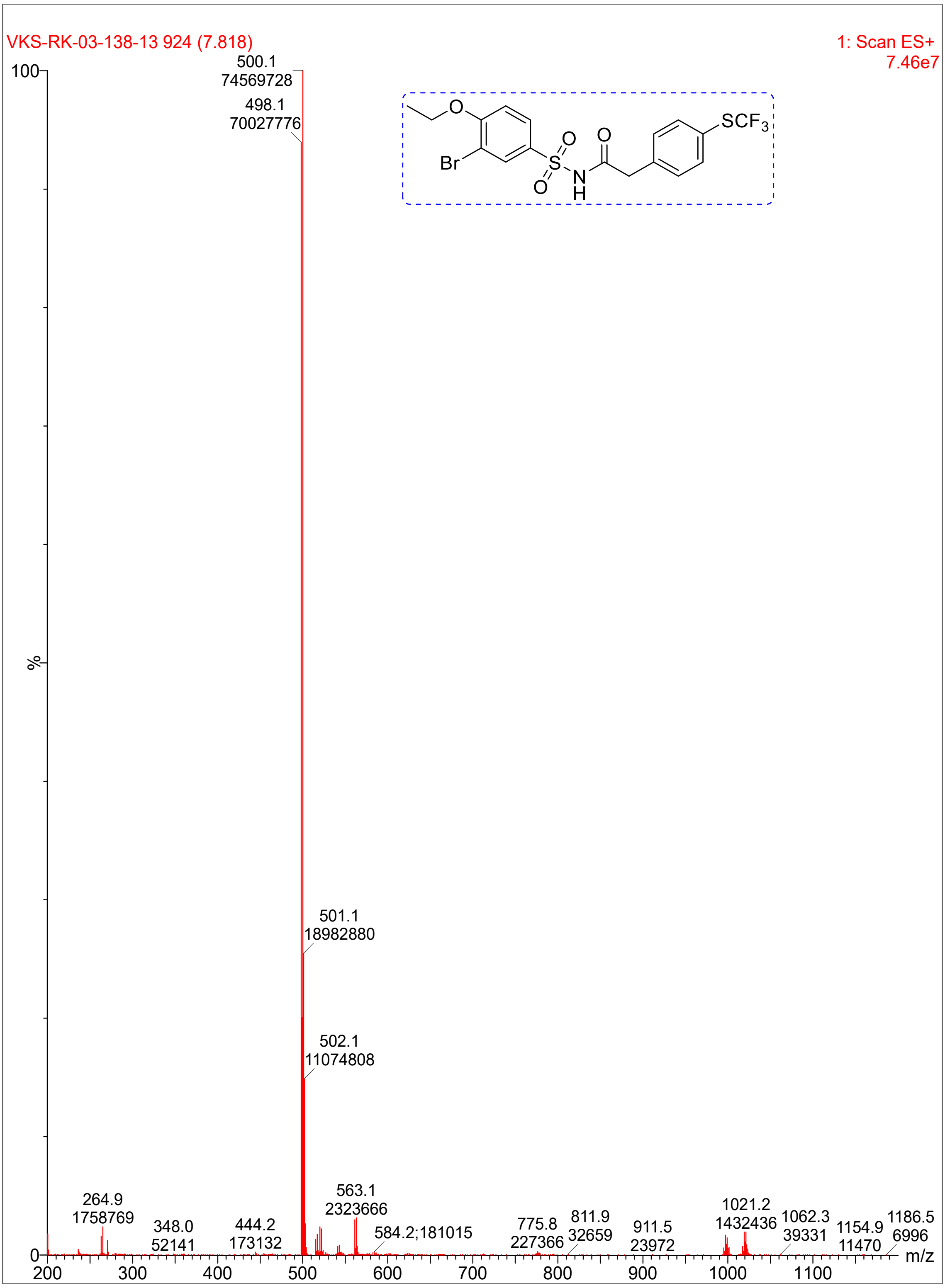
VKS-RK-03-138-13 924 (7.818)
1: Scan ES+
7.46e7
100
%
0
500.1
74569728
498.1
70027776
501.1
18982880
502.1
11074808
264.9
1758769
348.0
52141
444.2
173132
563.1
2323666
584.2;181015
775.8
227366
811.9
32659
911.5
23972
1021.2
1432436
1062.3
39331
1154.9
11470
1186.5
6996
200
300
400
500
600
700
800
900
1000
1100
m/z
Br
O
O
O
O
S
N
H
SCF3

**Compound 10**:

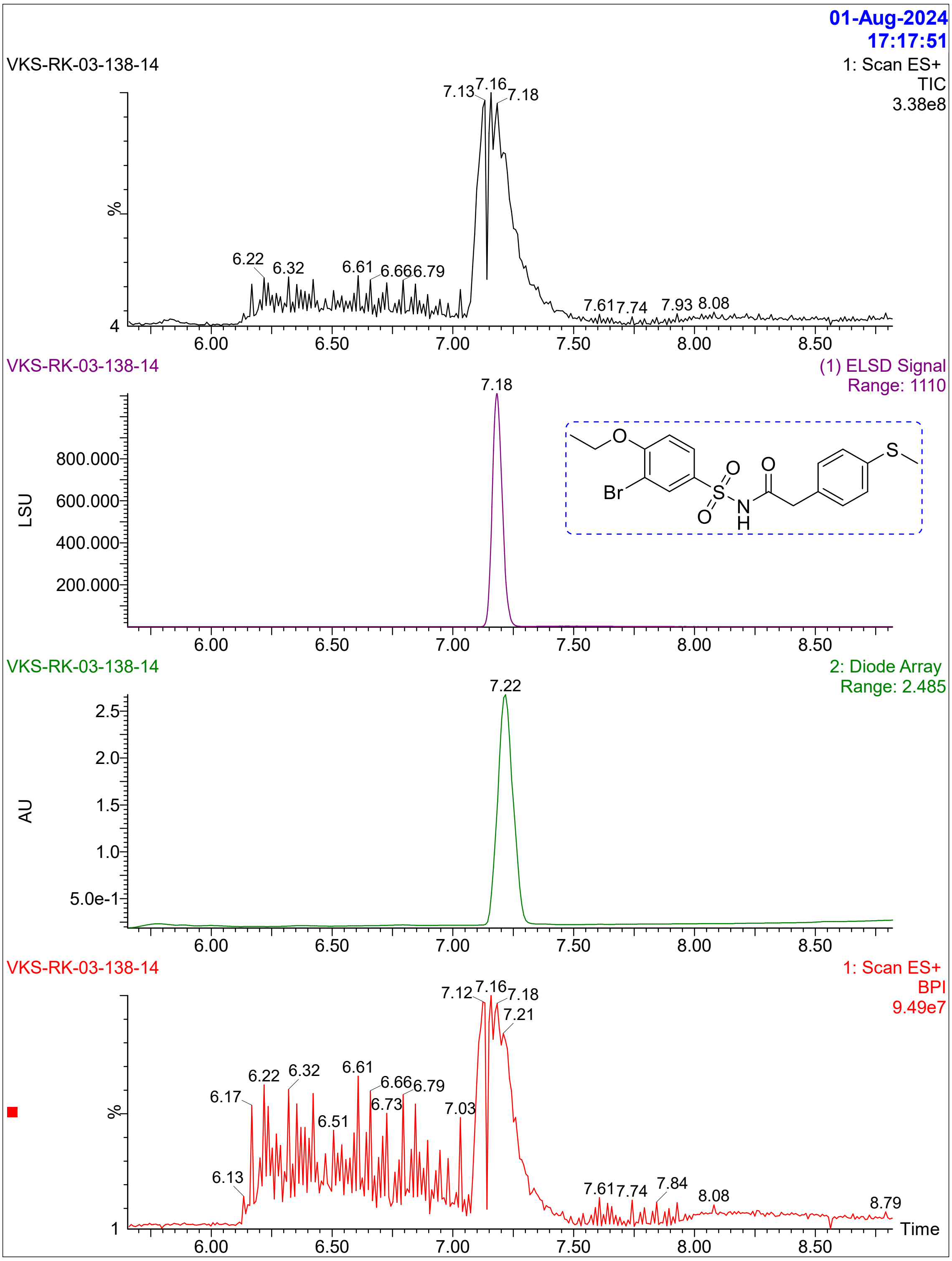
01-Aug-2024
17:17:51
VKS-RK-03-138-14
1: Scan ES+
TIC
3.38e8
7.13
7.16
7.18
6.22
6.32
6.61
6.666.79
7.617.74
7.93
8.08
%
4
6.00
6.50
7.00
7.50
8.00
8.50
VKS-RK-03-138-14
(1) ELSD Signal
Range: 1110
7.18
800.000
600.000
400.000
200.000
LSU
O
Br
S
O
O
N
H
O
S
VKS-RK-03-138-14
2: Diode Array
Range: 2.485
7.22
2.5
2.0
1.5
1.0
5.0e-1
AU
VKS-RK-03-138-14
1: Scan ES+
BPI
9.49e7
7.12
7.16
7.18
7.21
6.22
6.32
6.61
6.66
6.79
6.17
6.73
6.51
7.03
6.13
7.617.74
7.84
8.08
8.79
1
Time

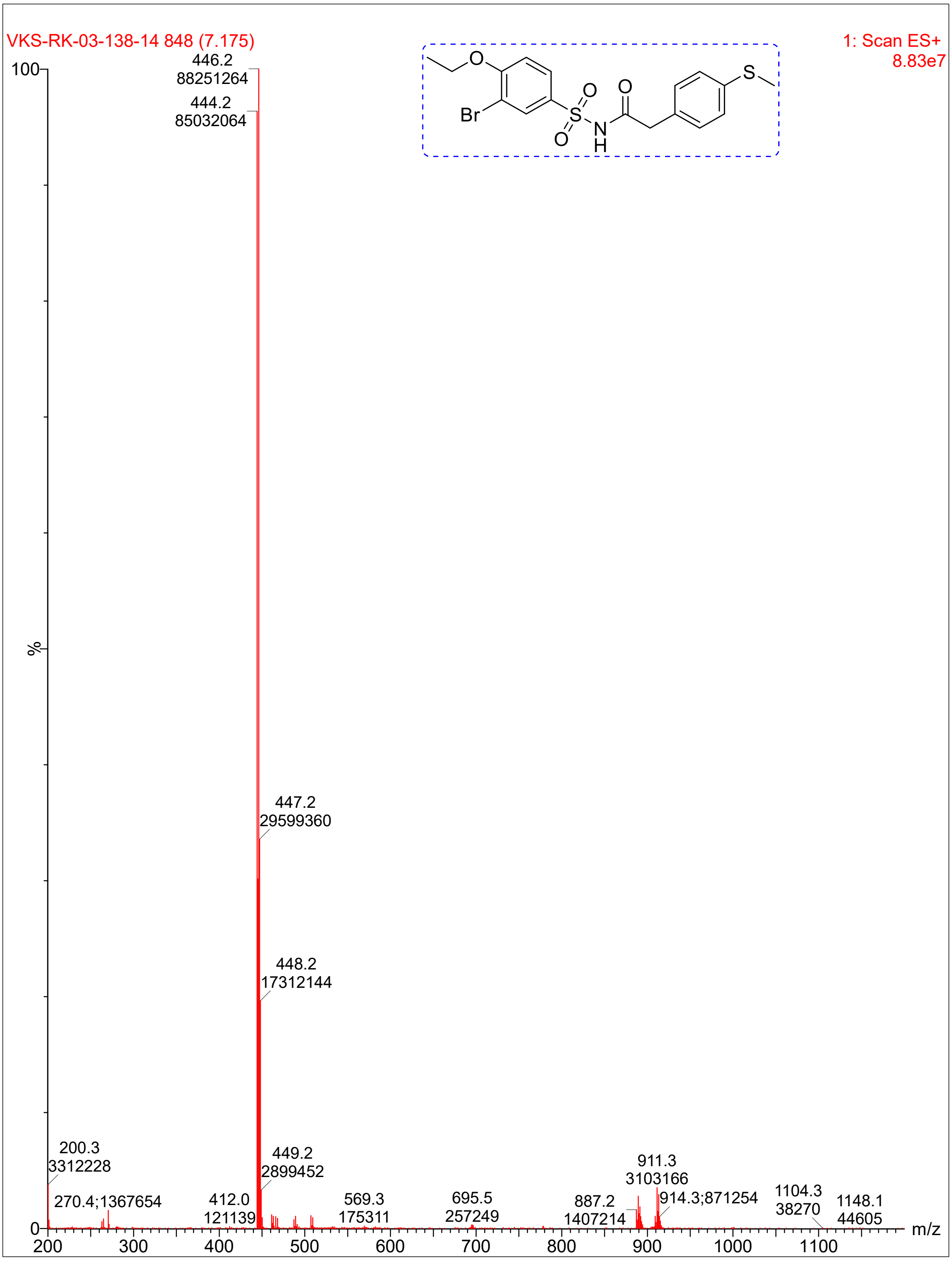

VKS-RK-03-138-14 848 (7.175)
1: Scan ES+
8.83e7
100
%
0
446.2
88251264
444.2
85032064
447.2
29599360
448.2
17312144
449.2
2899452
200.3
3312228
270.4;1367654
412.0
121139
569.3
175311
695.5
257249
887.2
1407214
911.3
3103166
914.3;871254
1104.3
38270
1148.1
44605
200
300
400
500
600
700
800
900
1000
1100
m/z
O
Br
S
O
O
N
H
O
S

**Compound 19:**

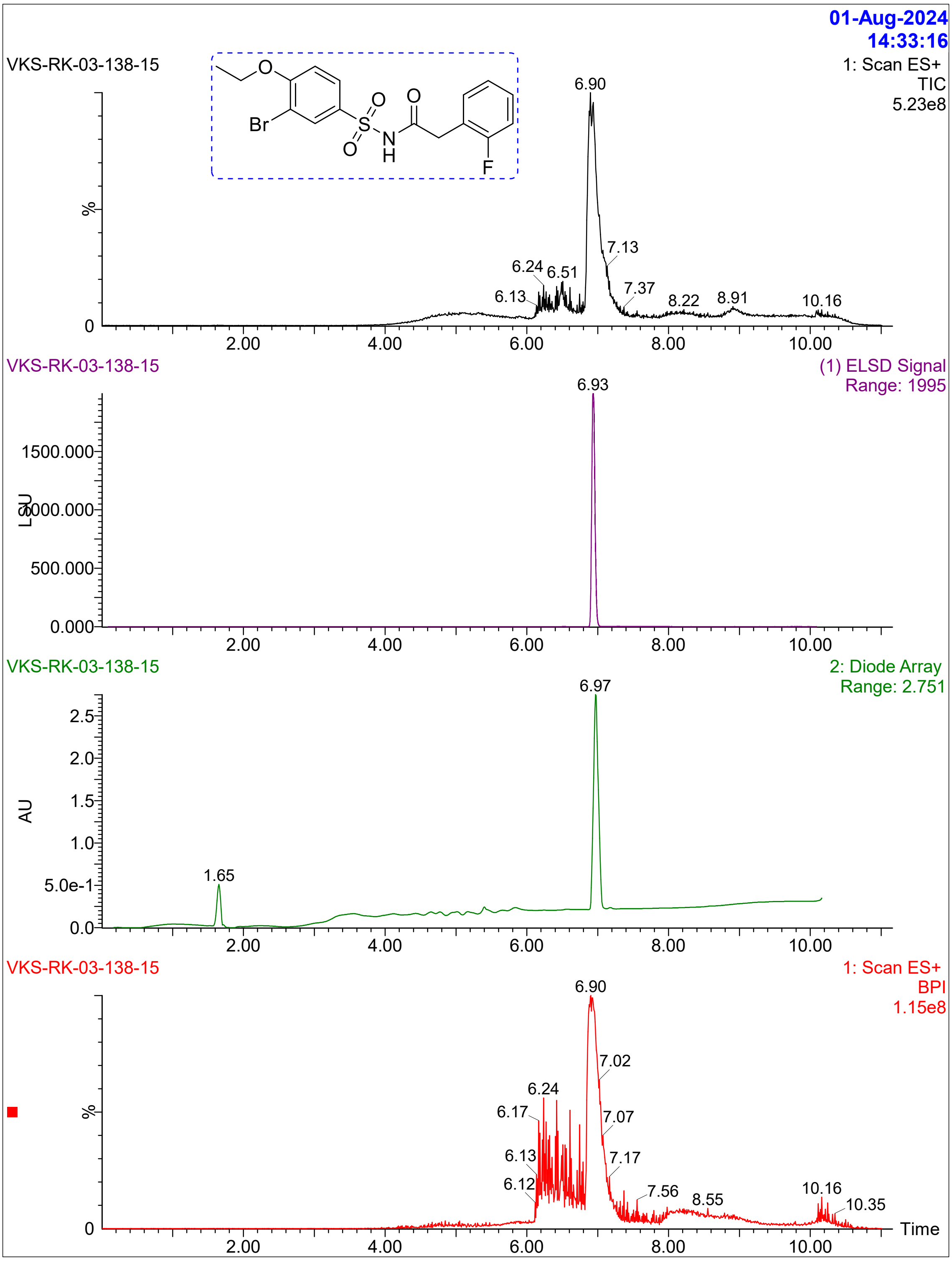

01-Aug-2024
14:33:16
VKS-RK-03-138-15
1: Scan ES+
TIC
5.23e8
O
Br
O
S
O
N
H
O
F
6.90
7.13
6.24
6.51
6.13
7.37
8.22
8.91
10.16
%
0
2.00
4.00
6.00
8.00
10.00
VKS-RK-03-138-15
(1) ELSD Signal
Range: 1995
6.93
1500.000
1000.000
500.000
0.000
LSU
VKS-RK-03-138-15
2: Diode Array
Range: 2.751
6.97
2.5
2.0
1.5
1.0
5.0e-1
0.0
AU
1.65
VKS-RK-03-138-15
1: Scan ES+
BPI
1.15e8
6.90
7.02
6.24
6.17
7.07
6.13
7.17
6.12
7.56
8.55
10.16
10.35
Time

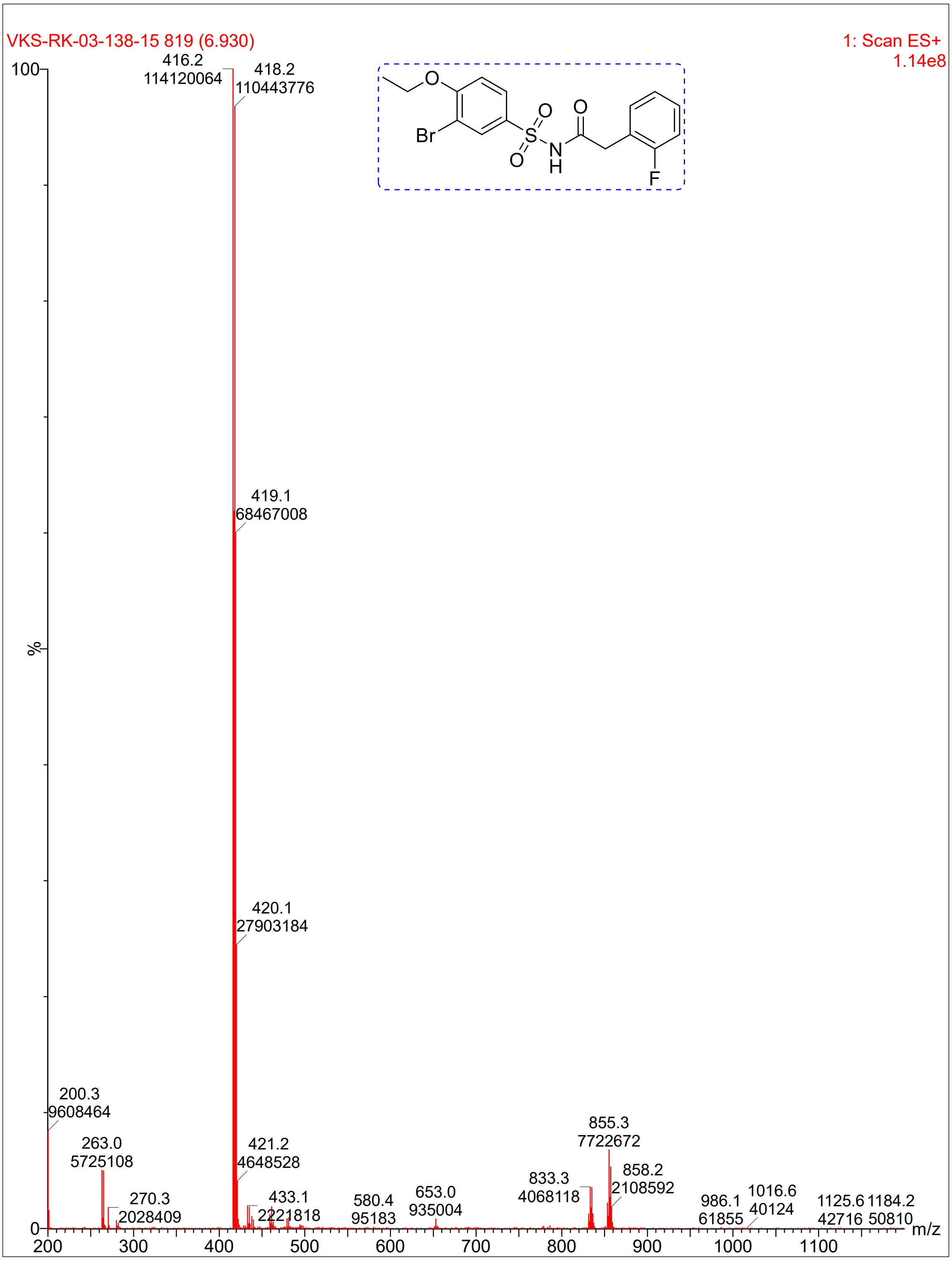
VKS-RK-03-138-15 819 (6.930)
1: Scan ES+
1.14e8
416.2
114120064
418.2
110443776
419.1
68467008
420.1
27903184
421.2
4648528
200.3
9608464
263.0
5725108
270.3
2028409
433.1
2221818
580.4
95183
653.0
935004
833.3
4068118
855.3
7722672
858.2
2108592
986.1
61855
1016.6
40124
1125.6
42716
1184.2
50810
%
m/z
Br
O
S
N
H
F

**Compound 9**:

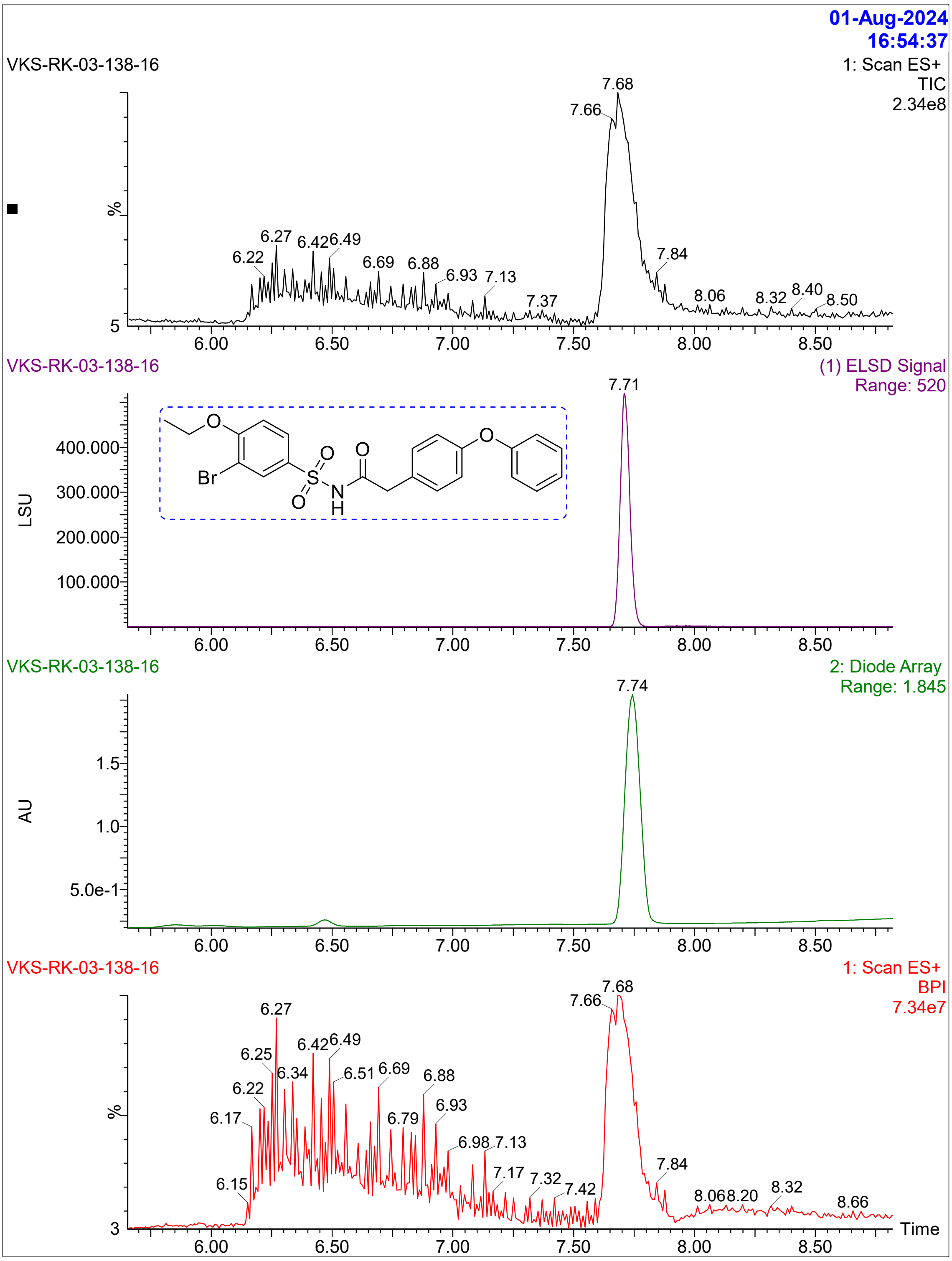
01-Aug-2024
16:54:37
VKS-RK-03-138-16
1: Scan ES+
TIC
2.34e8
%
VKS-RK-03-138-16
(1) ELSD Signal
Range: 520
LSU
VKS-RK-03-138-16
2: Diode Array
Range: 1.845
AU
VKS-RK-03-138-16
1: Scan ES+
BPI
7.34e7
%
Time

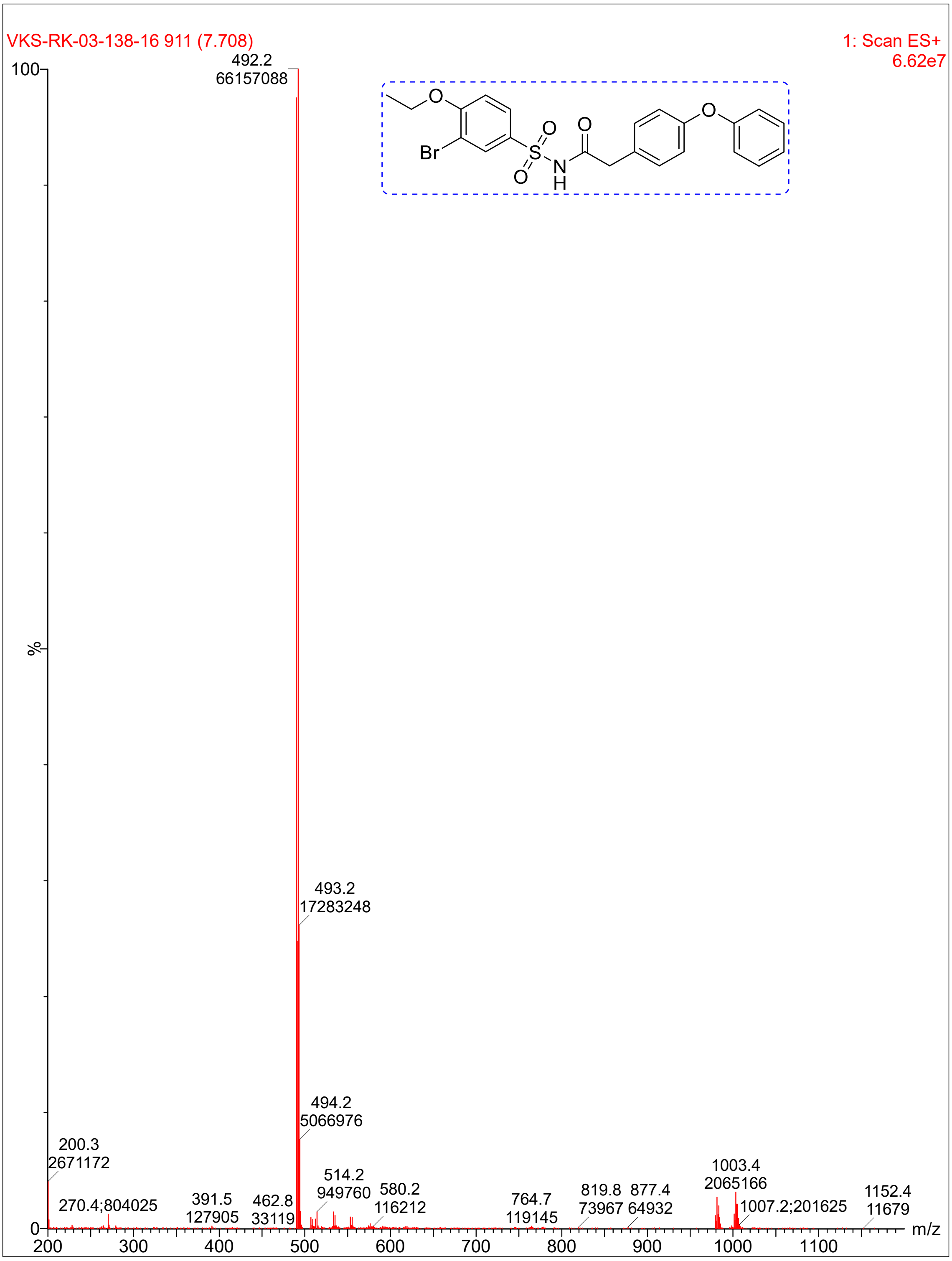
VKS-RK-03-138-16 911 (7.708)
1: Scan ES+
6.62e7
492.2
66157088
493.2
17283248
494.2
5066976
200.3
2671172
270.4;804025
391.5
127905
462.8
33119
514.2
949760
580.2
116212
764.7
119145
819.8
73967
877.4
64932
1003.4
2065166
1007.2;201625
1152.4
11679
Br
O
S
N
H
%
m/z

**Compound 8:**

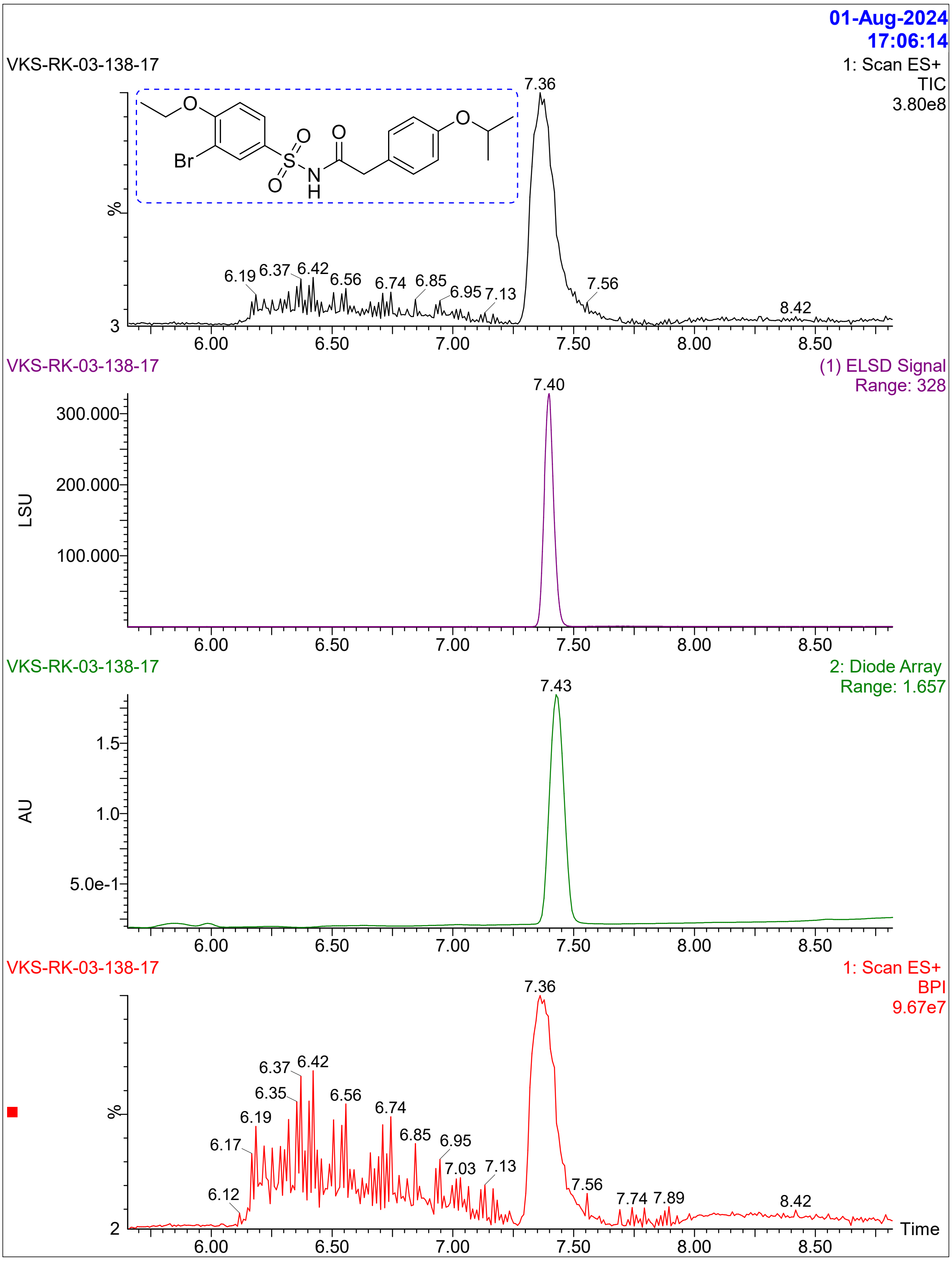

01-Aug-2024
17:06:14
VKS-RK-03-138-17
1: Scan ES+
TIC
3.80e8
Br
O
S
N
H
7.36
6.19 6.37 6.42 6.56 6.74 6.85 6.95 7.13 7.56 8.42
%
3
6.00 6.50 7.00 7.50 8.00 8.50
VKS-RK-03-138-17
(1) ELSD Signal
Range: 328
7.40
300.000
200.000
100.000
LSU
VKS-RK-03-138-17
2: Diode Array
Range: 1.657
7.43
1.5
1.0
5.0e-1
AU
VKS-RK-03-138-17
1: Scan ES+
BPI
9.67e7
7.36
6.12 6.17 6.19 6.35 6.37 6.42 6.56 6.74 6.85 6.95 7.03 7.13 7.56 7.74 7.89 8.42
%
2
Time

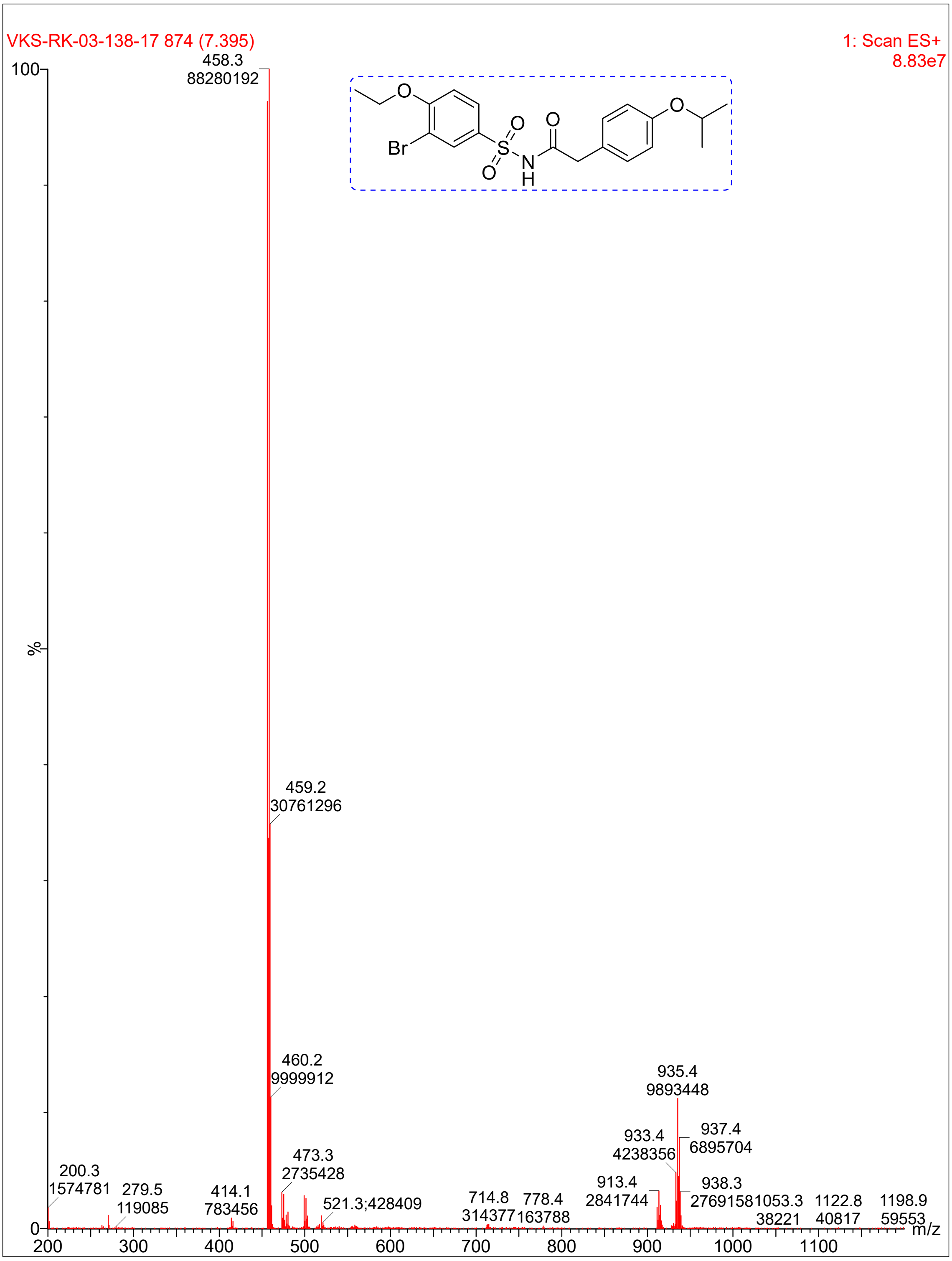
VKS-RK-03-138-17 874 (7.395)
1: Scan ES+
8.83e7
458.3
88280192
459.2
30761296
460.2
9999912
473.3
2735428
521.3;428409
200.3
1574781
279.5
119085
414.1
783456
714.8
314377
778.4
163788
913.4
2841744
933.4
4238356
935.4
9893448
937.4
6895704
938.3
2769158
1053.3
38221
1122.8
40817
1198.9
59553
%
m/z
Br
O
S
N
H

**Compound 42:**

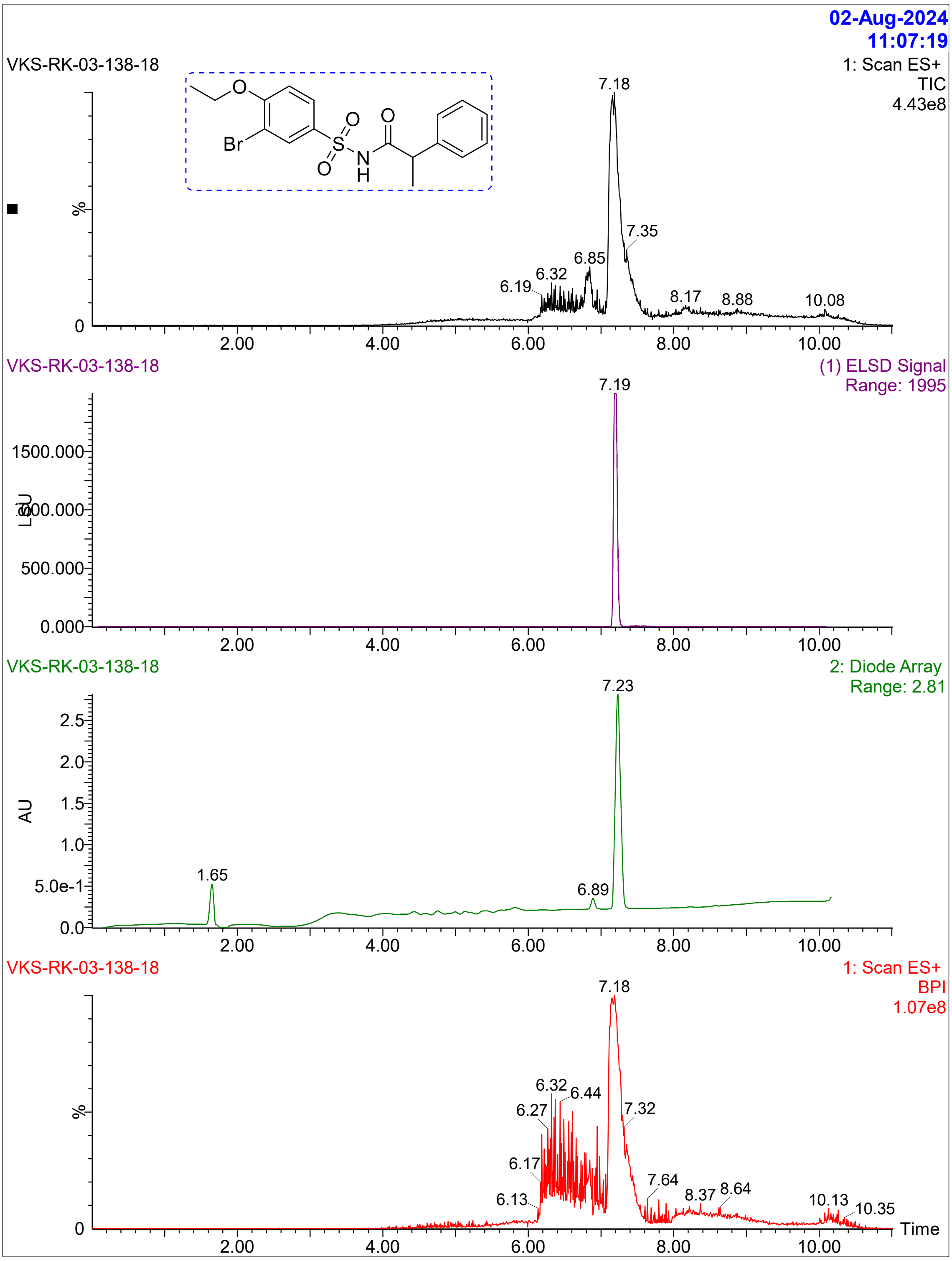

02-Aug-2024
11:07:19
VKS-RK-03-138-18
1: Scan ES+
TIC
4.43e8
Br
O
S
N
H
%
0
7.18
7.35
6.85
6.32
6.19
8.17
8.88
10.08
2.00
4.00
6.00
8.00
10.00
VKS-RK-03-138-18
(1) ELSD Signal
Range: 1995
7.19
1500.000
1000.000
500.000
0.000
LSU
VKS-RK-03-138-18
2: Diode Array
Range: 2.81
7.23
2.5
2.0
1.5
1.0
5.0e-1
0.0
AU
1.65
6.89
VKS-RK-03-138-18
1: Scan ES+
BPI
1.07e8
7.18
6.32
6.44
6.27
7.32
6.17
6.13
7.64
8.37
8.64
10.13
10.35
Time

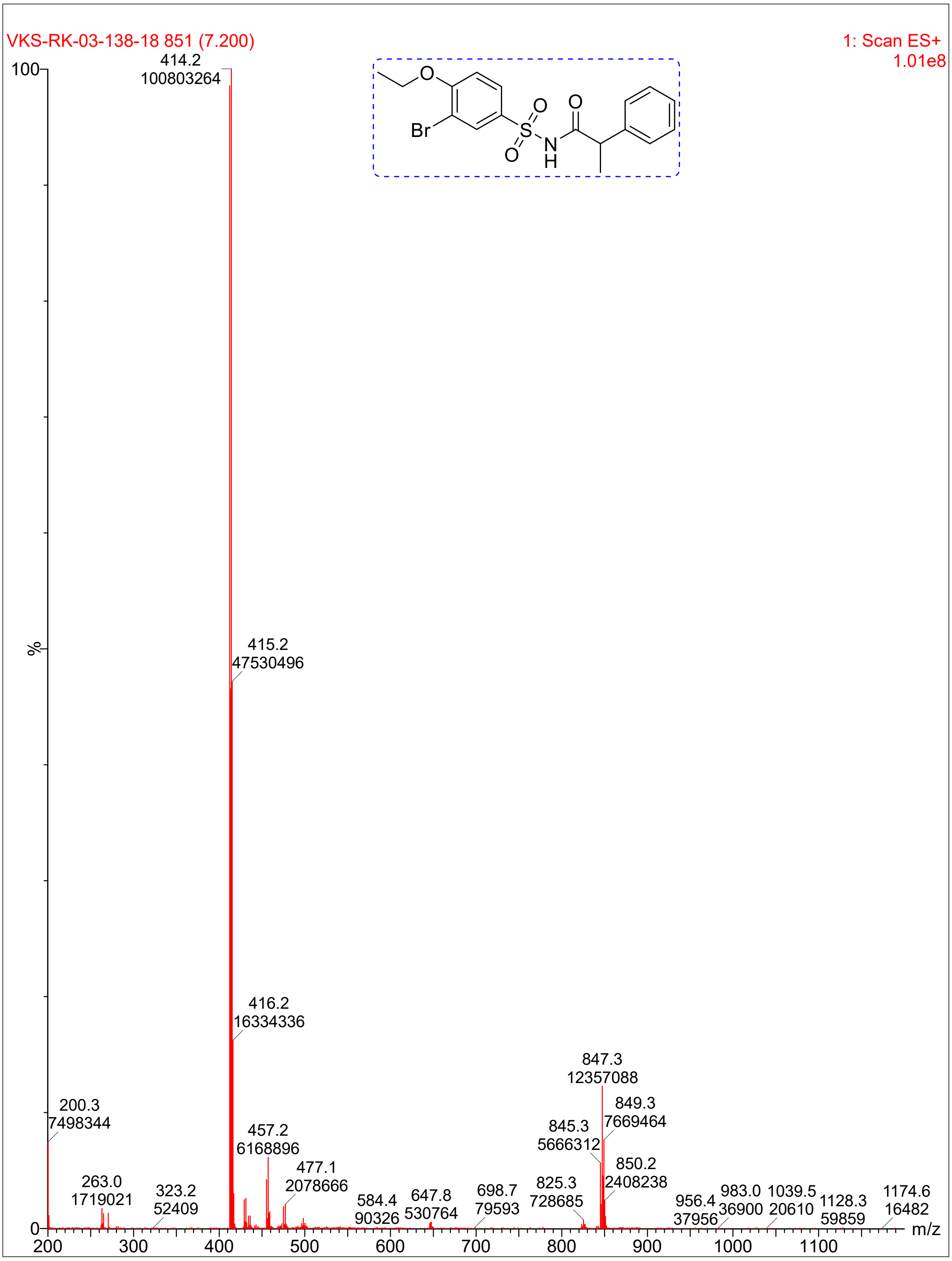

VKS-RK-03-138-18 851 (7.200)
1: Scan ES+
1.01e8
100
%
0
414.2
100803264
415.2
47530496
416.2
16334336
200.3
7498344
263.0
1719021
323.2
52409
457.2
6168896
477.1
2078666
584.4
90326
647.8
530764
698.7
79593
825.3
728685
845.3
5666312
847.3
12357088
849.3
7669464
850.2
2408238
956.4
37956
983.0
36900
1039.5
20610
1128.3
59859
1174.6
16482
m/z
200
300
400
500
600
700
800
900
1000
1100
Br
O
S
N
H

**Compound 34:**

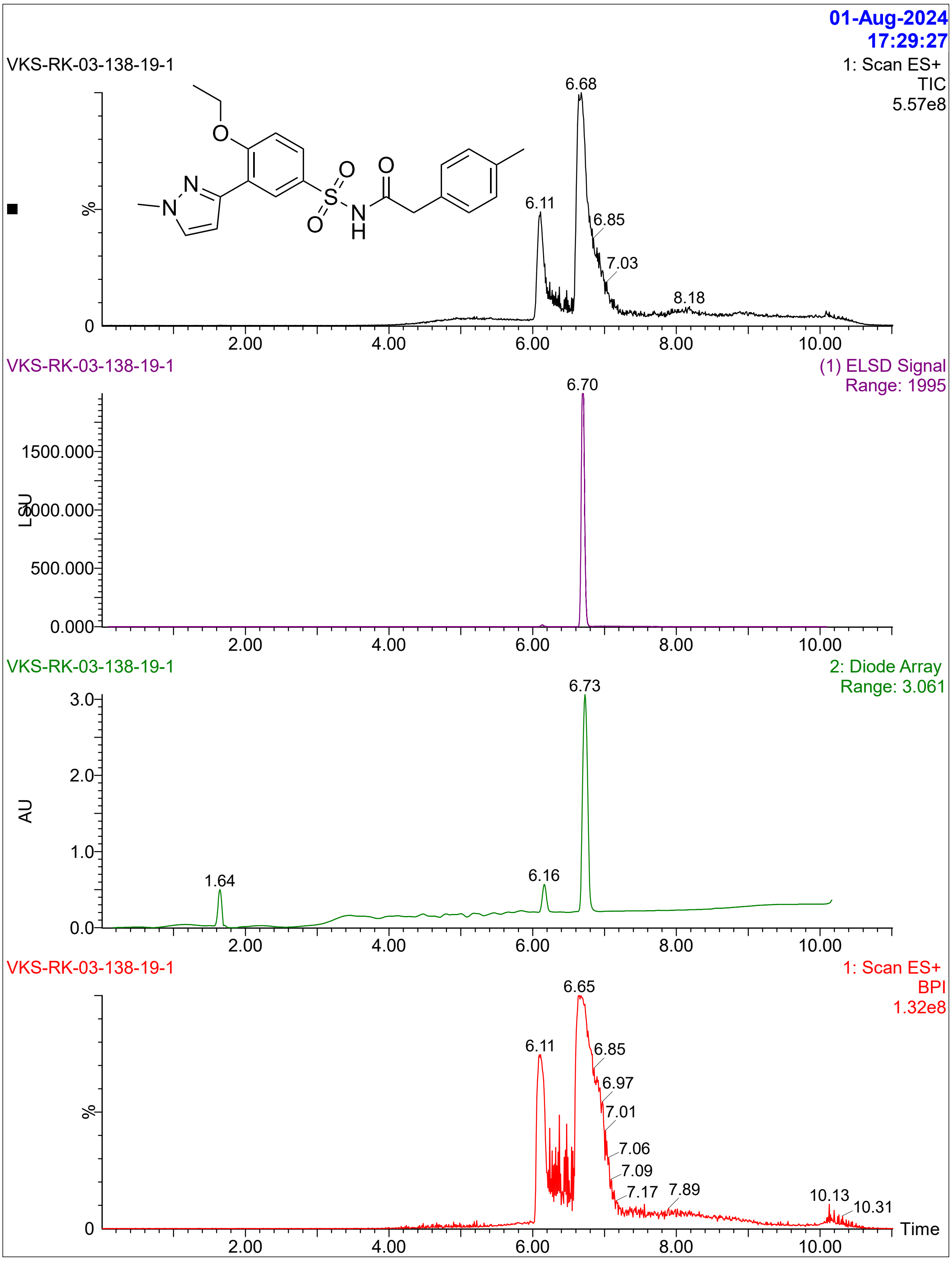

01-Aug-2024
17:29:27
VKS-RK-03-138-19-1
1: Scan ES+
TIC
5.57e8
%
6.68
6.11
6.85
7.03
8.18
2.00
4.00
6.00
8.00
10.00
VKS-RK-03-138-19-1
(1) ELSD Signal
Range: 1995
6.70
LSU
1500.000
1000.000
500.000
0.000
VKS-RK-03-138-19-1
2: Diode Array
Range: 3.061
6.73
AU
3.0
2.0
1.0
0.0
1.64
6.16
VKS-RK-03-138-19-1
1: Scan ES+
BPI
1.32e8
6.65
6.11
6.85
6.97
7.01
7.06
7.09
7.17
7.89
10.13
10.31
Time

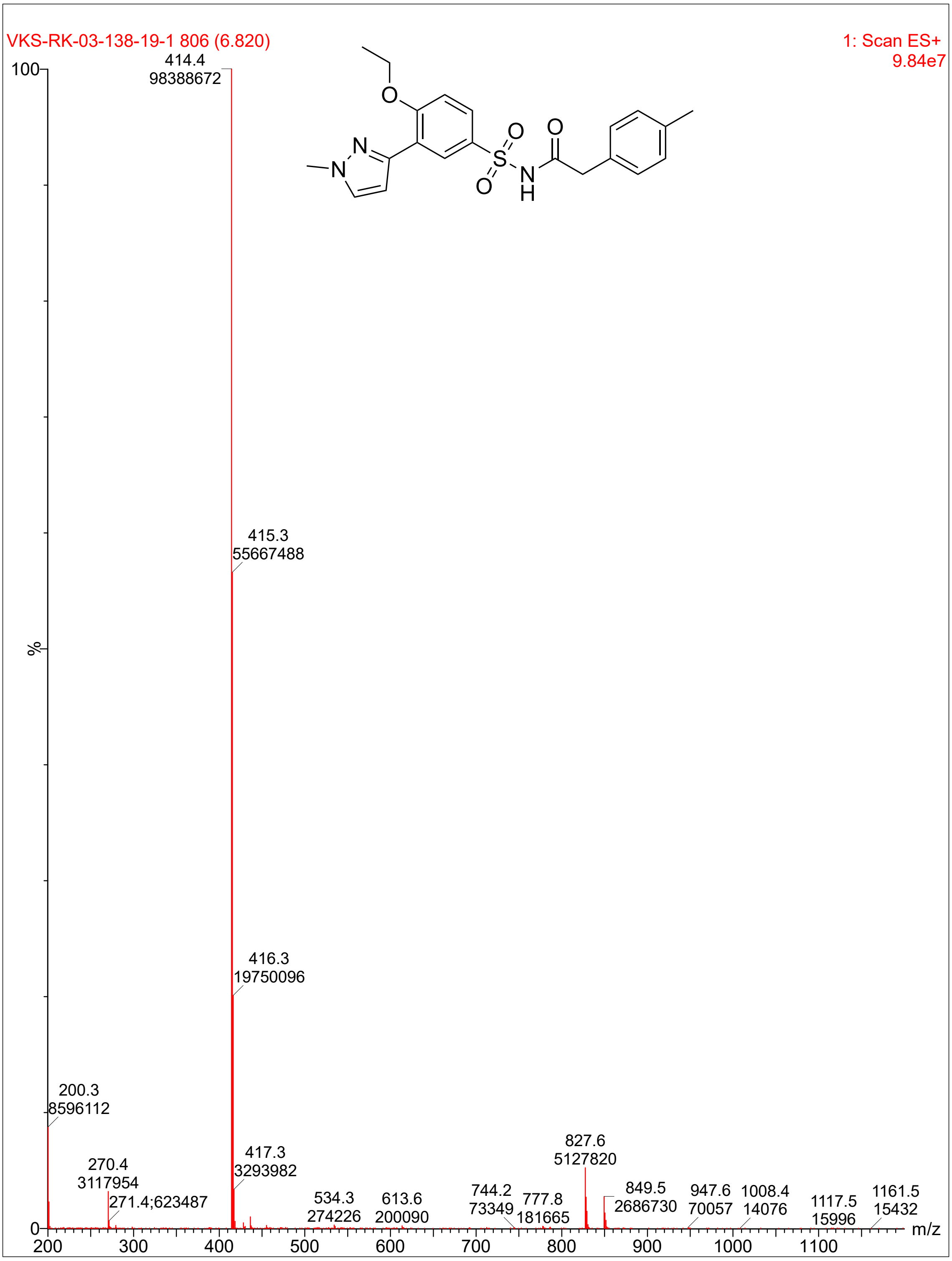

VKS-RK-03-138-19-1 806 (6.820)
1: Scan ES+
9.84e7
100
%
0
414.4
98388672
415.3
55667488
416.3
19750096
200.3
8596112
270.4
3117954
271.4;623487
417.3
3293982
534.3
274226
613.6
200090
744.2
73349
777.8
181665
827.6
5127820
849.5
2686730
947.6
70057
1008.4
14076
1117.5
15996
1161.5
15432
200
300
400
500
600
700
800
900
1000
1100
m/z
O
N
N
S
O
O
N
H
O

**Compound 12:**

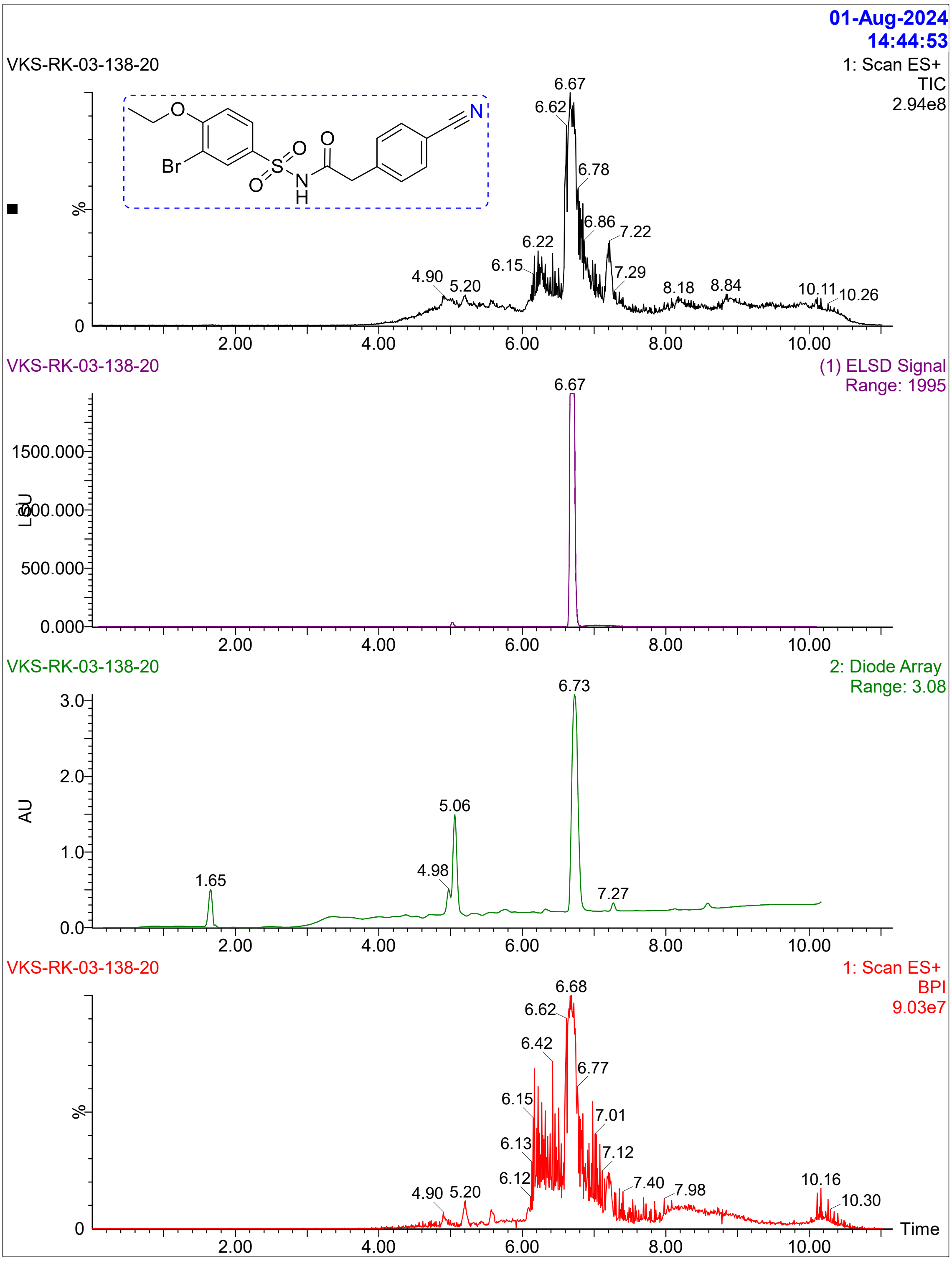

01-Aug-2024
14:44:53
VKS-RK-03-138-20
1: Scan ES+
TIC
2.94e8
6.67
6.62
6.78
6.86
7.22
6.22
6.15
7.29
4.90
5.20
8.18
8.84
10.11
10.26
%
0
2.00
4.00
6.00
8.00
10.00
VKS-RK-03-138-20
(1) ELSD Signal
Range: 1995
6.67
1500.000
1000.000
500.000
0.000
LSU
VKS-RK-03-138-20
2: Diode Array
Range: 3.08
6.73
3.0
2.0
1.0
0.0
AU
5.06
4.98
1.65
7.27
VKS-RK-03-138-20
1: Scan ES+
BPI
9.03e7
6.68
6.62
6.42
6.77
6.15
7.01
6.13
7.12
6.12
7.40
7.98
4.90
5.20
10.16
10.30
Time

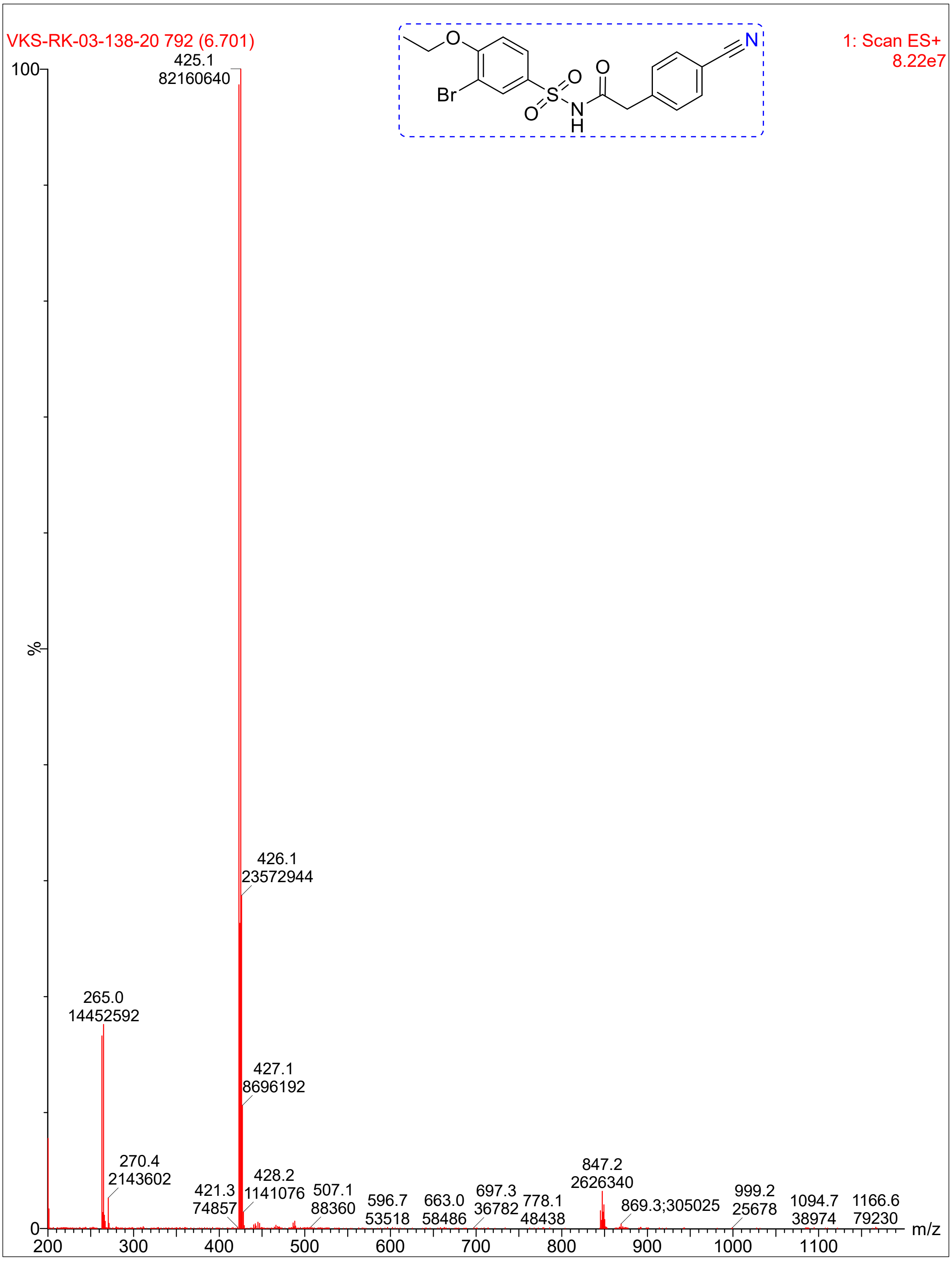
VKS-RK-03-138-20 792 (6.701)
1: Scan ES+
8.22e7
Br
O
S
N
H
N
100
%
0
425.1
82160640
426.1
23572944
265.0
14452592
427.1
8696192
270.4
2143602
421.3
74857
428.2
1141076
507.1
88360
596.7
53518
663.0
58486
697.3
36782
778.1
48438
847.2
2626340
869.3;305025
999.2
25678
1094.7
38974
1166.6
79230
200
300
400
500
600
700
800
900
1000
1100
m/z

**Compound 24:**

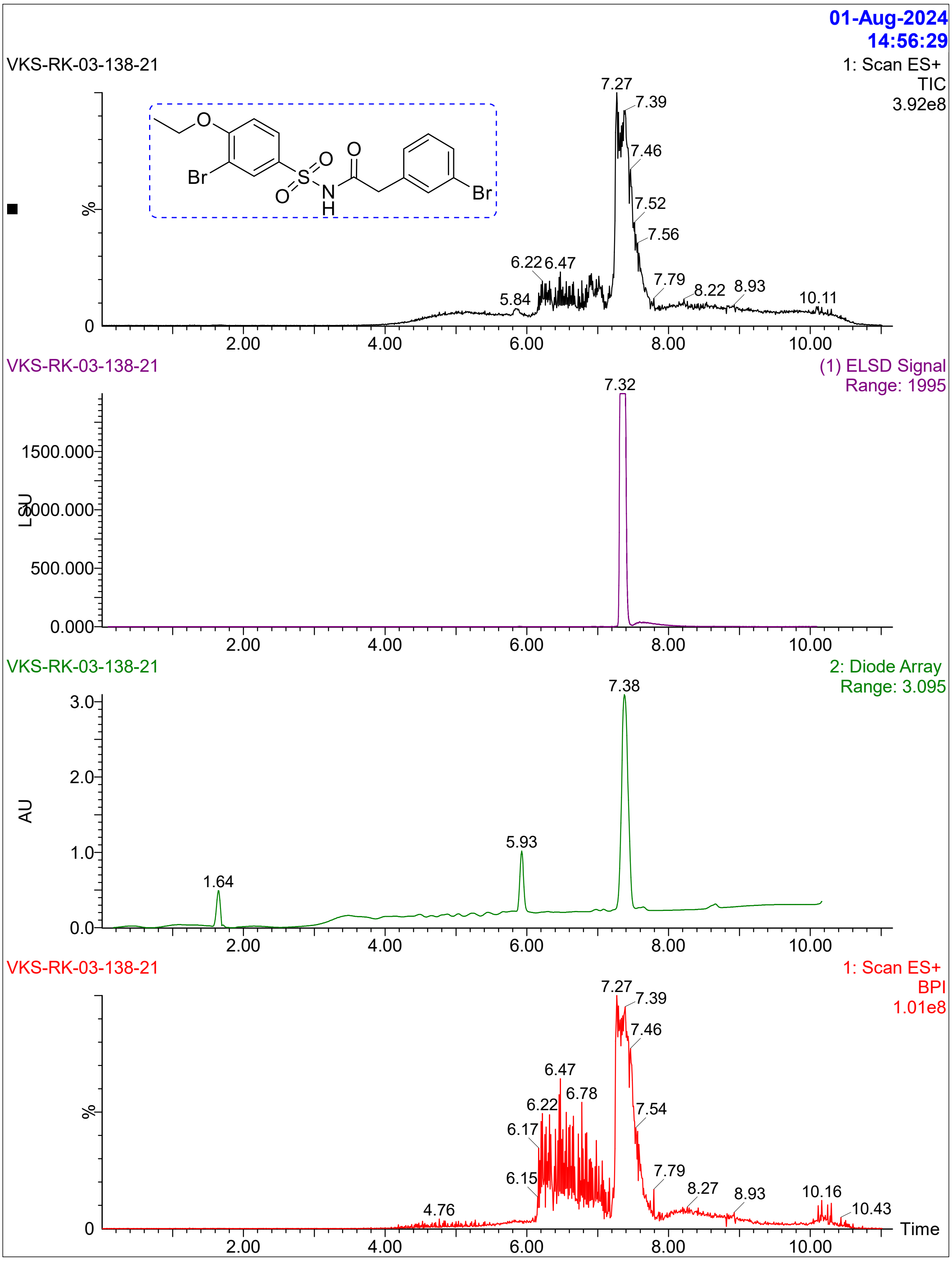
01-Aug-2024
14:56:29
VKS-RK-03-138-21
1: Scan ES+
TIC
3.92e8
Br
O
S
O
O
N
H
O
Br
%
7.27
7.39
7.46
7.52
7.56
6.22
6.47
5.84
7.79
8.22
8.93
10.11
VKS-RK-03-138-21
(1) ELSD Signal
Range: 1995
7.32
LSU
1500.000
1000.000
500.000
0.000
VKS-RK-03-138-21
2: Diode Array
Range: 3.095
7.38
AU
3.0
2.0
1.0
0.0
5.93
1.64
VKS-RK-03-138-21
1: Scan ES+
BPI
1.01e8
7.27
7.39
7.46
6.47
6.78
6.22
7.54
6.17
6.15
7.79
8.27
8.93
10.16
10.43
4.76
Time
2.00
4.00
6.00
8.00
10.00

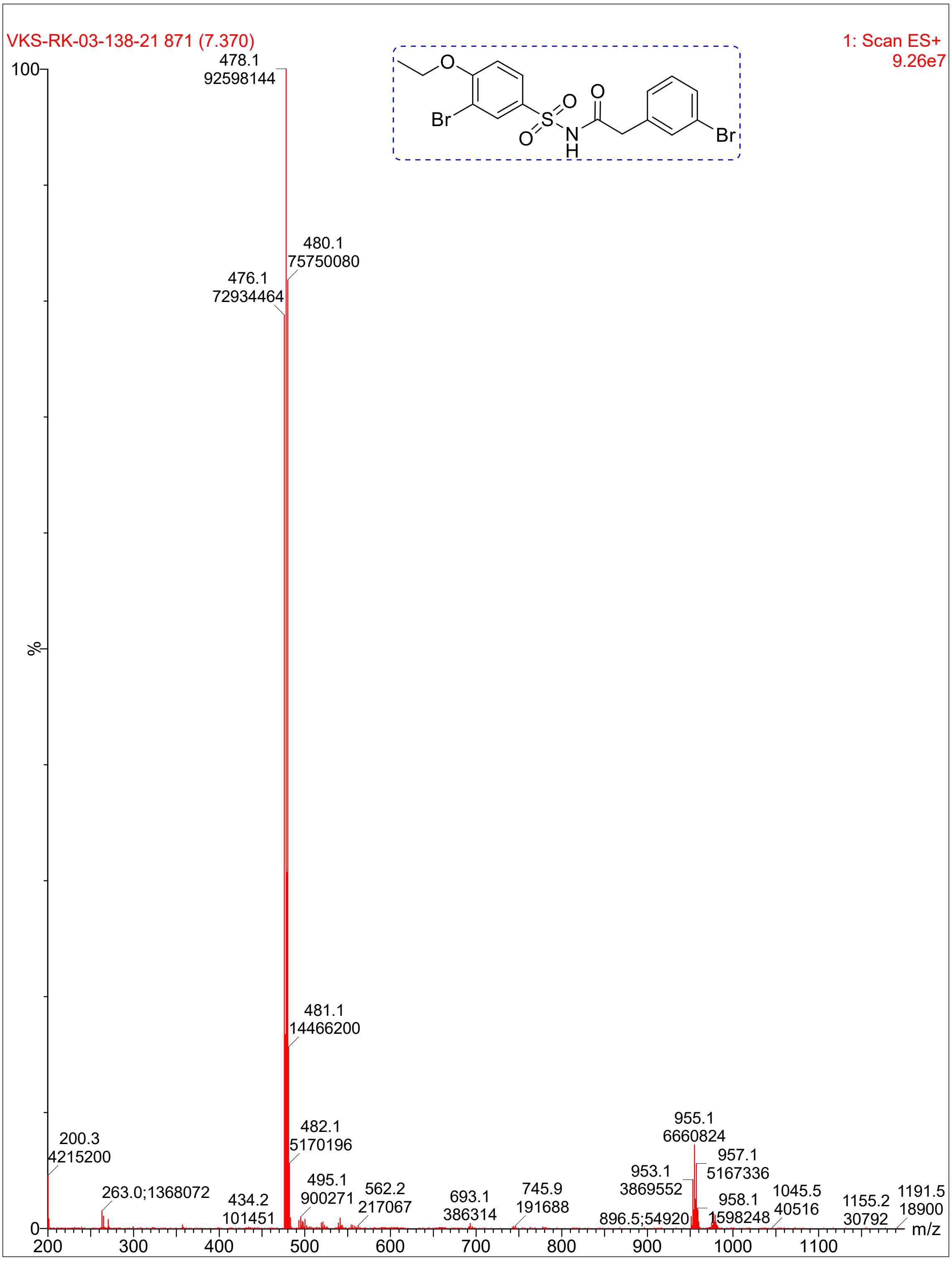

VKS-RK-03-138-21 871 (7.370)
1: Scan ES+
9.26e7
478.1
92598144
480.1
75750080
476.1
72934464
481.1
14466200
482.1
5170196
200.3
4215200
263.0;1368072
434.2
101451
495.1
900271
562.2
217067
693.1
386314
745.9
191688
896.5;54920
953.1
3869552
955.1
6660824
957.1
5167336
958.1
1598248
1045.5
40516
1155.2
30792
1191.5
18900
%
m/z
Br
O
S
N
H

### Compound 1:

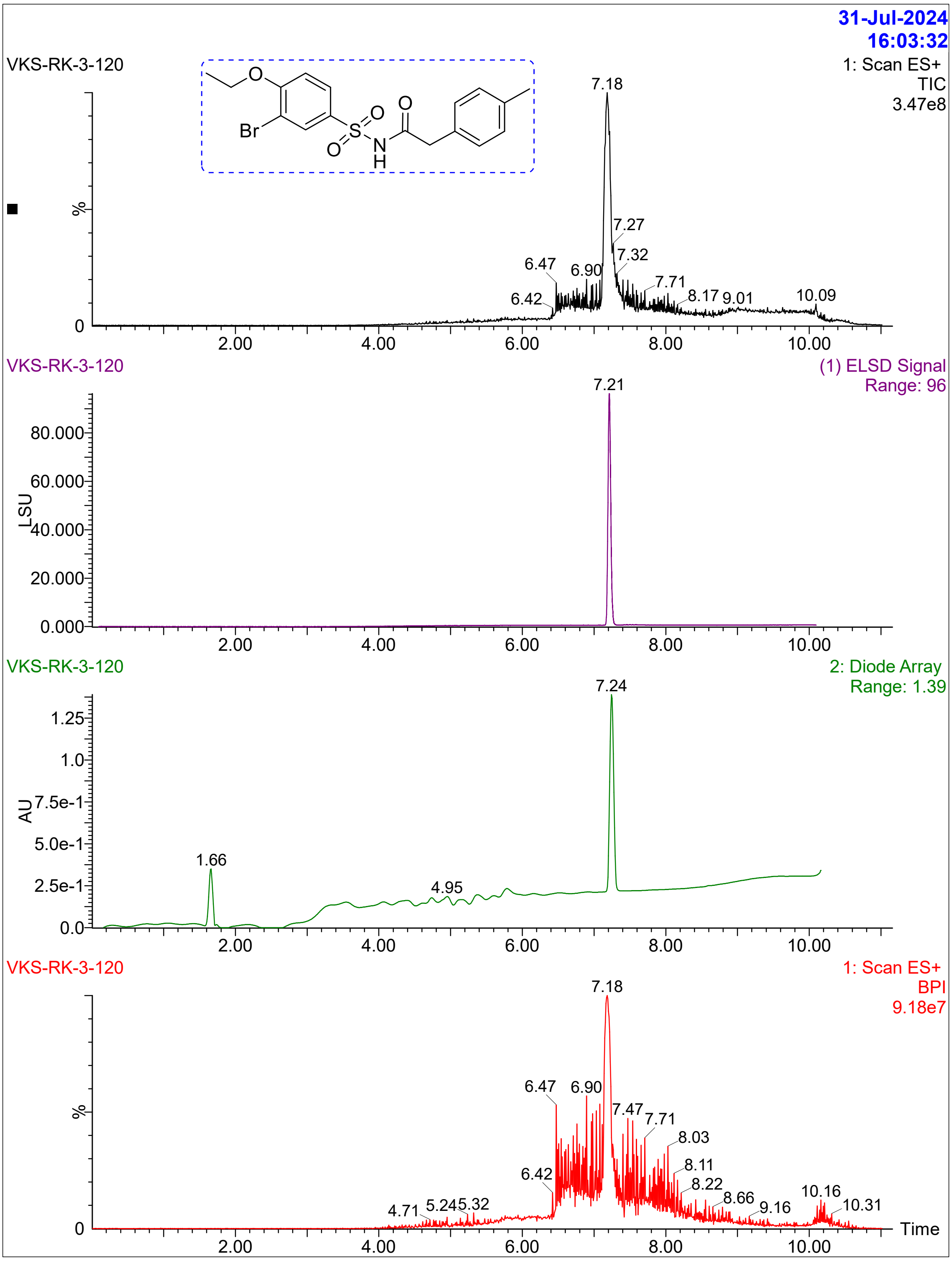
31-Jul-2024
16:03:32
VKS-RK-3-120
1: Scan ES+
TIC
3.47e8
%
7.18
7.27
7.32
6.47
6.90
6.42
7.71
8.17
9.01
10.09
2.00
4.00
6.00
8.00
10.00
VKS-RK-3-120
(1) ELSD Signal
Range: 96
LSU
7.21
80.000
60.000
40.000
20.000
0.000
VKS-RK-3-120
2: Diode Array
Range: 1.39
AU
7.24
1.66
4.95
1.25
1.0
7.5e-1
5.0e-1
2.5e-1
0.0
VKS-RK-3-120
1: Scan ES+
BPI
9.18e7
7.18
6.47
6.90
7.47
7.71
8.03
6.42
8.11
8.22
8.66
9.16
10.16
10.31
4.71
5.24
5.32
Time

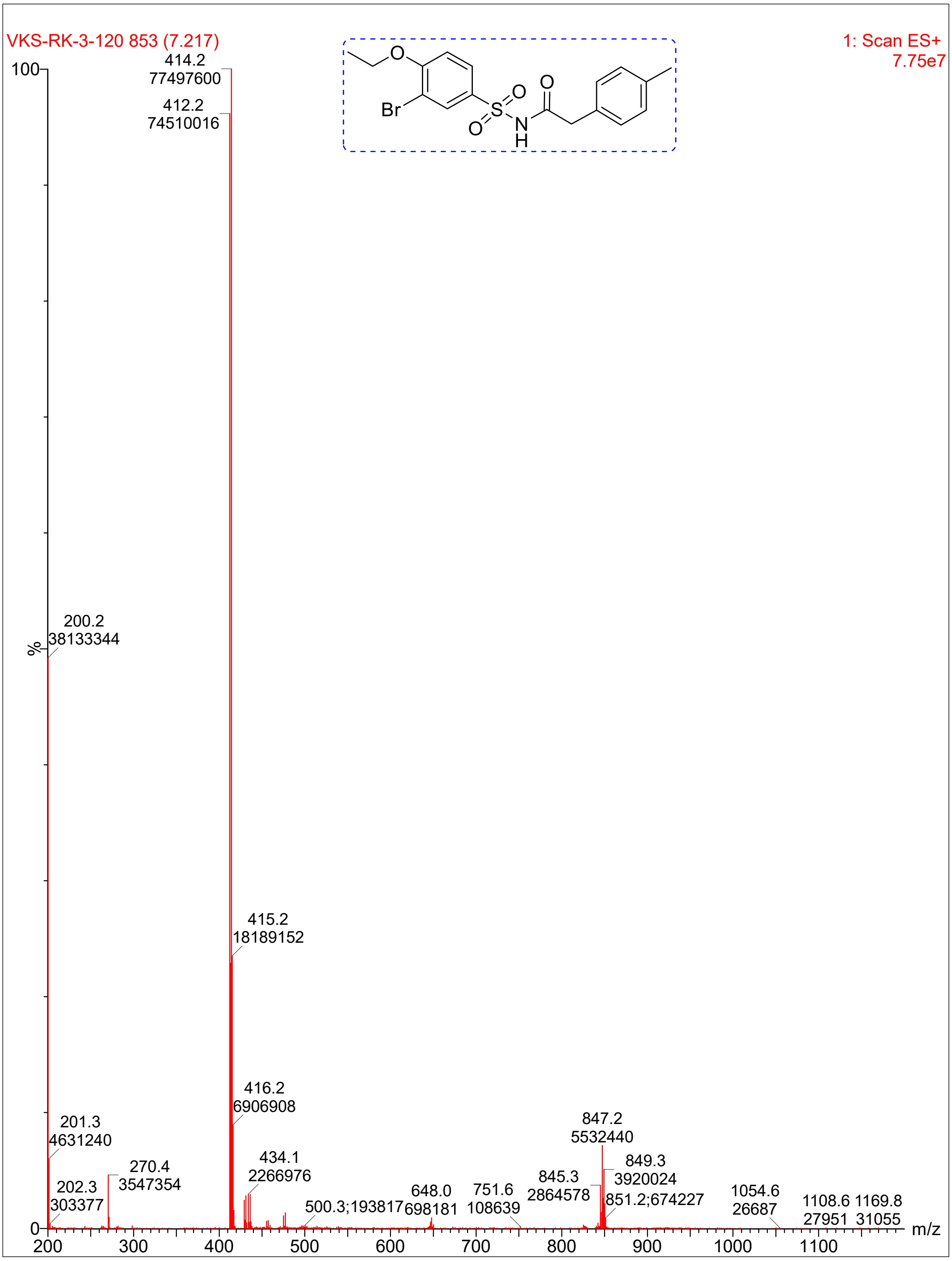

VKS-RK-3-120 853 (7.217)
1: Scan ES+
7.75e7
Br
O
S
N
H
414.2
77497600
412.2
74510016
200.2
38133344
415.2
18189152
416.2
6906908
201.3
4631240
270.4
3547354
202.3
303377
434.1
2266976
500.3;193817
648.0
698181
751.6
108639
845.3
2864578
847.2
5532440
849.3
3920024
851.2;674227
1054.6
26687
1108.6
27951
1169.8
31055
%
m/z

**Compound 3:**

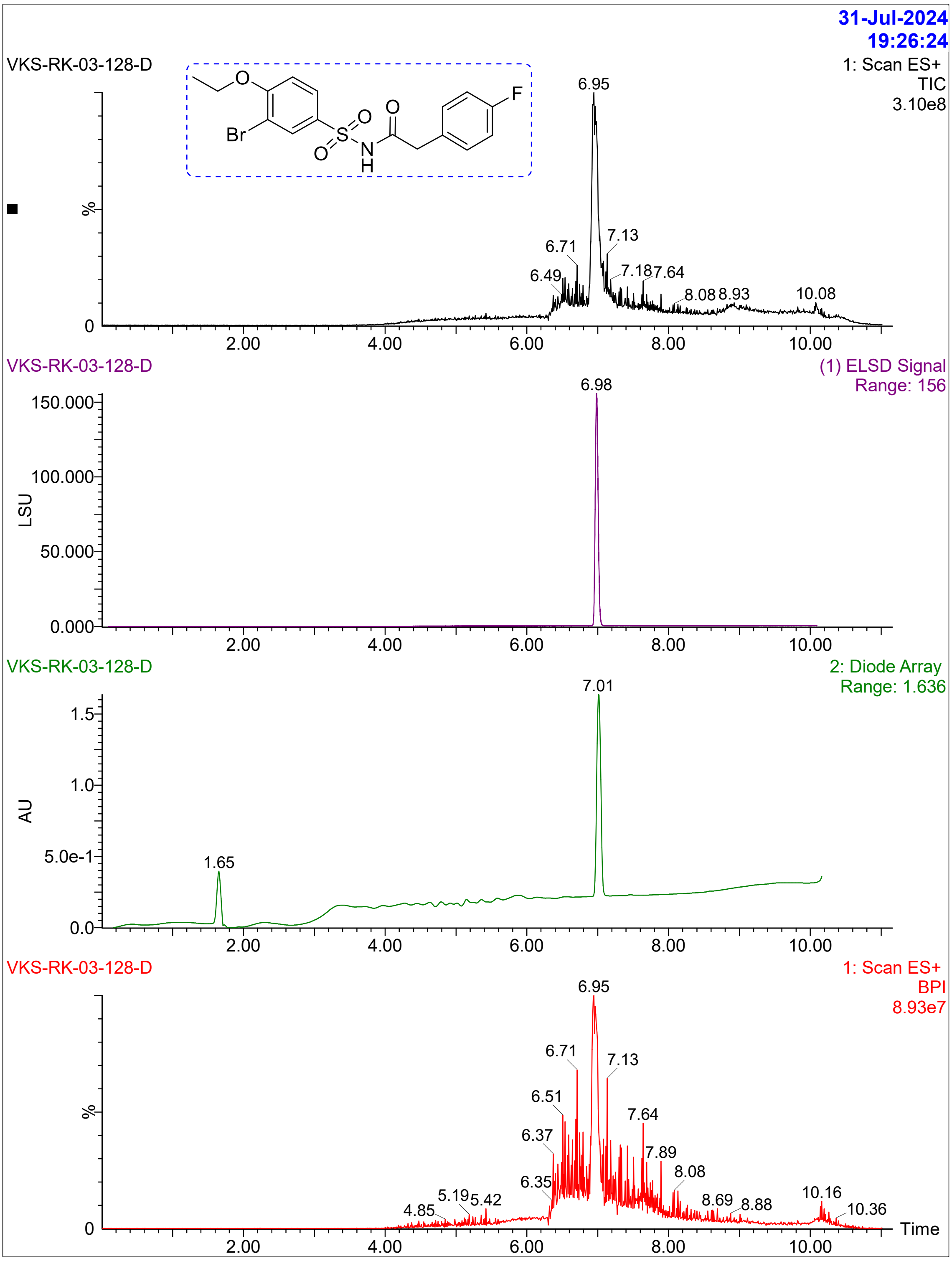
31-Jul-2024
19:26:24
VKS-RK-03-128-D
1: Scan ES+
TIC
3.10e8
6.95
6.71
7.13
6.49
7.18
7.64
8.08
8.93
10.08
%
0
2.00
4.00
6.00
8.00
10.00
VKS-RK-03-128-D
(1) ELSD Signal
Range: 156
6.98
150.000
100.000
50.000
0.000
LSU
VKS-RK-03-128-D
2: Diode Array
Range: 1.636
7.01
1.65
1.5
1.0
5.0e-1
0.0
AU
VKS-RK-03-128-D
1: Scan ES+
BPI
8.93e7
6.95
6.71
7.13
6.51
7.64
6.37
7.89
8.08
6.35
5.19
5.42
4.85
8.69
8.88
10.16
10.36
Time

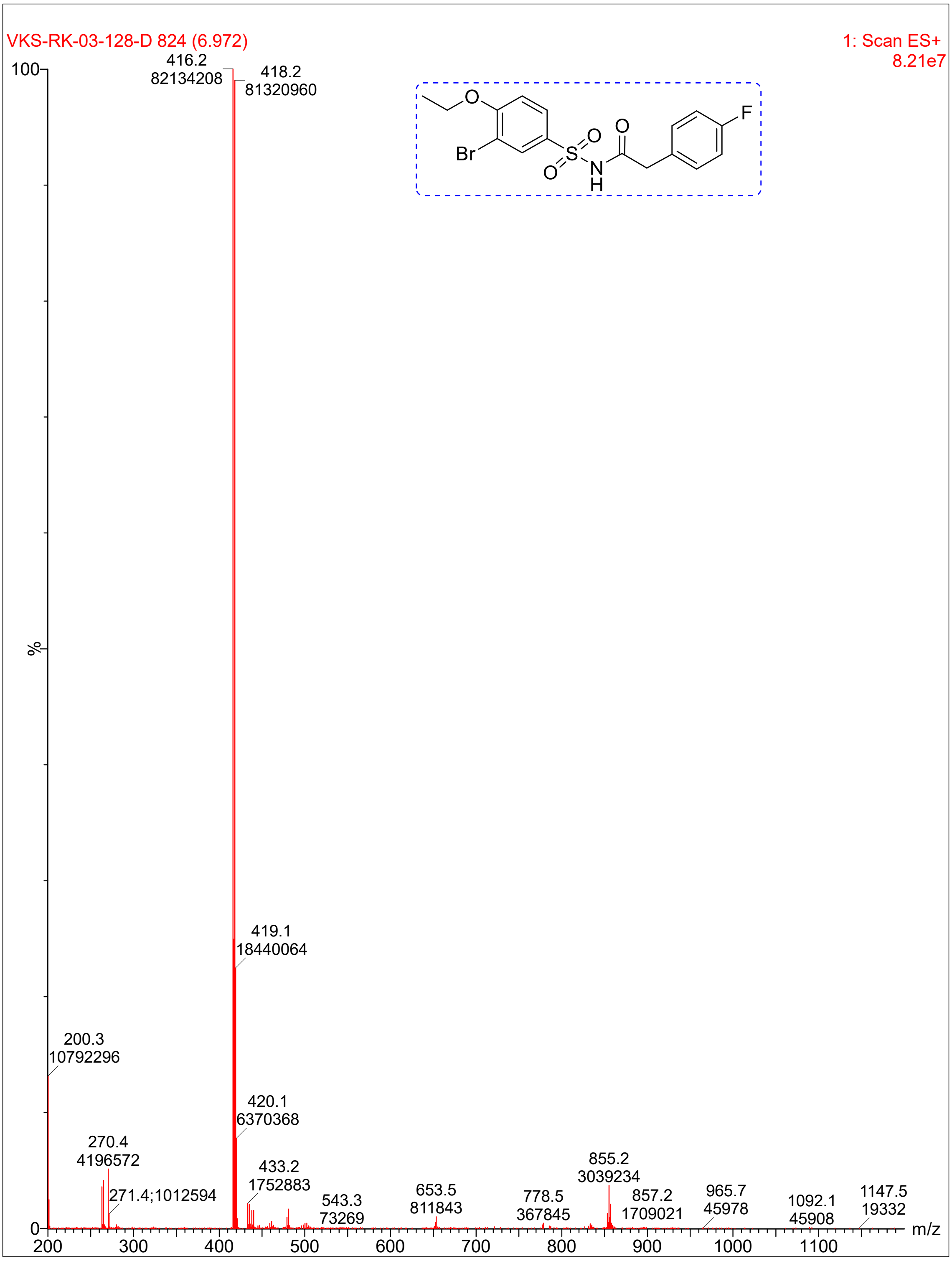
VKS-RK-03-128-D 824 (6.972)
1: Scan ES+
8.21e7
416.2
82134208
418.2
81320960
419.1
18440064
200.3
10792296
420.1
6370368
270.4
4196572
271.4;1012594
433.2
1752883
543.3
73269
653.5
811843
778.5
367845
855.2
3039234
857.2
1709021
965.7
45978
1092.1
45908
1147.5
19332
%
100
0
200
300
400
500
600
700
800
900
1000
1100
m/z
O
Br
S
O
O
N
H
O
F

**Compound 5:**

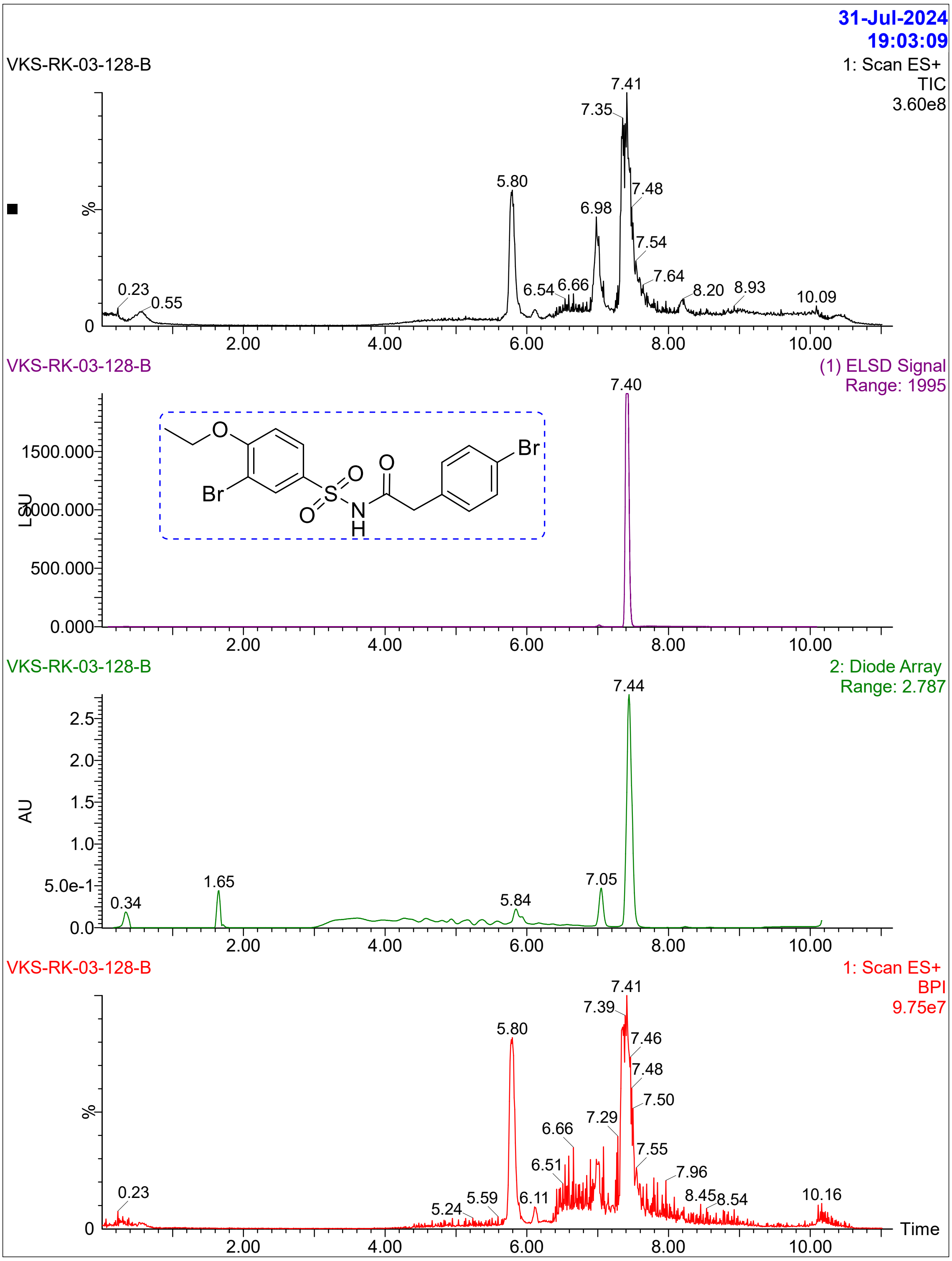
31-Jul-2024
19:03:09
VKS-RK-03-128-B
1: Scan ES+
TIC
3.60e8
7.41
7.35
5.80
7.48
6.98
7.54
7.64
0.23
0.55
6.54
6.66
8.20
8.93
10.09
%
0
2.00
4.00
6.00
8.00
10.00
VKS-RK-03-128-B
(1) ELSD Signal
Range: 1995
7.40
1500.000
1000.000
500.000
0.000
LSU
O
Br
S
N
H
Br
VKS-RK-03-128-B
2: Diode Array
Range: 2.787
7.44
2.5
2.0
1.5
1.0
5.0e-1
0.0
AU
0.34
1.65
5.84
7.05
VKS-RK-03-128-B
1: Scan ES+
BPI
9.75e7
7.41
7.39
5.80
7.46
7.48
7.50
6.66
7.29
6.51
7.55
7.96
0.23
5.24
5.59
6.11
8.45
8.54
10.16
Time

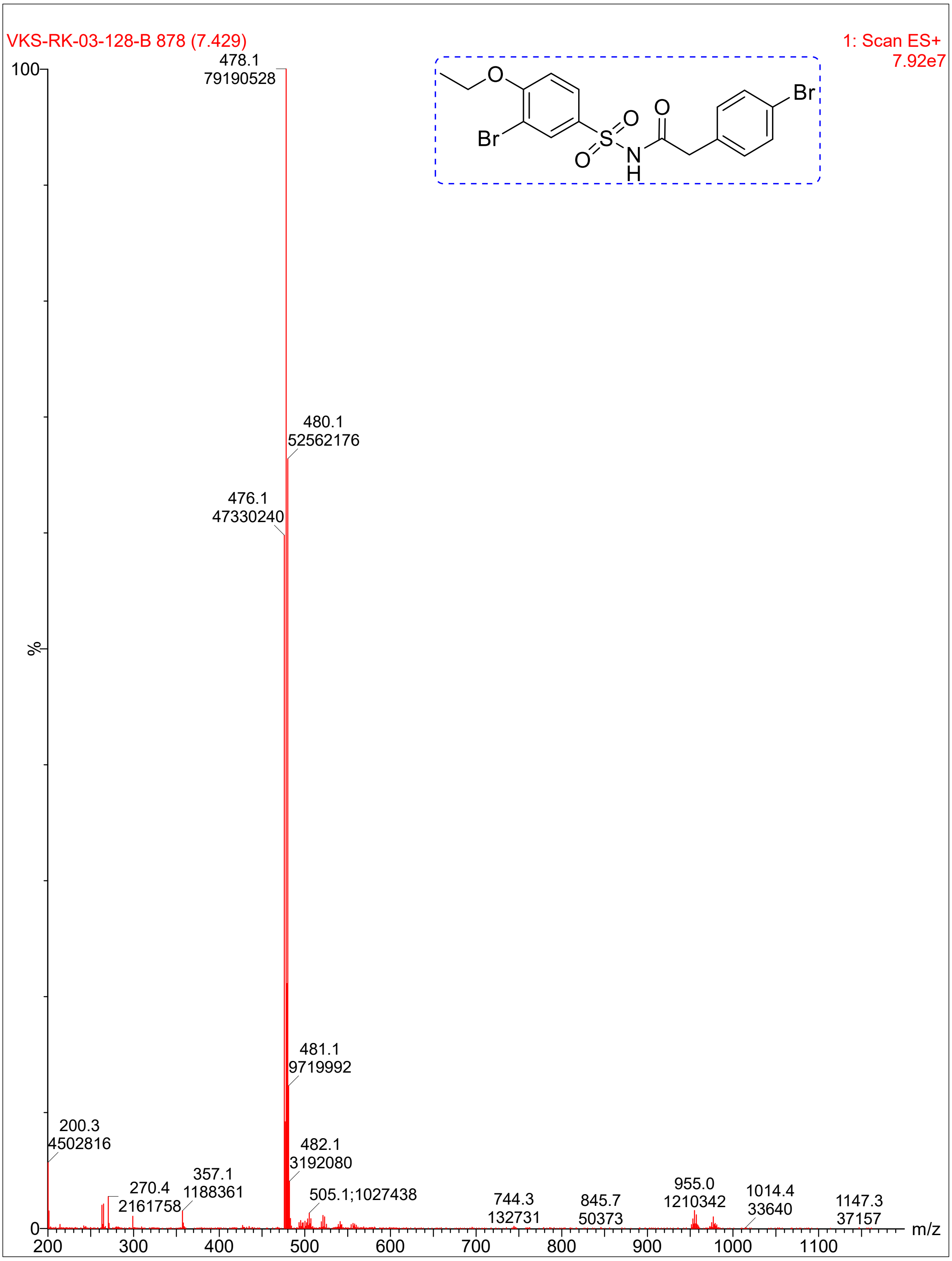
VKS-RK-03-128-B 878 (7.429)
1: Scan ES+
7.92e7
478.1
79190528
480.1
52562176
476.1
47330240
481.1
9719992
482.1
3192080
200.3
4502816
270.4
2161758
357.1
1188361
505.1;1027438
744.3
132731
845.7
50373
955.0
1210342
1014.4
33640
1147.3
37157
%
m/z
Br
O
S
N
H

### Compound 22

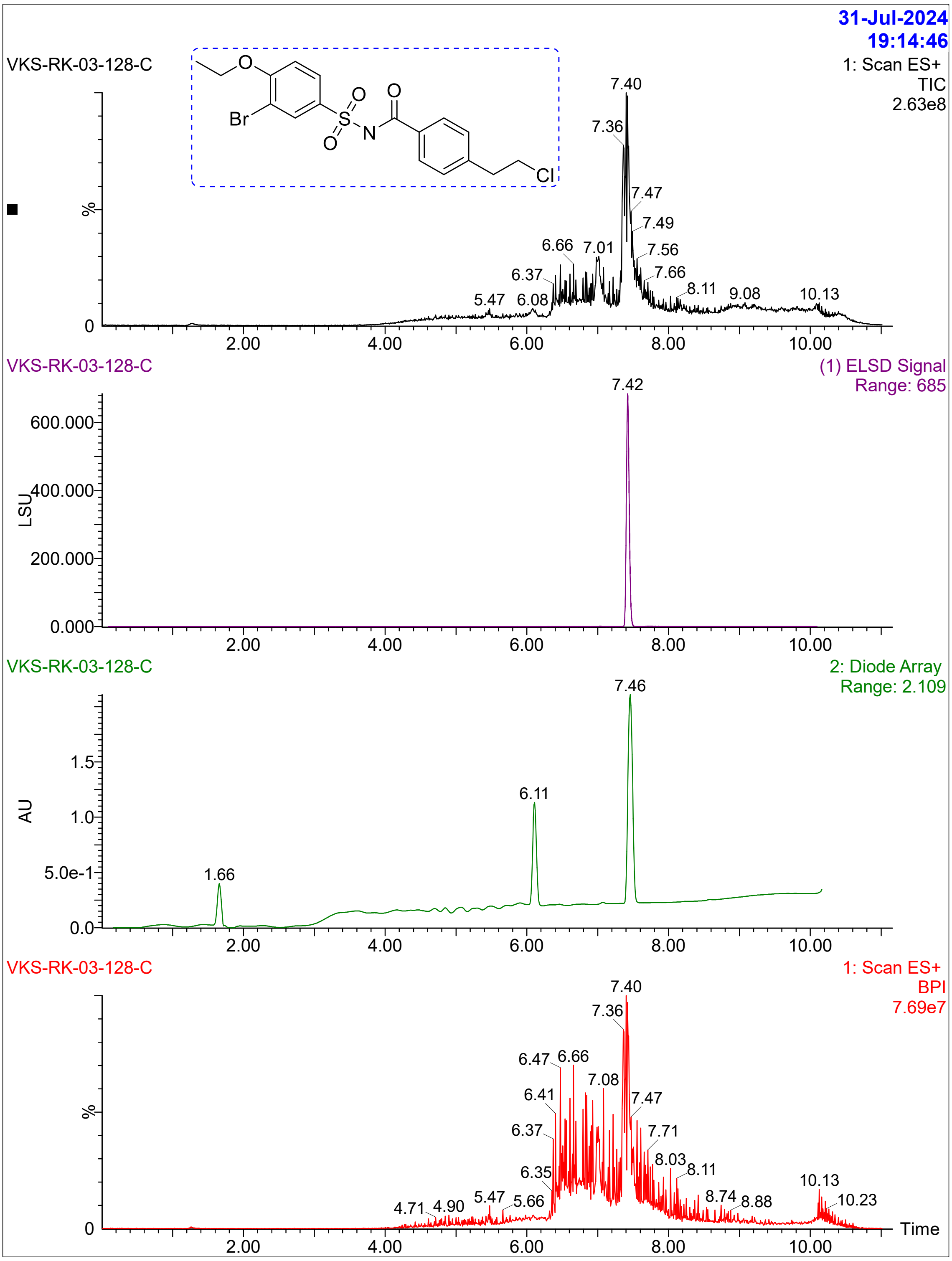

31-Jul-2024
19:14:46
VKS-RK-03-128-C
1: Scan ES+
TIC
2.63e8
Br
O
S
N
Cl
VKS-RK-03-128-C
(1) ELSD Signal
Range: 685
LSU
VKS-RK-03-128-C
2: Diode Array
Range: 2.109
AU
VKS-RK-03-128-C
1: Scan ES+
BPI
7.69e7
%
Time

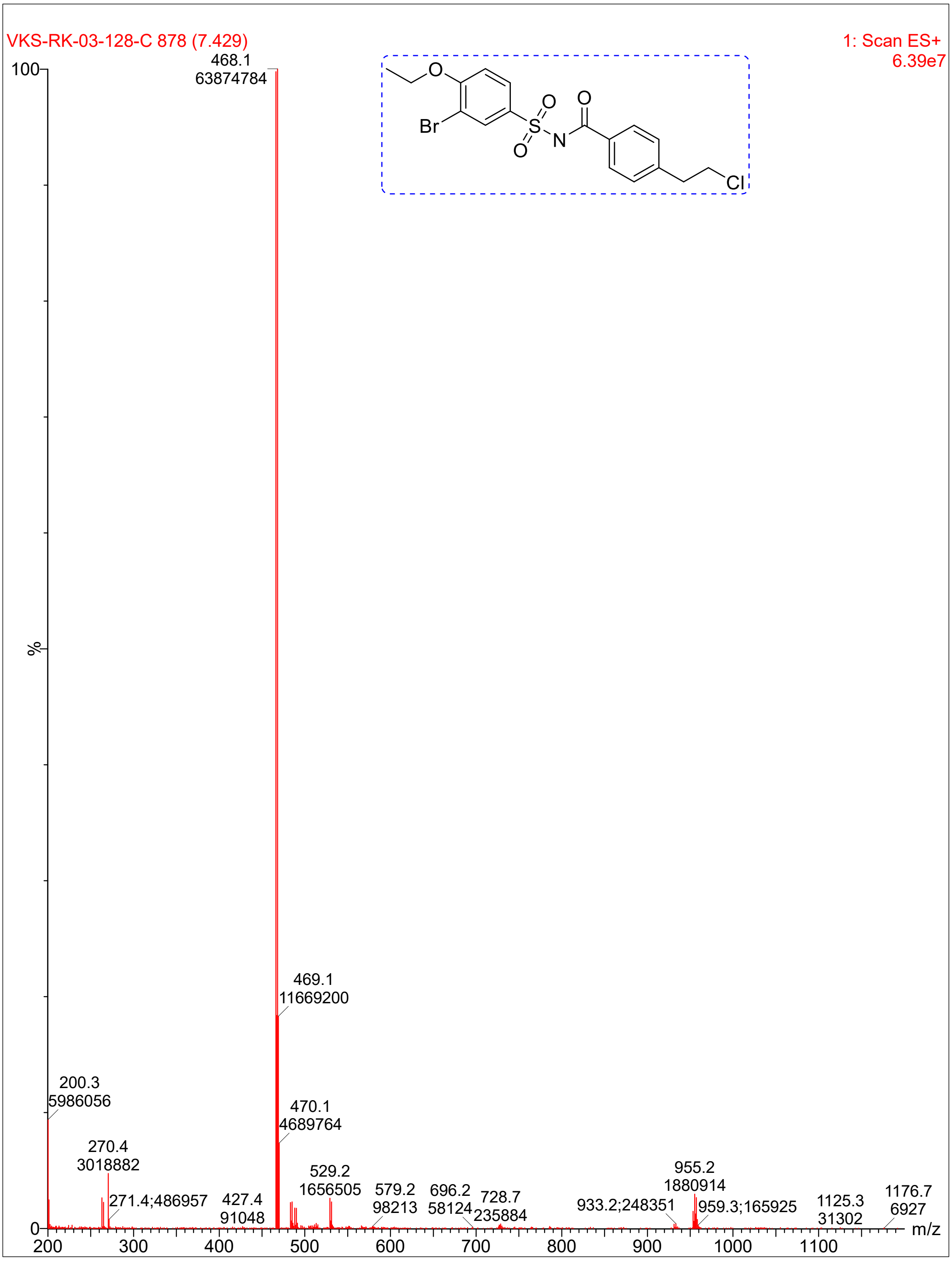

VKS-RK-03-128-C 878 (7.429)
1: Scan ES+
6.39e7
468.1
63874784
469.1
11669200
470.1
4689764
200.3
5986056
270.4
3018882
271.4;486957
427.4
91048
529.2
1656505
579.2
98213
696.2
58124
728.7
235884
933.2;248351
955.2
1880914
959.3;165925
1125.3
31302
1176.7
6927
%
m/z
O
Br
S
O
O
N
O
Cl

**Compound 41**:

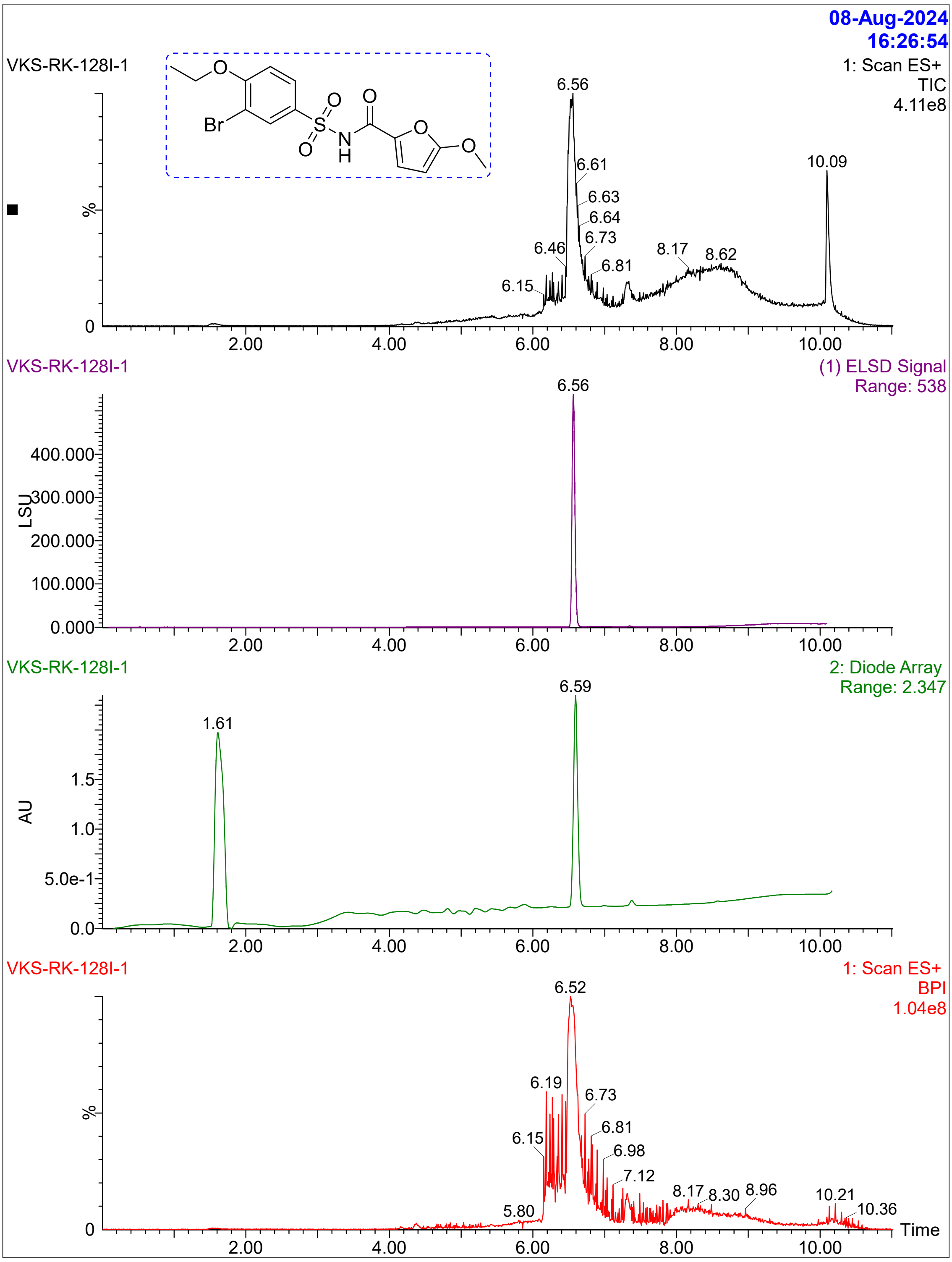

08-Aug-2024
16:26:54
VKS-RK-128I-1
1: Scan ES+
TIC
4.11e8
Br
O
S
N
H
6.56
6.61
6.63
6.64
6.73
6.81
6.46
6.15
8.17
8.62
10.09
%
0
2.00
4.00
6.00
8.00
10.00
VKS-RK-128I-1
(1) ELSD Signal
Range: 538
6.56
LSU
400.000
300.000
200.000
100.000
0.000
VKS-RK-128I-1
2: Diode Array
Range: 2.347
1.61
6.59
AU
1.5
1.0
5.0e-1
0.0
VKS-RK-128I-1
1: Scan ES+
BPI
1.04e8
6.52
6.19
6.15
6.73
6.81
6.98
7.12
5.80
8.17
8.30
8.96
10.21
10.36
Time

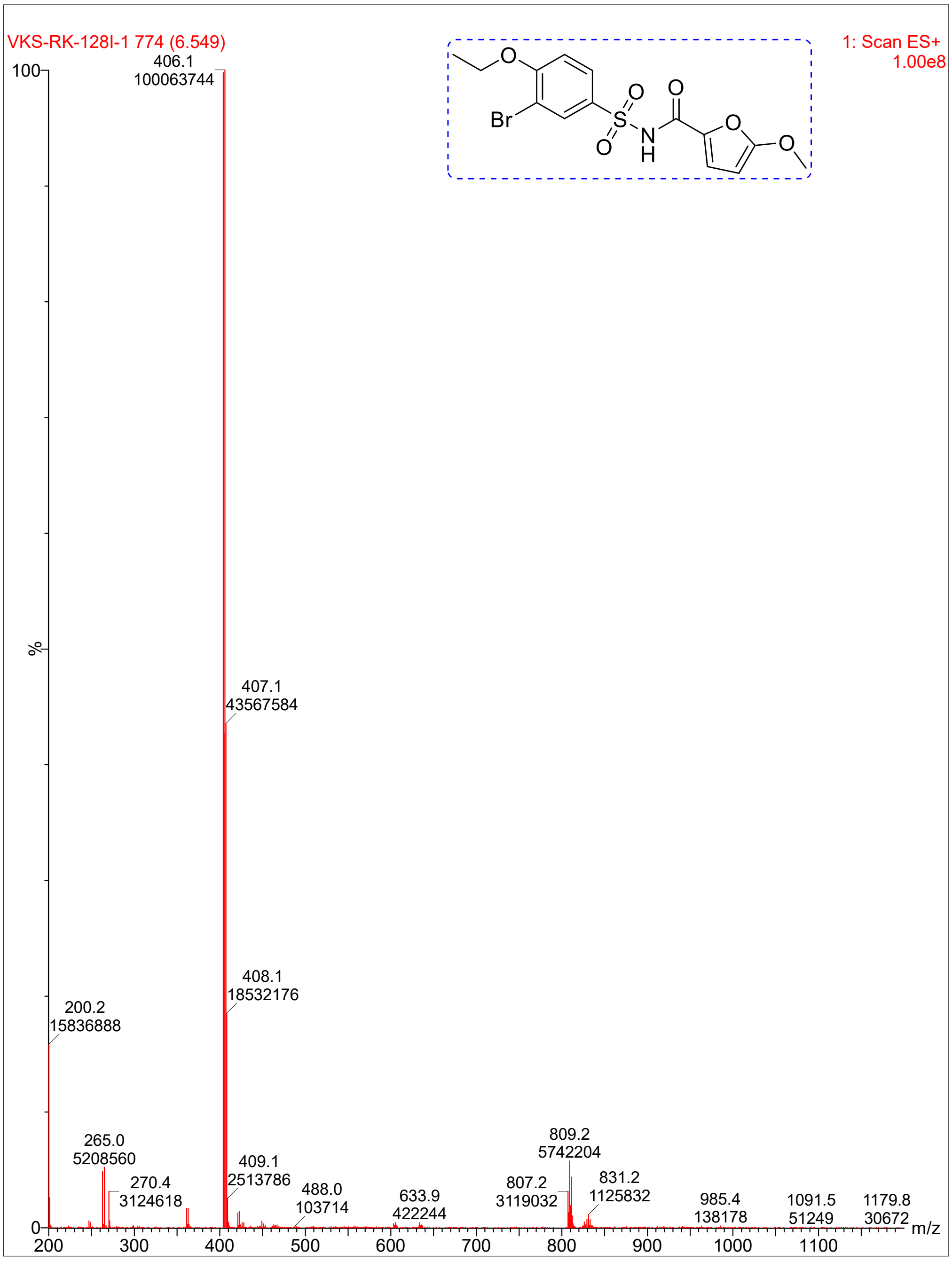


Reference:


(1) Lama, L.; Adura, C.; Xie, W.; Tomita, D.; Kamei, T.; Kuryavyi, V.; Gogakos, T.; Steinberg, J. I.; Miller, M.; Ramos-Espiritu, L.; et al. Development of human cGAS-specific small-molecule inhibitors for repression of dsDNA-triggered interferon expression. *Nature Communications* **2019**, *10* (1), 2261. DOI: 10.1038/s41467-019-08620-4.
